# A genetic history of the pre-contact Caribbean

**DOI:** 10.1101/2020.06.01.126730

**Authors:** Daniel M. Fernandes, Kendra A. Sirak, Harald Ringbauer, Jakob Sedig, Nadin Rohland, Olivia Cheronet, Matthew Mah, Swapan Mallick, Iñigo Olalde, Brendan J. Culleton, Nicole Adamski, Rebecca Bernardos, Guillermo Bravo, Nasreen Broomandkhoshbacht, Kimberly Callan, Francesca Candilio, Lea Demetz, Kellie Sara Duffett Carlson, Laurie Eccles, Suzanne Freilich, Ann Marie Lawson, Kirsten Mandl, Fabio Marzaioli, Jonas Oppenheimer, Kadir T. Özdogan, Constanze Schattke, Ryan Schmidt, Kristin Stewardson, Filippo Terrasi, Fatma Zalzala, Carlos Arredondo Antúnez, Ercilio Vento Canosa, Roger Colten, Andrea Cucina, Francesco Genchi, Claudia Kraan, Francesco La Pastina, Michaela Lucci, Marcio Veloz Maggiolo, Beatriz Marcheco-Teruel, Clenis Tavarez Maria, Cristian Martinez, Ingeborg París, Michael Pateman, Tanya Simms, Carlos Garcia Sivoli, Miguel Vilar, Douglas J. Kennett, William F. Keegan, Alfredo Coppa, Mark Lipson, Ron Pinhasi, David Reich

## Abstract

Humans settled the Caribbean ~6,000 years ago, with intensified agriculture and ceramic use marking a shift from the Archaic Age to the Ceramic Age ~2,500 years ago. To shed new light on the history of Caribbean people, we report genome-wide data from 184 individuals predating European contact from The Bahamas, Cuba, Hispaniola, Puerto Rico, Curaçao, and northwestern Venezuela. A largely homogeneous ceramic-using population most likely originating in northeastern South America and related to present-day Arawak-speaking groups moved throughout the Caribbean at least 1,800 years ago, spreading ancestry that is still detected in parts of the region today. These people eventually almost entirely replaced Archaic-related lineages in Hispaniola but not in northwestern Cuba, where unadmixed Archaic-related ancestry persisted into the last millennium. We document high mobility and inter-island connectivity throughout the Ceramic Age as reflected in relatives buried ~75 kilometers apart in Hispaniola and low genetic differentiation across many Caribbean islands, albeit with subtle population structure distinguishing the Bahamian islands we studied from the rest of the Caribbean and from each other, and long-term population continuity in southeastern coastal Hispaniola differentiating this region from the rest of the island. Ceramic-associated people avoided close kin unions despite limited mate pools reflecting low effective population sizes (2N_e_=1000-2000) even at sites on the large Caribbean islands. While census population sizes can be an order of magnitude larger than effective population sizes, pan-Caribbean population size estimates of hundreds of thousands are likely too large. Transitions in pottery styles show no evidence of being driven by waves of migration of new people from mainland South America; instead, they more likely reflect the spread of ideas and people within an interconnected Caribbean world.

---

Prior to European colonization, the Caribbean islands were a mosaic of archaeologically-distinct cultures reflecting intricate networks of interaction and multiple instances of cultural change since the first human occupation ~6,000 years ago. There is debate about the extent to which these changes reflect local developments and intra-Caribbean interactions or migrations from the American continents^1,2^, and the connections between demographic movements and three distinct archaeological periods, classified as the Lithic, Archaic, and Ceramic Ages remain unclear^3^ (Supplementary Information section 1).

The Lithic Age represents the earliest archaeological evidence of human occupation in the Caribbean (~6,000 years ago) and is found in Cuba, Hispaniola, and Puerto Rico. The Archaic Age is defined by the appearance of ground-stone artifacts as early as 5,000 years ago, and has been hypothesized to reflect a second spread of technology and possibly people from South America^4,5^. The Ceramic Age, beginning 2,500-2,300 years ago, is characterized by an agricultural economy and intensive pottery production; it is widely accepted as reflecting at least one migration of people from the mouth of the Orinoco River in Venezuela and the Guianas who spoke a language related to present-day Arawak languages. There are debates about the extent to which distinct ceramic styles (‘series’) correlate to the movements of people during the Ceramic Age^1–3^ (Supplementary Information section 1), as well as about pan-Caribbean population size during this time^6–8^.

Ancient DNA data from the Caribbean to date comes from mitochondrial DNA^9–13^ as well as genome-wide data from a 1,000 year-old individual from The Bahamas^14^ and two ancient individuals from Puerto Rico^13^. We screened 208 individuals for evidence of authentic ancient DNA (Supplementary Data 1) and generated genome-wide data passing standard criteria for authenticity for 184 unique ancient individuals who lived between ~3150-300 calibrated years (cal. yr) before the present (BP, taken as 1950 CE in accordance with radiocarbon calibration convention) in the Bahamas, Cuba, Hispaniola (which we separate into Haiti and the Dominican Republic in our analyses using present-day borders for higher geographic resolution), Puerto Rico, Curaçao, and northwestern Venezuela (Fig. 1; Supplementary Information section 2). We leverage these data and 52 new direct radiocarbon dates (Supplementary Information section 3; Supplementary Data 3) to address key debates about population history in the pre-contact Caribbean. We summarize these debates and how they are advanced by new genetic data in Table 1. In what follows, we use ‘Archaic’ to denote sites with an abundance of stone tools and/or site dates predating the spread of intensive ceramic use, and ‘Ceramic’ to denote sites with a preponderance of ceramics.

**Fig. 1:**
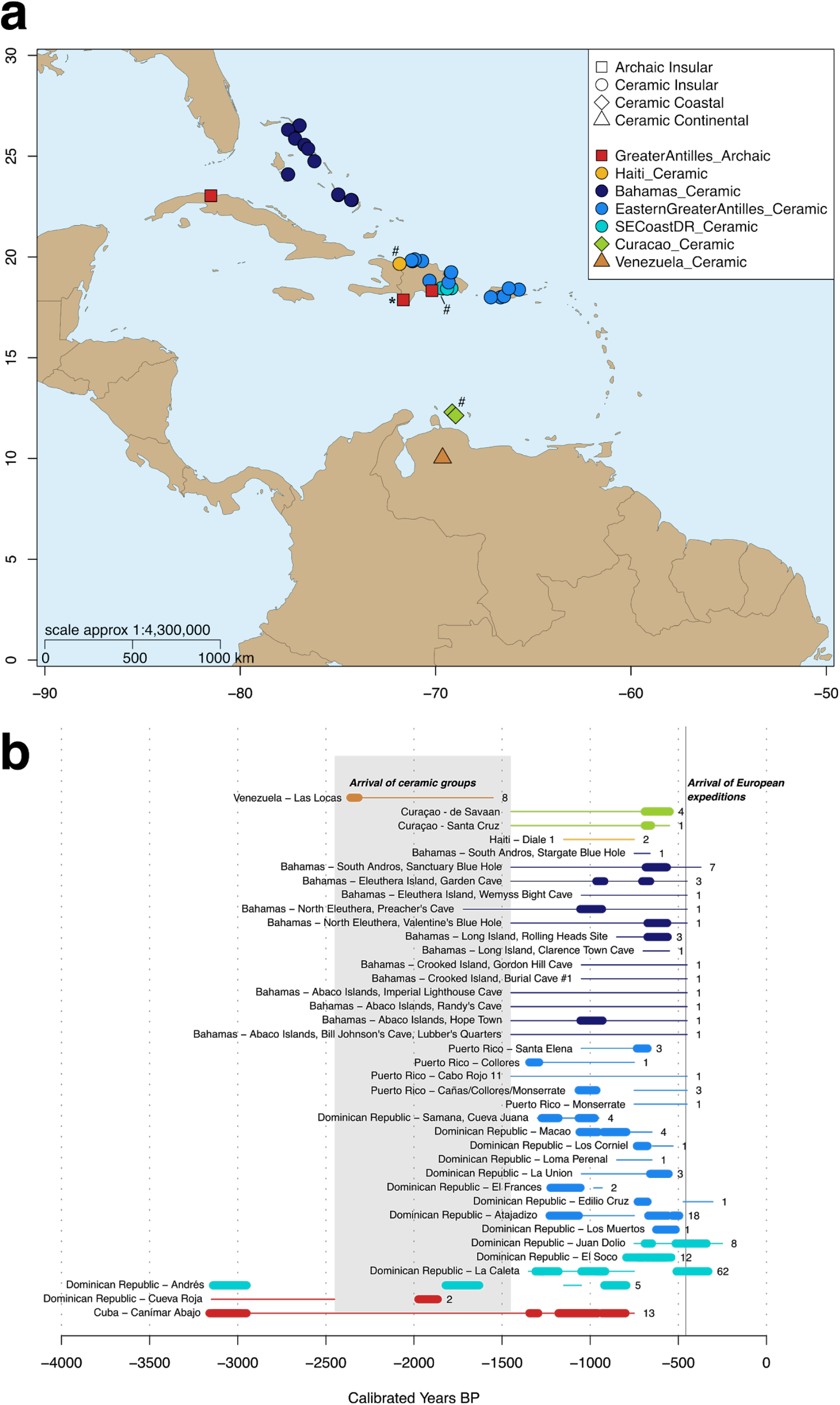
Analysed sites. **(a)** Geographical distribution. Shapes represent genetic clusters to which we assigned each sample; asterisk (*) denotes individuals from the Archaic site of Cueva Roja (Dominican Republic) excluded from main analyses due to low-coverage; hash (#) denotes presence of individuals who harbored mixtures of the main ancestry clusters. We separate sites in Hispaniola using the present-day border between Haiti and the Dominican Republic. (**b**) Temporal distribution. Numbers represent individuals from each site; thick lines denote direct C14 dates (95.4% calibrated confidence intervals); thin lines denote archaeological context dating; grey area identifies the first arrivals of ceramic-users in the Caribbean. Colors and labels are consistent with Fig. 2c.

**Table 1.** Archaeological debates addressed by genetic data generated in the present study.

| Key debates from archaeology | New genetic data exploring this debate |
| --- | --- |
| Geographic origin, timing, trajectory of Archaic Age migration into the Caribbean; number of migrations during the Archaic Age | Genome-wide data from 16 Archaic-associated individuals from Cuba and the Dominican Republic suggest that there was a homogenous Archaic ancestry profile throughout the Greater Antilles. This ancestry persisted with minimal admixture in some areas for over 2,000 years. The Archaic Age migration originated within a population that is not closely related to any present-day genotyped Indigenous groups, though we find a North American origin for this migration unlikely. We find no evidence of multiple migrations during the Archaic Age. |
| Geographic origin, timing, trajectory of Ceramic Age migration into the Caribbean; number of migrations during the Ceramic Age | Genome-wide data from 153 individuals across The Bahamas and Greater Antilles spanning 1,400 years point to a single wave of migration into the Caribbean in the Early Ceramic Age at least 1,800 years ago with a likely connection to the geographic region where Arawak-speaking peoples live today (Venezuela and the Guianas). This accords with the presence of Arawak languages in the pre-contact Caribbean. A caveat is that multiple waves of migration from source populations that were highly genetically similar (such as may have been widespread throughout northern South America in the past even though such populations are not widespread there today) could go undetected with ancient DNA. |
| The relationship between transitions of material culture styles ('series') and the movement of people during the Ceramic Age | We find a signal of genetic homogeneity during the Ceramic Age that is independent of ceramic typology. This increases the weight of evidence that changes in pottery styles throughout the Ceramic Age were not associated with multiple waves of migration of genetically-distinct peoples from the American continents, and instead suggests that cultural diffusion or movement of genetically homogenous people within the Caribbean drove stylistic change. We detect relatives ~75km apart on Hispaniola, providing direct evidence of mobility. Geography explains the subtle population structure that we do observe; for example, individuals from The Bahamas form a group relative to ceramic users from the Greater Antilles, and people from four sites on the southeast coast of Hispaniola form a group relative to people from other parts of the island. |
| Nature of interactions between Archaic-associated peoples of the Caribbean and ceramic users who moved into this region | We find instances of admixture between Archaic- and Ceramic-associated people in some places, and the parallel existence of Archaic- and Ceramic-associated populations with little if any admixture in others. Admixture occurred at the Ceramic-associated site of Diale 1 (Haiti), where two individuals had ~19% Archaic-related ancestry. An individual from La Caleta (Dominican Republic) had significantly more Archaic-related ancestry (12%) than other Ceramic Age people who lived in this part of Hispaniola. In contrast, unadmixed Archaic-related ancestry persisted for ~2,000 years at Canimar Abajo (Cuba). |
| Genetic distinction between people living in the early or late part of the Ceramic Age | Our data do not support genetic differentiation across time during the Ceramic Age. Instead, we identify long-term genetic continuity at sites such as Andrés (Dominican Republic), where a homogenous ancestry profile persists for almost 1,000 years. |
| Presence of pre-contact ancestry in modern Caribbean people | Our data reveal Ceramic-related, but not Archaic-related, ancestry in present-day peoples from Puerto Rico and parts of Cuba in amounts up to ~14%. We also identify a previously undocumented mtDNA haplogroup - a deep branch of C1d - that is unique to Caribbean people and is identified in a single individual from Puerto Rico from the 1000 Genomes Project, providing additional direct evidence of Indigenous ancestry in the Caribbean today. |
| Population size and social organization in the Caribbean prior to European colonization | We estimate that effective population sizes ranged around $2N_e=1000-2000$ people in the pre-contact Caribbean. Census population size is typically not more than ten times larger, suggesting that previous inferences of population sizes in the hundreds of thousands are overestimated. While Caribbean people avoided unions of first cousins or closer, they had a limited pool of potential mates, indicated by many unions as close as second or third cousins. |
| Origins of the people who colonized The Bahamas | Genome-wide data from 24 Ceramic Age individuals from five islands of The Bahamas show most genetic affinity to ceramic users from the Greater Antilles, providing evidence against their origin in Archaic populations from Cuba. We show that neither individuals from an Archaic-associated site in the western part of Cuba (Canimar Abajo), nor an Archaic Age individual from the Dominican Republic (Andrés) have genetic affinity to the ancient people of the Bahamas that we analyze. |
| Possible migration of Carib peoples around 800 CE | Our data point to genetic continuity throughout the Ceramic Age and provide no definitive genetic evidence of a Carib migration during this time, although we cannot rule out admixture proportions up to ~8%. Additional caveats are that *Venezuela_Ceramic* and *Arara* may be imprecise proxies for ancient Carib-related ancestry, that the limited number of ancient individuals studied here are not fully representative of the entire Ceramic Age Caribbean (where some places not studied here could have Carib-related ancestry), and that a widely-shared genetic profile throughout northern South America (possibly including the Caribs) at the time of these posited migrations makes them difficult to identify using ancient DNA data. |

## Ethics

We acknowledge the ancient individuals whose skeletal remains we analyzed, present-day people who identify as having an Indigenous legacy, the community representatives across the Caribbean who provided critical feedback and offered local perspectives, and the Caribbean-based scholars who were centrally involved as collaborators and co-authors. Permissions to perform ancient DNA analysis of the human skeletal remains in this study were documented through authorization letters signed by a custodian who assumed responsibility for the skeletal remains collected from a specific geographic region or site. The conclusions drawn by this project are intended to provide new information about the genetic ancestry of the people of the Caribbean prior to European colonization in the late 15^th^ century and the subsequent African slave trade, and were discussed prior to publication with members of Indigenous communities who trace their legacy to the pre-contact Caribbean; feedback from this discussion was then incorporated into our manuscript. While genetic data are one form of knowledge that contributes to understanding the past, oral traditions and other types of Indigenous knowledge can coexist with scientific data. Genetic ancestry should not be conflated with perceptions of identity, which cannot be defined by genetics alone. A full ethics statement is provided in Supplementary Information section 15.

## Genetic-based clustering

We performed principal component analysis (PCA), computing axes using present-day North, Central, and South American groups with no evidence of European or African ancestry^15^ (Fig. 2a; Supplementary Data 4; Supplementary Information section 4). Ceramic-associated individuals project in a cluster separate from Archaic-associated individuals, a distinction confirmed by clustering analysis using ADMIXTURE (Fig. 2b; Supplementary Information section 5). Archaic-associated individuals and ancient Venezuelans somewhat overlap on the PCA and are characterized by two primary components in the ADMIXTURE analysis, one maximized in modern-day Chibchan-speaking Cabécar and the other in Ceramic-associated Caribbean individuals. One exception to the genetic homogeneity observed within the majority of sites is at Andrés (Dominican Republic), where most individuals fall within the ceramic cluster (consistent with the primarily ceramic association of this site), but a single individual (I10126) is dated to the Archaic Age (3140-2950 cal. yr BP) and clusters with other Archaic-associated individuals in PCA. Individuals from Curaçao and the Diale 1 site (Haiti) are distinct from either of the two main clusters. We exclude one individual from each of the four pairs of first-degree relatives (Supplementary Information section 6) and the two Archaic-associated individuals from Cueva Roja (Dominican Republic) with low coverage (<~0.02X, or ~20,000 SNPs) from subsequent statistical analyses (Supplementary Data 1).

**Fig. 2:**
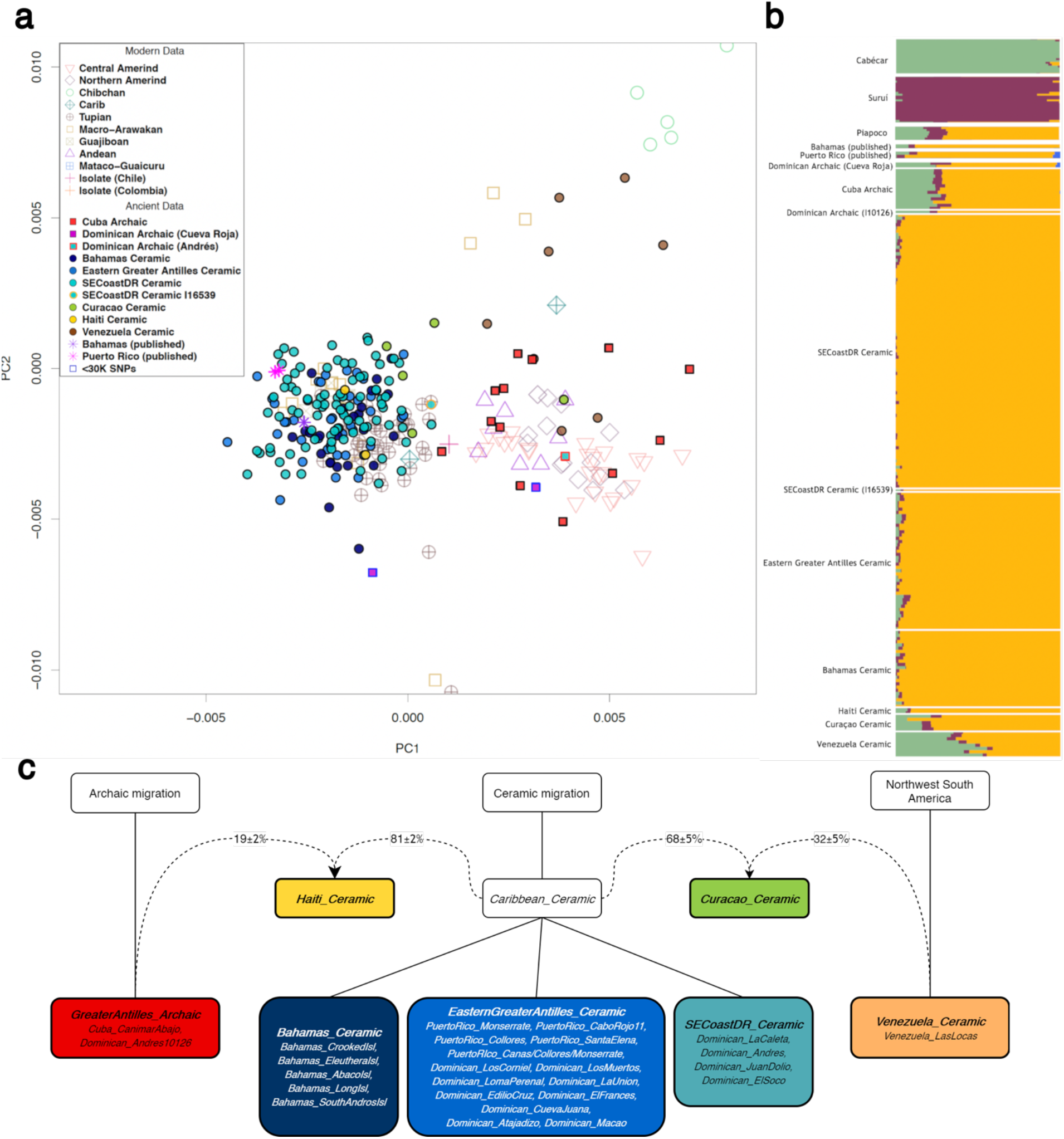
Overview of population structure. **(a)** PCA of ancient individuals projected onto present-day variation. Newly reported individuals are shown as solid symbols outlined in black, red, or blue (<30K SNPs). The plot here is zoomed for focus on ancient individuals and excludes some present-day populations; full plot provided in Supplementary Information section 4. **(b)** Visualization of model-based ancestry analysis using ADMIXTURE at five-fold cross-validation supported K=5 ancestral elements. Three modern-day populations are included for reference (Suruí, Cabécar, Piapoco); full results are provided in Supplementary Information section 5. **(c)** Relationships reconstructed with *qpWave*, Treemix, and *f*_*4*_-statistics (Supplementary Information section 7). Solid lines connect sub-groupings derived from a larger group; dashed lines represent ancestry contribution to admixed groups. Colored boxes represent the final sub-groupings.

To study genetic makeup independently from archaeologically-based material culture assignments (Supplementary Information section 2), we used a multi-step *qpWave*, Treemix, and *f_4_*-statistics-based workflow (Supplementary Information section 7). We sequentially grouped all individuals with increasing resolution based on their degree of allele sharing, starting with the identification of major groupings (‘clades’), and refining our understanding of the relationships between groups by assigning sub-groupings (‘sub-clades’) when they were supported. Clades and sub-clades were named following the formation of groups by combining the most specific geographic location encompassing all sites within that grouping along with ‘Archaic’ or ‘Ceramic’, based on chronology and/or the predominant artifacts at the site (Fig. 2c).

We identified three major clades that were significantly differentiated from each other by *qpWave*. *GreaterAntilles_Archaic* included all individuals from Canímar Abajo in northwestern Cuba spanning ~3150-800 cal. yr BP and individual I10126 from Andrés; two low coverage Archaic-associated individuals from Cueva Roja (~1900 cal. yr BP) also qualitatively fit into this clade (Fig. 2a, b). *Caribbean_Ceramic* comprised 153 individuals from The Bahamas, Dominican Republic, and Puerto Rico dating between ~1800-300 cal. yr BP. *Venezuela_Ceramic* comprised eight individuals from the site of Las Locas, dated to ~2350 cal. yr BP. Two remaining groups, *Haiti_Ceramic* and *Curacao_Ceramic*, were best modeled as having mixtures of ancestry related to the major clades (described below).

We next tested for finer population structure, identifying multi-site sub-clades within which individuals were significantly more closely related to each other on average than to individuals from other sites within the same clade (Supplementary Data 5). The sub-clade *SECoastDR_Ceramic* comprised four sites located along 50 kilometers of the southeast coast of the Dominican Republic (from west to east, La Caleta, Andrés, Juan Dolio, and El Soco) (Table S4). With radiocarbon dates spanning 1,500 years, this grouping documents a high degree of local and stable population structure over time and across changes in archaeological culture, indicating that shifts in ceramic style were not always associated with new spreads of people. All Bahamian sites (spanning ~600 years) separated into the sub-clade *Bahamas_Ceramic,* within which further substructure was detected specific to each of the five islands included in this study (Abaco Islands, Crooked Island, Eleuthera, Long Island, Andros Island), pointing to restricted gene flow among the islands; a possible signal of closer interactions is seen between the neighbouring Long and Crooked Islands (Table S3). The remaining sites from the *Caribbean_Ceramic* clade did not show any specific affinities and were merged to form *EasternGreaterAntilles_Ceramic*. We detected no between-site substructure within this sub-clade, except at Macao, where a signal of substructure is likely driven by the high proportion of related individuals analyzed. While we observe very limited differentiation in *Caribbean_Ceramic* spanning over a millennium, we use sub-clades to investigate the local history of ceramic users. We also tested all Ceramic-associated individuals individually for an excess of Archaic-related ancestry relative to others within their sub-clade using *f_4_*-statistics (Supplementary Information section 8; Supplementary Data 6) and identified individual I16539 from La Caleta (Dominican Republic) as showing significantly closer affinity to *GreaterAntilles_Archaic* than the rest of the *SECoastDR_Ceramic* sub-clade (Table S5); this individual was separated as *SECoastDR_Ceramic16539* for further analyses.

We called Y chromosome and mitochondrial (mtDNA) haplogroups (Supplementary Data 7; Supplementary Information section 9) and found that the majority of Y chromosome haplogroups belong to the Q-M3 lineage that is characteristic of Indigenous peoples of the Americas^16,17^, while all four major pan-American mtDNA haplogroups (A2, B2, C1, and D1^18,19^) are represented in our dataset. Consistent with previous work^13^, mtDNA haplogroup C1b2 was the most common haplogroup in *Caribbean_Ceramic*, and we find evidence of a previously undocumented deep branch of C1d present at a frequency of ~7% and in all sub-clades of *Caribbean_Ceramic* (Supplementary Information section 9). The discovery of this branch—which we do not detect in any ancient or modern individuals outside the Caribbean—indicates that ancestry of pre-contact people persists in the Caribbean today. It provides new evidence that Indigenous ancestry in the present-day Caribbean cannot simply be explained as imported from mainland groups after contact, as we find a single instance of this mtDNA haplogroup in a modern Puerto Rican individual in a screen of the 1000 Genomes Project dataset^20^ (Supplementary Information section 9).

## The early peoples of Cuba and the Dominican Republic

All Archaic-associated individuals from Canímar Abajo (Cuba) and I10126 form a single clade in *qpWave* despite spanning over 2,000 years. This points to a well-defined Archaic ancestry profile consistent with deriving from a single source spread across the Greater Antilles that persisted with minimal mixture in some regions well into the archaeologically-defined Ceramic Age. To investigate the affinities of *GreaterAntilles_Archaic* to populations from the American continents, we applied outgroup-*f*_*3*_ analysis (Supplementary Information section 10; Supplementary Data 8). *GreaterAntilles_Archaic* shares the most genetic drift with Indigenous groups from Central and northern South America (Fig. 3a) belonging to seven language families: Tupian, Macro-Arawakan, Cariban, Chibchan, Chocoan, Guajiboan, and Mataco-Guaicuru^21,22^. We find no evidence of excess allele sharing with people from any one family relative to the others (all |Z|<2.8, Fig. 3b; Supplementary Data 9).

**Fig. 3:**
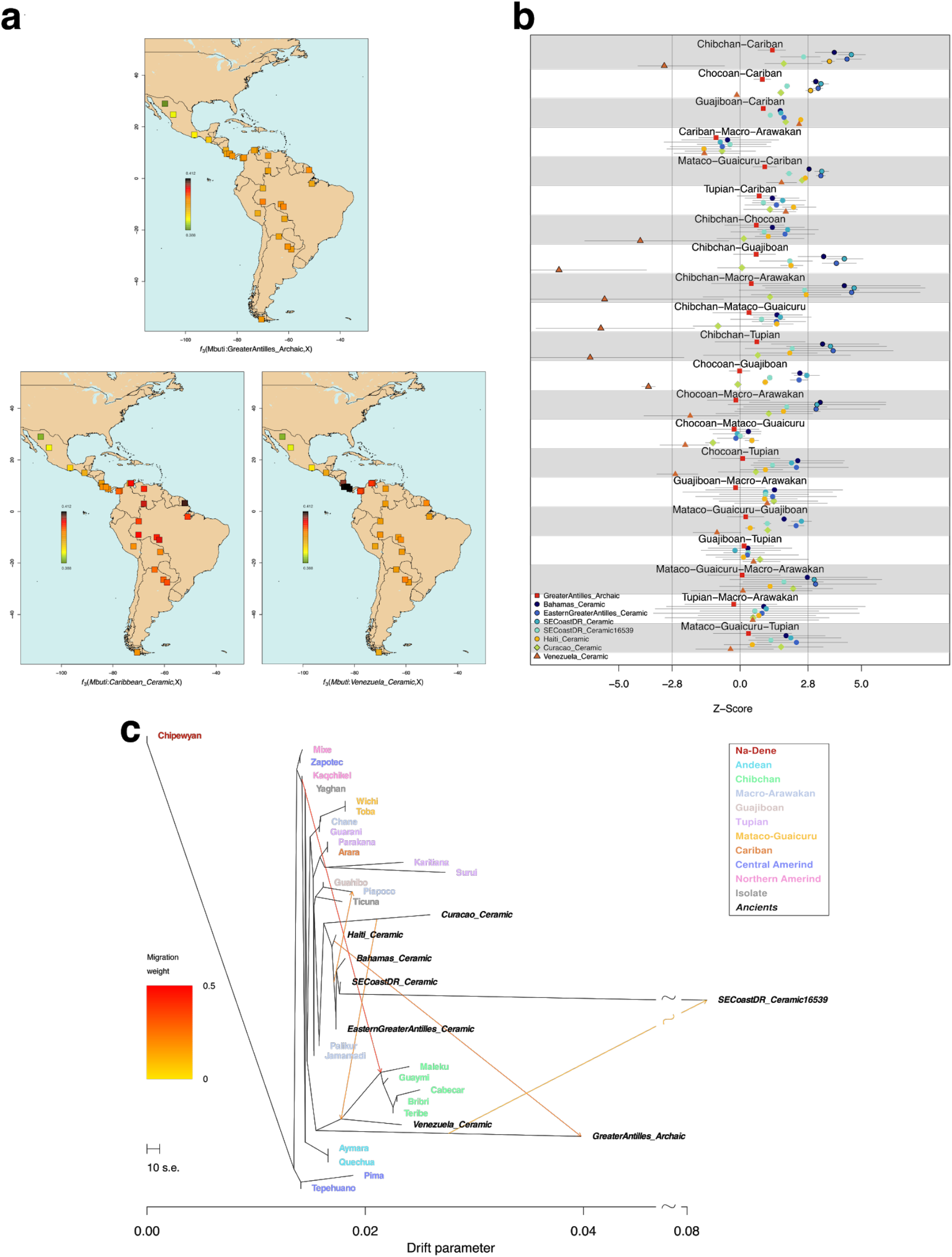
Affinities of the ancient populations from each region to modern Indigenous populations. (**a**) Outgroup *f*_*3*_-statistics performed on *GreaterAntilles_Archaic*, *Caribbean_Ceramic, and Venezuela_Ceramic*. Each square represents a modern population. (**b**) Symmetry tests using ancient sub-clades, analysing the average relatedness to non-pooled modern populations from seven language families found in South America. We show the average among symmetry tests of the form *f*_*4*_*(Mbuti, Test; LanguageGroup1Pop, LanguageGroup2Pop)*, where *LanguageGroupXPop* is a population from the corresponding language family; horizontal lines represent the range. Vertical lines represent statistical significance threshold of Z=±2.8 (corresponding to a 99.5% CI). (**c**) Maximum likelihood population tree from allele frequencies using Treemix showing the *Caribbean_Ceramic* sub-clades on the same branch as modern Arawak-speaking groups (Palikur, Jamamadi). Orange arrows represent admixture events, although the indicated direction of admixture might be inaccurate based on observations from analyses such as *qpAdm* admixture modeling - e.g., we believe it likely that there is *GreaterAntilles_Archaic* admixture into *Haiti_Ceramic* rather than the reverse direction of flow (Supplementary Information section 8).

We used *qpAdm* to explore the most likely source(s) of ancestry for *GreaterAntilles_Archaic* using published ancient DNA data older than 2000 years. We found a single model using individuals from southern Brazil as a source that passed a threshold of p>0.05 (*Brazil_Laranjal_6700BP*, p=0.343) (Supplementary Information section 8; Table S11). However, statistics of the form *f*_*4*_(*Mbuti*, *GreaterAntilles_Archaic*, *Brazil_Laranjal_6700BP, Test)* with *Test* as six other possible sources did not confirm significant affinities between *GreaterAntilles_Archaic* and *Brazil_Laranjal_6700BP* (|Z|<1.451), suggesting that the fit of *Brazil_Larajal_6700BP* as a source may reflect limited power to reject the model (Table S12). Using present-day populations, a similar result of no specific affinities was found (Table S13). In *qpGraph*, we fit *GreaterAntilles_Archaic* in several positions along a skeleton admixture graph from Posth et al.^23^, again reflecting poor power to differentiate between a Central or South American origin (Supplementary Information section 11). Likewise, in a maximum likelihood phylogenetic tree based on allele frequency covariances^24^, *GreaterAntilles_Archaic* is inferred to split before all South and some Central American populations (Fig. 3c). Together, these results point to the origin of *GreaterAntilles_Archaic* in a deeply divergent Native American population that is not particularly closely related to any sampled present-day groups.

## The spread of ceramic users in the Caribbean

When projected onto the PCA (Fig. 2a), all *Caribbean_Ceramic* individuals (dating ~1800-300 cal. yr BP) fall in a single cluster, as expected if a genetically homogeneous population moved into and throughout the Caribbean from a single source. We confirm this with pairwise F_ST_ <0.01 between the *Caribbean_Ceramic* sub-clades (very subtle absolute genetic differentiation). Values of F_ST_>0.1 (comparable to present-day inter-continental genetic distances) support the substantial genetic differentiation between this clade and *GreaterAntilles_Archaic* (Extended Data Fig. 1a).

Previous DNA studies have pointed to Arawak-speaking South American groups as being most closely related to Caribbean Ceramic-associated people^10,11,14,25^ (Supplementary Information section 1), consistent with the presence of Arawak languages in the Caribbean at the time of European contact^2^. ADMIXTURE analysis suggests that individuals from each *Caribbean_Ceramic* sub-clade are almost entirely composed of a component found in the highest proportion in modern Arawak-speaking Piapoco and Palikur (Fig. 2b), an affinity that is also supported by the position of all *Caribbean_Ceramic* sub-clades on the same branch as Piapoco and Palikur in a maximum likelihood tree allowing for admixture events (Fig. 3c). Similar to previous work^13,14^, we are unable to identify a statistically significant signal of closer relatedness to Arawak-than to Cariban- or Tupian-speaking populations using *f*_*4*_-statistics (Fig. 3b; Supplementary Information section 10; Supplementary Data 9), although we obtained a successful 1-way model with the Arawak-speaking Piapoco in *qpAdm* (p=0.519 or p=0.249, depending on the dataset used for Piapoco; Tables S13 and S14). Thus, our analyses also suggest an Arawak connection and highlight a genetic affinity between the Ceramic Age Caribbean and northeastern South Americans.

Aside from subtle population structure (described above) likely shaped by the geographic separation among the islands and potentially also influenced by different (but sub-significant) levels of admixture with Archaic groups (though a near-replacement of Archaic-related ancestry eventually did occur in many regions), we find no evidence that substantial proportions of different ancestry profiles were introduced to the Greater Antilles and The Bahamas throughout the Ceramic Age, in contrast to analyses of skeletal morphology that suggest a migration of Carib peoples from Venezuela ~1,150 years ago^26^. Our simulations show that ~2-8% Carib- or *Venezuela_Ceramic*-related ancestry would be required for detection, so we cannot rule out contributions less than this proportion; this analysis also requires that our selected proxies accurately represent Carib-related ancestry (Supplementary Information section 12). Thus, if a major Carib expansion occurred into the Antilles around 800 CE, it cannot have derived large proportions of ancestry from groups related to our present-day Cariban-speaking proxy or to ancient *Venezuela_Ceramic*-related people, and instead must have been derived to a mainland population from the northern coast of South America with an ancestry profile similar to that of present-day Caribbean Islanders. We are not aware of any modern groups from northern South America with such an ancestry profile, but the present-day distribution of Indigenous groups could plausibly be quite different from that of a millennium ago.

We find that only a minimal amount of Archaic-related ancestry may have persisted in ceramic-using populations, identifying isolated signals of Archaic-related admixture in three individuals from two Ceramic-associated sites in Hispaniola. *SECoastDR_Ceramic16539* (Dominican Republic) has 12.1±2.0% Archaic-related ancestry (Table S6), resulting from admixture that we estimate using the DATES software occurred an average of 39±14 generations before the individual lived (Z=2.77; Supplementary Information section 13), while two individuals from Diale 1 (Haiti) have 18.6±2.1% Archaic-related ancestry, which we estimate to have resulted from admixture ~11±5 generations before the individuals lived (Z=2.24; mixture proportions in Tables S7 and S8). In contrast, all individuals from the site of Canímar Abajo (Cuba) retained unadmixed Archaic-related ancestry spanning over 2,000 years to the limits of our resolution, consistent with archaeological^27^ and historical^28^ accounts that this region was home to people with a different language and cultural traditions than more easterly parts of Cuba as late as the Contact Period.

We model five ancient individuals from Curaçao as having 68.1±5.1% *Caribbean_Ceramic*-related ancestry and 31.9±5.1% *Venezuela_Ceramic*-related ancestry (p=0.317; Tables S9 and S10), indicating that people related to those who moved into the Greater Antilles also reached Curaçao and left a genetic legacy in the individuals we analyzed (we were unable to estimate dates for this admixture event due to limited statistical power). The *Venezuela_Ceramic* clade is distinct from any coastal or island population studied here, with ADMIXTURE suggesting a major component maximized in present-day Chibchan-speaking Cabécar (Fig. 2b); significant affinities with Chibchan speakers are also seen in statistics of the form *f*_*4*_*(Mbuti, Test; LanguageGroup1Pop, LanguageGroup2Pop)* (Fig. 3a, b). We are also able to model *Venezuela_Ceramic* using *qpAdm* with a single Chibchan-related component of ancestry from Cabécar (Tables S13 and S14). Thus, despite the location of *Venezuela_Ceramic* in a hypothesized source region for the peopling of the Caribbean and the dates of the analyzed individuals near the beginning of the Caribbean Ceramic Age, our analysis increases the weight of evidence for the Ceramic expansion having more easterly origins in northern South America. These results point to a scenario in which Curaçao’s Ceramic Age population was derived from admixture of two groups: one that was related to the population that spread to the Antilles at the beginning of the Caribbean Ceramic Age, and the other associated with the Dabajuroid ceramic style that linked Venezuelan sites like Las Locas to Curaçao.

## Insights into social structure and demographic history

We assessed kinship between the ancient individuals and identified a total of 43 first-, second-, or third-degree relationships. The majority were within the site of La Caleta, where 34 out of 62 individuals studied had relatives in our dataset, although the proportion of related pairs was not significantly larger than that from other sites (32 related out of 1891 pairs tested, 1.69%, 95% CI 1.16%-2.38%, versus six related out of 302 within-site pairs tested in other Caribbean Ceramic sites, 1.99%, 95% CI 0.73%-4.25%; Fig. 4a; Supplementary Information section 6). We also identified relatives (all genetic males) buried ~75 kilometers apart in the southern Dominican Republic: father I17906 and son I17903 from Atajadizo were second and third-degree relatives of I15601 from La Caleta. This inter-regional kinship provides direct evidence of mobility and connectivity during the Ceramic Age.

**Fig. 4:**
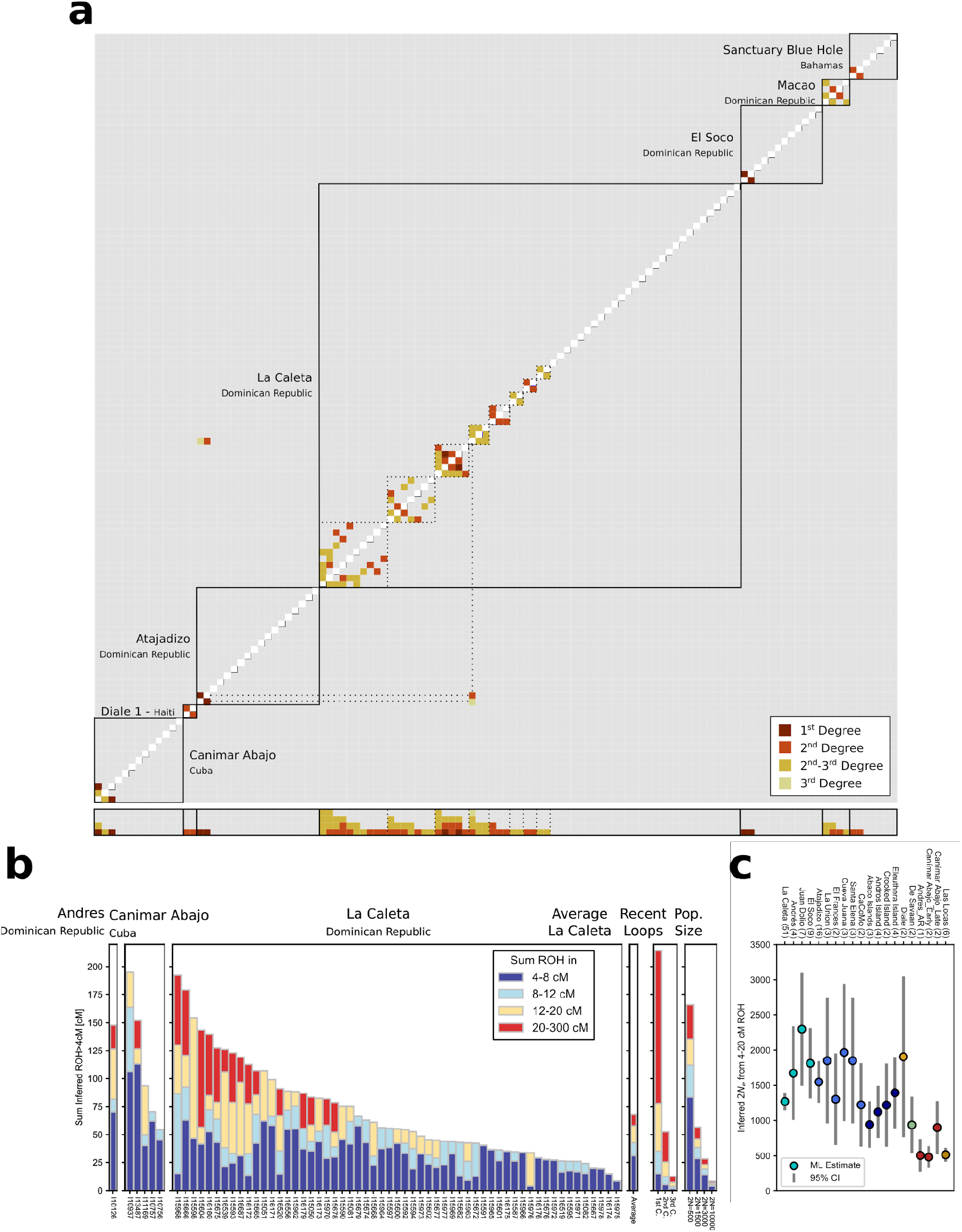
(**a**) Pairwise kinship estimates for all individuals from sites where relatives were identified. Dotted lines identify family clusters and inter-site relationships. Bottom rows correspond to relationships per individual. (**b**) Sums of inferred ROH larger than 4cM, per individual, for the Archaic-associated site of Canímar Abajo (Cuba) and the Ceramic-associated site of La Caleta (Dominican Republic). Remaining sites shown in Figure S17. “Recent loops” and “Pop. Size” represent expected distributions from simulated data for parental relationships as shown (C., cousin), and varying population sizes, respectively. (**c**) Inferred effective population sizes for individuals grouped per site (note that we here differentiate between an earlier and later occupation at Canímar Abajo). These estimates are derived from ROH blocks 4-20cM and a likelihood model (Supplementary Information section 6). Colors as per sub-clades, numbers in brackets denote sample size with sufficient coverage for ROH estimation).

We screened 154 individuals with at least 400,000 SNPs covered for runs of homozygosity (ROH) >4cM^29^ (Supplementary Information section 6; Supplementary Data 10; Figure S17). Large sums of long ROH blocks (> 20cM) indicate recent parental relatedness (Fig. 4b), whereas an abundance of shorter ROH is a signature of background parental relatedness, a consequence of restricted mating pools^30^. We identified only 2 of 154 individuals with at least 100cM of their genome in ROH>20cM blocks (~135cM is the average value for blocks of this size in offspring in first cousins, Fig. 4b, Figure S17). This shows that close kin unions were rare in the pre-contact Caribbean; instead, many unions took place between people as close as second and third cousins. However, we detected a general abundance of short and mid size ROH across our ancient samples (Fig. 4b). To quantify this signal, we applied a maximum likelihood method to estimate effective population size Ne using the length distribution of all ROH within 4-20cM, which arise from co-ancestry within the last few dozen generations (see Supplementary Information section 6 for details). We infer that population sizes for Ceramic Caribbean sites are larger (2N_e_ values ranging 1000-2000) than population sizes for Archaic individuals older than 2000 BP (2N_e_~500 for two sites) (Fig. 4c; Extended Data Table 1), pointing toward an increase of population density with the arrival of agricultural subsistence strategies, an observation in agreement with conditional heterozygosity levels (Extended Data Fig. 2). However, effective population size estimates across the Ceramic Caribbean remain low throughout the Ceramic Age, within the range of previous estimates made using a single ancient genome from The Bahamas^14^ (2N_e_=3200) and using coalescence rates of Caribbean mtDNA clades in Puerto Rico13 (pre-contact female 2N_e_<10,000). While short ROH measures population sizes of the past few dozen generations, the consistent signal across all Caribbean sites (including large and small islands) indicates that these estimates share some signal of larger meta-populations, as expected due to a high rate of mobility (and as confirmed by our detection of relative buried ~75 kilometers apart). For humans, census population sizes are typically up to an order of magnitude larger than effective population sizes^31^, with a factor of three estimated for inferences based on shared sequence blocks^32^. Even using the larger ratios, our results indicate that the population size estimates of hundreds of thousands on major Caribbean islands such as Hispaniola prior to European colonization that are discussed in the literature^6^ are substantial overestimates. At the same time, our results confirm the magnitude of population collapse following European colonization and the killing of Indigenous people. The severity of this collapse is evidenced, for example, in the frequencies of mitochondrial haplogroups such as B2 and D1 that were common in our data but are rare in present-day Caribbean populations^33^.

## Pre-Contact Ancestry Persists in Modern Caribbean Populations

We used genotype data from modern Caribbean populations^25^ to test for affinities between the Indigenous ancestry found in present-day Caribbean groups and the ancient Caribbean individuals. For three different populations (Cuba, Dominican Republic, and Puerto Rico) as *Test*, we measured affinity to the Archaic-associated individuals from Cuba versus Ceramic-associated individuals by computing the statistic *f*_*4*_(*European*, *Test*; *Cuba*_*Archaic*, *Caribbean_Ceramic*). Despite reduced statistical power due to relatively low proportions of Indigenous ancestry in present-day populations, we obtained a significant signal (Z=3.0) for Puerto Rico of greater relatedness to Ceramic-associated individuals, which we also replicated using Puerto Ricans from the 1000 Genomes Project (Z=4.1) (Supplementary Data 11). We performed an empirical power analysis (Supplementary Information section 14) for the present-day Cubans and determined that our results are consistent with entirely Ceramic-related ancestry but not with entirely Archaic-related ancestry. We also carried out the same test separately for present-day individuals from all 15 provinces of Cuba^34^ and found three provinces and five municipalities with weakly significantly Ceramic-related ancestry (2<|Z|<3.1) but none with significant evidence of Archaic-related ancestry (|Z|<2), including the region of northwestern Cuba that was the source of our ancient individuals (Supplementary Data 11). Thus, while our ancient DNA data from northwestern Cuba do not identify any Ceramic-related ancestry by ~900 cal. yr BP, such ancestry did eventually reach the island (Extended Data Tables 2 and 3). We also find support for the persistence of Indigenous ancestry in uniparental haplogroups, where major Indigenous-specific haplogroups identified in our ancient individuals are still found in the Caribbean today^33,35–37^, despite the introduction of European and African haplogroups over the last ~500 years. Detection of our newly discovered deep branch of haplogroup C1d (the only haplogroup in our dataset unique to the Caribbean) in a modern individual from Puerto Rico from the 1000 Genomes Project adds to the weight of this evidence.

## Discussion

Our analysis provides insight into debates surrounding the timing, trajectory, and geographic origin of demographic movements into the pre-contact Caribbean and the role of such movements in changes in technology and material culture that took place in this region over the past 6,000 years. First, we show that the ancestry in the Greater Antilles during the Archaic Age was consistent with deriving from a single continental source, though a limitation of our data is that the individuals we were able to analyze at high resolution were recovered from just two sites, albeit importantly coming from two different islands (Cuba and Hispaniola). While we cannot currently distinguish between models of a Central or South American origin, our analyses suggest that a North American origin for these Archaic-associated Caribbean peoples was unlikely.

Second, in line with evidence suggesting that the people who introduced intensive ceramic usage and agriculture into the Greater Antilles and The Bahamas moved from a source along the Orinoco River basin in Venezuela and the Guianas and arrived in the Caribbean Islands ~2,500 years ago^38^, our data are consistent with a movement from a single source accompanying the transition in material culture from predominant lithic-usage to a preponderance of ceramics. Ceramic-associated individuals from the parts of the Caribbean that we analyzed show an affinity to present-day Arawak speakers, consistent with an origin in northeastern South America as evidenced by archaeological and linguistic data^39^. Consistent with hypotheses that proto-Arawak-speakers experienced population splits at river junctions as they migrated from Amazonian South America toward the northeast part of South America, with some groups moving further along the Orinoco and into the Antilles and others moving toward the western Venezuela coast^35^, Ceramic-associated individuals from Curaçao have some ancestry related to that found in the *Caribbean_Ceramic* clade and additional ancestry such as is seen in the ceramic-users from Las Locas, suggesting that these populations came back together through admixture in the ancestry of the studied individuals in Curaçao.

Third, we find no evidence of differential ancestry during the Ceramic Age across changes in ceramic styles that have been hypothesized to reflect new migrations (though we present several caveats in Table 1). A recent study of facial morphology^26^ proposed a second expansion of Carib peoples to the Greater Antilles ~1,150 years ago, but we do not detect significant shifts in genetic ancestry throughout the entirety of the Ceramic Age, despite the fact that genome-wide analysis of hundreds of thousands of independent SNPs should in principle have more power to detect population substructure than morphological analysis, which at most can analyze only dozens of independent characters. More generally, we find no clear association between our genetic clusters within the Ceramic Age and the traditional ceramic stylistic typologies (Saladoid, Ostionoid, Meillacoid, Chicoid; Supplementary Information section 1), thereby providing no evidence for the culture-history model that considered stylistic transitions as the result of major movements of new people replacing established groups. Instead, we show that the ancestry profile of the southeastern coastal region of the Dominican Republic spans more than a millennium across major transitions in material cultural styles. While we cannot rule out scenarios of migrations of genetically very similar populations among the islands helping to drive some of the observed cultural changes, our findings increase the weight of evidence that exchange among established groups within the Caribbean was a key driver of stylistic changes.

Fourth, by assembling the first dataset from the Caribbean from both the Archaic and Ceramic Ages, we shed light on the interaction between representatives of both ancestry types. Our data from Canímar Abajo confirms that the direct descendants of people of the Archaic Age were contemporaries with ceramic-users, users consistent with the archaeological evidence of continuity of Archaic Age cultures in western Cuba. Furthermore, the presence of admixed individuals in Hispaniola (one from La Caleta and two from Diale 1), documents that a complete replacement of local Archaic-related ancestry did not immediately take place in all regions.

Fifth, we find that peoples living in some parts of the Caribbean (including Puerto Rico and certain areas of Cuba) today carry a signal of Ceramic-related ancestry despite centuries of admixture from European and African individuals. In Cuba, our results (combined with linguistic and physical anthropological evidence) suggest the possibility of the persistence of Archaic groups up until the Contact Period; however, the Indigenous admixed ancestry found in Cuba today is most likely not derived from this source. This could reflect post-colonial movement of Indigenous people between Caribbean islands or a more complex landscape of ancestry in Cuba prior to European colonization. Denser sampling of ancient individuals in the centuries surrounding colonization and from under-sampled locales such as Jamaica will provide insight on the events that contributed to the current distribution of Indigenous ancestry in Caribbean peoples.

Finally, our data provide new insights into social structure and demography. We identify instances of mobility, with related male individuals buried ~75 kilometers apart. We observe an active avoidance of unions between close relatives during both the Archaic and Ceramic Ages and large amounts of cumulative ROH across most of the Caribbean sites analysed compared to almost all ancient cultures studied to date with ancient DNA^29^. This reflects limited pools of mates due to small effective population sizes (1,000-2,000) averaged over the millennium prior to the time the individuals lived that are difficult to reconcile with theories that pre-contact population sizes in the Caribbean were extremely large. Even allowing for the fact that census sizes can be an order of magnitude larger than effective population sizes, our results point to a census size for the population of which Hispaniola was a part being no more than 10,000-20,000, which is far less than estimates of over 100,000 or even over 1,000,000 that have been discussed in the literature^3,40^. While our estimates are lower than some previous ones, the devastating impact that European colonization, expropriation, and systematic killing of Indigenous people had on Caribbean populations is clear. Today, modern Caribbean people are mixtures in different proportions of three groups, all of which have contributed importantly to present-day populations: Indigenous populations that experienced extreme post-colonial bottlenecks (~4% on average in Cuba, ~6% in Dominican Republic, and ~14% in Puerto Rico according to our estimation by *qpAdm*), immigrant Europeans (~70% in Cuba, 56% in the Dominican Republic, and 68% in Puerto Rico), and immigrant Africans who arrived in the course of the trans-Atlantic slave trade (~26% in Cuba, ~38% in the Dominican Republic, and ~18% in Puerto Rico) (Extended Data Tables 2 and 3). Altogether, these results reveal dynamic webs of biological and cultural connectivity in the ancient Caribbean, transformed over the last 500 years by demographic upheaval, but nevertheless persisting in many of the present-day people of this region.

## METHODS

No statistical methods were used to predetermine sample size. The experiments were not randomized, and the investigators were not blinded to allocation during experiments and outcome assessment.

### Ancient DNA analysis

We generated powder from the skeletal remains of all individuals. Powder was produced from a cochlea^41,42^, tooth, phalanx, or ossicle^43^ from each individual in a clean room facility at Harvard Medical School (Boston, USA), University College Dublin (Dublin, Ireland), or the University of Vienna (Vienna, Austria). See Supplementary Data 2 for the skeletal element used for each individual and location of powder preparation.

We extracted DNA in dedicated ancient DNA laboratories at Harvard Medical School or the University of Vienna following published protocols^44–46^. From the extracts, we prepared dual-barcoded double-stranded^47^ or dual-indexed single-stranded libraries^48,49^, both treated with uracil-DNA glycosylase (UDG) to reduce the rate of characteristic ancient DNA damage^50^. Double-stranded libraries were treated in a modified partial UDG preparation^47^ (‘half’), leaving a reduced damage signal at both ends (5’ C-to-T, 3’ G-to-A). Single-stranded libraries were treated with *E. coli* UDG (USER from NEB) that inefficiently cuts the 5’ Uracil and does not cut the 3’ Uracil. For a subset of samples, we increased coverage by preparing multiple libraries; see Supplementary Data 2 for the number of libraries analyzed for each individual.

To generate SNP capture data, we used in-solution target hybridization to enrich for sequences that overlap the mitochondrial genome and ~1.24 million genome-wide SNPs^51–54^ (“1240k”), either in two separate enrichments or simultaneously (Supplementary Data 2). We then added two 7-base-pair indexing barcodes to the adapters of each double-stranded library and sequenced libraries using either an Illumina NextSeq500 instrument with 2×76 cycles or an Illumina HiSeqX10 instrument with 2×101 cycles and reading the indices with 2× cycles (double-stranded libraries) or 2×8 cycles (single-stranded libraries).

Prior to alignment, we merged paired-end sequences, retaining reads that exhibited no more than one mismatch between the forward and reverse base if base quality was ≥20, or 3 mismatches if base quality was <20. A custom toolkit was used for merging and trimming adapters and barcodes (available at https://github.com/DReichLab/ADNA-Tools). Merged sequences were mapped to the reconstructed human mtDNA consensus sequence (RSRS)^55^ and the human reference genome version hg19 using the samse command in BWA v.0.7.15-r1140^56^ with the parameters −n 0.01, −o 2 and −l 16500. Duplicate molecules (those exhibiting the same mapped start and end position and same stand orientation) were removed after alignment using the Broad Institute’s Picard MarkDuplicates tool (available at http://broadinstitute.github.io/picard/). We trimmed two terminal bases from UDG-half libraries to reduce damage-induced errors.

We evaluated the authenticity of the isolated DNA by retaining individuals with a minimum of 3% of cytosine-to-thymine substitutions at the end of the sequenced fragments^47^, point estimates of mitochondrial DNA (mtDNA) contamination below 5% using contamMix v.1.0-12^51^, and point estimates of X chromosome contamination (in males) below 3%^57^ (Supplementary Data 2); three individuals (I7977, I7594, and I16540) were slightly below the set threshold for one authenticity metric, most likely due to low coverage (0.02-0.10X nuclear coverage and <23X mtDNA coverage), but passed the threshold for the other two metrics. SNPs were determined by randomly sampling an overlapping read with minimum mapping quality of ≥10 and base quality of ≥20. Individuals with less than 20,000 covered SNPs were excluded from quantitative analyses. One individual from each pair of first-degree relatives in the dataset was excluded from population genetics analysis; in all cases, we retained the higher coverage individual, listed in Supplementary Data 1.

### Radiocarbon dates

We report 52 new radiocarbon (^14^C) dates on bone fragments generated using accelerator mass spectrometry (AMS) (Supplementary Data 3). Most dates (n=48) were generated at the Pennsylvania State University (PSU) Radiocarbon Laboratory, and the remainder (n=4) were generated at the Center for Isotopic Research on Cultural and Environmental heritage (CIRCE). The sample preparation methodology at PSU was carried out as previously reported^58^, where bone collagen was extracted and purified using a modified Longin method with ultrafiltration^59^ (>30 kDa gelatin); if collagen yields were low, a modified XAD process^60^ (XAD amino acids) was used. Sample preparation at CIRCE was carried out following the lab-adapted Longin method^61^. Supplementary Data 3 lists the preparation method used for each sample.

We provide conventional radiocarbon dates in years BP and three calibrated dates for each conventional age in Supplementary Data 3. The first date is calibrated using the IntCal13 curve^62^; the second date is calibrated using a mix of the IntCal13 and Marine13^62^ curves in a 75%:25% ratio, accounting for a 25% marine protein contribution to the diet; and the third date is calibrated using a mix of the IntCal13 and Marine13 curves in a 50%:50% ratio, accounting for a 50% marine protein contribution to the diet. Supplementary Information section 3 discusses the impact of the marine reservoir effect on these dates. The differences between the median dates obtained for the three different calibration scenarios range from 190-360 years (median 265 years), and so we view this as a measure of the degree of systematic uncertainty in the ^14^C dates we obtained. We report all dates in the main manuscript as calibrated years before present, using the IntCal13 curve data for the calibrated dates.

### Dataset assembly

We merged the 184 ancient individuals that passed screening into a base dataset that included 61 previously published ancient American individuals^13,14,23,63–65^, and 36 modern Indigenous American groups sourced from single nucleotide polymorphism (SNP) array genotyping datasets or whole genome sequencing datasets (Extended Data Table 4):

- ‘1240K SNPs’, whole genome sequencing data restricted to a canonical set of 1,233,013 SNPs^51–54,66,67^
- ‘Human Origins dataset’, 597,573 SNPs^68–70^
- ‘Illumina dataset’ (unmasked/unadmixed individuals only), 352,432 SNPs^15^

All comparative analyses involving present-day Indigenous American populations were performed on the Illumina dataset, whereas for *qpAdm* and *qpWave*’s set of outgroup populations (“Right”) we used the Human Origins dataset for increased coverage. All genome-wide analyses were performed on autosomal data.

### Uniparental haplogroups

We determined mtDNA haplogroups using bam files, restricting to reads with MAPQ ≥ 30 and base quality ≥ 20. We constructed a consensus sequence with samtools and bcftools version 1.3.1 using a majority rule and then determined the haplogroup with HaploGrep2, using Phylotree version 17. We determined Y chromosome haplogroups using sequences mapping to 1240K Y-chromosome targets, restricting to sequences with MAPQ ≥ 30 and base quality ≥ 30. We called haplogroups by determining the most derived mutation for each sample, using the nomenclature of the International Society of Genetic Genealogy (ISOGG; http://www.isogg.org) version 14.76 (April 2019).

Mutational differences and corresponding mtDNA haplogroups, and Y chromosome haplogroups and their supporting derived mutations are found in Supplementary Data 7. Discussion of mtDNA and Y chromosome haplogroup distribution in the Caribbean is found in Supplementary Information section 9.

### Kinship, Consanguinity and Conditional Heterozygosity

We assessed kinship for every pair of individuals (including individuals from different sites and islands) using a previously described method^71^, and we present results for first-, second-, and third-degree relatives in Table S2. A number of the identified kin relationships are labeled as “2^nd^-3^rd^ degree”, due to uncertainty in the assessment of mismatch rates caused, for example, by increased homozygosity in the population overall.

In our dataset of 184 ancient individuals, we identified 49 individuals sharing 43 unique pairwise kin relationships up to the third degree. Four pairs of individuals were identified as first-degree relatives, while 16 pairs were definitively second-degree relatives, and 1 pair was probable third-degree relatives. The remaining 22 relationships were considered “2^nd^-3^rd^-degree.”

We identified Runs of Homozygosity (ROH) within our ancient dataset using the Python package *hapROH* (https://test.pypi.org/project/hapROH/). Following a previously described method^29^, we used 5008 global haplotypes from the 1000G haplotype panel^20^ as the reference panel. Following the recommendations for datasets with genotypes for ~1.24 million SNPs, we applied our method to ancient individuals with at least 400,000 SNPs covered (n=154) and ran the method on the pseudo-haploid data to identify ROH longer than 4 centiMorgans (cM). We used the default parameters of *hapROH*, which are optimized for ancient data genotyped at a similar number of sites. For each individual, we group the inferred ROH into four length categories: 4-8 cM, 8-12 cM, 12-20 cM and >20 cM and report the total sum in these bins (Supplementary Data 10; Figure S17).

To estimate effective population size from ROH, we applied a maximum likelihood inference framework (for derivation of the likelihood see Supplementary Information section 6). We fit all genome-wide ROH lengths between 4 and 20 cM long, and infer the effective population size that maximizes the likelihood for ROH lengths observed in a set of individuals. Estimation uncertainties are obtained from the curvature of the likelihood (Fisher Information matrix). Tests on simulated data confirmed the ability of the estimator to recover Ne estimates from genome-wide ROH of few individuals (Figures S18 and S19).

We used popstats^70^ to compute conditional heterozygosity for all clades and sub-clades, which we compared with contemporaneous groups from continental South America, such as from the Peruvian Middle and Late Horizon periods^72^. As previously described^73,74^, we restricted the analysis to transversion SNPs ascertained in a Yoruba individual; see Extended Data Fig. 2.

### PCA

We performed principal component analysis (PCA) with smartpca^75^, using the option ‘lsqproject: YES’ to project ancient individuals onto the eigenvectors computed from modern individuals. The approach of projecting each ancient sample onto patterns of variation learned from modern samples enables us to use data from a large fraction of SNPs covered in each individual and therefore maximize the information about ancestry that would be lost in approaches that require restriction to a potentially smaller number of SNPs for which there is intersecting data across lower coverage ancient individuals. We used the option ‘newshrink: YES’ to remap the points for the samples used to generate the PCA onto the positions where they would be expected to fall if they had been projected, thereby allowing the projected and non-projected samples to be appropriately co-visualized. We projected three previously published ancient individuals^13,14^ and 184 new ancient individuals onto the first two principal components computed using 61 individuals from 23 present-day populations (Fig. 2a). See Supplementary Data 4 for all individuals included in PCA and values of PCs 1 and 2. For PCA by archaeological site, non-zoomed PCA, and PCA excluding CpG sites, see Figures S13-S15.

### Unsupervised analysis of population structure

We used the software ADMIXTURE^76,77^ to perform unsupervised structure analysis on a dataset comprised of SNPs that overlap between the 1240k and Illumina dataset and were pruned in PLINK1.9^78^ using --indep-pairwise 200 25 0.4. This left 269,091 SNPs for the analysis. We ran five random-seeded replicates for each K in the interval between 2 and 10 with cross-validation enabled (--cv flag) to identify the runs with the lowest cross-validation error (Table S1). For each value of K, the replicate with the lowest cross-validation error was plotted and the results were compared. We choose to present K=5 as Fig. 2b as we found that the model with five components had the lowest cross-validation error in four out of five replicates and differentiated the components in a useful way for visualization. Results for the other values of K are presented as Figure S16 in Supplementary Information section 5.

### Estimation of F_ST_ coefficients

To measure pairwise genetic differentiation between two groups of individuals, we estimated average pairwise F_ST_ and its standard error via block-jackknife over 1000 markers using the function “average_patterson_fst” from the package “scikit-allel” (version 1.2.1, DOI 10.5281/zenodo.3238280). We removed the individual with lower coverage of each pair of first degree relatives, as well as ancestry outliers (see main text). For each ancient individual we analyzed one allele per pseudo-haploid call; see Extended Data Fig. 1.

### Clade grouping framework with *qpWave*, Treemix and *f*_*4*_-statistics

We used a multi-step framework involving *qpWave*, Treemix, and *f*_*4*_-statistics to group sites and individuals, and considered this information together with admixture profiles and proportions from *qpAdm* to produce **Fig. 2c**(detailed methodology in Supplementary Information section 7). We started by using *qpWave* to identify major clades based on shared ancestry and then used Treemix and *f*_*4*_-statistics to investigate the existence of sub-clades. Once all sub-clades were identified, we used *f*_*4*_-statistics to investigate further substructure between sites within each clade. Geographic and chronological information such as island or cultural affiliation was not considered for these analyses, ensuring all clades and subclades were based solely on genetic information. We examined the association between genetic data and archaeological cultural complexes only after considering the genetic and archaeological information separately, following a previously published example^79^.

The software *qpWave*^15^ from ADMIXTOOLS^68^ estimates the minimum number of ancestry sources needed to form a group of test populations (“Left”), relative to a set of differentially related reference populations (“Right”). If the “Left” group contains two populations, *qpWave* will evaluate if they can be modelled as descending from the same sources, and hence will determine whether they form a clade. We used 12 present-day Indigenous American populations from the Human Origins dataset^69^ plus Yukpa from ^66^ representing different language groups and ancestries from the American continent as our “Right” reference population set:

Chipewyan, Zapotec, Mixe, Mixtec, Suruí, Cabécar, Piapoco, Karitiana, Yukpa, Quechua, Wayuu, Apalai, Arara

The argument ‘allsnps: NO’ was used. We ran two consecutive steps of *qpWave* analyses, starting with the identification of major groupings (step 1), or clades, and then reassessed the relationships between members within those clades by running the same tests in a “model competition” approach (step 2), such as is implemented in the related software *qpAdm*. A significance threshold of p>0.01 was set for accepting a clade between two sites or individuals.

After identifying the major clades and/or pairs of sites that uniquely formed a clade with one another, we ran Treemix with these clades and 27 previously published present-day Indigenous populations^15^ (Extended Data Table 4) to identify within-clade site structure (step 3) by generating a maximum likelihood tree. We excluded four Chibchan, Chocoan and Arawak-speaking populations possibly admixed with each other from this analysis. We ran Treemix, grouping the SNPs in windows of 500 (flag - k 500) to account for linkage disequilibrium, setting Chipewyan as root (-root), allowing random migration events (-m), and disabling sample size correction (-noss) in order to include sites or populations represented by a single-individual. By running Treemix and allowing consecutive random migration/admixture events, we identified the ancient Caribbean sites that consistently shared the same relationships. We then used *f*_*4*_-statistics to evaluate if they formed a sub-clade to the exclusion of the other sites by following the tree’s structure. For each identified intact node among all Treemix runs we used each downstream pair of site(s) as Test1 and Test2 and investigated their relationship to upstream sites or pools of sites (step 4). If an upstream node was unchanged in all runs, the sites composing it were pooled. However, once the first inconsistency was identified in an upstream node, all sites beyond that node were pooled together. A combination of three statistics per relationship allowed us to evaluate the Treemix structure of the sites being tested:

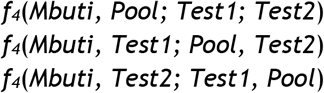

With Test1 and Test2 expected to be closer to each other than to Pool, the tested relationship finds support if the first test is statistically non-significant and at least one of the other two are significant. We used a Z-score threshold of 2.8 (associated with a 99.5% CI) to assess significance. These sites were then merged into a sub-clade inside the major Ceramic clade for further analysis.

After this clading analysis, we used *f*_*4*_-statistics to further investigate potential substructure between sites within each sub-clade (step 5). For each pairwise site comparison, we randomly divided each site into two groups of individuals, and used a statistic of the form *f*_*4*_(Site1_subset1, Site2_subset1; Site1_subset2, Site2_subset2) to identify positive statistics suggesting substructure within the same clade. This randomization step was repeated 10 times, and the average Z-score was calculated. If a site was composed of a single individual we instead computed statistics of the form *f*_*4*_(*Mbuti*, Site1_subset1; Site2_singleIndividual, Site1_subset2), intended to evaluate if individuals within Site1 were closer to each other than to the single individual from Site2. No statistics were computed if both sites being tested contained only one individual.

#### qpAdm

We used *qpAdm*^53^ from ADMIXTOOLS^68^ with ‘allsnps: NO’ to identify the most likely sources of ancestry and admixture for our populations/clades. First, we investigated if the outliers *SECoastDR_Ceramic16539* and *EasternGreaterAntilles_Ceramic7969,* as well as the individuals comprising the sub-clades *Haiti_Ceramic* and *Curacao_Ceramic*, could be modelled as admixed between the major ancestries represented by *GreaterAntilles_Archaic* (composed of all individuals from the Cuban site of Canímar Abajo and I10126), *Caribbean_Ceramic* (composed of *Bahamas_Ceramic*, *EasternGreaterAntilles_Ceramic* and *SECoastDR_Ceramic*), and *Venezuela_Ceramic.* We used this information to complete Fig. 2c. Then, based on this admixture information, we attempted to obtain more detailed admixture models using the sub-clades from within *Caribbean_Ceramic* and the sites from within *GreaterAntilles_Archaic* as possible sources. Lastly, we attempted to identify more distal sources of ancestry by using previously published ancient individuals from the Americas^23,63–65^, in this case for *qpWave*’s three major clades/groups. The base “Right” set used was the same used for *qpWave*. We also tested all 1-, 2-, and 3-way models using these “Right” populations as sources by moving them to the “Left” as necessary, and confirmed the results with the same unmasked/unadmixed populations from the Illumina dataset.

#### qpGraph

Due to the lack of significant *f*-statistics and attraction of *GreaterAntilles_Archaic* to any other modern or ancient sequenced population, we investigated where *GreaterAntilles_Archaic* would fit in the skeleton tree of previously published ancient American populations^23^ using *qpGraph*. Detailed methodology is provided in Supplementary Information section 11.

### Admixture simulations

We investigated the sensitivity of *qpWave* in detecting Carib-related ancestry in the *Caribbean_Ceramic* sub-clades by generating artificially admixed individuals with *Caribbean_Ceramic* ancestry mixed with increasing amounts of proxy Carib-related ancestry (1, 2, 5, 8, 10, 20, 30, 40, and 50%), and then assessing at what admixture threshold we were able to reliably detect the latter ancestry type (Supplementary Information section 12; Figure S28). To generate these admixed individuals, we identified common SNPs between the two sources, randomly selected genotypes from the Arara individuals from the Human Origins and Illumina SNP array datasets, corresponding to each of the nine percentages to be tested, and added the remaining SNPs from a random individual from *Bahamas_Ceramic*, *EasternGreaterAntilles_Ceramic* and *SECoastDR_Ceramic* with over 800,000 SNPs. We then ran *qpWave* with *Bahamas_Ceramic*, *EasternGreaterAntilles_Ceramic*, *SEoastDR_Ceramic* and each of the simulated admixed individuals on the “Left”, while using the default 13 “Right” populations, as described in Supplementary Information section 7, plus the Carib population (Arara) used to generate those individuals.

### Dating admixture

We used the method *DATES* (Distribution of Ancestry Tracts of Evolutionary Signals^58,80^) version 3520 to estimate the time of admixture in admixed individuals from Haiti and Curaçao as well as admixed individual I16539. This method measures the decay of ancestry covariance to infer the time since mixture and estimates jackknife standard errors. Details of *DATES* analysis is found in Supplementary Information section 13.

### Relatedness of ancient individuals to present-day admixed Caribbean populations

We computed relative allele-sharing between present-day admixed Caribbean populations (via their Indigenous ancestry) and ancient Archaic-associated versus Ceramic-associated individuals through the statistic *f*_*4*_(*European*, *Test*; *Cuba*_*Archaic*, *Caribbean_Ceramic*). In order to evaluate statistical power, we compared results for present-day Cubans alone to results obtained by adding one ancient individual from either the *GreaterAntilles_Archaic* or *Caribbean_Ceramic* clade to the Cuban test population. Full details found in Supplementary Information section 14.

### Testing for an Australasian link

We tested for a signal of relatedness to present-day Australasian populations^66,70^ (“Population Y” signal), using the statistic *f*_*4*_(*Mbuti*, *Onge/Papuan*; *Mixe*, *Archaic/Ceramic*). Here, Mixe is representative of a population that harbors no Population Y signal. When Onge was used as the Australasian proxy, several of our ancient groups showed weak positive statistics (Z > 2), but only the Archaic individual I10126 from the site of Andrés in the Dominican Republic (Z = 3.4) surpassed our threshold of Z > 2.8 (Extended Data Table 5). The signal was also weaker when Papuan was used as the Australasian proxy (Z < 2.6). The lack of a clear population Y signal is consistent with prior studies that also have not found this signal in ancient individuals from this region^14^ and other areas of South America^23^.

## Code availability

The custom code used in this study is available from https://github.com/DReichLab/ADNA-Tools.

## Data availability

The aligned sequences are available through the European Nucleotide Archive under accession number PRJEB38555. Genotype data used in analysis are available at https://reich.hms.harvard.edu/datasets. Any other relevant data are available from the corresponding authors upon reasonable request.

## Acknowledgements

We acknowledge the ancient people who were the source of the skeletal samples analyzed in this study as well as modern people from the Caribbean who have a genetic or cultural legacy from some of the ancient populations we analyzed. This work was supported by a grant from the National Geographic Society to Michael Pateman to facilitate analysis of skeletal material from The Bahamas. D.R. was funded by NSF HOMINID grant BCS-1032255, NIH (NIGMS) grant GM100233, the Paul Allen Foundation, the John Templeton Foundation grant 61220, and the Howard Hughes Medical Institute. We thank Juan Avilés, Juan Acayaguana Delvalle, Jorge Estevez, Dianne T. Golding Frankson, Jenna Gregory, Lynne A. Guitar, Lisa Kelly, Gerald Alexander Lopez Castellano, Kalaan Robert Nibonri, and Orlando Patterson for comments on an early version of this manuscript and discussions improving the presentation of this work; Vanessa A. Forbes-Pateman and Nancy Albury for their assistance compiling descriptions for archaeological sites in The Bahamas; Eadaoin Harney and Robert Maier for help with data processing; and Nick Patterson for advice on analysis. We dedicate this article to the memory of Fernando Luna Calderon, who would have been a co-author had he not passed away in the course of the work for this study.

## Author Contributions

W.F.K., A.Cop., M.L., R.P., and D.R. supervised the study. J.S., O.C., C.A.A., E.V.C., R.C., A.Cuc., F.G., C.K., F.L.P., M.L., M.V.M., C.T.M., C.M., I.P., M.P., T.S., C.G.S., and M.V. provided skeletal materials and/or assembled and interpreted archaeological and anthropological information. C.A.A., E.V.C., C.K., M.V.M., C.T.M., C.M., I.P., M.P., T.S., and C.G.S. contributed local perspectives to the interpretation and contextualization of new genetic data. B. M.-T. provided data from present-day populations. N.R., M.M., S.M., N.A., R.B., G.B., N.B., O.C., K.C., F.C., L.D., K.S.D.C., S.F., A.M.L., K.M., J.O., K.Ö., C.S., R.S., K.St., and F.Z. performed ancient DNA laboratory and/or data-processing work. B.J.C., L.E., F.M., F.T. and D.J.K. performed radiocarbon analysis. D.F., K.Si., H.R., M.M., S.M., I.O., and M.L. analysed genetic data. D.F., K.Si., W.F.K., and D.R. wrote the manuscript with input from all co-authors.

## Additional information

**Extended data** is available for this paper at xxxxxxxx.

**Supplementary information** is available for this paper at xxxxxxx.

**Reprints and permissions information** is available at xxxxxx.

---

**Extended Data Fig. 1.**
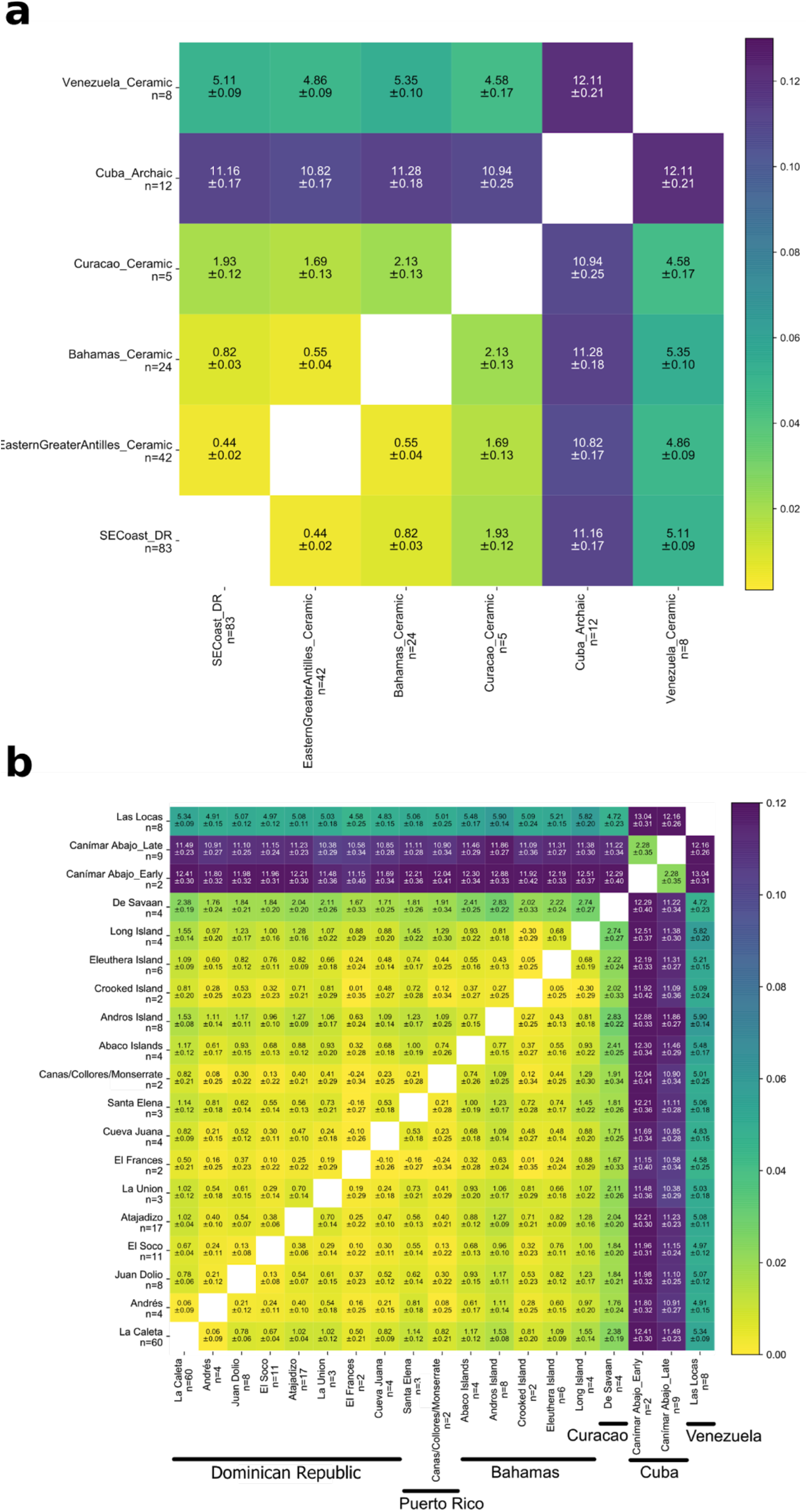
F_ST_ distances by site and clades. Average pairwise FST distances (x100) between **(a)** clades and **(b)** sites with more than two unrelated individuals, demonstrating both overall high levels of genetic similarity between the *Caribbean_Ceramic* sub-clades and the sites composing them, as well as the magnitude of genetic differentiation between those and the groups with Archaic- and Venezuela-related ancestries.

**Extended Data Fig. 2:**
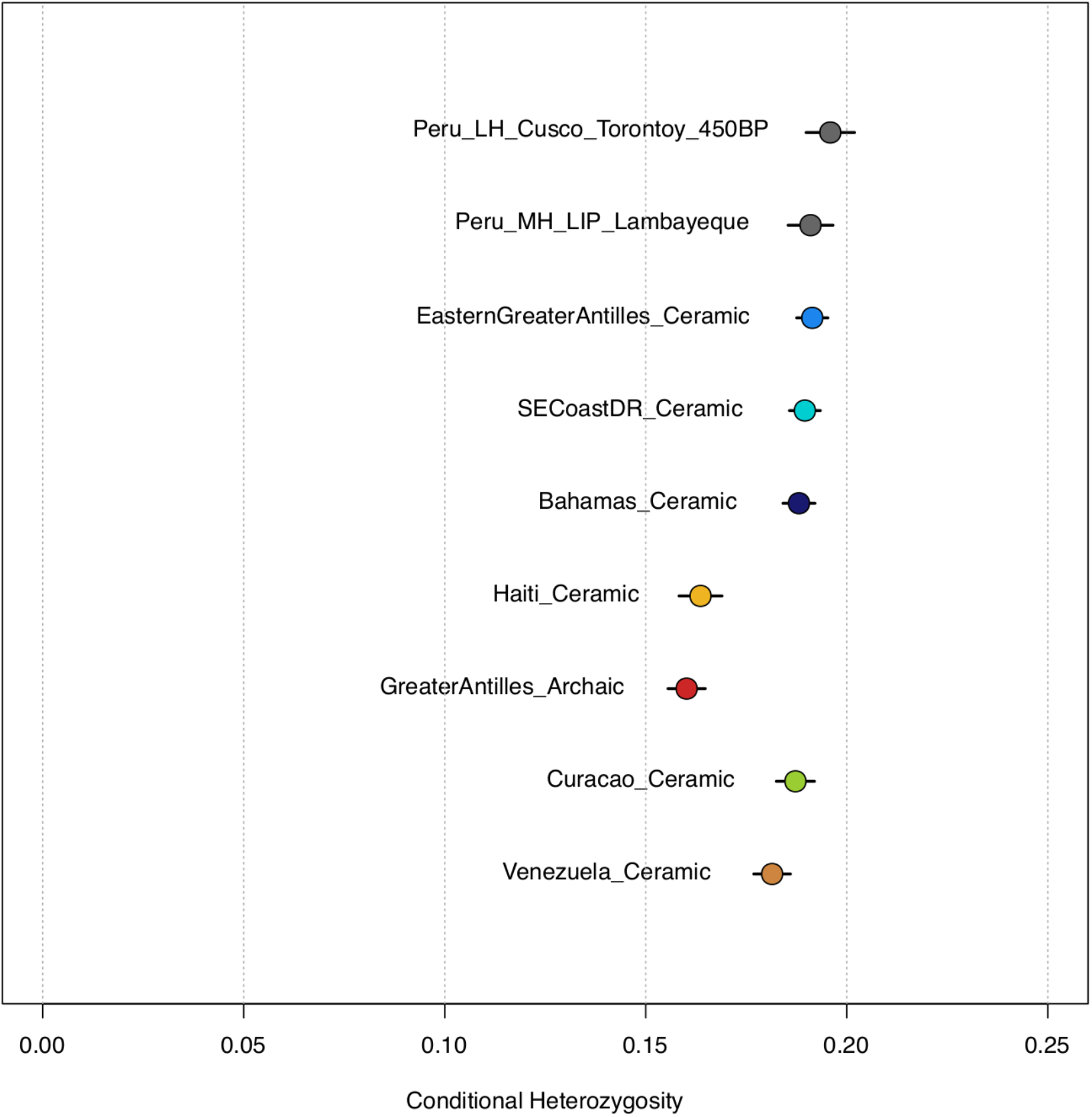
Conditional heterozygosity by clade. Conditional heterozygosity in the ancient Caribbean was similar to that of contemporaneous groups from Peru, except for the Archaic-associated groups, *Venezuela_Ceramic*, and admixed Ceramic-associated populations (Nakatsuka et al. 2020). Bars represent 3 standard errors.

**Extended Data Table 1:** 2N_e_ estimates for each site. Table includes all individuals where ROH analysis is possible and excludes individuals with more than 50cM sum of 20cM long ROH.

| $2N_e$ estimate | STD | 95% CI (low) | 95% CI (high) | n | Site (Country) | Clade |
| --- | --- | --- | --- | --- | --- | --- |
| 12670 | 63 | 1146 | 1394 | 51 | La Caleta (Dominican Republic) | Caribbean_Ceramic |
| 1673 | 340 | 1007 | 2339 | 4 | Andrés (Dominican Republic) | Caribbean_Ceramic |
| 2298 | 411 | 1493 | 3104 | 7 | Juan Dolio (Dominican Republic) | Caribbean_Ceramic |
| 1814 | 255 | 1314 | 2314 | 9 | El Soco (Dominican Republic) | Caribbean_Ceramic |
| 1549 | 152 | 1252 | 1846 | 16 | Atajadizo (Dominican Republic) | Caribbean_Ceramic |
| 1850 | 455 | 958 | 2741 | 3 | La Union (Dominican Republic) | Caribbean_Ceramic |
| 1302 | 332 | 652 | 1953 | 2 | El Frances (Dominican Republic) | Caribbean_Ceramic |
| 1966 | 498 | 990 | 2942 | 3 | Cueva Juana (Dominican Republic) | Caribbean_Ceramic |
| 1850 | 455 | 959 | 2742 | 3 | Santa Elena (Puerto Rico) | Caribbean_Ceramic |
| 1223 | 303 | 629 | 1816 | 2 | Canas/Collores/Monserrate (Puerto Rico) | Caribbean_Ceramic |
| 943 | 169 | 612 | 1273 | 3 | Abaco Islands (Bahamas) | Caribbean_Ceramic |
| 1123 | 189 | 753 | 1493 | 4 | Andros Island (Bahamas) | Caribbean_Ceramic |
| 1220 | 302 | 628 | 1812 | 2 | Crooked Island (Bahamas) | Caribbean_Ceramic |
| 1394 | 259 | 886 | 1902 | 4 | Eleuthera (Bahamas) | Caribbean_Ceramic |
| 1906 | 582 | 765 | 3048 | 2 | Diale 1(Haiti) | Haiti_Ceramic |
| 937 | 205 | 535 | 1339 | 2 | De Savaan (Curaçao) | Curacao_Ceramic |
| 504 | 119 | 272 | 737 | 1 | Andrés (Dominican Republic) | GreaterAntilles_Archaic_10126 |
| 482 | 78 | 328 | 636 | 2 | Canimar Abajo: Early (Cuba) | GreaterAntilles_Archaic |
| 899 | 193 | 521 | 1277 | 2 | Canimar Abajo: Late (Cuba) | GreaterAntilles_Archaic |
| 514 | 50 | 417 | 612 | 6 | Las Locas (Venezuela) | Venezuela_Ceramic |

**Extended Data Table 2:**
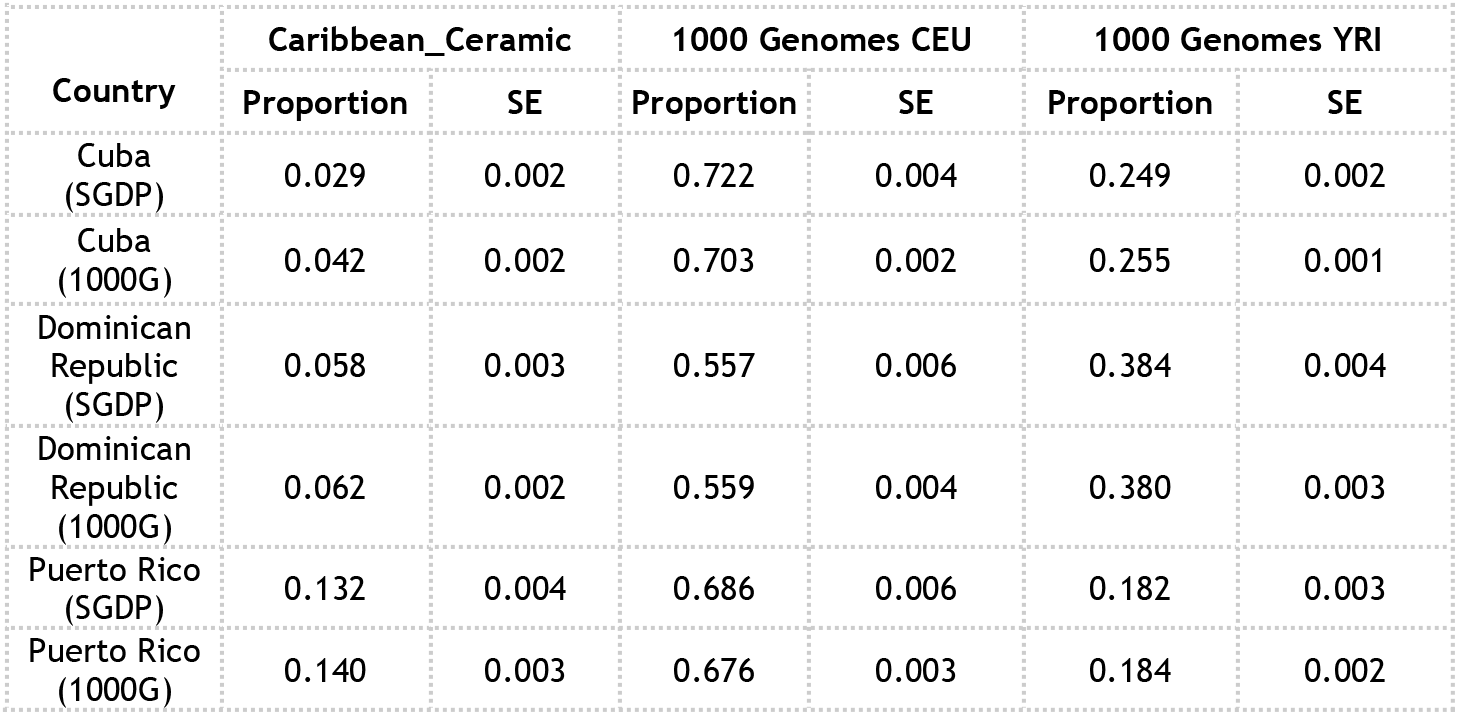
Ancestry proportion estimates with *qpAdm* of Indigenous, European, and African sources in present-day Caribbean individuals from Cuba, Dominican Republic, and Puerto Rico from the GOAL study (Moreno-Estrada et al. 2013). SGDP = Simons Genome Diversity Project outgroup populations Karitiana, Mixe, Yakut, Ulchi, Papuan, Mursi, and Mbuti; 1000G = 1000 Genomes outgroup populations PEL, PJL, JPT, and MSL.

| Country | Caribbean_Ceramic |  | 1000 Genomes CEU |  | 1000 Genomes YRI |  |
| --- | --- | --- | --- | --- | --- | --- |
|  | Proportion | SE | Proportion | SE | Proportion | SE |
| Cuba (SGDP) | 0.029 | 0.002 | 0.722 | 0.004 | 0.249 | 0.002 |
| Cuba (1000G) | 0.042 | 0.002 | 0.703 | 0.002 | 0.255 | 0.001 |
| Dominican Republic (SGDP) | 0.058 | 0.003 | 0.557 | 0.006 | 0.384 | 0.004 |
| Dominican Republic (1000G) | 0.062 | 0.002 | 0.559 | 0.004 | 0.380 | 0.003 |
| Puerto Rico (SGDP) | 0.132 | 0.004 | 0.686 | 0.006 | 0.182 | 0.003 |
| Puerto Rico (1000G) | 0.140 | 0.003 | 0.676 | 0.003 | 0.184 | 0.002 |

**Extended Data Table 3:** Ancestry proportion estimates with *qpAdm* of Indigenous, European, African, and East Asian sources in present-day individuals across different Cuban provinces. Outgroup populations used: PEL, PJL, JPT, MSL and GIH.

| Province | 1000 Genomes CEU |  | 1000 Genomes YRI |  | Caribbean_Ceramic |  | 1000 Genomes CHB |  |
| --- | --- | --- | --- | --- | --- | --- | --- | --- |
|  | Proportion | SE | Proportion | SE | Proportion | SE | Proportion | SE |
| Artemisa | 0.834 | 0.005 | 0.100 | 0.003 | 0.038 | 0.004 | 0.028 | 0.004 |
| Camaguey | 0.617 | 0.004 | 0.297 | 0.002 | 0.074 | 0.003 | 0.013 | 0.003 |
| Ciego_de_Avila | 0.788 | 0.004 | 0.145 | 0.002 | 0.057 | 0.003 | 0.010 | 0.003 |
| Cienfuegos | 0.740 | 0.004 | 0.220 | 0.003 | 0.028 | 0.003 | 0.012 | 0.003 |
| Granma | 0.567 | 0.003 | 0.271 | 0.002 | 0.144 | 0.003 | 0.018 | 0.002 |
| Guantanamo | 0.549 | 0.003 | 0.363 | 0.003 | 0.083 | 0.002 | 0.004 | 0.002 |
| Holguin | 0.655 | 0.003 | 0.237 | 0.002 | 0.095 | 0.002 | 0.013 | 0.002 |
| La_Habana | 0.694 | 0.003 | 0.257 | 0.002 | 0.033 | 0.002 | 0.015 | 0.002 |
| Las_Tunas | 0.725 | 0.007 | 0.161 | 0.004 | 0.113 | 0.005 | 0.001 | 0.005 |
| Matanzas | 0.818 | 0.003 | 0.140 | 0.002 | 0.016 | 0.003 | 0.026 | 0.003 |
| Mayabeque | 0.889 | 0.005 | 0.940 | 0.003 | 0.012 | 0.004 | 0.005 | 0.004 |
| Pinar_del_Rio | 0.727 | 0.003 | 0.227 | 0.002 | 0.036 | 0.002 | 0.010 | 0.002 |
| Sancti_Spiritus | 0.809 | 0.003 | 0.108 | 0.002 | 0.065 | 0.003 | 0.018 | 0.003 |
| Santiago_de_Cuba | 0.501 | 0.003 | 0.417 | 0.002 | 0.076 | 0.002 | 0.006 | 0.002 |
| Villa_Clara | 0.813 | 0.003 | 0.106 | 0.002 | 0.066 | 0.002 | 0.016 | 0.002 |

**Extended Data Table 4:** Present day Indigenous American populations used per analysis, their broad language family attribution, and dataset of origin. IL= Illumina, HO = Human Origins.

| Population | Language Family | Included in PCA | Included in ADMIXTURE | Included in qpWave/qpAdm | Included in Treemix | Included in outgroup-f3 | Included in language group f4-statistics |
| --- | --- | --- | --- | --- | --- | --- | --- |
| Chipewyan | Na-Dene | - | Yes (IL, n=2) | Yes (HO, n=28),<br>Yes (IL, n=2) | Yes (IL, n=2) | - | - |
| Aymara | Andean | Yes (IL, n=4) | Yes (IL, n=4) | - | Yes (IL, n=4) | Yes (IL, n=4) | - |
| Quechua | Andean | Yes (IL, n=1) | Yes (IL, n=1) | Yes (HO, n=7),<br>Yes (IL, n=1) | Yes (IL, n=1) | Yes (IL, n=1) | - |
| Arara | Carib | Yes (IL, n=1) | Yes (IL, n=1) | Yes (HO, n=4),<br>Yes (IL, n=1) | Yes (IL, n=1) | Yes (IL, n=1) | Yes (IL, n=1) |
| Apalai | Carib | - | - | Yes (HO, n=4) | - | - | - |
| Yukpa | Carib | Yes (1240K, n=1) | Yes (1240K, n=1) | Yes (HO, n=1) | - | - | - |
| Mixtec | Central Amerind | Yes (1240K, n=1) | Yes (1240K, n=1) | Yes (HO, n=10) | - | - | - |
| Zapotec | Central Amerind | Yes (IL, n=5) | Yes (IL, n=5) | Yes (HO, n=10),<br>Yes (IL, n=5) | Yes (IL, n=5) | Yes (IL, n=5) | - |
| Nahua | Central Amerind | Yes (1240K, n=2) | Yes (1240K, n=2) | - | - | - | - |
| Pima | Central Amerind | Yes (IL, n=14) | Yes (IL, n=14) | - | Yes (IL, n=14) | Yes (IL, n=14) | - |
| Tepehuano | Central Amerind | Yes (IL, n=2) | Yes (IL, n=2) | - | Yes (IL, n=2) | Yes (IL, n=2) | - |
| Bribri | Chibchan | Yes (IL, n=2) | Yes (IL, n=2) | - | Yes (IL, n=2) | Yes (IL, n=2) | Yes (IL, n=2) |
| Cabecar | Chibchan | Yes (IL, n=24) | Yes (IL, n=24) | Yes (HO, n=6),<br>Yes (IL, n=24) | Yes (IL, n=24) | Yes (IL, n=24) | Yes (IL, n=24) |
| Guaymi | Chibchan | Yes (IL, n=5) | Yes (IL, n=5) | - | Yes (IL, n=5) | Yes (IL, n=5) | Yes (IL, n=5) |
| Kogi | Chibchan | Yes (IL, n=4) | Yes (IL, n=4) | - | - | Yes (IL, n=4) | Yes (IL, n=4) |
| Maleku | Chibchan | Yes (IL, n=2) | Yes (IL, n=2) | - | Yes (IL, n=2) | Yes (IL, n=2) | Yes (IL, n=2) |
| Teribe | Chibchan | Yes (IL, n=2) | Yes (IL, n=2) | - | Yes (IL, n=2) | Yes (IL, n=2) | Yes (IL, n=2) |
| Waunana | Chocoan | Yes (IL, n=3) | Yes (IL, n=3) | - | - | Yes (IL, n=3) | Yes (IL, n=3) |
| Embera | Chocoan | Yes (IL, n=5) | Yes (IL, n=5) | - | - | Yes (IL, n=5) | Yes (IL, n=5) |
| Guahibo | Guajibuan | Yes (IL, n=6) | Yes (IL, n=6) | - | Yes (IL, n=6) | Yes (IL, n=6) | Yes (IL, n=6) |
| Jamamadi | Macro-Arawakan | Yes (IL, n=1) | Yes (IL, n=1) | - | Yes (IL, n=1) | Yes (IL, n=1) | Yes (IL, n=1) |
| Palikur | Macro-Arawakan | Yes (IL, n=2) | Yes (IL, n=2) | - | Yes (IL, n=2) | Yes (IL, n=2) | Yes (IL, n=2) |
| Piapoco | Macro-Arawakan | Yes (IL, n=6) | Yes (IL, n=6) | Yes (HO, n=4),<br>Yes (IL, n=6) | Yes (IL, n=6) | Yes (IL, n=6) | Yes (IL, n=6) |
| Wayuu | Macro-Arawakan | Yes (IL, n=3) | Yes (IL, n=3) | Yes (HO, n=1) | - | Yes (IL, n=3) | Yes (IL, n=3) |
| Chane | Macro-Arawakan | Yes (IL, n=2) | Yes (IL, n=2) | - | Yes (IL, n=2) | Yes (IL, n=2) | Yes (IL, n=2) |
| Toba | Mataco-Guaicuru | Yes (IL, n=2) | Yes (IL, n=2) | - | Yes (IL, n=2) | Yes (IL, n=2) | Yes (IL, n=2) |
| Wichi | Mataco-Guaicuru | Yes (IL, n=4) | Yes (IL, n=4) | - | Yes (IL, n=4) | Yes (IL, n=4) | Yes (IL, n=4) |
| **Kaqchikel** | Northern Amerind | Yes (IL, n=1) | Yes (IL, n=1) | - | Yes (IL, n=1) | Yes (IL, n=1) | - |
| **Maya** | Northern Amerind | Yes (IL, n=1) | Yes (IL, n=1) | - | - | - | - |
| **Mixe** | Northern Amerind | Yes (IL, n=9) | Yes (IL, n=9) | Yes (HO, n=10),<br>Yes (IL, n=9) | Yes (IL, n=9) | Yes (IL, n=9) | - |
| **Guarani** | Tupian | Yes (IL, n=3) | Yes (IL, n=3) | - | Yes (IL, n=3) | Yes (IL, n=3) | Yes (IL, n=3) |
| **Karitiana** | Tupian | Yes (IL, n=13) | Yes (IL, n=13) | Yes (HO, n=16),<br>Yes (IL, n=13) | Yes (IL, n=13) | Yes (IL, n=13) | Yes (IL, n=13) |
| **Parakana** | Tupian | Yes (IL, n=1) | Yes (IL, n=1) | - | Yes (IL, n=1) | Yes (IL, n=1) | Yes (IL, n=1) |
| **Surui** | Tupian | Yes (IL, n=24) | Yes (IL, n=24) | Yes (HO, n=12),<br>Yes (IL, n=24) | Yes (IL, n=24) | Yes (IL, n=24) | Yes (IL, n=24) |
| **Ticuna** | Ticuna (isolate) | Yes (IL, n=5) | Yes (IL, n=5) | - | Yes (IL, n=5) | Yes (IL, n=5) | Yes (IL, n=5) |
| **Yaghan** | Yaghan (isolate) | Yes (IL, n=1) | Yes (IL, n=1) | - | Yes (IL, n=1) | Yes (IL, n=1) | - |

**Extended Data Table 5:**
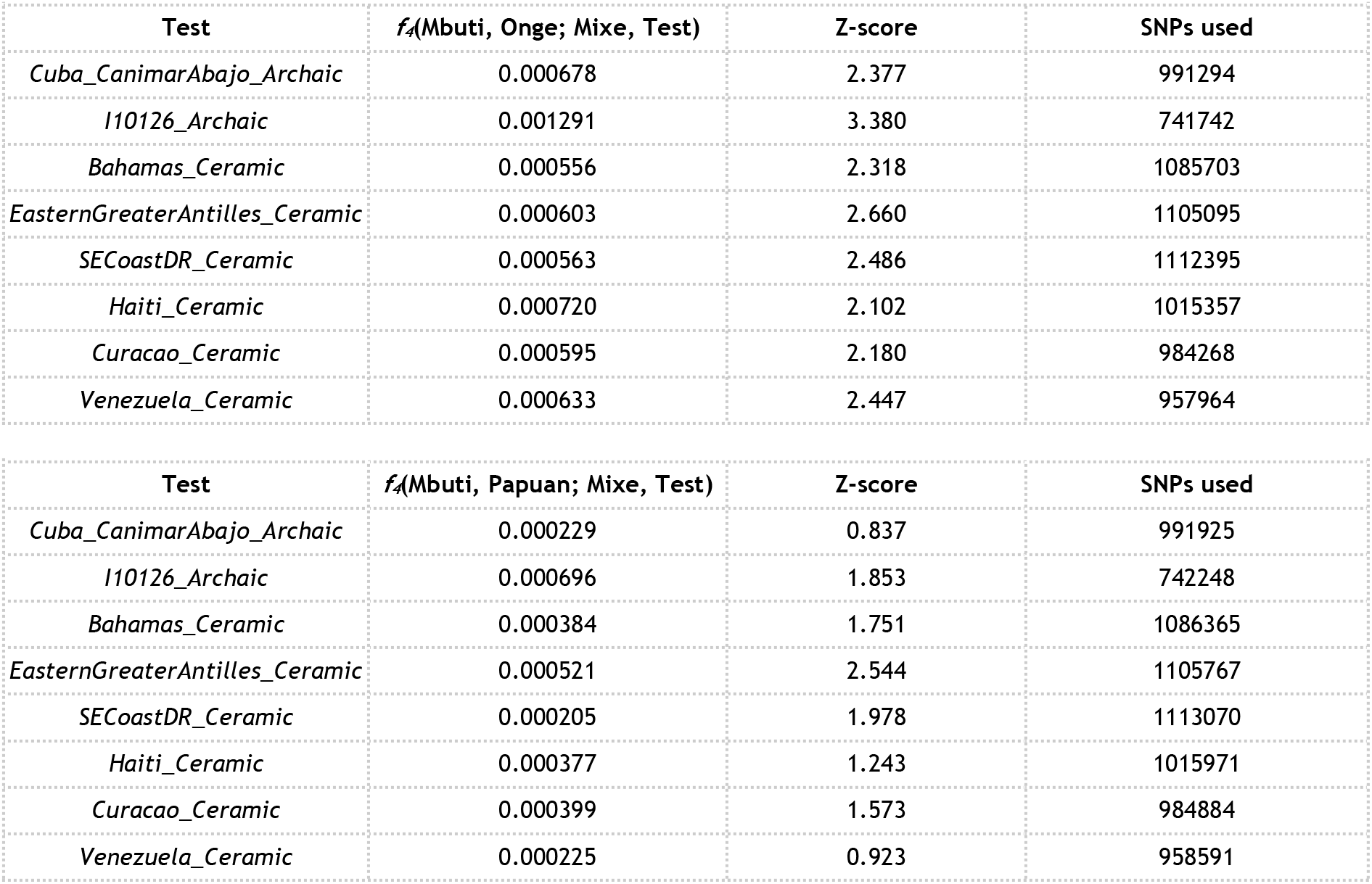
Statistics testing for an Australasian link.

