## Supplementary Information for "A genetic history of the pre-contact Caribbean"

### TABLE OF CONTENTS

**SI section 1.** Overview of archaeological series and culture history 3

**SI section 2.** Archaeological site information 16

**SI section 3.** Newly reported direct $^{14}C$ dates 47

**SI section 4.** Principal component analysis 49

**SI section 5.** Unsupervised population structure analysis 52

**SI section 6.** Kinship, consanguinity and conditional heterozygosity 54

**SI section 7.** Clade Grouping and Substructure Analysis with *qpWave*, Treemix and $f_4$-statistics 60

**SI section 8.** Admixture modeling and estimates of ancestry proportions 71

**SI section 9.** Uniparental haplogroups 81

**SI section 10.** $f$-statistics and relatedness to modern language groups 86

**SI section 11.** *qpGraph* 87

**SI section 12.** Ability to detect a Carib migration into the Caribbean: simulations and *qpWave* 88

**SI section 13.** DATES 90

**SI section 14.** Relatedness of ancient individuals to present-day admixed Caribbean people 91

**SI section 15.** Ethics statement 92

**References** 94

**SI1 - Overview of archaeological series and culture history in the Caribbean and northern South America**

**The Caribbean**

Similarities and differences in material culture are arranged in classification systems to identify patterns in the archaeological record of a region. Two taxonomic systems are employed for the Caribbean islands. The first and most general is an “Age” system based on the introduction of significant changes in technology. The second is the time-space periodization of internal developments as reflected through changes in style (“Series”). These systems are hierarchical in that “internal” stylistic changes are subsumed under “external” technological introductions.

The “Age” system is divided into Lithic, Archaic, Ceramic, and Historic Ages (Rouse 1992). Irving Rouse, a foundational figure in Caribbean archaeology, developed the system to recognize the most significant changes in technology observed in the region. Technological change occurs in three main ways: independent invention, diffusion of ideas, or the movement of people who carry the new technology. The latter (the arrival of new immigrants from the mainland bearing new technology) was assumed when developing this nomenclature. Working within this frame of reference, the pattern of human movement(s) into the islands was reconstructed primarily through comparisons of technologies between Caribbean archaeological sites and mainland material assemblages (Wilson 2007). In this approach, technological change is viewed as the product of external influences, and the introduction of new technologies supposedly leads to the physical and/or cultural displacement of previously established practices (Rouse 1992). In reality, even if cultural practices changed, existing technology(s) usually continued in use, albeit often in different ways (Rodríguez Ramos 2010).

We begin by providing a brief description of three distinct Ages of Caribbean occupation (we do not focus in detail on the Historic Age here). These Ages began and ended at different times on different islands; therefore, we provide the earliest date for each Age. For ease of interpretation, we discuss archaeological periods and corresponding ceramic typological styles in this section and the following section using BCE/CE dates; in the main manuscript, we discuss dates using years before present (BP, taken as AD 1950 in accordance with radiocarbon calibration convention; we use calibrated years BP (cal. yr BP) when direct radiocarbon ($^{14}C$) dates are available). We provide an explanation for the presentation of dates in this paper in Supplementary Information section 3.

**Lithic Age (4000 BCE)**

The Caribbean Islands were first settled by humans around 4000 BCE, although this earliest phase of human occupation is not well studied or adequately dated. The majority of sites are in Cuba, Haiti, Dominican Republic, and Puerto Rico (Pantel 1988), with evidence of sites in the Lesser Antilles and ABC islands (Aruba, Bonaire, Curaçao) as well (Napolitano et al. 2019). The Lithic Age is characterized by flaked-stone technologies most often associated with the production of large blades. It has been argued that the first colonizers of the Caribbean came from Central America/Yucatán because

Antillean Casimiroid blades (Casimiroid is named for the Casimira site, Dominican Republic) resemble blades produced in Belize at about the same time (Wilson et al. 1998). However, there is not universal agreement about this geographic localization (Callaghan 2001). Little is known about these first inhabitants of the Antilles. Most of the archaeological sites are quarries where the raw material for tool manufacture was procured. It is assumed they were Paleo-Indian hunter-gatherers who lived in small bands and lacked both cultigens and pottery (Rouse 1992). In fact, all that is known is that humans arrived in the Antilles by 4000 BCE and the only surviving element of material culture is flaked-stone tools (Keegan and Hofman 2017).

**Archaic Age (3000 BCE)**

The beginning of the Archaic Age is marked by the appearance of ground-stone tools, shell implements, and the production of freehand flakes using multidirectional flaking formats (Rouse 1992). Archaeological sites from this time are distributed from Cuba in the Greater Antilles through the Leeward Islands of the northern Lesser Antilles, although none have been identified in Jamaica (Keegan 2019). The most common interpretation of this distribution is that the spread of this technology reflects a new migration of people originating from Trinidad, and that these Ortoiroid immigrants (Ortoiroid is named for sites at Ortoire, Trinidad) encountered Casimiroid peoples in the Greater Antilles and effectively ended the Lithic Age (Rouse 1992; Boomert 2013). Interestingly, no Archaic Age artifacts have been found in the Windward Islands of the southern Lesser Antilles (Keegan and Hofman 2017) even though they are widely distributed from Panama (in Central America) to Trinidad (Rodríguez Ramos 2008). There is evidence for fires at this time in this region (Siegel et al. 2015), but no evidence that humans were responsible. Either the Windward Islands were bypassed by Ortoiroid canoes, or Archaic Age practices reached the islands by independent invention within the Caribbean and diffusion among the islands.

The absence of substantial quantities of pottery at Archaic Age sites has led to the belief that they too were hunter-gatherers who lived in small mobile bands. A key adaptation for this group was expanding their diet to include plants processed with grinding tools and a variety of marine foods, especially mollusks (Davis 2000). A problem with the Age system, especially for the Lithic and Archaic Ages, is the tendency to assume that these societies were culturally static, and that they were all isolated groups lacking both pottery and farming. However, recent investigations demonstrate that cultigens, including maize and beans, were grown from early in the Archaic Age (Pagán Jiménez 2013; Smith et al. 2018); hutia (*Capromyidae*) may have been domesticated (Colten and Worthington 2019); pottery was made and used, albeit in small quantities and for special purposes (Rodríguez Ramos 2008); and some of the groups developed hierarchical social formations (Rivera-Collazo 2011).

Recent research has shown that Archaic Age practices persisted much longer than initially presumed. There is evidence that the Archaic inhabitants of Hispaniola prevented the early Ceramic Age inhabitants of Puerto Rico from settling on the island in large numbers for about 1,000 years. Archaic and Ceramic Age sites overlap for some time on Puerto Rico (Rodríguez Ramos 2010; Rivera-Collazo 2011) and probably other islands. Archaic Age societies in Cuba have been reported to have persisted uninterrupted at the Camínar Abajo site until as late as 950 calibrated years CE (calCE; Roksandic et al. 2015) and in the Banes Archaeological Zone of northeastern Cuba until after 900 calCE (Persons 2013). Autochthonous Guanahatabey and Ciboney groups appear to have remained in the western and central parts of Cuba (respectively) for such an extended period of time that the first historical European accounts describe peoples of a different language and cultural tradition than contemporaneous ceramic-using groups (Lovén 1935; Chinique de Armas et al. 2016; Keegan and Hofman 2017).

**Ceramic Age (500-200 BCE)**

The Ceramic Age is distinguished by the introduction of pottery and agriculture almost certainly by Arawak-speaking peoples from South America. Although Archaic Age peoples planted cultigens and made pottery, their practices differed substantially from those of the Early Ceramic Age. The former is highly distinct from the latter, which is why the apparent misnomer of Ceramic Age remains in use. We use it in this study to be consistent with the literature.

In what follows, we move from the external Age system to the internal "Series" taxonomy (see Rouse 1972, 1992; Curet 2004; Bérard 2019). The finest-grained unit is the "style," which is identified as a shared set of common technical and decorative "modes" (e.g., vessel shape, paste, decoration). Styles are named for the first archaeological site at which this collection of modes were observed and are therefore not necessarily the site most representative of the style. Islands that are in close proximity to each other typically have at least one local style that shares modes with a local style on another island; these are grouped into a larger taxonomic unit called a "series," which Rouse (1992) interpreted as representing "peoples or cultures." Series also were named for the first site at which the specific collection of modes was identified, and were designated with the suffix '-oid.' Subseries, with the suffix '-an,' were later added to emphasize when series were thought to evolve from a single tradition (Rouse 1992). For example, the Cabo Rojo site, located at Punta Ostiones, Puerto Rico, is the first at which a particular style of redware pottery was observed; thus, the local style was named "Ostiones." Other local styles with similar pottery (e.g., Santa Elena in Puerto Rico, Anadel in Dominican Republic, Little River in Jamaica) were deemed similar enough to be grouped into an Ostionoid series.

Rouse (1992) identified four series for the Antilles: Saladoid, Ostionoid, Meillacoid, and Chicoid, ordered sequentially (we do not discuss the Troumassoid series specific to the Lesser Antilles in detail, as no individuals from this part of the Caribbean are included in this study). As with changes in technology in the Age system, the development of every new series completely replaced the previous series, such that the preceding series only survived on the margins. When other archaeologists suggested that each of the series represented separate migrations from the mainland into the islands (see below), Rouse (1992) introduced the concept of subseries, denoted by the suffix "-an," to emphasize his belief that all of the ceramic series evolved from a single Ostionoid tradition.

His original series were subsequently converted to Ostionoan Ostionoid, Meillacan Ostionoid, and Chican Ostionoid.

Rouse (1992) envisioned the Ceramic Age as the product of a single population expansion (migration) that began on the banks of the Orinoco River in eastern Venezuela and proceeded “island by island” through the Lesser Antilles and on to the Greater Antilles (Willey 1971:364-365). His singular tradition taxonomy should be viewed as a theory that requires appropriate testing to assess its validity. Detailed discussions of the competing models that have been proposed to explain the relationships and transitions among the Ceramic Age cultures are presented in *Caribbean before Columbus* (Keegan and Hofman 2017) and *Handbook of Caribbean Archaeology* (Keegan et al. 2013). A brief introduction to these models with specific attention to human mobility is discussed below.

*Early Ceramic Age (500-200 BCE to 900 CE): Saladoid, Huecoid, and Saladoid/Barrancoid*

The Early Ceramic Age (ECA) migrants originated in lowland South America and reached the Antilles sometime between 500-200 BCE (Lathrap 1970; Rouse 1986; Keegan and Hofman 2017). The ECA is often described solely in terms of Saladoid series pottery (Saladoid is named for the Saladero site on the lower Orinoco River in eastern Venezuela) and the cultural practices inferred from Saladoid sites. The pottery exhibits distinctive complex vessel forms, white-on-red painting (W-O-R), incised designs, and zoomorphic adornos (animal-shaped appendages) that link it to the Orinoco basin in present-day Venezuela and the Guianas (Rouse 1992; Boomert 2013), a region which today is home to some Indigenous Arawak-speaking groups. The Saladoid tradition has been identified throughout the Lesser Antilles, Puerto Rico, and possibly eastern Dominican Republic, but it does not occur in Haiti, Jamaica, Cuba, the Bahama archipelago, or the ABC Islands of Aruba, Bonaire, and Curaçao. Peoples carrying this tradition may have absorbed or mixed with autochthonous Archaic populations of the islands (Wilson 2007), though previously established communities may have remained as independent entities at least in some areas (Keegan and Hofman 2017; the results in the present study also provide support for the existence of Archaic-associated groups well into the Ceramic Age).

Although Saladoid was the first ECA tradition identified, two additional and contemporaneous cultural traditions have since received widespread acceptance: 1) the Huecoid pottery series (Chanlatte 2013), which has distinctive vessel shapes but lacks painted motifs, is centered on eastern Puerto Rico and nearby Vieques Island (Huecoid is named for the La Hueca site, Vieques Island, located off of eastern Puerto Rico) and occurs at archaeological sites throughout the Lesser Antilles; and 2) the Saladoid/Barrancoid tradition, which is found is in the Windward Islands (Keegan and Hofman 2017; Rouse 1992). It is important to understand that these different traditions are identified primarily by differences in pottery styles. Small differences may also reflect local group identities subsumed by broader similarities that serve to unify widely dispersed autonomous communities. The manner in which ECA peoples came to settle the Lesser Antilles and Puerto Rico has been the subject of intense debate.

The first model was proposed prior to the general availability of radiocarbon dates and at a time when Saladoid was the only cultural tradition. The movement of Saladoid people at the onset of the ECA was proposed as a general 'stepping-stone' movement ("island by island") from South America to Trinidad, through the Lesser Antilles, and eventually into Puerto Rico (Boomert 2013; Rouse 1986; Willey 1971:364-365). There are two major problems with this model. First, humans cannot reproduce fast enough to settle every island in turn and still reach Puerto Rico at an early date (Keegan 1995). Thus, either some islands were bypassed or population expansion occurred in waves. Expansion in waves fits the proposed separate arrival of Huecoid possibly earlier than the arrival of Saladoid (Chanlatte Baik 2013).

The second problem is that the earliest radiocarbon dates for Saladoid and Huecoid (circa 500-200 BCE) all cluster in the northern Lesser Antilles and Puerto Rico (Fitzpatrick et al. 2010; Napolitano et al. 2019). The evidence suggests that the northern Lesser Antilles and Puerto Rico were settled in a direct jump across the Caribbean Sea. Pottery styles similar to the Saladoid series are distributed along coastal South America (e.g., Huecoid exhibits similarities with the Río Guapo style), while various shell and exotic stone ornaments are similar to objects from the Isthmo-Colombian area (Rodríguez Ramos 2013). Based on material similarities, it is not clear precisely where in South America the ECA colonists originated; however, linguistic studies suggest they spoke an Arawak language, consistent with pottery styles connected to eastern Venezuela.

The first evidence for people in the Windward Islands is dated to the first centuries CE (Napolitano et al. 2019). Saladoid sites in the southern Lesser Antilles exhibit stronger affiliation with Barrancoid series pottery, which is characterized by an abundance of finely made zoomorphic adornos (Keegan and Hofman 2017). It is possible that this reflects yet another wave of colonists from South America, this time from the Orinoco delta region. Alternatively, because these later sites share W-O-R painted pottery with earlier Saladoid sites to the north, other archaeologists have argued that these cultures do not necessarily reflect a separate migration.

Saladoid pottery has been described as a "veneer," meaning that widely shared modes highlighted regional integration and masked local variability (Keegan 2004). These modes begin to disappear around 500-600 CE, especially in Puerto Rico, although they continue in use until around 900 CE on some islands (Versteeg and Schinkel 1992). In the Greater Antilles these new styles are classified as part of the Ostionoid series, while emerging styles in the Lesser Antilles are classified as Troumassoid (Ostionoid is named for sites near Punta Ostiones, western Puerto Rico, and Troumassoid is named for Troumassee site, St. Lucia). Troumassoid is considered an *in situ* regional development that did not involve the arrival of peoples from the mainland.

*Ostionoid (600-900 CE)*

Ostionoid series pottery is characterized by distinctive red surface treatments, the disappearance of W-O-R painting, simple lugs and rim projections, and incised designs (with regional variation).

Ostionoid and Saladoid are conventionally interpreted as demographically continuous, in the sense that people descended from the same demographic movement (likely from the South American continent) practiced both. However, it remains difficult to differentiate between the movement of people and cultural diffusion using archaeological data alone (Keegan and Hofman 2017). The most common explanation is that Ostionoid developed from Saladoid within the islands. It is associated with a resumption of population expansion to the west following a 1,000-year period of adaptation to island conditions (Rouse 1992). In this regard it is viewed as a new wave of inter-island mobility that arrived first in Hispaniola (present-day Haiti and the Dominican Republic), then in Jamaica (with the first permanent human inhabitants of that island), and finally in Cuba (although Cuban archaeologists do not recognize Ostionoid pottery). It then moved north ~700 CE to Grand Turk in the Turks and Caicos Islands via seasonal visitors from Hispaniola (Carlson 1999).

It is noteworthy that Ostionoid pottery appears simultaneously in Puerto Rico, Hispaniola, Jamaica, and the southern Bahamas. This may reflect the rapid pace at which the culture expanded, or the limits of dating methods to confirm directionality. An alternative perspective, based on the abundance of crab claws in Saladoid sites and of mollusk shells in Ostionoid sites (or their respective absence), is that these series represent separate migrations, although no homeland was proposed for the Ostionoid expansion (Rainey 1940). Rather than resulting from a separate migration, Keegan (2006) and Rodríguez Ramos (2005) have suggested that Ostionoid pottery is a form of Archaic Age pottery that originated during the Archaic Age, belonging to the Pre-Arawak Pottery Horizon (Rodríguez Ramos 2008). This would account for its sudden appearance, wide distribution, and rapid disappearance. In Jamaica, for example, it is thought that new immigrants bearing Meillacoid series pottery quickly replaced Ostionoid, and there is no evidence that the two interacted (Keegan 2019). In sum, the humans associated with Ostionoid pottery would have a Saladoid ancestry (Rouse 1992), an Archaic ancestry (Keegan 2006, Rodríguez Ramos 2008), or a different but unidentified ancestry (Rainey 1940). Ancient DNA data provides power to distinguish between these hypotheses.

*Meillacoid (800-1500 CE)*

The next major material culture change during the Ceramic Age commences around 800 CE. The earliest evidence of this tradition is in central Hispaniola near the border between Haiti and the Dominican Republic. Meillacoid series pottery is characterized by narrow V-shaped incisions (often unsmoothed) combined in oblique-parallel-line motifs, crosshatching, punctations, appliqué, and constructed adornos. Meillacoid represented the third wave of inter-island cultural expansion, which Rouse (1986) ultimately traced to the original Saladoid settlement of the islands. Using subseries terminology, Meillacan belonged to the Ostionoid, and reflected the incorporation of design motifs copied from Archaic stone bowls. Meillacoid pottery spread across Hispaniola, and then diffused to Jamaica and Cuba sometime after 900-1000 CE (Rouse 1992). It is supposed to have disappeared in Hispaniola around 1200 CE following the spread of Chicoid pottery, although it continued in Jamaica and Cuba until European colonization.

Marcio Veloz Maggiolo (1972, 1993) and Alberta Zucchi (1985) have suggested a different history of the Meillacoid tradition. According to their reports, there were other pottery-making traditions in Hispaniola that preceded Meillacoid (and Ostionoid) pottery styles by perhaps 1,000 years (Veloz Maggiolo and Ortega 1996) and are contemporaneous with Saladoid in Puerto Rico. These styles, culminating in Meillacoid, reflect a direct connection to the South American mainland. A study of pre-Columbian facial morphology suggested that people living in Hispaniola, Jamaica, and The Bahamas formed a single cluster (Ross et al. 2020). Meillacoid pottery is the one element of material culture shared across these islands, and the V-shaped incisions and appliqué in the design of the pottery are also associated with “Carib” pottery in coastal South America (Lathrap 1970; Meggers et al. 1965). It is possible that even if it does represent a movement of people, the Carib expansion could have involved the infiltration of Indigenous, Arawak-speaking communities, by small numbers of people, thus explaining the persistence of Arawak languages throughout these regions.

*Lucayan Palmetto Ware (800-1550 CE)*

Locally produced pottery in the Bahamian archipelago (today including the independent Commonwealth of The Bahamas and the British Overseas Territory, Turks and Caicos Islands) is called Palmetto Ware, which is named for the Palmetto Grove site on the island of San Salvador in The Bahamas. It is a “ware,” and not a series, because it is represented by only one type. There are no additional Palmetto styles. Palmetto Ware is a redware made using local clay and calcified shells from the conch (*Lobatus gigas*). It is so different from Ostionoid and Meillacoid pottery that Rouse was unwilling to attribute it to either tradition. Most Palmetto Ware is undecorated, but when it is, it exhibits Meillacoid motifs. The recent accelerator mass spectrometry (AMS) dating of 60 Lucayan skeletons from across The Bahamas returned dates in the 900-1550 CE range (Schulting et al., *in preparation*), although it is impossible to be certain that the earliest archaeological site in a region has been identified and dated.

The earliest well-dated archaeological site in the Bahamian archipelago is the Coralie site on Grand Turk. At this site, short-term occupations spanning 700-1100 CE are associated with seasonal procurement of tortoises (now extinct), iguanas, fish, birds, and sea turtles (Carlson 1999). All of the pottery is Ostionoid-based, indicating that it was brought to this location from Hispaniola. As such, it does not necessarily reflect the human colonization of these islands. Nevertheless, proximity to Hispaniola (circa 150 kilometers), the Ostionoid and Meillacoid frontier model (Rouse 1986), settlement patterns, and the civic-ceremonial site on Middle Caicos (MC-6) have been interpreted as supporting evidence for the conclusion that the earliest inhabitants of the archipelago arrived from Hispaniola (Sears and Sullivan 1978; Keegan 1992). The recent study of facial morphology provides additional support for this conclusion (Ross et al. 2020).

Historic toponyms (Granberry 1991), a few early radiocarbon dates in the 700-900 CE range from the central Bahamas (i.e., San Salvador, New Providence, and Eleuthera), and certain pottery characteristics were used to propose that Cuba was the original source of migrants to The Bahamas

(Berman and Gnivecki 1995; Sears and Sullivan 1978). This Cuban connection was based on the assumption that the Ceramic Age reached Cuba by 700 CE. Local, “pre-Arawak” pottery was made in Cuba for at least 2000 years (Ulloa Hung and Valcárcel Rojas 2002), but Ceramic Age pottery did not arrive until after 900 CE (Persons 2013; Valcárcel Rojas 2002). Thus, if people from Cuba colonized The Bahamas, the colonists were likely of Archaic ancestry. The data presented in this paper do not support a biological connection between Ceramic Age peoples from The Bahamas and Archaic inhabitants of Cuba.

*Chicoid (1200-1450 CE)*

Around 1200 CE, a new pottery series emerged in eastern Dominican Republic. Chicoid (Chicoid is named for the Boca Chica site in the southeastern Dominican Republic) series pottery has complicated vessel shapes with fine, hard, and smoothed surfaces. Elaborate motifs including continuous scrolls, lines ending in dots, flat and prismatic lugs, and modelled anthropomorphic (“bat face”) lugs facing each other above the rim. The pottery is completely different from Meillacoid and Ostionoid and appears to be a local development. Chicoid pottery is associated with the contact-era “Classic Taínos” (Rouse 1992; Keegan et al. 2013). It has been found at archaeological sites across the entire island of Hispaniola. In Rouse’s (1992) classification it completely replaced Meillacoid pottery on the island, but new evidence suggests that both were in use until European contact (Keegan and Hofman 2017). To the east, Chicoid pottery occurs at outposts in Puerto Rico and in the Lesser Antilles (Hoogland and Hofman 1999; Rouse 1992). To date, it has not been identified in Jamaica, its motifs were copied by local artists in Cuba, and tradewares of this style are found on some islands, including The Bahamas, although the nature of this contact is poorly understood.

**Historic Age (1492 CE)**

It has sometimes been assumed that European colonization led to the rapid and total disappearance of Indigenous Caribbean peoples. However, the survival and continuing legacy of this group is widely recognized today, in part through the study of genetic ancestry (Martínez-Cruzado 2010, 2013, Schroeder et al. 2018, Nieves-Colón et al. 2020). Newly generated data in this study confirm the persistence of pre-contact Indigenous ancestry in present-day admixed people of the Caribbean.

**Northern South America and the Arawak Diaspora**

The archaeological record of South America was originally described in much the same way as that described for the Caribbean Islands. An initial “Paleo-Indian Period” (Lithic Age) was followed by a “Meso-Indian Period” (Archaic Age), then a “Neo-Indian Period” (Ceramic Age), and finally the “Historic Period” (Willey 1971). The coastal zone, stretching from western Venezuela to Brazil, was initially occupied by autonomous communities whose economy was based on gathering plants, fishing, and collecting marine mollusks for thousands of years. Over time, some adopted the use of pottery to a limited degree, and some may have practiced small-scale incipient farming. Cultural practices reflected local conditions, and major cultural changes were not apparent until “Tropical Forest” communities began to expand along interior river valleys and out onto the coast. The degree to which these Indigenous communities were displaced or assimilated is a question of local history that remains unclear.

Tropical Forest societies began to develop sometime before 2000 BCE near the confluence of the Amazon, Madeira, and Negro Rivers (Lathrap 1970). Their economy was based on the cultivation of bitter manioc, which fueled a rapid increase in population that led to expansion along the major rivers' drainages and then out to the coast. An abundance of ceramics is associated with the development of these societies. Lathrap (1970) proposed a single ceramic homeland, which affiliates all of the subsequent changes in ceramic styles to common ancestry (Willey 1971). A significant boundary in the ceramic series of the Northwest quadrant of Amazonia occurs at the middle Orinoco (Rouse and Cruxent 1963). To the west, Tocuyanoid is the earliest series, and forms the foundation on which later ceramic series developed.

The Santa Ana style identified at the Las Locas site (for which ancient DNA data are reported here) is an early expression of expansion into the Quíbor Valley (around 500 BCE), which appears to be related to the Dabajuroid series expansion that spread north to the central coast and then west to Lake Maracaibo and out to the islands of Aruba, Bonaire, Curaçao and the Las Aves Archipelagos (Haviser 1987; Oliver 1997; Antczak and Antczak 2015). The Dabajuroid expansion is juxtaposed to the Barrancoid expansion to the east, although whether or not they share genetic ancestry has not been addressed previously. The Dabajuroid series and the Tierroid series (Lara) are associated with the historical Caquetio, a people large in number and widespread as far south as the Barinas plains and Rio Cojedes (Arcaya 1916; Oliver 1989). The Caquetio lived along rivers and streams and on the plains where suitable agricultural soils were available. They had a versatile diet, including agricultural production, hunting of game, and exploiting marine resources such as reef fish and shellfish. When living in locations without access to maritime food sources, they would trade crops and game for fish with other peoples (see Federmann 1557). Caquetio settlements on Curaçao are located on low hills at about 500 meters distance from a bay with sea access, close to soils suitable for agriculture and gullies where rainwater comes together after heavy rainfall (Haviser 1987). The Caquetio were polygamous, or at least their caciques were (Oliver 1989:280, Federmann 1557). Federmann (1557) describes a Caquetio household as consisting of a man with his wife and children. A daughter of cacique Manaure went to live with her cacique husband after marriage, suggesting patrilocality (Oliver 1989:281).

To the east, linguistic evidence suggests that around 1000 BCE, Arawak communities began to move rapidly through the Negro and Orinoco floodplains and onto the Caribbean and Guiana coasts (Heckenberger 2002; Zucchi 2002). The center of dispersion is not certain, but appears to be the “ceramic homeland” in the Northwest Amazon (Lathrap 1970; Rouse 1992). The first wave of this expansion is characterized by the painted motifs of the Saladoid series, which spread downriver from the middle Orinoco to the eastern Caribbean coast of Venezuela, the Guianas, and then into the

Antilles. It was soon followed by a Barrancoid expansion into the same areas, but which differs from Saladoid in its elaborate incised motifs and relative absence of painting (Lathrap 1970).

The early Arawak expansion, including the colonists who migrated into the Antilles, shared a system of meaning and continuity in the broader cultural pattern. Summarizing Heckenberger (2002), Arawak speakers reproduce a *habitus* predisposed to perpetuate an ethos of settled village life, commonly coupled with large, fixed populations, fairly intensive subsistence economies, and landscape alteration (rather than mobility and low impact); institutional social ranking based on bloodline and birth order; and regional integration (particularly coupled with a social preoccupation with exchange and a cultural aesthetic that places great symbolic value upon foreign things) and a foreign policy commonly characterized by accommodation and acculturation of outsiders. This description is best suited to the Piapoco and their Arawak-speaking neighbors who share many additional cultural similarities with Indigenous Caribbean communities.

**Modern Arawak Language and Culture**

Anthropologists have long recognized that there is no necessary correlation between ancestry and culture (Boas 1940). An excellent example is a study of material culture along 700 kilometers of coastal New Guinea encompassing 55 languages and belonging to at least eight major language families. This study revealed that similarities and differences in material culture were better predicted by geographical than by linguistic affiliation (Welsch et al. 1992). Indigenous societies in South America exhibit similar linguistic diversity, with at least three major language families (i.e., Tupian, Cariban, and Arawakan) and hundreds of individual languages. The diversity of languages many have developed among autonomous lineage-based villages when population densities were low (Steward and Faron 1959). As groups associated with the different language families expanded, it has been hypothesized that they produced “multiethnic and multilingual regional sociopolitical systems” (Heckenberger and Neves 2009). A similar process could explain why in Papua New Guinea groups more closely resembled their neighbors who spoke different languages than they resemble more remote speakers of the same language. Moreover, the high degree of mobility and exogamous marriage contributed to widely shared cultural practices (Lowie 1948a). The practice of lineage and clan exogamy contributed to biological connectivity between communities, which suggests that neighbors should exhibit a closer genetic relationship irrespective of the language that they speak.

Language families continue to be used as an organizing principle for the recognition of ethnic communities, including in the Caribbean (e.g., “Arawak speakers”), and shared languages do establish cultural linkages between groups of people and raise the possibility of substantial amounts of shared ancestry as language shifts in pre-state societies are usually propelled by movements of people (Bellwood 2001). In light of examples like the study from New Guinea, however, it is incumbent on investigators to demonstrate empirically the degree to which genetic, linguistic, and material culture classifications are correlated. Confusion can also arise from the use of similar names in different taxonomies and from changes or differences in interpretation. For example, it was initially assumed that the "Caribs" or "Island Caribs" of the Lesser Antilles (today called Kalinago) spoke a Cariban language (reviewed in Rouse 1992:21-22), but detailed linguistic analysis has demonstrated that they actually spoke an Arawak-based language (Granberry and Vescelius 2004). As Salzano and colleagues caution (2005:S126): "Analogies between linguistic and genetic variability should be performed with caution."

In light of the fact that it cannot be assumed that shared language necessarily expresses a close genetic relationship, we considered the results of the present study which show that by ADMIXTURE, Treemix and *qpAdm* analyses (albeit not replicated with $f_4$-statistics), the main group of individuals from the Ceramic Age in the ancient Caribbean showed greatest affinity with present-day Indigenous South Americans who speak Arawak languages (Fig. 3). This finding suggests that the spread of the ceramic-using genetic cluster was carried out at least in part by Arawak speakers, as Arawak languages were widely spoken in the pre-contact Caribbean. Arawak languages are known to be concentrated in populations to the north of the Amazon River, while Tupi speakers are concentrated in the south, and Carib-speaking groups cluster along the coast (Salzano et al. 2005; Walker et al. 2012).

In northern South America, Arawak-speaking Indigenous groups tend to focus on controlling riverine habitats distinguishing them from their interfluvial upland neighbors who spoke Cariban and other languages (Steward and Faron 1959). The largest number of Arawak languages are recorded in the Northwest Amazon, which led to the conclusion that this was likely the Arawak homeland, and would be consistent with the interpretation that the Arawak-speakers expanded down the Amazon and Orinoco Rivers (Heckenberger 2002). However, it also is possible that the demographic catastrophe that resulted from European conquests led to migrations upriver to the point where further westward progress was impeded by the Andean foothills and the absence of navigable rivers, and thus the resulting distribution of Indigenous languages today might be qualitatively different from the pre-contact one (Lathrap 1970).

Today, some Arawak-speakers live in close geographical proximity to areas from which the initial Ceramic Age colonists of the Caribbean Islands likely originated more than two millennia ago. Some also live in western Venezuela, which is significant because it is where V-shaped incised and appliqué pottery motifs from the west (Araquinoid and Dabajuroid series) abut U-shaped incised and painted motifs (Barrancoid and Saladoid series, respectively) from the east (Zucchi 2002). Dabajuroid series pottery is relevant to this study as it is the only ceramic tradition identified for Curaçao, where we infer about two-thirds ancestry from the main Caribbean Ceramic genetic cluster, and Meillacoid pottery from the western Greater Antilles and The Bahamas has been affiliated to V-shaped incised and appliqué pottery traditions (Zucchi 1985; Ross et al. 2020). In what follows, we specifically discuss similarities between material culture of Caribbean islanders as inferred from archaeological evidence, and ethnographic information for the two genotyped Arawak-speaking groups to which they show strongest affinity according to Treemix and *qpAdm* models (Supplementary Information sections 7 and 8).

*Piapoco*

Ethnographic research indicates that the Piapoco occupied the middle course of the Guaviare River, María River, and Cuinada River in the vicinity of Lake Maracaibo in western Venezuela and eastern Colombia. Gregorio Hernandez de Alba (1948) describes cultural practices in this region with a particular focus on the Achagua, and includes the Piapoco in this description. Most of these features also are found in the Caribbean Islands (Keegan and Hofman 2017; Rouse 1992). The communities in this region were palisaded villages built to protect the inhabitants from raids by Carib communities of the interior uplands. The palisade contained large communal dwellings (up to 500 people) as well as a special men's house. Dugout canoes were an important mode of transportation. Gardens were prepared by men and cultivated by women. Maize (*Zea mays*) along with sweet and bitter manioc (*Manihot esculenta*) were the most important cultigens. Pottery was used for food preparation and storage, and calabashes for water. They produced baskets and woven matts, made cotton fishing nets and hammocks, and short women's skirts were the only article of clothing. Objects of wood included stools and large hollow-log drums. The most important ornaments were shell beads, which on occasion were sacrificed to mark successful fishing (see Carlson 1993 for a similar practice at GT-2 on Grand Turk, Turks and Caicos Islands). They painted their faces for luck in hunting.

Social organization is characterized as patrilineal sibs (i.e., clans), and individuals "had to go to distant villages to marry" (Hernandez de Alba 1948: 404). Polygyny was practiced, and every village had a chief. They fought with neighboring Caribs and the bow and arrow and a war club (*macana*) were the main weapons. Shaman were healers who extracted an object from the patient, and were intermediaries with the spirit world. They inhaled a powdered drug, using crossed bird bones, to facilitate divination. The narcotic is identified as *Piptadenia* sp., which was called *cohoba* in the Greater Antilles. Of note is that the use of *Piptadenia* may have been restricted to Western and Northwest Amazonia (Cooper 1948:537). Frogs symbolized the "lords of the water" (Hernandez de Alba 1948: 410). They did not produce idols, but instead represented the spirit world with masks.

*Palikur*

The Arawak-speaking Palikur, or Pa'ikwené, inhabit the northeast coast along the border between Brazilian and French Guiana, along the Oyapack and Vaçá River drainage. They express a hereditary enmity with the neighboring Carib-speaking Galibi, although the Carib and Arawak tribes of the Guianas share many features (Gillin 1948). Their historic homeland of Amapá, between the Amazon and Oyapock Rivers, was in the past characterized by "a profusion of diverse of ethnic groups, clans, and languages (Arawak, Carib, Tupi) out of which there seems to have developed a unified (though not homogeneous) culture, entailing peaceful and interdependent relations and interethnic trade, festivities, and marriage (Passes 2002:178). Their territory is swampland with small clusters of beehive huts (with walls and roof merged) constructed on forested islands. Movement between these clusters is hampered by swampy conditions, especially during the midsummer rainy season, so log causeways are built across the muddy terrain (Lowie 1948a). They make a crude pottery that is less refined than the urns they once made. These urns were used for burials, and each moiety had its own cemetery. They are known for the realistic representation of turtles on clay vessels. Their material culture is not distinctive, but they share the use of wooden shields and fall traps for capturing game with the Arawak-speaking Island Caribs (Cooper 1948; Métraux 1948a). They also used the bow and arrow. Their sides of their canoes may have been built up with planks. They were propelled with poles, or paddled with "crutch handle" elongated leaf-shaped blade paddles (Gillin 1948:837-838).

Each of the original 18 clans had their own territory and spoke their own language. The remaining seven or eight Pa'ikwené clans are the product of accretion and consolidation. A process that began in the 17$^{th}$ century with elements of depleted groups progressively absorbed into Pa'ikwené clans, including the Carib-speaking Paragoto (Passes 2002). Every village had a headman or "chief" whose authority was based on consensus. His main task was greeting visitors and representing the community. Social organization is characterized as moieties divided into seven patrilineal clans. However, it is possible that they originally were matrilineal and that current practices developed in relation to neighboring groups. Marriage distances are short, women exert considerable authority (even with regard to a husband's personal possessions; Lowie 1948c:356), and residence is described as both uxorilocal (residing with the wife's family) and ambilocal (residing with the parents who offer the best accommodation). Their moieties are agamic, meaning there are no restrictions on inter-clan marriage. Like the Carib, premarital license is accepted, although the Palikur are more strictly monogamous than neighboring tribes. Boys experienced scarification and flagellation during puberty rites, after which they wore cotton bands on the arms and legs (Metraux 1948b:377). The later practice is described for the Island Carib. Shamans were responsible for healing the sick, and could serve as "master of ceremonies" during feasts (Lowie 1948a). Like the Island Carib they celebrated the recovery of the sick person with a feast; and like the Taíno, made food offerings to the spirits (Métraux 1948c:578). There is no mention of drug use other than tobacco.

While the convergence of genetics, linguistics, archaeology and ethnography connecting Arawak-speakers of the Caribbean islands to Arawak-speakers in northern South America is striking, our $f_4$-statistic analysis failed to replicate a specific association to Arawak-speakers. However, support for that inference comes from ADMIXTURE, Treemix, and $f_4$-statistics-based *qpAdm*. It is possible that the inconsistent genetic signal reflects admixture and complexity in the history both of present-day Arawak speakers on the South American mainland, and in the people who settled the Caribbean during the Ceramic Age. Thus, there is also some evidence in our data for there not being a one-to-one relationship between genetics and archaeological style.

**SI2- Archaeological site information for newly reported individuals**

**BAHAMAS:**

The Bahama Archipelago (comprising the Commonwealth of The Bahamas and the Turks and Caicos Islands, a British Overseas Territory) is located in the Atlantic Ocean extending south from Florida to Hispaniola (over 1,000 kilometers). It consists of over 700 islands and cays. The archipelago comprises a land area of approximately 14,000 kilometers$^2$ laid out in a northwest to southeast direction. The elevation throughout the archipelago is less than 60 meters above sea level and, in most places, no more than 20 meters above sea levels. The islands are the result of coral reefs, which became dry land when the sea level dropped hundreds of centuries ago. The islands are mostly flat with miles of white and pink sandy beaches. Solution features in the limestone have created a karstic landscape with sharp pinnacles, crevices, caves, and sinkholes. Additionally, there are no freshwater rivers located in any of the islands of the archipelago (Sealey 1995).

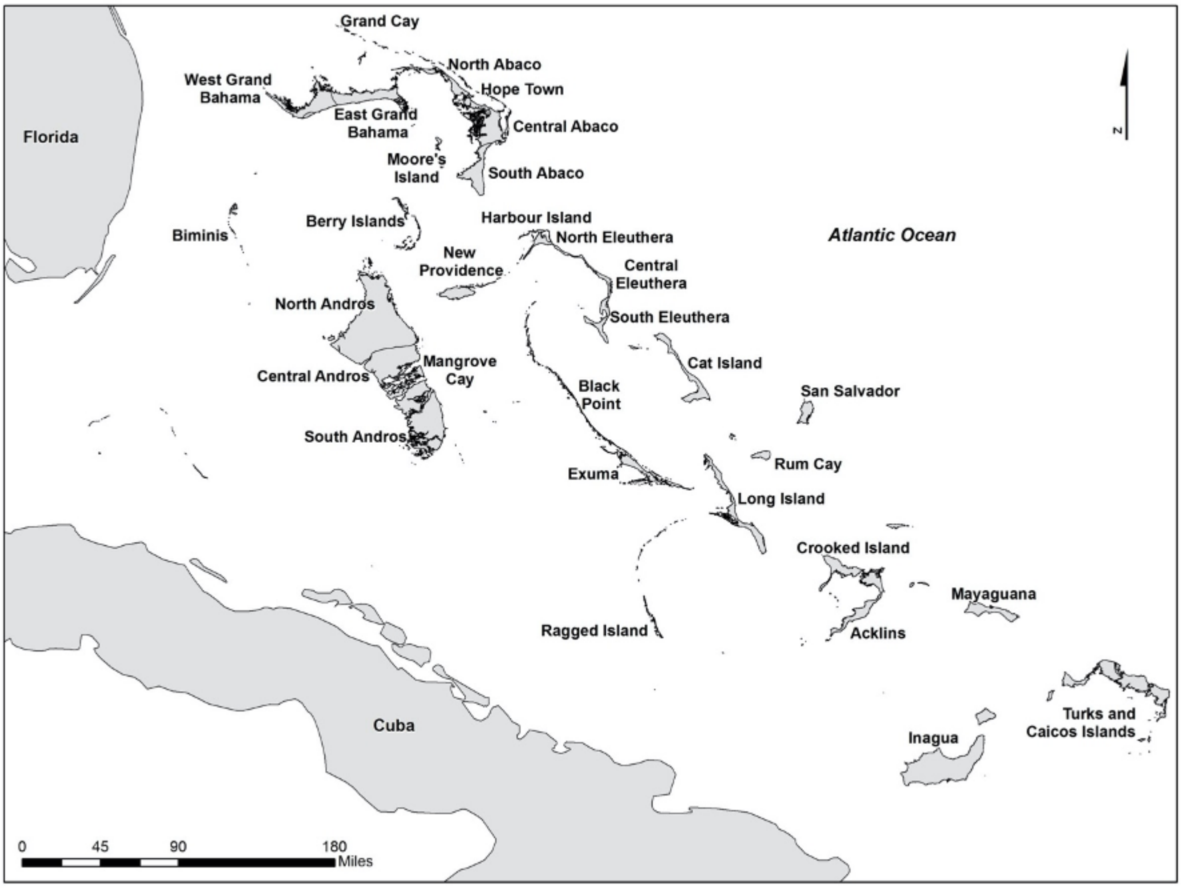

**Figure S1:** Map of the Bahama Archipelago.

The first skeletal research in The Bahamas was conducted by Brooks (1888), describing three crania found in caves from throughout the islands. Caves played a vital role in Taíno lifestyle and spiritual beliefs, and as such, it played an essential role in that of the Lucayans. Therefore, it should be no surprise that caves represent a significant aspect of the archaeological record of The Bahamas. These caves exist in two forms, wet (including blue holes and caves with a direct connection to the water table) and dry. These caves contain a variety of artifacts which have not been preserved at open sites such as human burials, petroglyphs and pictographs, faunal and botanical remains, and a variety of wooden artifacts (examples in De Booy 1913; Rainey 1934; Hoffman 1973; Keegan 1982; Pateman 2007; Pateman and Keegan 2019). The majority of Lucayan burials are known from caves from throughout the archipelago.

The human remains included in this study were curated at either the Yale Peabody Museum of Natural History (Peabody Museum) or at the National Museum of The Bahamas. All of the human remains in the Peabody Museum from The Bahamas used in this study were collected in 1934 by Froelich G. Rainey. In January 1934, Rainey visited The Bahamas to locate and excavate archaeological sites and examine several cultural materials found by locals on many islands (Rainey 1934). Rainey recovered skeletons from thirteen dry cave sites from the Abaco Islands, Eleuthera, San Salvador, Rum Cay, Long Island, and Crooked Island (Rainey 1934). Although Rainey referred to the survey and excavations in The Bahamas in his field diary, he did not write a report about the excavations or provide detailed field notes of the recoveries (Granberry 1978) and therefore their context and antiquity of the remains are not well documented. Some of the locations where human remains were recovered are now lost to science. Several of these remains were found by cave earth diggers and thrown on the outside of the caves. Similarly, numerous human skeletal remains have been recovered in wet caves by cave divers and explorers with little regard to proper archaeological methods and protocols (Pateman and Keegan 2019). Previous skeletal studies have shown a population high in dental disease as well as multiple traumas, either the result of accidents or violence (Pateman 2007; Schaffer 2015) and a relatively young population at death (Pateman and Keegan 2019). The Bahamian human remains from the Peabody Museum have been the subject of several publications and are most systematically described by Schaffer (2015).

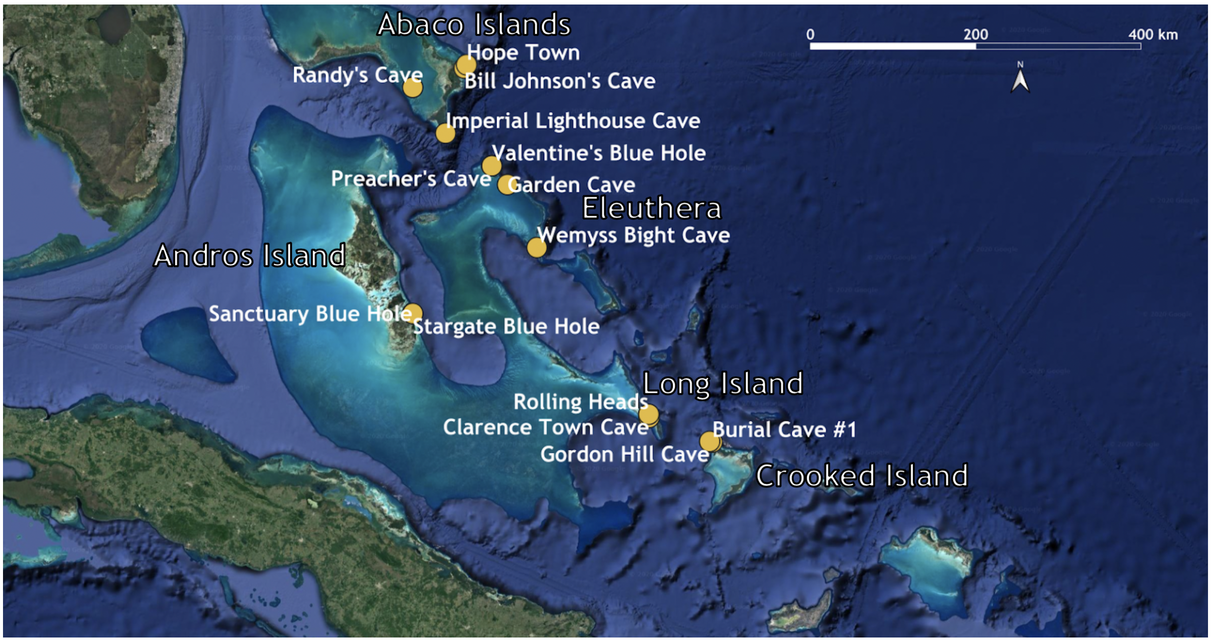

**Figure S2:** Location of 14 archaeological sites in The Bahamas included in this study. Map created using QGIS Geographic Information System v3.6 (http://qgis.org); basemaps from Google Earth.

##### The Abaco Islands

The Abaco Islands lie in the northern Bahamas and include the main islands of Great Abaco and Little Abaco along with numerous cays and consist mainly of tropical marine wet and dry climatic zones (Sealey 1995). Walker's Cay of the Abacos is the northernmost point of the Bahama archipelago. The combined landmass of the Abacos is approximately 2,009 kilometers$^2$, and Caribbean pine (*Pinus caribaea*) dominates the modern vegetation. Research into the paleo-environment showed that the landscape consisted of lush tropical forests (Steadman et al. 2007). Hurricane Dorian (2019) razed the Abaco Islands, flooding inundating the main island and irrevocably changing the coastline due to erosion. There are four burial locations in the Abacos with four individuals as part of this study.

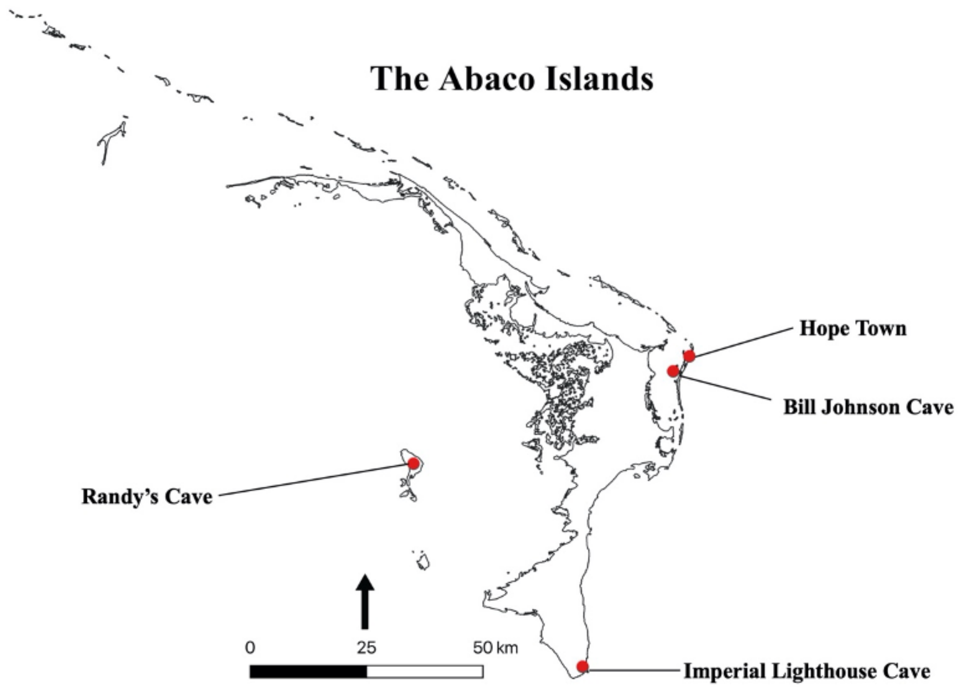

**Figure S3:** Map of the Abaco Islands with burial locations involved in the study

**Bill Johnson's Cave, Lubber's Quarters**

Bill Johnson's Cave (AB-10) is on the southeast side of a cay known as Lubbers Quarters, which is situated southwest of Elbow Cay, facing Tilloo Cut to the east. It is 10 meters due north of the house owned by the property owner, Bill Johnson. This flank margin cave is a large single chamber situated in the south flank of a sheer cliff face with a narrow single cave passage that penetrates the cliff approximately 15 meters into the hillside. The cave's entrance is about 25 meters west of the shoreline and 4 meters above sea level. The entrance chamber is 3 meters high and runs 10 meters along the cliff face. The cave entrance is partly filled with yard debris on the west side of the chamber, but otherwise there is almost no sediment in the interior passages. Other than a small section of flowstone, there is no evidence of bats or guano. Numerous broken conch shells, many with small round holes (Lucayan-modified), are scattered downslope of the cave's entrance chamber. Approximately 3 meters inside the eastern dripline, Nancy Albury collected a small sample from dry sediment, including an iguana (*Cyclura* sp.) jawbone, bone fragments from fish, bat, tortoise, and human bone fragments.

**Hope Town**

There is not much known about the provenience of this individual. Found by residents in Hope Town, Elbow Cay, The Abacos during excavation for a water cistern, this individual was recovered by amateur archaeologists in 1990. The remains from Hope Town, represent the first, non-cave burial found in The Bahamas (as noted above, the majority of Lucayan burials are in caves from throughout the archipelago). The remains consist of a nearly complete individual female, age 21 to 25. There were associated artifacts found with the burial, but it is challenging to say if it is a funerary offering or a separate deposit. Objects include a large limestone fragment (possible pounding stone), five Palmetto Ware sherds, one worked conch tool, parrotfish bones, fire-cracked rock, turtle bones, and various shell fragments. As an archaeologist did not excavate this, reports are lacking.

**Imperial Lighthouse Cave**

While Rainey was in Abaco (1934), he investigated an ocean sinkhole on the southern point of Abaco approximately 1.6 kilometers from the Hole in the Wall Imperial Lighthouse. There he recovered cranial and postcranial bones of a child, aged 5 to 10, along with five undecorated Meillacoid sherds, fish bones, conch shells charcoal, bird bones, and hutia bones (Keegan 1982). However, Rainey's notes do not provide further information.

**Randy's Cave**

Randy's Blue Hole (AB-55) is located on Moore's Island, south of Great Abaco Island, approximately 550 meters northeast of the nearest mangrove shoreline. The small blue hole is an eroded subsurface feature with geologic characteristics of a flooded vertical pit cave, having a subaerial diameter of three to five meters. Surface water lies three meters below ground level with a small subsurface chamber extending laterally 15 meters to a depth of eight meters. Brian Kakuk and Kenneth Broad conducted a dive in July 2007. They discovered a partial human skull of a Lucayan, one human molar, a hutia sacrum, and of tortoise bone fragments. Subsequent trips by Albury noted that bulldozers infilled the blue hole during the clearing and construction of a nearby athletic field.

**Andros Island**

Although Andros physically comprises multiple islands, politically, it is considered one island. Andros is located in the central Bahamas and consists of the main islands of North Andros, Mangrove Cay, and South Andros, along with numerous cays. Andros' three main islands are separated by trifurcated estuaries, connecting the island's east and west coasts. Collectively the fifth largest in the West Indies, the islands are 6,000 kilometers$^2$, 167 kilometers long, and 64 kilometers at the widest point. Andros has a rare combination of marine features, ecosystems, and the world's largest collection of blue holes. It is flanked on the east by the Tongue of the Ocean, a 6,600-foot deep trench. The world's sixth longest atoll, The Andros Barrier Reef, runs for 225 kilometers, and to the west, northwest and south lies the Great Bahama Bank. Paleoclimate and vegetation studies show that during the late Holocene the environment supported a dry shrubland environment (Kjellmark 1996). Andros' higher levels of rainfall, cooler temperatures, and pine forests are very different from the other islands of the central Bahamas. Andros exhibits greater faunal and flora biodiversity than any other Bahamian island (Campbell 1978). The island's land ecosystem is hugely varied, including pineyard, scrub, hardwood coppice, saltwater marsh, rocky and sandy beaches, palm savannas, and mangroves (Randolph 1994). There are two burial locations in Andros with eight individuals as part of this study.

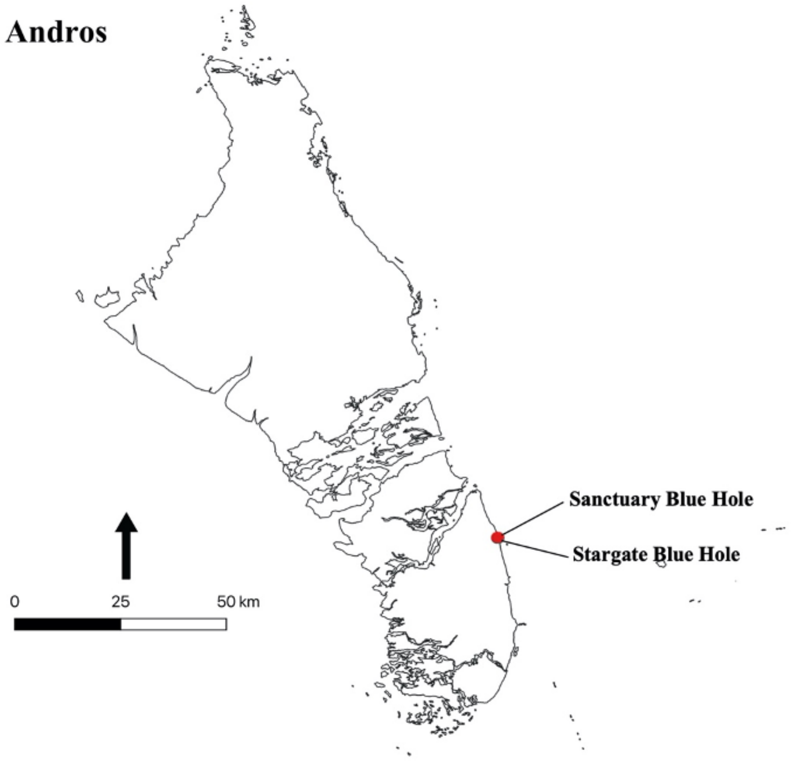

**Figure S4:** Map of Andros with burial locations involved in the study

**Sanctuary Blue Hole**

Sanctuary Blue Hole (AN-12) is about 500 meters inland from the east coast of South Andros Island on the eastern side of the settlement of The Bluffs village. It is part of a significant north-south slump fracture zone paralleling the underwater escarpment separating the Tongue of the Ocean and the Great Bahama Bank. This slump fracture extends for tens of kilometers and formed as a result of glacio-eustatic sea-level changes and gravitational forces along the edge of the limestone banks (Palmer 1986a, b). The entrance consists of a collapse-floored fissure extending down beneath a bedrock ledge to a pool about 5 meters long by 2 meters wide. A rift drops vertically beneath the water surface and opens out at a depth of 25 meters into a significant fissure passage that extends in both directions. Water depths in the cave do not exceed 60 meters. Between 1990 and 1991, cave diver Rob Palmer recovered 17 sets of skeletal remains. In 2009, as part of a National Geographic expedition, another individual was retrieved (Pateman and Keegan 2019). Osteological studies reveal a population that was short in stature and relatively active, probably due to subsistence activities. Indicators of health show that they had a relatively good childhood and adult health, although it appears there may have been some dietary stress (Pateman 2007). Existing radiocarbon data show a site range between 1200 to 1400 CE (Pateman and Keegan 2019).

**Stargate Blue Hole**

Stargate Blue Hole (AN-13) is about 500 meters inland from the east coast of South Andros Island, on the western side of the settlement of the Bluffs village. It is a part of the same system. The cave's entrance is a partially roofed-over cavern with a vertical drop of 6 meters to the water level. The restricted nature of the access limits organic input, and as such, the surface water is relatively clear. Underwater, a shaft drops vertically to depths over 80 meters, while rift-like passages extend north and south. A gossamer layer of fine, brown sediment covers breakdown blocks on the floor of the cave; speleothems are present at all depths (Palmer 1986a, b). Divers recovered a ceremonial canoe from a ledge at 20 meters (the canoe is less than 2 meters long). The shelf also contained human remains, but these were not retrieved. During a National Geographic expedition in 2009, divers encountered two additional sets of human remains, removing only one. These remains date to 1210 to 1390 CE (Hastings et al. 2014).

**Crooked Island**

Geographically positioned southeast of Long Island, Crooked Island is 238 kilometers$^2$ and sits in the central region of a tropical moist forest zone (Stokes 1998). It is both an island and a district, the largest of a group comprising a large shallow lagoon known as the Bight of Acklins. Crooked Island is a part of The Bahamas' cotton belt, settled by Loyalists in 1780, and it also suffered extreme deforestation by plantation economies that collapsed in 1825 (Craton and Saunders 1992). In 2015 Crooked Island was catastrophically damaged when hurricane Joaquin's eyewall passed over.

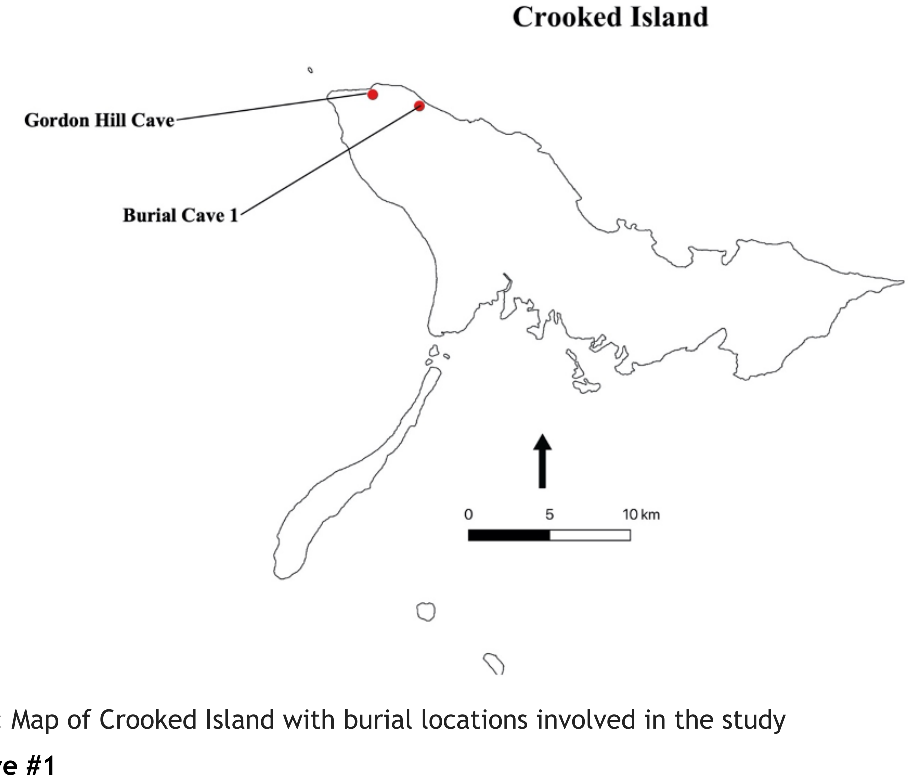

**Figure S5:** Map of Crooked Island with burial locations involved in the study

**Burial Cave #1**

Rainey (1934) recovered the remains of four individuals that had been excavated by cave diggers. The exact location of the site is unknown but is on the northern end of the island (Rainey 1934, Stokes 1998). It may be part of the cave system (Crossbed, 1702, and Owl Roost caves) recently investigated by David Steadman and Nancy Albury who recovered crocodile, hutia, and tortoise bones but do not mention human or other cultural remains (Steadman et al. 2017). The animal bones from these caves are dated between 1450-1620 CE. Features of four individuals were recovered, one male, one female, and two undetermined, all between the ages of 17 and 25 (Keegan 1982; Stokes 1998).

**Gordon Hill Cave**

According to Rainey's diary (1934), the only site he systematically excavated was Gordon Hill Cave. Gordon Hill Caves are located on the northwest end of the island about 450 meters from the coast, among a series of caves along a limestone bluff. Gordon Hill Caves represent Rainey's most productive excavations, both culturally and osteologically (Granberry 1978). Seven caves were excavated, four of which were previously partially excavated for cave earth. Two chambers were excavated in the Gordon Hill Burial Cave, each containing a single burial. One burial was very fragmentary, lying on its left side with the legs partly flexed on a small rock shelf on the cave floor just beneath the surface with finger bones covering the pubic region. A second disturbed burial was found in the south side of the cave, where it sloped up to the entrance. This burial also lay on the rock floor of the cave. In Chamber one, near the burial were the bones of hutia and various birds. Another cave, described as the residential cave, was extensively excavated with a variety of artifacts discovered, and had evidence of multiple fires. The material recovered was classified as Meillac-like (Meillacoid) with Carrier-Like (Chicoid) traits (Granberry 1955), pottery styles found as trade ware in The Bahamas.

**Long Island**

Long Island, located in the central Bahamas, is approximately 130 kilometers long and 6 kilometers wide with a landmass of 596 kilometers$^2$. The Tropic of Cancer splits the island, which is acclaimed for its caves and surrounded by smaller islands, bays, and inlets. Its orientation, geology, topography, and climate share commonalities with the other central islands. Studies of the northern Bahamas reveal that "a late Holocene dry period altered the limnology and supported only dry shrubland" (Kjellmark 1996). The island experienced catastrophic deforestation during the Loyalist Period by land clearance of cotton plantations and the harvesting of valuable hardwoods between 1783 and 1788 (Craton and Saunders 1992). Long Island's terrain now varies widely, including white salt flats (salina), swamplands, beaches, and northern sloping and southern low hills. Hurricanes Joaquin (2015) and Dorian (2019) severely damaged the island's coastline.

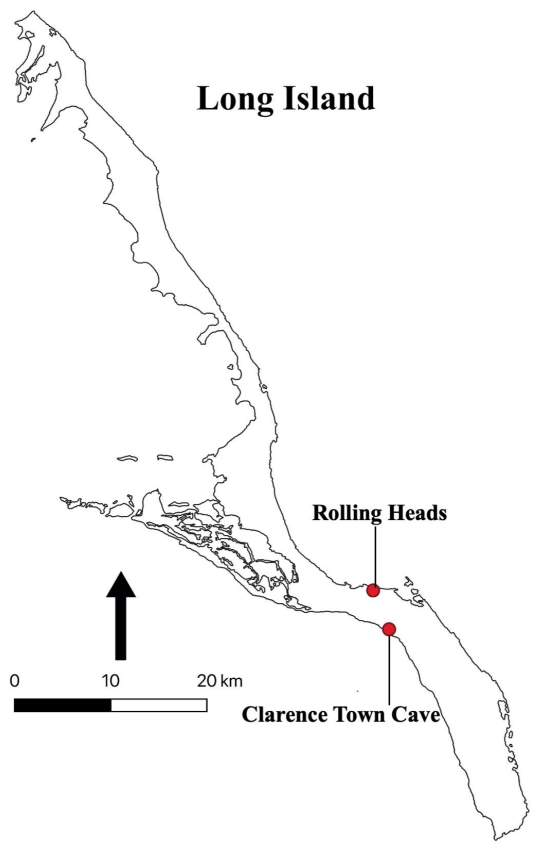

**Figure S6:** Map of Long Island with burial locations involved in the study

**Clarence Town Cave**

In 1934 Rainey located and excavated one cave on southern Long Island near Clarence Town (Stokes 1998). The Clarence Town cave contained a main chamber with smaller branched compartments that indicated evidence of guano excavation. The material culture found in chamber one included skeletal remains and an undescribed piece of pottery. Rainey assumed the remains were one individual, but Keegan (1982) subsequently determined the skeletal remains were those of two individuals. The first was an under 14-year-old juvenile represented by a humerus, radius, small temporal bone, and section of the ilium. The second is most likely a female between 20 and 30 years old evidenced by the maxillary, a third molar, and part of the ilium. The adult female from chamber one was dated to 1175-1295 calCE (Stokes 1988).

**Rolling Heads**

After Hurricane Joaquin devastated Long Island in September 2015, residents Nick Constantakis, Nick Maillis, and Anthony Maillis found two fronto-occipital modified Lucayan skulls on Lowe's Beach. They identified two places exposing human bones in the dune face. Between October 17-19, 2016, Pateman and Keegan excavated three sets of remains from what is the first multiple Lucayan burials found outside of a cave environment by archaeologists. The graves had no associated cultural material (Pateman and Keegan, 2019). Further testing revealed two seasonal occupation areas with associated earth ovens located 80 meters east of burial 1 (LN101A) and 20 meters west of burial 2 and 3 (LN-

101B) (LeFebvre et al. 2019). The burials are dated between 1075-1405 calCE without marine diet correction.

**Eleuthera**

Eleuthera refers to the main island, its associated chain of smaller islands, including Harbour, Russel, Royal, and Windermere, and numerous cays. Eleuthera is located on the Great Bahama Bank in the central Bahamas. The main island is 180 kilometers long, little more than 1.6 kilometers at its narrowest, and has a landmass of 457.4 kilometers$^2$. The islands also were part of the Bahamas' cotton belt and faced extreme deforestation due to plantation economics. The hardwoods harvested during the Loyalist occupation included ten tons of Brasilwood sent as part of an endowment for Harvard University (Craton and Saunders 1992). Hurricane Andrew (1992) severely damaged The Eleutheras; the category five storm carried immense wind speeds, and an 18-foot tidal surge inundated the coastline. The islands' current topography differs extensively, ranging from pink sand beaches, while also including sizable ancient coral reefs, caves, and other geological features.

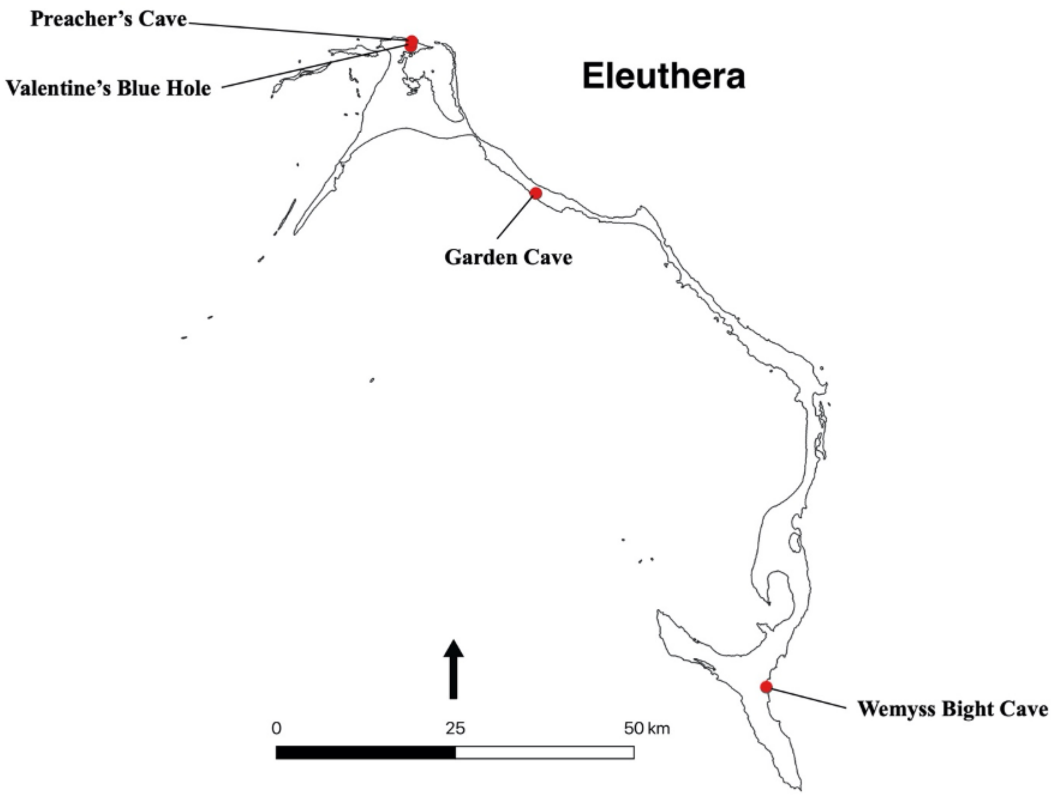

**Figure S7:** Map of Eleuthera Island with burial locations involved in the study

**Garden Cave**

Garden Cave (EL-229) is a shallow flank margin cave (see Mylroie and Mylroie 2013) developed in an inland but seaward-facing cliff in the region of Hatchet Bay, Eleuthera. The cave is a short distance from the more famous Hatchet Bay Caves. Various caves riddle the entire ridgeline in the area. In 2006, officials from the National Museum of The Bahamas were made aware of the site, because of reported looting, and recovered two exposed skulls. Local eyewitnesses reported that when local explorers first discovered the site, there were 12 skulls arranged with other human bones. In July 2017, Keegan, Pateman, and Maurice White excavated the remains of at least six individuals (Keegan et al. 2018). Paleontologists Nancy Albury and David Steadman also studied the cave. They conducted excavations in the cave to recover animal bones as part of a larger project to understand the indigenous fauna of The Bahamas (Steadman et al. 2017).

**Preacher's Cave**

The Preacher's Cave site is on the northern coast of Eleuthera adjacent to Jean's Bay. Preacher's Cave is characteristic of a sea cave, mostly horizontal and lacking speleothems. It received its name from the original Puritan settlers who shipwrecked there in 1648 (Craton and Saunders 1992). Recovered from within the cave were a total of seven Lucayan burials (Carr et al. 2006; Pateman and Keegan 2019). These burials are among the most complete archaeologically documented Lucayan burials in the Bahama archipelago. Two of the burials show evidence of binding with plaited matting, and one of the individuals has associated grave goods comprised of a triton (*Charonia variegata*) shell, 29 sunrise tellin (*Tellina radiata*) shells, red ochre, and a fishbone pin. Existing radiocarbon dates show a site range for the burials of ~800 to 1250 CE.

**Valentine's Blue Hole**

"Valentine's Cave" (EL-179; also known as "Bat Cave") is situated about 1.7 kilometers southwest of Preachers Cave and 900 meters south of the north shoreline of Eleuthera. The cave is named for Valentine Yacht Club on Harbour Island, whose members surveyed it many years ago and left lines and markers. Approximately 20 meters south of the road, an east-west fissure is open to a large subsurface flank margin cave. The small entrance (approximately 2 meters x .5 meters) is partially filled with a ficus tree root system and opens into a large dry cave chamber ~25 meters in diameter and 3 meters high. A shallow lake (approximately 3 meters depth) fills the back of the dry chamber and a small area of the central chamber. Sediments are almost absent from the dry portion of the cave, but black organic residue and guano are interspersed between rocks and boulders, covering the bottom of the flooded portion. A roosting colony of about 200 Buffy flower bats (*Erophylla sezekorni*) was noted in February 2018. The surface saltwater is crystal clear, crisp, without hydrogen sulfide. A flooded tunnel leads away from the main chamber to the southwest reaching a maximum penetration of approximately 200 meters and maintains a depth of 3 meters. In August 1993, cave divers removed human skulls and other bones from this site. Osteological analysis of these remains reveals at least five adult individuals, four males and one female (Keegan et al. 2018). Further investigations by Nancy Albury revealed several hutia bones in the shallow cave sediments and the shaft of a human bone, both within the flooded portion of the main chamber. Brian Kakuk collected these; the unassigned leg bones are the shaft (missing proximal and distal ends) of a juvenile.

**Wemyss Bight Cave**

Rainey located a cave two and a half miles inland from Wemyss Bite, southern Eleuthera. Upon discovering the site, excavation revealed a low cave that contained a surface burial. Upon inspection, it contained no artifacts, only scattered remains mixed with dirt and organic decay. A cranium bought from Wemyss settlement fits the mandible found during excavation (Keegan 1982). The skeletal remains represent at least two individuals. When Keegan (1982) analyzed the bones, they were determined to be male, but indistinguishable due to similarities in appearance and size. His count indicated two separate individuals represented by two left temporal bones, two right humeri, two right scapulae, one almost complete cranium and one left temporal, and one lumbar vertebra (Stokes 1998).

**CUBA:**

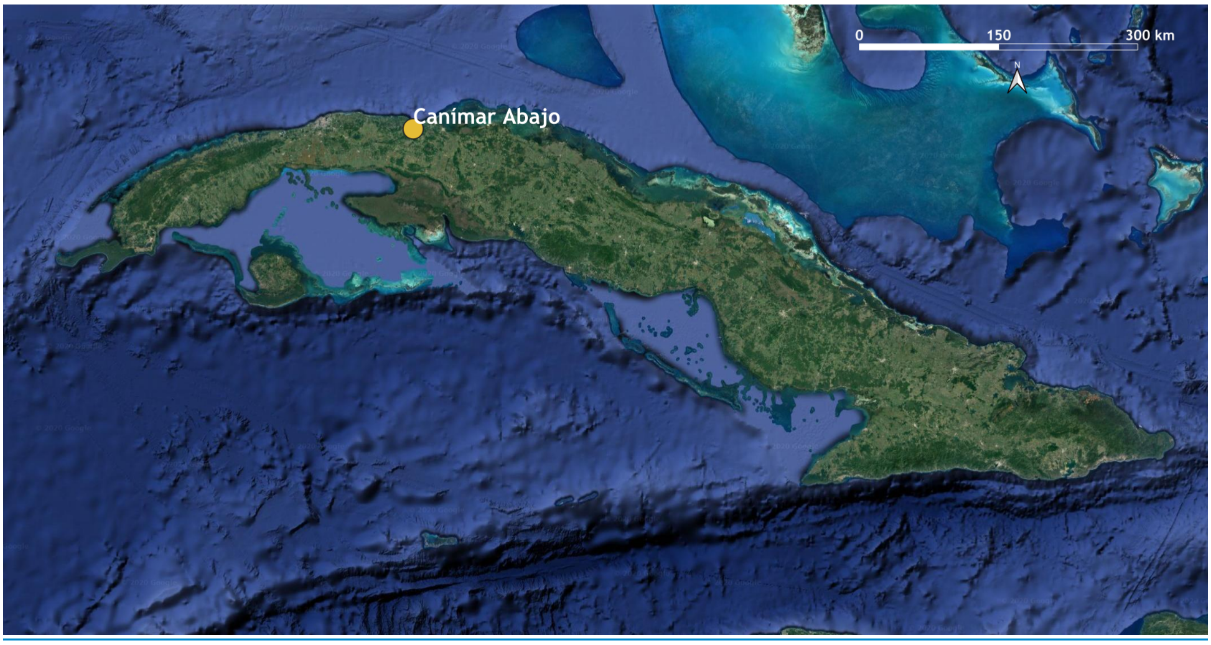

**Figure S8:** Location of archaeological site in Cuba included in this study. Map created using QGIS Geographic Information System v3.6 (http://qgis.org); basemaps from Google Earth.

**Canímar Abajo (Matanzas Province, Cuba)**

Canímar Abajo is an early pre-Columbian archaeological site situated on the south-western side of the Canímar River (Rodríguez Suárez et al. 2006). Primarily identified in the 1960s by tourists, Cuban archaeologists from the Montané Anthropological Museum and the Faculty of Biology of the University of Havana have investigated the site intermittently since 1984 (Martínez López et al. 2007, 2009). While most of the archaeological and anthropological findings are stored in the Montané museum, it is supposed that some anthropological remains have been lost to other institutions in the country based on unpublished site reports from Dr. Rivero de la Calle, one of the original excavators in the 1980s (Dumas León 2009).

Canímar Abajo, a protected site at the bottom of a limestone outcrop, is largely a funerary site, containing at least 199 burials in two cemeteries that are separated in time by approximately 1,500 years. The stratigraphy of the site highlighted three major elements labelled S1 to S3. S1 is a more recent burial area (Young Cemetery, YC) located at a depth spanning 0.60 meters to the surface level (360-950 calCE), while S3 is an older burial area (Old Cemetery, OC) located at a depth of 1.80-1.50 meters (1380-800 calBCE) (Martínez López et al. 2009; Morales Valdes 2009). Between these two burial levels is S2 (1.50-0.60 meters), which is composed of different settlement phases showing evidence of rituals or food-processing, with faunal remains and traces of burning, as well as a shell midden (Martínez López et al. 2009; Morales Valdes 2009). The site is roughly 40 meters from the Canímar River, and the surrounding vegetation consists mainly of bushes, mangroves, and, further off, semi-deciduous forest (Dumas León 2009). This abundant greenery, and the food sources accessible from the river itself, allow a variety of sustenance activities (Chinique de Armas et al. 2008, 2015), while the karst outcrop provided a protected rock shelter (Dumas León 2009) for the inhabitants of the area.

With a total of at least 213 individuals, including 83 adults and 130 sub-adults, Canímar Abajo is one of the largest pre-Columbian cemeteries excavated in Cuba. Since 2010, research has been conducted within a joint Cuban/Canadian project in collaboration with the University of Winnipeg, adopting a multidisciplinary approach (Chinique de Armas et al. 2015; Roksandic et al. 2015, 2016).

Radiocarbon dates from charcoal at the site have provided dates as early as 5590-4622 BP (Cooper 2010; Rodríguez Suárez et al. 2006), while more recently acquired dates from skeletal material indicate that the site spans at least 3000-1250 BP (Rodríguez Suárez et al. 2010; Roksandic et al. 2015). Although more dates are required, especially considering the difficulty of taphonomic interpretation of the site chronology due to disturbance and the reuse of burial space, these dates make Canímar Abajo one of the oldest sites in Cuba, particularly Western Cuba (Cooper 2010; Dumas León 2009; Martínez López et al. 2007, 2009; Rodríguez Suárez et al. 2006). In this study, we include samples from both the older (OC) and the more recent (YC) cemeteries.

**CURACAO:**

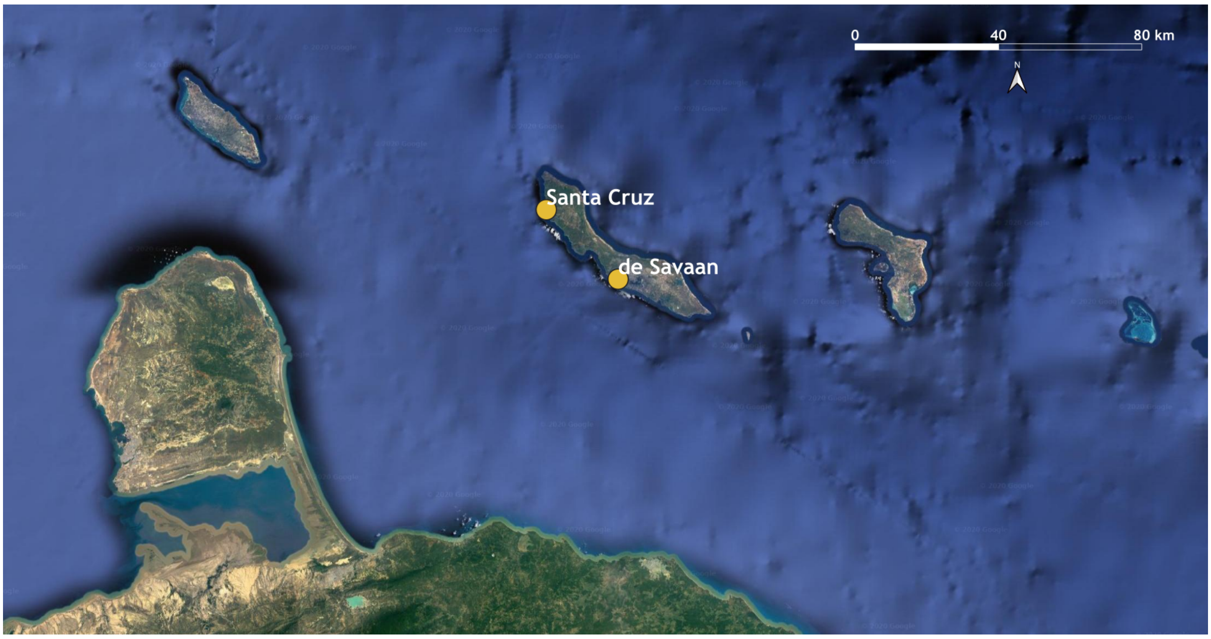

**Figure S9:** Location of two archaeological sites in Curaçao included in this study. Map created using QGIS Geographic Information System v3.6 (http://qgis.org); basemaps from Google Earth. .

Curaçao is the second largest island in a chain of small islands and cays parallel to the coast of Venezuela (called the "Southern Caribbean Region" (Keegan and Hofman 2017)). The island is semi-arid, with a mean annual precipitation of ~550 millimeters and a near constant average annual temperature of ~27ºC. The strong trade wind blows for almost the entire year from the northeast, and hurricanes are infrequent. Previous work (Beers et al. 1997) recognized seven different landscape types with vegetation specific to geology and soil types.

Archaeological investigations on Curaçao were initiated by the amateur archaeologist Antonius J. van Koolwijk between 1878 and 1880 and have continued to the present. A total of 97 Amerindian sites were recorded in 1987. The earliest evidence for human presence on the island is dated between 3735-2895 BCE for the Rooi Rincon site. Jay Haviser (1987) describes the characteristics of five early sites as reflecting Archaic Age practices, with some continuation of Lithic Age technologies. The St. Michielsberg site especially has a material culture that is very similar to El Heneal on the north coast of Venezuela (see Willey 1971:366). The influence of these material cultures disappeared following the arrival of agriculture-associated Ceramic Age material culture about 1,500 years ago. The new arrivals manufactured pottery with motifs that are classified as Dabajuroid and which show direct connections to western Venezuela (Oliver 1997). This pottery has distinctive patterns of parallel incised lines, punctations, and applique. In addition, pottery painted in Dabajuroid motifs on Curaçao reflects continuity with the Second-Painted Horizon of Colombia even further west, and it is not affiliated with painted pottery to the east (i.e., Saladoid). Given that our genetic analysis models about 1/3 of Ceramic Curaçao ancestry as related to that of the individuals we have studied from ~400 BCE - 200 CE from Las Locas in western Venezuela (who were also a ceramic-using), it is parsimonious to identify that the arrival of Dabajuroid pottery in Curaçao about 500 CE was related to the same events that brought this ancestry component to Curaçao. In this context it is striking that the other ~2/3 of ancestry in the Curaçao individuals we analyzed (all from ~1300 CE) can be well-modeled as coming from the Caribbean Ceramic cluster, suggesting that there was major gene flow from the Antilles that affected Curaçao in the centuries after the arrival of Dabajuroid pottery. An important topic for future research will be to identify the archaeological correlates of these events.

Human remains are described as scarce in Curaçao (Tacoma 1990). Haviser (1987) describes the characteristics of seven Ceramic Age sites, including the two sites from which the human remains in this study were recovered (i.e., de Savaan and Santa Cruz). The individuals analysed for this study come from two Caquetio settlements, associated archaeologically with large quantities of Dabajuroid ceramics.

**De Savaan**

The De Savaan site (C-0021) is located in south-central Curaçao about 500 meters from the northern shore of Piscadera Bay, and about 2-3 kilometers from Schottegat Bay (East) and St. Michiels Bay (West). All three bays have access to the sea in the South (Bonaire Basin). The site is located on a hilltop close to gullies where water concentrates after heavy rainfall, and soil is good for agriculture. The settlement and its agricultural lands encompass an area of at least 2-3 hectares (Haviser 1987). From 1980 on, several archaeological excavations have taken place at the site, some of which have been published (Tacoma 1990; Haviser 1987). The finds of NAAM fieldwork in 2016 are still in the process of being analysed. In total, 14 graves have been recovered from De Savaan site. In general, individuals were buried with legs flexed and arms folded in a grave, although there is some evidence of secondary burials. In some cases, individuals were covered with inverted ceramic vessels (Haviser 1987; Kraan et al. 2016). Its context date of 1160-1500 CE and the abundant presence of Dabajuroid ceramics indicate the site was a Caquetio village.

**Santa Cruz**

The Spanish reported the Santa Cruz site (C-0004) as the village of a local Caquetio chief. The site is located on a high hill about 500 meters from the Santa Cruz Bay with access to the sea, close to gullies and with soil suitable for agriculture (Haviser 1987). The settlement encompasses about 2-3 hectares without associated agricultural plots. During two salvage excavations, one in the 1980s (Haviser 1987) and another in 2004 (currently unpublished), a total of three graves were documented, and another identified but not excavated. In general, the individuals were buried in graves with flexed arms and legs. In some cases, grave goods were identified. There was no evidence of urns. Recent research resulted in a context date of 1443-1522 CE (Kraan et al 2016).

**HISPANIOLA:**

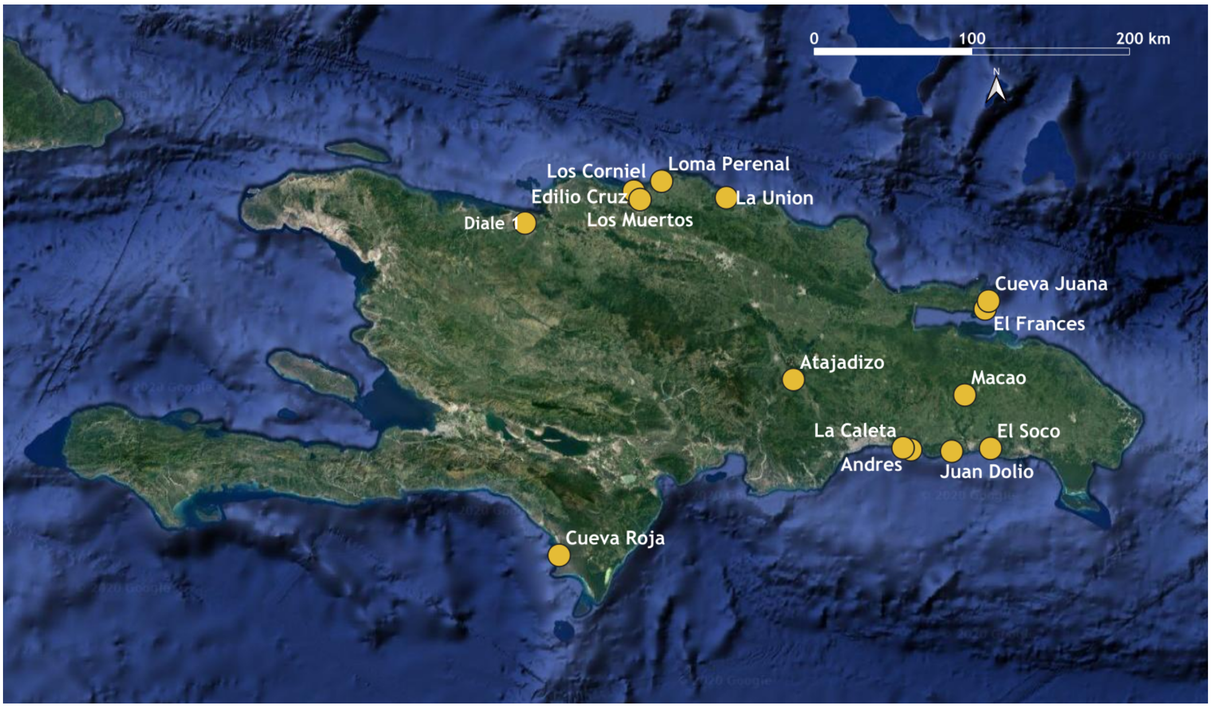

**Figure S10:** Location of 15 archaeological sites on Hispaniola included in this study. Map created using QGIS Geographic Information System v3.6 (http://qgis.org); basemaps from Google Earth.

Sites are discussed beginning with Diale 1 in present-day Haiti and moving clockwise around the island.

**Diale 1**

The Diale 1 and Diale 2 sites are located on the western side of Ft. Liberté Bay on the northeast coast of present-day Haiti. The site is on a peninsula that juts slightly into the bay at the point where the entrance opens into the bay. It is about 5 kilometers north of the modern town of Faiton. Diale 2 is immediately to the south. It differs from Diale 1 (Meillacoid pottery) in being smaller and having Carrier style (Chicoid) pottery. The sites were first investigated by Froelich Rainey from Yale University in 1934 (Rainey 1941). Rainey dug one 18 x 2 meter trench through midden 17 at Diale 1. At the time he was assisted by Irving Rouse, then a student.

After completing excavations at other sites in 1935, Rouse directed the excavations at Diale 1. The site is described as 29 “middens,” “which appear as perceptible mounds,” extending in a roughly north-south direction, sloping gently upward from the shore to a low ridge (Rainey 1941:42-43). The shore is rocky and lined with mangroves except for two small beaches around which the mounds are clustered. Rouse excavated trenches in mounds 1, 2, 5, 6, and 8. The mounds are composed of marine shells; the bones of fishes, manatee, sea turtle, and hutia; and land crabs. Cultural materials include large quantities of Meillacoid pottery, shell and stone implements, and stone and shell beads. The maximum depth of any midden was 1.5 meters, although most are less than one meter deep.

All of the human burials were recovered from mound 5, which has the most burials of any site investigated in the area (Rainey 1941: figures 1-19). The mostly primary burials are in a shell refuse layer at depths between about 25-40 cm. Most are disturbed and partially disintegrated. The remains of at least 7 individuals were recovered. Burial 1 contained two skeletons, side by side; the primary burial of an adult flexed on its left side, and a youth. Burial 3 was an adult male on his back with his head pushed forward and the knees drawn up to the chest. Burial 4 was one meter from Burial 3; it is described as a small child placed on its back in a prone position with legs extended. Described as a secondary burial, the bones in Burial 5 are scattered across a one-meter square area at a shallower depth (about 16 cm below surface).

The human skeletal remains included in this study (ANTPA.000159) were acquired by Rainey in 1935 [according to the museum accession files (Diale 1 site. Section B [*sic* 8], Trench B, Burial 2)]. Burial 2 was in mound 5, section 8, at a depth of 25-30 cm below surface, four meters from Burial 1, and eight meters from Burial 3. Burial 2 contained two skeletons with additional bones from secondary burials. A complete bowl was positioned between the skeletons close to the feet. These primary, flexed burials were disturbed and slightly disintegrated.

Our detection of admixture (10-15% Archaic-related, and 80-85% Caribbean-Ceramic-related) in both of the Diale 1 individuals is striking in light of the genetic findings in the Dominican individuals in our dataset, all dating after ~100 calCE from the same island of Hispaniola including from sites not too far to the east from Diale along its north coast. Of these 121 Dominican ceramic-using individuals, all but one are consistent with having no Archaic-related admixture (the only exception is outlier individual I16539 at La Caleta is inferred to have ~12-14% Archaic-related admixture). These results suggest that Hispaniola may have included within it a cline of admixture between Archaic and Ceramic-related ancestry with sampling proportions of Archaic-related ancestry to the west of the island (present-day Haiti) where we have no sampling except for Diale 1 where we do detect such admixture. Much further to the west in western Cuba at Canímar Abajo the ancestry we detect is entirely Archaic-related even after ~900 calCE. An important topic for future research will be to carry out dense sampling of ancient DNA from western Hispaniola (Haiti) and eastern Cuba to determine the geographic, temporal, and genetic characteristics of this admixture cline and to correlate it to the archaeological evidence for interaction between east and west in this period.

**Edilio Cruz (Rancho Manuel)**

The site (RD/PR/02) is located on the northern side of the road that goes from Rancho Manuel toward Gregorio and the crossroads for Punta Rucia. The area where archaeological materials were found extends for 6000 square meters, while its height above sea level corresponds to 70 meters. The site was identified during the July 2006 Survey conducted by the Italian Archaeo-Anthopological Mission in the Dominican Republic in collaboration with the Museo del Hombre Dominicano. A short excavation test was conducted on the site with the opening of a 2x2 meter trench which led to the identification of a burial. The excavation also produced faunal remains, in particular hutia and manatee, as well as a fair number of fish bones and gastropods.

The archaeological material identified during the excavation mainly includes fragments referable to decorated and undecorated ceramic containers. Most of them refer to the Chicoid phase with the presence of bowls and plates characterized by handles with zoomorphic and anthropomorphic depictions according to the typical style of this phase. A smaller percentage refers instead to the production of Meillacoid culture, typical of the northern coast of the island, and characterized by the presence of low cups and cups decorated with plastic elements depicting zoomorphic depictions and engraved geometric patterns. In addition, the portion of the site investigated produced a good amount of *buren* fragments which are clay plates for domestic use, in particular for cooking food.

**Los Corniel (Rancho Manuel)**

The site (RD/PR/01) is located behind and towards the south of Rancho Manuel. After careful observation of the limits of dispersion of surface materials, the total extension of the site was assessed, which corresponds to 1000 square meters, while the height above sea level corresponds to 57.5 meters. The site was identified during the July 2006 Survey conducted by the Italian Archaeo-Anthropological Mission in the Dominican Republic in collaboration with the Museo del Hombre Dominicano. The mounds appear lined up in a double row on the crest of the hill. Twenty-two mounds were counted.

In order to understand the size of the site and above all to estimate the archaeological deposit and to distinguish exactly the occupation, a verification trench was opened in the area which showed a greater quantity of ceramics and remains of shells.

The trench followed the standard for rapid checks and measured four square meters. Also in this trench the remains of a burial have been identified as well as several elements that shed light on the consistency of the site. Faunal remains, fish bones and gastropods represent the maximum percentage among the finds. The ceramic fragments refer to both the Chicoid and Meillacoid phases, and correspond to highly fragmented material referable to plates and bowls. In addition, some examples of polished stone tools such as millstones, strikers, and axes have been collected.

**Los Muertos**

The Los Muertos site (RD/PR/21) is located on the north slope of the northern cordillera. The entrance is on the southern side of the road from Rancho Manuel to Hestero Hondo. The site consists of three lines of mounds of which the northern and the southern run along the limits of the ridge. Thick vegetation covers the whole area, which includes probably 10 mounds with a size of about 20 meters in diameter and oriented in an east-west direction. A burial was excavated from a depth of 0.60 centimeters (Individual A). It was in a crouched position, oriented with the head to the north and facing west. Fragmentary remains of three other individuals (B, C and D) were also recovered.

**Loma Perenal (Puerto Plata)**

Archaeological research was carried out in 1998/1999 in the locality of Loma Perenal. This site is located on a plateau overlooking the alluvial valley of the Rio Bajabonico just before it flows into the bay of La Isabela. It is situated on the north coast of the Dominican Republic, a few kilometers from the town of La Isabela, in the province of Puerto Plata. This archaeologically-rich area covers the southern portion of the plateau in the direction of the river at approximately 70-80 meters above sea level, and was rich in cultural materials associated with Indigenous peoples. Excavation campaigns in 1998 and 1999 were conducted jointly by the Museo del Hombre Dominicano and the University of Rome "La Sapienza," thanks to the commitment of the Costa Foundation and with the collaboration of the Dominican Dirección Nacional de Parques.

The November/December 1998 campaign focused on Mound 3. The mound has an oval shape with a long axis oriented north-south and dimensions of approximately 16 x 12 meters. The height of its peak measured from the base varies in the southern half of the mound from 66 centimeters (compared to the south/east margin) to 124 centimeters (compared to the south/west margin).

The first layer (US 5) present on the surface consists of reddish-brown earth that is rich in humus as well as in roots, with numerous stones of different sizes variously distributed both on the surface and within the layer. During the excavation of US 5, some layers of ash appeared, light gray in color, of limited extension and with irregular edges. Three similar formations (US 9, 10, 11) were documented and removed in different locations. It does not seem that combustion took place *in situ* to create these layers, but rather that ashes were thrown on the mound before US 5 was formed, after or during the deposition of the underlying layer. Ceramics present include both Meillacoid and Chicoid, with these contexts dated to between 900 and 1500 CE.

There are numerous lithic tools, made mostly of roughly worked stones. The majority of the tools are rather simple, but there are some more elaborate examples. Relevant materials include some personal ornaments, made from bone and shell, as well as a fragment of a spatula. It should be emphasized that the artifacts found are all Indigenous-associated, without any indication of Spanish origin. Animal bones were found (though less frequently), almost all from small rodents or birds. In addition, there are some fish vertebrae.

In May 1999 at approximately 60-80 meters to the southeast of Mound 3, a burial was found. Called Burial 1, it was determined to be a tomb partially disturbed by recent excavations. The excavation was therefore carried out on a disturbed context in which only a few portions of the skeleton were in the original position. The human bones were in poor condition. The rather loose soil, gray-brown in color, was full of fragmented ceramics, lithics, and shells.

**La Union**

The site of La Union was studied as part of an emergency excavation in July 1972 by the Museo del Hombre Dominicano, and consisted of a single trench (Veloz Maggiolo et al. 1972; 1973). It was determined to be the site of a community of fishermen with evidence of the presence of agriculture. It included ceramics of the Chicoid series, with very few Melliacoid ceramics, and was context dated to ~1435 CE. Twenty burials were excavated, 19 of which were associated with funerary goods. Four tombs only contained postcranial skeletal material and were devoid of skulls, consistent with a Taíno practice of keeping the skulls of ancestors in their houses (Veloz Maggiolo 1977). At the La Union site, the large volume of fishing net weights and marine gastropods (*Cittarium pica*) suggested the development of an exchange network characterized by local adaptations in which coastal communities specialized in fishing and collecting marine resources exchanged products with the more agricultural settlements of the interior (Veloz Maggiolo 1993).

**Cueva Juana**

The archaeological site was located at a karst rock spur approximately 100 meters from the east coast of the Cape of Samanà. The archaeological deposit was identified in the portion covered by the shelter and was investigated by the archaeologists of the Museo del Hombre Dominicano in the 1970s. The archaeological material identified during a recent survey is linked to the Ostionoid phase. Four human burials were discovered during the archaeological explorations, allowing us to consider this site as linked to the funeral aspects of the community.

**El Frances**

The research activities on the El Frances site (Samaná peninsula) were conducted in three excavation campaigns during 2018 and 2019 by the Italian Archaeological-Anthropology Mission in the Dominican Republic (Sapienza University of Rome, Museo del Hombre Dominicano, and the Shelley Foundation). During these three campaigns, nine excavation areas were analysed. Both habitation and funerary areas were excavated. Seven excavated primary burials were dated between 788-874 calCE and 968-1023 calCE, and a context date of 532-584 calCE was also obtained from the site.

The site is of particular importance in the history of the development of the cultures in the eastern part of Hispaniola especially with regard to contacts with the nearby island of Puerto Rico, from which a sequence of cultural development subsequently spread throughout the eastern part of the island. The study of ceramics and radiometric dating attribute the site to an ancient phase of the island's ceramic population. This phase coincides with the introduction of the Saladoid material culture, which is found beneath a stratum containing Ostionoid materials. The Saladoid ceramics show clear morphological and stylistic characteristics that distinguish them from the original context, including surface treatment and the white-on-red paint as the main decorative element. During the last excavation, a whale vertebra was found for the first time in a site in the Dominican Republic.

**Macao, El Morro (La Altagracia)**

Punta Macao is a multi-component, but predominantly Late Ceramic Age, habitation site on the north-eastern coast of the easternmost province of La Altagracia. The site is situated on a rocky promontory called El Morro and is near the modern-day town of Macau. The site is mentioned in Las Casas' Apologética Historia, in which he claims to have visited the town and notes that a large population inhabited the area where almost 100 mounds were identified (de las Casas 1992; Olsen 2004; Veloz Maggiolo 1972; Veloz Maggiolo and Ortega 1972). Excavations at the site were undertaken by De Booy in 1915, Rainey in the 1940s, Veloz Maggiolo and Ortega in 1972, and a team from the Museo del Hombre Dominicano in 2004 (Atiles 2004; Olsen 2004; Tavarez María 2004).

The excavations conducted in 1972 showed two levels of occupation, an older Level I comprising an Ostionoid horizon, and a more recent Level II comprising a Chicoid horizon. Some evidence of Saladoid ceramics also was found at the base of the stratigraphy. The "Monticulos" are agricultural mounds, some of which contain burials, and most of which are disturbed by illegal excavations. The subsistence economy of the Level I population was mainly agricultural, supplemented by gathered plants and to a lesser extent fishing. Agriculture continued in Level II, but fishing increased compared to gathering. In general, more than 50% of marine-type foods were oysters and crabs collected from shallow water. Land snails represent 90% of the animal protein collected on land.

The skeletons included in this study come from 15 burial pits containing the remains of 26 individuals that were excavated by the Museo del Hombre Dominicano in 2004 (Atiles 2004; Olsen 2004; Tavarez María 2004). Most burial pits contained the remains of a number of individuals, who seemed to have been deposited simultaneously in primary depositions. The individuals were primarily males (n=11) and juveniles (n=8), with only six females present (Tavarez María 2004). The bodies were generally interred in small round or oval burial pits in supine positions or reclined on one side. All of the skeletons were in a flexed position, with the legs drawn up to the chest. Burial 2 was interred with a large Chicoid ceramic vessel placed upside down on the head, a practice observed in other contemporary cemeteries. The majority of the burials were located in the southern part of the site, leading the investigators to suggest that this part of the site comprised the cemetery area. Many burials remain unexcavated in this part of the site, which has since been transformed into a golf course (Atiles 2004; Olsen 2004; Tavarez María 2004). Three radiocarbon dates, ranging from 1240±40-790±60 BP (uncalibrated) were obtained for three skeletons excavated in 2004, though it is unclear which three (Hofman et al. 2007). These dates, along with ceramic finds associated with the burials, indicate a predominantly Ostionoid and Chicoid chronology for the burials. However, some Spanish colonial ceramics (*majolica*) were found in the cemetery area of the site, perhaps indicating that the cemetery was still in use during the early contact period. The precise relationship between these ceramics and the burials is unclear (Atiles, 2004).

**El Soco**

The Boca del Soco site was excavated in 1975 and 1980 by researchers from El Museo del Hombre Dominicano. The site is located in the province of San Pedro de Macoris approximately 75 km east of Santo Domingo (Luna Calderon 1985; Veloz Maggiolo 1972, 1993). Two occupational phases are present at Boca del Soco. The first is an Ostionoid period occupation called the Margarita Phase and dating to around 700-800 CE. This occupation was excavated in 1975 with the name of El Soco I. The second phase of occupation is characterized by Chicoid ceramics (called the El Soco Phase and dating to around 1000-1500 CE). It was excavated in 1980 under the name of El Soco II. The burials are in both primary and secondary contexts, and are accompanied by grave offerings such as pottery and dogs. Burial was collective during the Margarita phase.

A total of 158 individuals were recovered from Boca del Soco, 118 belonging to the Margarita Phase and 40 belonging to the El Soco Phase (Coppa et al. 1995). A high incidence of disease, along with infections, caused a high mortality rate among the sub-adult portion of the population, mainly infants in the Margarita Phase. This is in contrast to the high rate of adolescent death in the El Soco Phase (Luna Calderon 1985). In general, the Boca del Soco population was not in good health. Comparatively, the mortality of the adult population is higher than in the Juan Dolio cemetery, located about 20 kilometers away (Coppa et al. 1995). The isotopic data of El Soco, from eight samples from the Margarita Phase and only one from the El Soco Phase, suggest that the diet was based more on terrestrial proteins than Juan Dolio's samples (Stokes 1998).

**Juan Dolio**

Juan Dolio is a Late Ceramic Age site, located on the southern coast of the Dominican Republic, approximately 70 kilometers to the east of the capital city of Santo Domingo. The site was excavated by numerous researchers (Boyrie Moya 1960; Boyrie Moya and Cruxent 1955). It is known as one of the key sites of the "Boca Chica" (Chicoid) ceramic style in the southern and south-eastern Dominican Republic, however, as excavations at the site focused predominantly on the cemetery area, little is known about the house structures and material culture at the site. The archaeological remains appear in two different, separate humiferous layers, clearly detailing two important Indigenous occupations (Veloz Maggiolo 1972:157).

Juan Dolio was occupied as late as European contact as evidenced by the abundance of Spanish and other European pottery sherds in upper levels and in some burials (Garcia Arévalo 1978, 1990; Goggin 1960; Keehmen 2019; Ortega and Fondeur 1978; Veloz Maggiolo 1993).

Skeletal remains of 102 burial contexts were identified during excavations at Juan Dolio under the direction of Fernando Luna Calderón in 1974; however, many of these individuals comprised only a few fragmentary bones. Further analysis of the skeletal assemblage yielded an estimated minimum number of individuals of 78 persons, of which 31 were adult males, 29 were adult females, 18 were individuals of unknown sex, and 11 were juveniles (Drusini et al. 1987; Veloz Maggiolo 1972). A subsequent, more detailed study identified the skeletal remains of 108 individuals (Coppa et al. 1995). A large number of skeletal remains uncovered at the site during excavations in 1974 reportedly date predominantly to the late 15th century (Drusini et al. 1987; Veloz Maggiolo 1972).

As at Boca del Soco, cranial deformation was common (Drusini et al. 1987). In comparing the skeletal material from Boca del Soco and Juan Dolio, Coppa et al. (1995) found that living conditions improved from the early Taíno period (El Soco) to the period directly preceding European contact (Juan Dolio). The analyses conducted on the morphology and morphometry of the teeth of this necropolis in comparison with the pre-Ceramic one of Cueva Roja and with El Soco had highlighted marked differences such as different mobility patterns at the origin of the two cultural and chronological phases (Coppa et al. 1995). These differences have subsequently been confirmed on a larger sample and using more sophisticated multivariate statistics (Coppa et al. 2004,, 2008; Cucina et al. 2003) and can be further tested using paleogenomic data.

A study on syphilis showed that this population like the others of the Ceramic Age were endemically affected by this disease, while the Archaic population of Cueva Roja shows no evidence of this pathology (Rothschild et al. 2000). Isotopic analyses for the study of diet have shown a mixture of marine and terrestrial diet with a more significant presence of the terrestrial component (Stokes 1998).

**Andrés**

The archaeological site of Andrés is situated on a narrow sand spit projecting into the Bahia de Andrés on the Caribbean Sea near the Dominican town of Boca Chica. It is about 25 kilometers east of the capital city of Santo Domingo and directly adjacent to the sugar warehouses of the Compania Azucarera Boca Chica and the adjoining Dominican village of Andrés. A first exploration of Andrés was carried out in 1922, with the excavations directed by Mr. Amado Franco Bido. The materials were studied by Dr. Narciso Alberti y Bosch, who in 1920 had also recovered many materials from the area. The entire bay appears to be a single large Taíno site, probably composed of multiple villages. In 1929, the American Museum of Natural History (AMNH) in New York City arranged a visit to the site. Construction of a new sugar warehouse during the previous year had exposed a substantial quantity of ceramics and human skeletons. Through this expedition, the AMNH obtained a collection of skulls, entire skeletons, and earthenware vessels, along with the skulls from excavations made at the cemetery directly in front of the warehouse (Krieger 1931). In 1930, new excavations were undertaken by the Smithsonian Institution (Krieger 1931). Somewhat later, Dr. Narciso Alberti y Bosch of the National Museum of the Dominican Republic went to Boca Chica and collaborated in further excavations on the sandy beach in front of the sugar warehouse and obtained additional earthenware vessels and a large number of skulls (Alberti y Bosch 1932).

The skeletons used in the present study come from materials excavated and published by Alberti y Bosch (1932). Complete vessels and pottery sherds exhibit Chicoid motifs associated with the Late Ceramic Age Taíno (Krieger 1931). The AMS dates obtained from most of the human bones are consistent with the cultural materials, however one skeleton returned a date of 2890±20 BP. This pre-Ceramic individual clearly represents an earlier occupation of the area, but there is insufficient site documentation to determine its particular context. There is, however, substantial evidence that this area continuously was exploited by humans since their initial arrival on the island (Veloz Maggiolo 1972, 1993). The specific location of the Andrés burials has been explained as reflecting limited access to sandy soils (Krieger 1931). It is therefore not surprising that individuals from different time periods were buried in the same location.

**La Caleta**

La Caleta is a multi-component habitation site with a large burial population situated in the town of La Caleta, approximately 17 kilometers east of Santo Domingo. The site was first inhabited during the Archaic Age, with radiocarbon dates indicating an early occupation around 545 BCE that continued throughout the Early and Late Ceramic Ages, with the most recent radiocarbon dates around 1280 CE (Morbán Laucer 1979; Ortega 2005). The site belongs mainly to the Boca Chica cultural horizon ("Chicoid"), but there is also evidence of Saladoid ceramics from the early Ceramic Age (Veloz Maggiolo 1972). During the excavation of the site, but not in association with the burials, European-made objects also were found (Veloz Maggiolo et al. 1976; Keehnen Floris 2019).

Numerous excavations have been undertaken at the site over the years, uncovering at least 373 human skeletal remains (Morbán Laucer 1979). A first set of burials was excavated during the 1944-1945 campaigns (Herrera Fritot and Youmans 1946). This collection includes 14 skeletons, five of which were brought to Cuba and two to the Dartmouth College Museum in Hanover, New Hampshire. These are primary depositions in a flexed position. Further excavations were conducted in the following years (Boyrie de Moya and Herrera Fritot 1948). Most of the other burials in the collection were excavated in 1970–1971 by Chanlatte Baik and Morbán Laucer among others (Veloz Maggiolo 1972).

Individuals were interred both as primary and secondary depositions in the midden area and other parts of the site. Primary interments consisted mainly of flexed, supine skeletons, with the legs drawn up to the chest. In a small number of cases, individuals were interred in a seated, flexed position. A large number of subadult skeletons (fetuses and infants) were recovered from the site. In one case, two juveniles pertaining to the Ostionoid occupation of the site were found buried with seven ceramic vessels of different sizes and a dog. Other grave goods include sherds of Ostionoid pottery, stone axes, shell amulets and vomiting spatulas, and the remains of marine foods. Secondary burial of juveniles sometimes consisted of interment in a ceramic vessel. Some individuals, both adults and juveniles, were found buried with a ceramic vessel placed over their head and/or face (Morbán Laucer 1979). The skeletons used in the present study come from the excavations of 1970-1971.

**Atajadizo (La Altagracia)**

The El Atajadizo site is located near the southeastern coast at the mouth of the Yuma River. It was partially investigated in 1974 by a field party from the Museo del Hombre Dominicano.

The beginning of the ceramic period occupation in El Atajadizo probably occurred around 800 CE. Veloz Maggiolo and colleagues obtained a radiocarbon date of 840±80 CE for this phase (I-8649, Veloz Maggiolo et al. 1976b). The necropolis has two occupation phases called the Atajadizo Phase (840-1000 CE) and Guayabal Phase (1000-1400 CE) on the basis of a series of radiocarbon dates made at the time of the excavations in 1975 (Veloz Maggiolo et al 1976b; Wilson, 2007). European-made objects were also found during site excavation, but not in the burials (Veloz Maggiolo et al. 1976b; Keehnen Floris 2019).

Mounds (Monticulos) 1 and 2 are placed in the Atajadizo Phase. A skeleton in Monticulo 1 was recovered associated with Chicoid pottery fragments. In deeper levels, fragments of Ostionoid ceramics were found. Mounds 4 and 5 are placed in the Guayabal Phase. Mound 3 contained no artifacts, and Mound 6 was disturbed by looters who left the skeletons *in situ*. These mounds cannot be attributed to either phase due to the lack of archaeological material.

In addition, there were burials in Trench 1 from the Plaza Sud Ouest (Luna Calderón 1976a, 1976b; Veloz Maggiolo et al. 1976a, 1976b). A bioanthropological study of 47 burials was conducted. Several of the burials in Mound 4 (Guayabal Phase) show signs of violence, which was interpreted as evidence of inter-tribal struggles (Luna Calderón 1976a, 1976b). Finally, the burial 6 of an adult in Mound 5 (Guayabal Phase) is missing its skull (as also seen in the La Union site), which could be a consequence of the Taíno practice of curating human skulls in their houses (Veloz Maggiolo 1993).

Subsistence during the older Atajadizo Phase was based on the gathering of terrestrial foods (land snails and hutia) and, to a lesser degree, fishing and gathering of seafood from mangroves. For the more recent Guayabal Phase, there is an increase in the gathering of molluscs from the mangroves as well as fishing. Gathering was also reported for both phases to be of greater importance than agricultural activities. The faunal data show that the marine resources comprise on average about 50% of the sample, including littoral molluscs and crustaceans (Rímoli 1976). The presence of both Chicoid and Ostionoid ceramics indicates that this site was occupied during a transition in pottery traditions.

**Cueva Roja (Pedernales)**

The excavation of the site of Cueva Roja took place in 1978, financed by the National Geographic Society and the Universidad Central del Este. The deposits are in a small cave near the sea and comprise skeletal remains with some associated cultural materials. The skeletal remains of 98 individuals in secondary deposition were recovered, of which 54 were adults over the age of 25 years, 10 were between 10 and 24 years of age, and 34 were under 10 years old. Some remains have been cremated.

The preliminary analyses conducted on the morphology and morphometry of the teeth from this necropolis identified marked differences in comparison with those of later ceramic users from El Soco and Juan Dolio. These differences generated the hypothesis that different mobility patterns are reflected between the two cultural and chronological phases (Coppa et al. 1995). The high fracture index at Cueva Roja is compatible with a mobile, transhumant way of life. This hypothesis subsequently was confirmed on a larger sample using more sophisticated multivariate statistics. The analysis also showed that the population of Cueva Roja exhibited a high biological affinity with the pre-Ceramic Guanahatabey of Cuba (Coppa et al. 2004, 2008; Cucina et al. 2003). Our genetic analysis of both Cueva Roja individuals which identifies entirely Archaic-related ancestry in both individuals is strongly consistent with this inference. An examination for evidence of syphilis showed that this population, as opposed to those of the Ceramic Age, was not affected by this pathology (Rothschild et al. 2000).

**PUERTO RICO:**

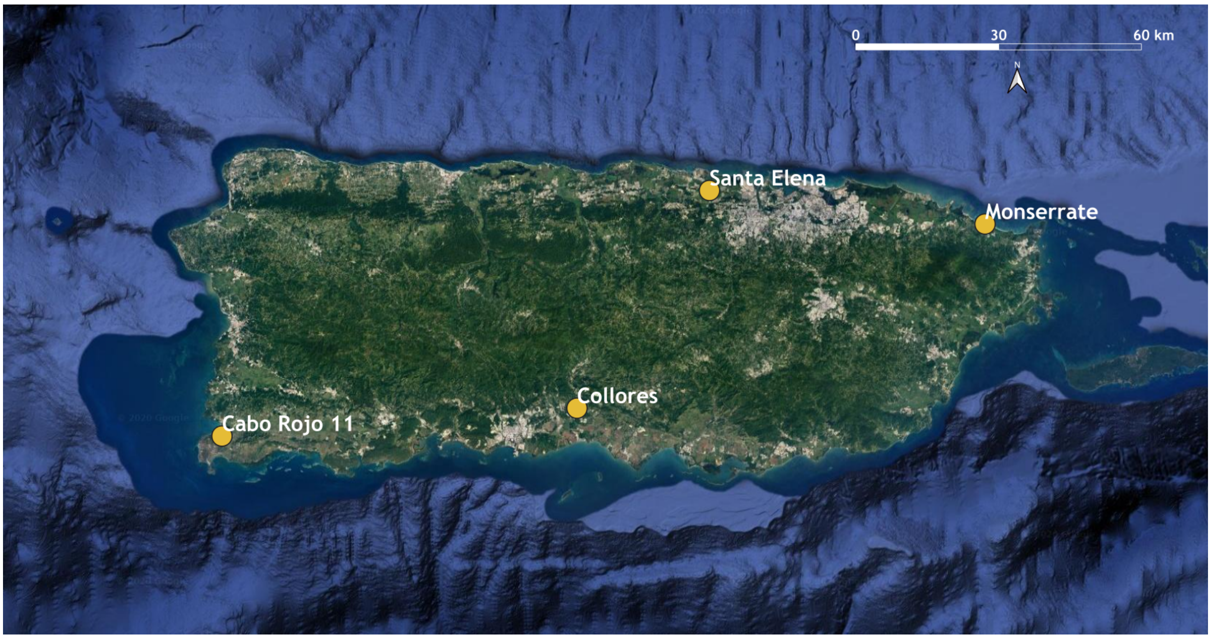

**Figure S11:** Location of four archaeological sites with known locations in Puerto Rico included in this study. Map created using QGIS Geographic Information System v3.6 (http://qgis.org); basemaps from Google Earth.

The remains from Puerto Rico were acquired by Rainey in 1934 and in 1935, (ANTPA.000161, ANTPA.000163, and ANTPA.000165) and in 1938 by Irving Rouse (ANTPA.000150, ANTPA.000151, ANTPA.000152, ANTPA.000153). The human remains from Puerto Rico were described by Drew (2009). Rouse (1952a, 1952b) refers to the Puerto Rican sites by more than one name, including several sites that have the same name and a numerical suffix. For example, Cabo Rojo has four such variations in publications (Rouse 1952a:311) and seventeen in unpublished notes.

**Collores**

Collores is located on Puerto Rico's mid-Southern coast, in the foothills approximately 2 kilometers northwest of the town of Juana Díaz (Rouse 1952b:532). Shell debris covered approximately half an acre of the site. Modern agriculture had impacted the site, but two large middens, each "50 centimeters high", were still present when Rainey and Rouse examined the site (Rouse 1952b:532). A modern road cut through one of the middens (Midden A), but the other midden (Midden B) was intact when Rainey encountered it. Rainey excavated two sections of Midden A. Excavation area 1 contained pottery of the Cuevas style (late Saladoid) and Ostiones and Santa Elena styles (Ostionoid) distributed throughout, with Ostiones pottery in the highest density. Rouse found the mixture of styles to be "unusual," and likely a disturbed deposit (Rouse 1952b:533). According to Rouse (1952b:532-533), "Charcoal, shells, animal bones, and potsherds, probably of the Ostiones style, were observed in the black loam of both middens."

ANTPA.000161 was recovered from the Collores site (Juana Díaz 1). This site was excavated by Rainey in 1934, and Rouse (1952b:532) says that it is located in “almost the exact center of the south-coast area” of Puerto Rico. Rouse visited the site on September 15, 1936. Describing the earlier excavations, Rouse (1952b:535) says: “Two burials were encountered in the lowest, "red culture" layer ... The first, which lay at a depth of 150 centimeters in section Dl, consisted of an adult female skeleton lying flexed on its left side. Little more than a meter away, and at a slightly lower depth, were the remains of a baby, too badly disintegrated to determine the position of the body.”

Rouse sorted excavation area 2 into two divisions based on ceramic styles: a Cuevas division (late Saladoid) and an Ostiones (Ostionoid) division. Both of the burials were found in the Cuevas division. After evaluating Rainey’s excavation notes and artifacts from area 2, Rouse felt it possible that the lowest layer, which contained the burials, were deposited “*in situ*”, and that the rest of the material “eroded down the hillside at a later date.” (Rouse 1952b:537).

**Monserrate**

The Monserrate site (Luquillo 1) was excavated by Rainey in 1934 and 1935. The site is located near the mouth of a small lagoon on Punta Embarcadero in Barrio Mameyes of the municipality of Luquillo (Rouse 1952a:419), east of San Juan near the northeast corner of the island. Rainey noted five distinct mounds, rising 1-1.5 meters above ground, and abundant surface ceramics at the site. ANTPA.000163 was one individual in a group of three burials in section B-5, level 1.25 meters, Mound A, Barrio Monserrate, according to museum catalogue records. Rainey (1940:78) states that this section of the mound contained many flexed burials, and in the level where this individual was found “...the deposit was so filled with skeletons that some sections appeared to contain a massed burial.” Shell refuse was abundant in the mound, along with the remains of other animals and artifacts. The most abundant pottery in Mound A was Ostionian Ostionoid (Rouse 1952:421).

**Cañas/Unknown**

Cañas is located three kilometers north of the Caribbean Sea on the east bank of the Río Cañas (Rouse 1952b:552), in a plain between the ocean and sea on the southwest coast of Puerto Rico near Ponce (Rainey 1940:7). Cañas was Rainey’s “principal site” where he first defined his Crab (red-on-white Saladoid) and shell (incised) pottery (Ostionoid) cultures (Rouse 1952b:522). According to Rainey (1940:7), the archaeological material at Cañas was well known to locals and prior to his excavations antiquities collectors would gather artifacts after field plowing exposed them. Rainey excavated in an overgrown land plot that contained “several mounds of considerable elevation (Rainey 1940:7).” Cultural refuse was abundant on the surface of the site. Rainey excavated the largest mound and put test pits in two additional mounds. Two sections of the large mound were excavated; we analyzed an individual from section # 2 (PA 164; I13542), the southwestern area of the mound. Nine burials were encountered in this area. According to Rainey (1940:12), seven were from a substratum of land-crab shells but were so poorly preserved that they could not be adequately studied. We analyzed one of the two individuals preserved well enough to be characterized osteologically (a morphologically female adult, Drew 2005); however, the individual failed to produce working ancient DNA data.

In addition to the failed Cañas individual, we analyzed two individuals of uncertain provenience in the Yale Peabody collections that could potentially be from Cañas. Rouse inherited a collection of materials from Rainey. The collection contained human remains, a large assortment of high-quality pottery, and other artifacts (Drew 2005). According to Drew (2005:528), a piece of highly-polished redware labelled "Cañas #2 G-4 1.00" was in the materials (probably Ostionoid), which may suggest a provenience for the materials. Drew (2005:528) also surmises that "the state of preservation of these remains is consistent with coastal locations where the abundance of shell fragments inhibited decomposition." The individuals we analyzed were part of a set of multiple co-mingled individuals comprising at least three adults and two infants. They are cataloged as being from Canas, Collores or Monserrate, three different sites in Puerto Rico.

**Cabo Rojo 11**

From 1937-1938 Rouse surveyed and excavated a number of sites around the Cabo Rojo municipality. ANTPA.000150 was recovered from the Cabo Rojo 11/Llanos Tuna site (Rouse 1952a:391), the only definite ball court site Rouse excavated during his 1930s fieldwork. This site is located in Barrio Llanos Tuna of the municipality of Cabo Rojo, about four kilometers southeast of the town of Cabo Rojo in the western part of the island. This site is near an unnamed stream, believed to flow northward and westward into the Rio Guanajibo and the bay of Mayagüez. Most of the pottery from this site was of the Ostiones type (Ostionoid). According to Rouse (1952a:391), Ostiones sherds were common on the surface along with other artifacts, but not charcoal, ash or animal bones. Rouse excavated a 4 x 4 meter test at Cabo Rojo. One burial was encountered in this test unit. Rouse (Rouse 1952a:393) describes the burial as:

> "A skeleton of an adult was found in section Bl of level 2, apparently flexed and lying on its left side. This skeleton was much disintegrated partially because the shells were closely packed and partially because it had been penetrated by the roots of a tree. Only fragments of the skull, parts of the pelvis, and the long bones remained. The latter were inclined, some one way and some the other, as if they had been moved out of position. The burial was directly in the refuse, without associated artifacts."

Drew (2005:525) determined the individual to be morphologically an adult male.

**Toa Baja 2**

The site of Toa Baja 2 (sometimes referred to as Santa Elena) is located on Puerto Rico's north coast. The large village site is situated where the tracks of a cane railroad crossed the valley of the La Plata river, about 1.5 kilometers above Toa Baja and 5 kilometers from the sea near the north, central shore of the island. ANTPA.000151, ANTPA.000152, and ANTPA.000153 were recovered from this site (Rouse 1952a:426). Rouse (1952a) noted that the site was well known to locals prior to archaeological investigation. According to Rouse (1952a), the site was located on a bench above a river floodplain and about five acres in size. Rouse excavated four 2x2 meter units at the site during the 1937-1938 field seasons. The ceramics at the site suggested it had both a pre-contact occupation as well as a contact and post-contact occupation (European ceramics were found on the uppermost layers). Sherds in the Precontact layers were primarily Santa Elena style (Elenan Ostionoid).

Rouse (1952a:428-429) encountered three burials in the pre-contact strata. The first burial was found at the lowest shell stratum in the northwestern corner of section A2, at an average depth of 118 centimeters. The individual was in a flexed position lying on its right side. The bones were in good condition and there were no associated artifacts. The second burial was found in section Al, at a depth of from 136 to 160 centimeters. The burial included the remains of an infant that were badly decomposed and included the skull, vertebrae, ribs, and a few long bones. This infant was between the legs and arms of an adult. Most of the adult skeleton was outside of the excavation unit, but it lay on its left side in a flexed position with no associated artifacts. The third burial was found about 20 centimeters under the second burial, and also included an infant associated with an adult. The infant remains were in poor condition whereas the adult was better preserved. The adult was buried in a flexed position on its right side with no associated artifacts.

**VENEZUELA:**

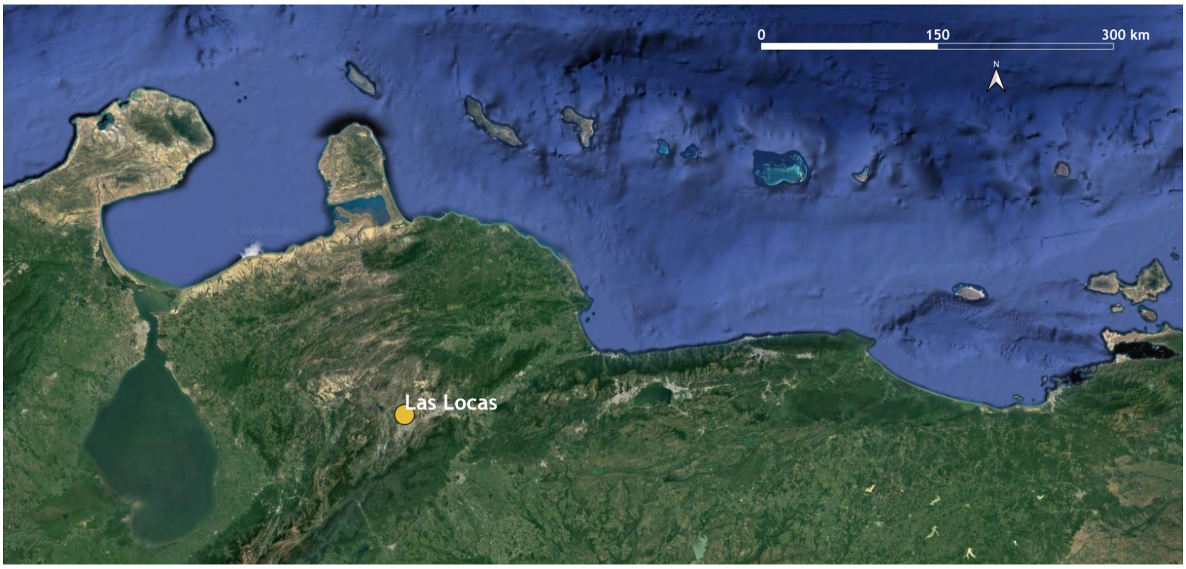

**Figure S12:** Location of the archaeological site in Venezuela included in this study. Map created using QGIS Geographic Information System v3.6 (http://qgis.org); basemaps from Google Earth.

**Las Locas (Quíbor Valley, Jiménez Municipality, Venezuela)**

Las Locas is the oldest necropolis described in the Quíbor Valley, located in the west-central region of Venezuela. It is located 11 kilometers northwest of the city of Quíbor in the Jiménez Municipality in Lara. The cemetery was first excavated by Mario Sanoja and Iraida Vargas as part of the Archaeological Project of the West of Venezuela (Sanoja and Vargas 1967). The site was estimated to date to around 500 BCE, based on the time range established for the Tradition Santa Ana style of the ceramics collected in the cemetery (Sanoja and Vargas 2007). We obtained slightly later dates on bone from two genetically homogenous individuals directly analyzed in this study: 2360-2315 cal. yr BP (PSUAMS-7365) for I17892, 2360-2330 cal. yr BP (PSUMAS-7364) for I17889. These calibrations do not account for any marine reservoir correction (see Supplementary Information section 3 and Supplementary Data 3 for additional information on newly generated radiocarbon dates).

The skeletal remains studied as part of this work are deposited in the "Gonzalo Rincón Gutiérrez" Archaeological Museum of the University of Los Andes, Mérida-Venezuela. The archaeological remains found in the necropolis (mortuary treatment associated with exuberant funerary ceramics) provide evidence concerning the emergence of hierarchical societies in northwestern Venezuela.

Our genetic data from Las Locas are notable in two ways. First, while the Las Locas individuals were members of a community that used ceramics, this does not exclude the possibility that the region where they lived was a possible point of origin for the populations who moved into the Caribbean at a much earlier time during the Archaic Age and established an ancestry profile at least in the Greater Antilles that we examine in this study. Population structure analysis revealed similar ancestry patterns between the Las Locas individuals and our Archaic-related individuals from the Greater

Antilles, while in maximum likelihood trees the latter are positioned in a node leading to Las Locas and present-day Chibchan-speaking groups. This position, however, is not clearly confirmed with $f_4$-statistics as we could not find a closer affinity of the Archaic-related individuals to Las Locas or any Chibchan-speaking group when compared to other present-day or ancient populations. While additional ancient DNA from the region from earlier in time would allow further testing of a scenario where the Archaic Age movement into the Caribbean involved people living in this part of South America in groups ancestral (several millennia before) to the people of Las Locas, it is possible that the Las Locas individuals we analyzed (who date from later periods) would be best seen as an Archaic-related population impacted by later admixture, despite their intensive ceramic use. Second, a striking genetic inference from Las Locas is that the ancestry of the individuals we analyzed is a good proxy source for about 1/3 of the ancestry of the individuals we analyzed from Curaçao (see also a discussion of this in the section on Curaçao). Combined with the archaeological evidence of the spread of Dabajuroid pottery around ~500 CE (which represents a material connection between Curaçao and the region of west Venezuela where the site of Las Locas is found), our results suggest that Las Locas-related ancestry spread to Curaçao from the mainland in association with the spread of Dabajuroid material culture, largely displacing whatever ancestry was previously present in Curaçao (as the other component of ancestry in the Curaçao individuals we analyzed was primarily associated with that seen in people living in the Greater Antilles and The Bahamas during the Ceramic Age).

**SI3- Newly Reported Direct 14C Dates**

Direct radiocarbon dates obtained from skeletal material from the Caribbean have the potential to improve our understanding of a range of topics of interest, including the timing of population movements into and around the Caribbean as well as the onset of new cultural traditions and transitions between traditions. Radiocarbon dates require calibration to account for changes in the global radiocarbon concentration over time (Bronk Ramsey 2008; Taylor 2009), and skeletal samples from the Northern Hemisphere are most often calibrated using the Intcal13 curve (Reimer et al. 2013), which represents the mid-latitude Northern Hemisphere atmospheric reservoir. However, it has been previously argued that radiocarbon dates from insular biomaterials should undergo correction for the marine radiocarbon reservoir effect, as $^{14}C$ is not equally distributed across the biosphere and marine ecological zones leading to an offset in $^{14}C$ age between contemporaneous organisms from the terrestrial environment and organisms deriving carbon from marine environments; this offset averages around 400 years (Aitken 2013). It has been shown that the marine reservoir effect can influence the $^{14}C$ concentration of radiocarbon dates determined from humans that consumed marine-derived proteins (e.g., Ascough et al. 2012), as would be expected for populations of the Caribbean.

The use of isotopic information from ancient individuals (specifically a $\delta^{13}C$ value and a $\delta^{15}N$ value) to gain insight into the proportion of diets dependent on marine protein sources has been argued to be an important part of accurately calibrating radiocarbon dates, while cruder corrections accounting for a certain percentage of diet estimated to originate from marine sources (considered correcting for the Marine Reservoir Effect, or MRE) are often applied when isotopic information is lacking (e.g., Napolitano et al. 2019). This involves calibrating dates using the Marine13 radiocarbon calibration curve (Reimer et al. 2013), or a mix of the IntCal13 and Marine13 calibration curves that accounts for a specified percentage of diet being marine-based. It is important to note that the Marine13 curve represents a hypothetical "global" marine reservoir, which serves as a baseline for regional oceanic variations (Reimer et al. 2013), and that a local reservoir value (ΔR) must then be applied to take into account any differences between the global average marine reservoir offset and local variations (Cook et al. 2015; Russell et al. 2015).

While the majority of skeletal samples included in this study derive from the insular Caribbean, we do not automatically apply a marine correction to our dates presented in this manuscript for several reasons. First, the Caribbean lacks ΔR values for nearly all of the islands. These corrections may have a strong influence on the chronologies of some islands (Fitzpatrick and Rick 2015). Second, the generation of diet mixing models that accurately reflect the consumption of sources of food from terrestrial and marine environments in the Caribbean presents a particular challenge. Estimates of percent contributions of marine and terrestrial foodstuffs to the diet of ancient Caribbean peoples remains under debate, and it is likely to have varied significantly though space and time. Furthermore, the analysis of isotopic information obtained from bone collagen alone (generated alongside radiocarbon dates) is likely only to reflect the protein component of diet (see Pestle 2013;

Stokes 1998), with isotopes from bone apatite also required to distinguish whether the carbohydrates consumed were a mix of $C_3/C_4$ (and in what proportion) or whether individuals consumed a monoisotopic diet (Ambrose and Norr 1993).

With the introduction of agriculture into the Caribbean at the onset of the Ceramic Age, the contribution of plant foods to the diet increased substantially. While manioc was the staple crop in South American tropical forests (and probably the Caribbean, along with sweet potato), maize also arrived early to the islands (probably with the earliest inhabitants; Pagán-Jiménez et al. 2015) and seems to be especially important because of its high protein content. However, maize quickly depletes tropical soils due to a high nutrient requirement and therefore is a secondary crop outside river drainages where soils are replenished by annual floods. Maize was certainly a part of Caribbean diets, though this percentage varied by location and environmental conditions (Figueredo 2015). However, stable isotopes are largely unable to aid in the quantification of maize consumption in the ancient Caribbean. This is because maize (a $C_4$ plant) has an $\delta^{13}C$ signature of around 10 per mil, similar to the $C_4$-like signature observed in fish and mollusks from Caribbean coral reef environments (Keegan and DeNiro 1988). It is therefore difficult to determine the proportion of $\delta^{13}C$ contributed by carbohydrates (maize) versus protein (marine organisms) using bone collagen alone. Though fish could potentially be a major component of diet for people on an island, many of the Greater Antillean sites included in this study are not immediately on the coast, and so lacked direct access to marine proteins. Furthermore, the success of fishing is limited by weather, seasonal and other conditions that limit catches, and there is also evidence for fairly rapid local resource depletion, including the size of fish caught, even at low population densities (Carlson and Keegan 2004; Fitzpatrick and Keegan 2007).

We provide conventional radiocarbon dates in years BP and three calibrated dates for each conventional age in Supplementary Data 3. The first date is calibrated using the IntCal13 curve (Reimer et al. 2013); the second date is calibrated using a mix of the IntCal13 and Marine13 (Reimer et al. 2013) curves in a 75%:25% ratio, accounting for a 25% marine protein contribution to the diet; and the third date is calibrated using a mix of the IntCal13 and Marine13 curves in a 50%:50% ratio, accounting for a 50% marine protein contribution to the diet. We report all direct $^{14}C$ dates in the main manuscript as calibrated years (cal. yr) before present (BP), using the IntCal13 curve data for the calibrated dates.

**SI4 - Principal Component Analysis**

Here we provide the same PCA as in Fig. 2a, but we separate the projected ancient individuals from the Caribbean and Venezuela by site. For The Bahamas, we cluster all sites from each island together.

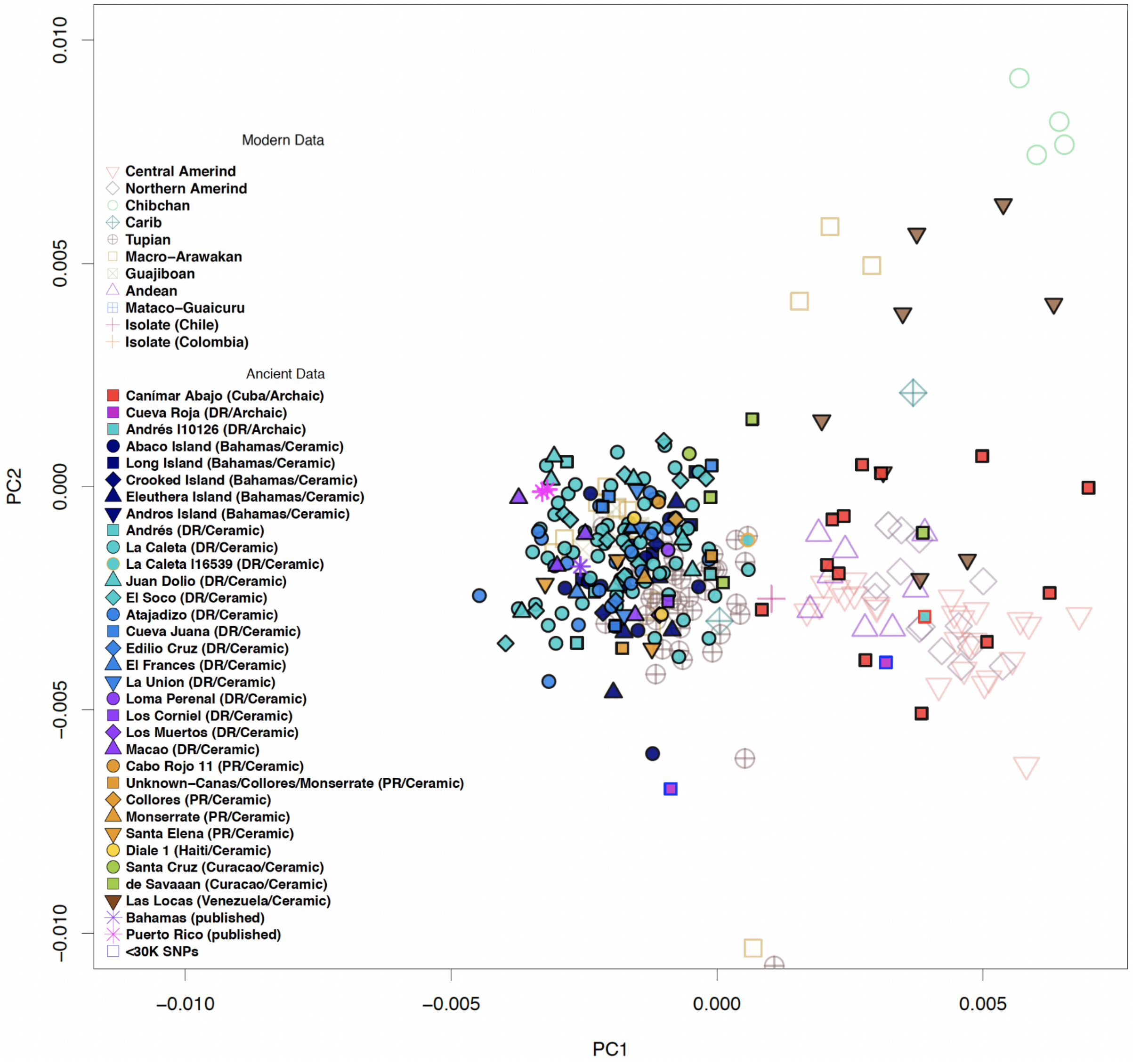

**Figure S13:** PCA with 184 new ancient samples separated by site. Information in parentheses includes the present-day country where the site is located (DR, Dominican Republic; PR, Puerto Rico) and predominant associated technology.

In order to visualize the Ceramic-associated cluster in greater detail, the PCA in Fig. 2a is zoomed for focus on this cluster specifically. Here we provide a non-zoomed version of the same PCA in order to visualize the extent of genetic diversity of the 23 modern populations used to calculate the PC axes (Figure S14).

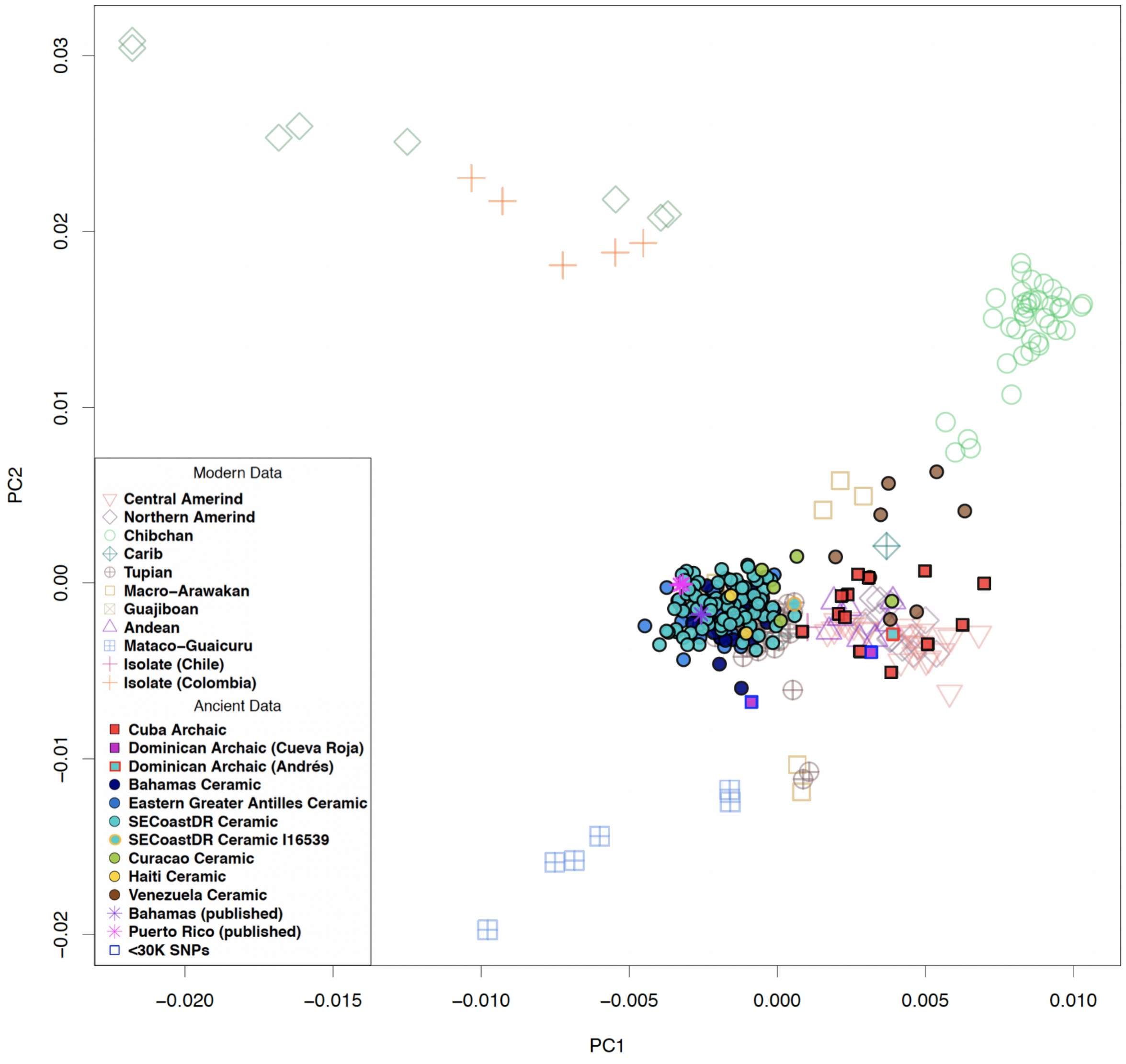

**Figure S14:** Non-zoomed version of PCA in Fig. 2a.

We confirmed that our qualitative results were not substantially affected by errors induced by ancient DNA damage by performing PCA with a dataset excluding CpG sites (Figure S15). We evaluate this result as visually similar in layout to the non-zoomed PCA created when CpG sites are included (Figure S14), despite a reverse orientation of samples along PC1 .

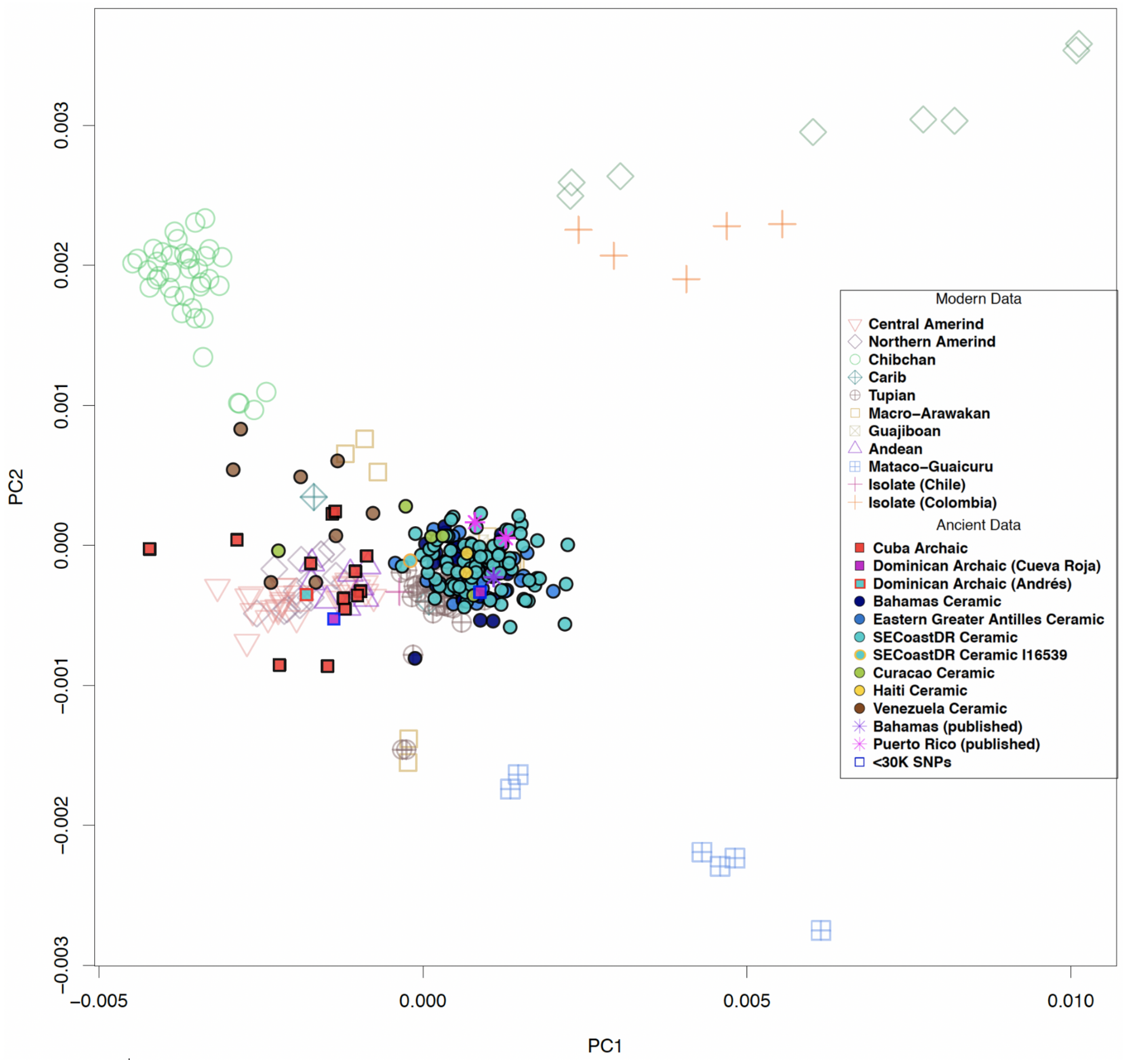

**Figure S15:** Full PCA (as in Figure S14) excluding CpG sites.

#### SI5 - Unsupervised population structure analysis

We performed unsupervised population structure analysis using ADMIXTURE (Alexander et al.; Alexander and Lange 2011) using a reference panel of 286 modern individuals from populations spread across the world, and three published ancient individuals from the Caribbean with genome-wide data (Schroeder et al. 2018; Nieves-Colón et al. 2020). We display K=5 in the main manuscript (Fig. 2b), selected based on the following table of cross-validation errors for five replicates of K=2-10 (Table S1).

**Table S1**: Cross-validation errors for five replicates of K=2-10.

| | | Run 1 | Run 2 | Run 3 | Run 4 | Run 5 |
|---|---|---|---|---|---|---|
| K value | 2 | 0.66416 | 0.66417 | 0.66410 | 0.66411 | 0.66412 |
| | 3 | 0.65713 | 0.65722 | 0.65720 | 0.65902 | 0.65755 |
| | 4 | 0.65064 | 0.65237 | 0.65233 | 0.65049 | 0.65586 |
| | 5 | 0.64590 | 0.64576 | 0.64569 | 0.65390 | 0.64575 |
| | 6 | 0.64835 | 0.64857 | 0.64884 | 0.64871 | 0.64829 |
| | 7 | 0.64708 | 0.64707 | 0.64736 | 0.64660 | 0.64706 |
| | 8 | 0.64605 | 0.65209 | 0.65310 | 0.64556 | 0.64641 |
| | 9 | 0.65170 | 0.65229 | 0.65123 | 0.65184 | 0.65240 |
| | 10 | 0.65314 | 0.65249 | 0.65180 | 0.65278 | 0.65243 |

In Figure S16, we visualize results for each value of K between 2 and 10, using the data generated from the replicate that produced the lowest cross-validation error for each value. Here we provide a visualization for the full reference panel with colors of modern-day American populations corresponding to language groups in Fig. 3c of the main manuscript.

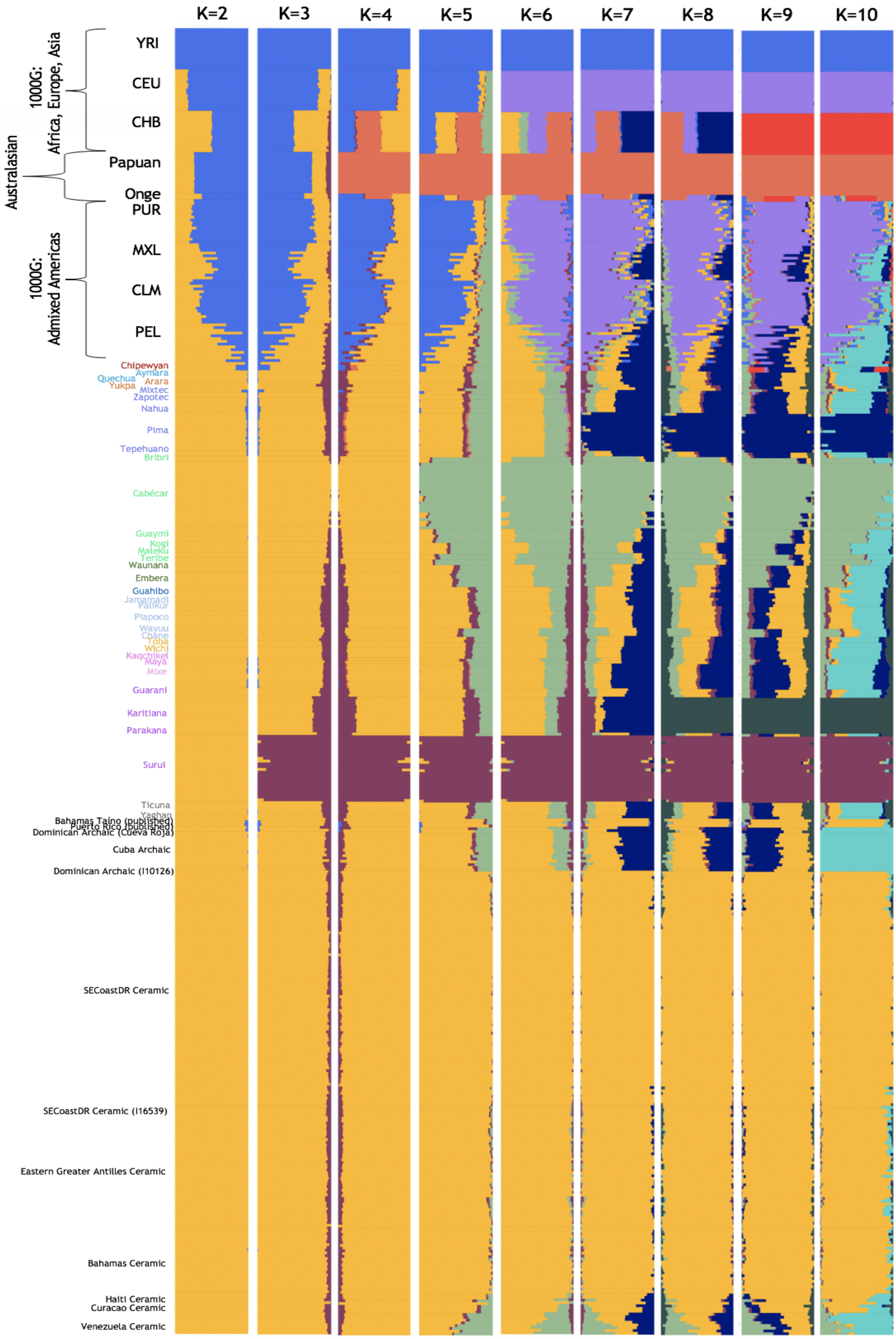

**Figure S16:** Results of model-based ancestry analysis using ADMIXTURE for values of K between 2 and 10.

#### SI6 - Kinship, Consanguinity, and Conditional Heterozygosity

##### Kinship analysis

We find that the majority of individuals who had a kin relationship (n=34) were from the site of La Caleta (Table S2). While this site has the highest number of individuals (n=62) of any site analyzed, we note that an unusually high proportion of individuals (~55%) share a kin relationship with another individual included in this study. Other sites also exhibited high frequencies of related individuals: we find that the two studied individuals from Diale 1 in Haiti were related ($2^{nd}$-degree relatives), as were all four individuals from the site of Macao in the Dominican Republic (1 pair of $2^{nd}$-degree relatives and two pairs of $2^{nd}$-$3^{rd}$-degree relatives). It is possible that this could be explained by non-random sampling (for example, selecting two individuals buried near to each other), as well as relatively small population sizes at these locales.

We identify a pair of $2^{nd}$-degree relatives with one individual buried at different sites in the southern Dominican Republic (I17906 is buried at Atajadizo, while I15601 is buried at La Caleta) (Fig. 4a). We note that I17906 also has a $1^{st}$-degree relative at Atajadizo (I17903), and I15601 also shares a $3^{rd}$-degree relationship with I17903. All three individuals were molecularly-sexed as males (Supplementary Data 2). While the numbers are too few to test for a statistically significant difference between behaviors of males and females, these results suggest that males engaged in intra-island mobility and connectivity.

##### Consanguinity analysis

ROH blocks longer than >12cM provide a clear indication of recent parental relatedness (up to 5 generations ago), as recombination quickly breaks up blocks back in time, making this signal independent of demographic processes deeper in time. In Figure S17 we visualize our results, grouped per archaeological site (data in Supplementary Data 10).

**Table S2:** Identified pairs of relatives per site. Number in parentheses is the number of individuals studied from the site. Gray cells identify inter-site relatives from La Caleta (I15601) and Atajadizo (I17903 and I17906).

| La Caleta (N=62) | | |
|---|---|---|
| I15590 | I16687 | 1st degree |
| I15050 | I16540 | 2nd degree |
| I15051 | I15604 | 2nd degree |
| I15082 | I16181 | 2nd degree |
| I15590 | I15598 | 2nd degree |
| I15598 | I16687 | 2nd degree |
| I15599 | I15964 | 2nd degree |
| I15675 | I15678 | 2nd degree |
| I15964 | I15977 | 2nd degree |
| I16172 | I16174 | 2nd degree |
| I16175 | I16181 | 2nd degree |
| I16520 | I16539 | 2nd degree |
| I16556 | I15681 | 2nd degree |
| I15051 | I16540 | 2nd-3rd degree |
| I15590 | I16520 | 2nd-3rd degree |
| I15594 | I16180 | 2nd-3rd degree |
| I15595 | I15678 | 2nd-3rd degree |
| I15597 | I15601 | 2nd-3rd degree |
| I15597 | I15602 | 2nd-3rd degree |
| I15598 | I16520 | 2nd-3rd degree |
| I15601 | I15602 | 2nd-3rd degree |
| I15604 | I15970 | 2nd-3rd degree |
| I15671 | I15678 | 2nd-3rd degree |
| I15671 | I15964 | 2nd-3rd degree |
| I15671 | I15965 | 2nd-3rd degree |
| I15671 | I15977 | 2nd-3rd degree |
| I15674 | I15973 | 2nd-3rd degree |
| I15965 | I15977 | 2nd-3rd degree |
| I15971 | I16173 | 2nd-3rd degree |
| I15977 | I16556 | 2nd-3rd degree |
| I16180 | I16540 | 2nd-3rd degree |
| I16520 | I16687 | 2nd-3rd degree |

| Canimar Abajo (N=13) | | |
|---|---|---|
| I17592 | I11170 | 1st degree |
| I10756 | I11170 | 2nd-3rd degree |

| Macao (N=4) | | |
|---|---|---|
| I7973 | I7976 | 2nd degree |
| I7972 | I7974 | 2nd-3rd degree |
| I7972 | I7976 | 2nd-3rd degree |

| Atajadizo (N=18) | | |
|---|---|---|
| I17903 | I17906 | 1st degree |

| Diale 1 (N=2) | | |
|---|---|---|
| I12575 | I12576 | 2nd degree |

| El Soco (N=12) | | |
|---|---|---|
| I13189 | I13203 | 1st degree |

| Sanctuary Blue Hole (N=7) | | |
|---|---|---|
| I13558 | I14883 | 2nd degree |

| Inter-Site: La Caleta - Atajadizo | | |
|---|---|---|
| I15601 | I17906 | 2nd degree |
| I15601 | I17903 | 3rd degree |

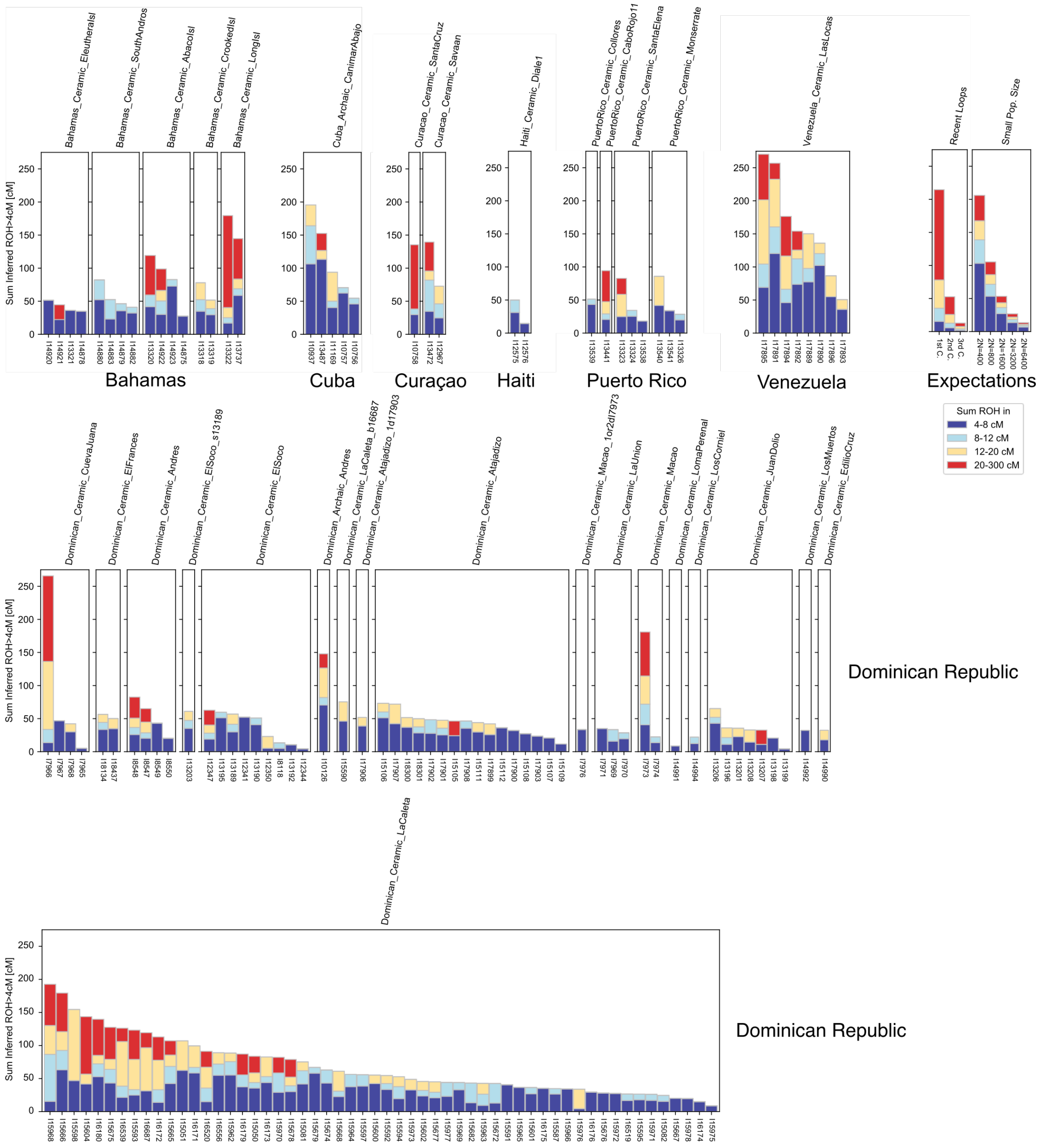

**Figure S17:** Runs of Homozygosity calls in ancient individuals. We depict inferred ROH for 154 ancient individuals grouped by location. Each bar represents one ancient individual, and we depict the total sums of ROH that fall into four length categories: 4-8, 8-12, 12-20, >20 cM for each individual (colored, stacked bars). The legend (lower right) shows analytical expectations, calculated using the formulas reported in Ringbauer et al. 2020. Letters “b”, “s”, and “1d” at the end of labels represent “brother of”, “sister of”, and “1st-degree relative of”, respectively.

We also depict the expected ROH for close cousin relationships and panmictic populations of small Ne, calculated using the formulas reported in Ringbauer et al. (2020), which show that the degree of inbreeding in La Caleta corresponds to the expectation for an effective population size of ~1000-1500 (census size could realistically be up to 10 times higher). We stress that for most populations, ancestry spreads geographically when tracing it backward in time (Ralph and Coop 2013), so effective population size values should be seen as comparison in an idealized case. Moreover, these are expectations, per realized individual there is considerable variation due to randomness of recombination (Ringbauer et al. 2020), for instance, 50-500 cM ROH in total are plausible for first cousins.

We also evaluated heterozygosity levels and found lower genetic diversity in the individuals from *GreaterAntilles_Archaic* and *Haiti_Ceramic* than in any of the other Ceramic-related (sub-)clades from the Greater Antilles, Curaçao, and Venezuela (Extended Data Fig. 2). These results agree with the median sums of ROH suggesting lower genetic diversity and a small population pool in the Archaic-related than in the Ceramic-related individuals. However, the Ceramic-associated Caribbean groups here studied were found to have an overall similar genetic diversity to that of some contemporary groups from continental South America, such as from the Peruvian Middle and Late Horizon periods (Nakatsuka et al. 2020).

**Maximum Likelihood Estimation of effective population size using Runs of Homozygosity**

We developed a method to fit the effective population size $N_e$ from observed ROH lengths, using the observed lengths of ROH blocks **l** = $l_1$, ...$l_n$ for a given set of individuals $i_1$.. $i_k$. The method is based on a maximum likelihood inference scheme, calculates the likelihood Pr(**l** | $N_e$) for the length distribution of a single individual and then estimates the population size that maximizes the product of the likelihoods. The likelihood is based on a well-known formula for the probability density of observing ROH (or IBD blocks) of length $x$, $b(x|t)$, between a pair of haplotypes originating from time t in the past; see Browning and Browning (2015) and Ringbauer, Coop, and Barton (2017) for details:

$$b(x|t)\,dx = \left( \underbrace{(G-x)(2t)^2 \exp(-2tx)\,dx}_{(i)} + \underbrace{2(2t)\exp(-2tx)\,dx}_{(ii)} \right) \psi(t) \tag{1}$$

Here $\psi(t)$ denotes the probability of coalescence $t$ generations ago, $G$ the length of the haplotype, and (i) describes blocks in the interior, amd (ii) blocks at one of the two ends of the haplotype. For a continuous panmictic population consisting of N haplotypes the coalescence probability time t ago, we plug in

$$\psi_N(t) = \exp\left(-\frac{t}{N}\right)\frac{1}{N}. \tag{2}$$

Integrating over all $t$ yields a straightforward analytical solution:

$$f_N(x)dx = \left(\frac{8(G-x)}{N}\frac{1}{(2x+\frac{1}{N})^3} + \frac{4}{N}\frac{1}{(2x+\frac{1}{N})^2}\right)dx \tag{3}$$

For multiple chromosomes, this formula needs to be summed over all chromosomes. For each small length intervall $\Delta x$, the product $f_N(x)\Delta x$ gives the expected number of ROH within this length interval. Following the approach of Ralph and Coop (2013), applied and explained in detail in (Ringbauer, Coop, and Barton 2017), we arrive at an approximate likelihood by binning shared blocks into small length bins $x_1, \ldots, x_l$ of width $\Delta x$ by modelling ROH counts in each of these bins as independent Poisson counts with expected rates $f_N(x_i)\Delta x$. We then calculate a composite likelihood of the observed data given $N_e$ using this Poisson model, and computationally maximize this overall likelihood. Estimates for the uncertainty of the estimator can be obtained via the curvature of the likelihood function (Fisher information matrix), or also via bootstrap over individuals or chromosomes. An implementation of this method is available via the Python package *hapROH* (https://test.pypi.org/project/hapROH/), which uses speed ups of the calculations explained in Ringbauer, Coop, and Barton (2017) that finish optimization and calculation for ROH of 10 individuals within much less than a second on a standard CPU.

**Testing the $N_e$ Inference Method**

To test the inference method, we generated simulated ROH for a panmictic population. We used the software *msprime* (Kelleher, Etheridge, and McVean 2016) and simulated all autosomes, with lengths determined between the genetic map difference between the first and last 1240k SNP on each autosome (as used in our inference scheme). We simulated four population sizes (2N=500, 1000, 2000, 4000) and validated the simulations by comparing the averages to our analytical expectations (obtained from integrating Formula (3); Figure S18). We then ran our inference method on genome-wide ROH of groups of 10 individuals, testing two scenarios: data where ROH blocks are i) determined by each true recombination event (recorded using the full ARG), and ii) by continuous tracts of co-ancestry within the last 100 generations - that merge “ineffective recombination events” and are detected in empirical data as continuous stretches (Chiang, Ralph, and Novembre 2016). Our results demonstrate that our method can robustly recover the population size used in the simulations. Moreover, the confidence intervals (obtained from the Fisher Information) accurately reflect the estimator uncertainty (Figure S19). Merging non-detectable recombination events starts having an observable effect for $2N_e$ below 1000, causing a downward bias of estimates (as more ROH are expected than expected from true recombination). Above $2N_e$=1000 this effect is negligible, and down to circa $2N_{e.}$=500 the observed bias remains small (<20%). For the empirical analysis, we filtered individuals who are possibly offspring of closely related parents, with more than 50cM of their genome in ROH>20cM blocks.

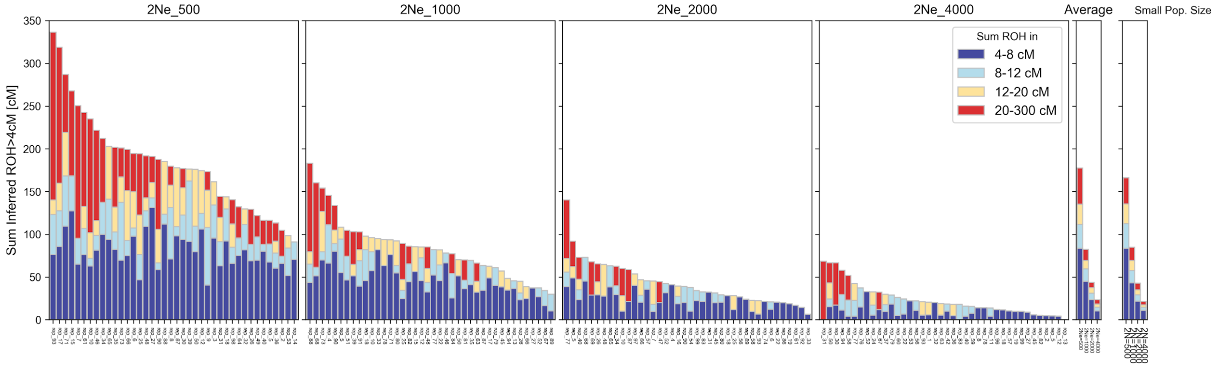

**Figure S18: Simulated ROH for four population sizes**. We visualize simulated ROH distribution on all autosomes. Each bar is ROH of one individual (40 per population size). The "average" panel gives the empirical average for each of these groups, and the "Small Pop. Size" panel gives the analytical average calculated from formula (3).

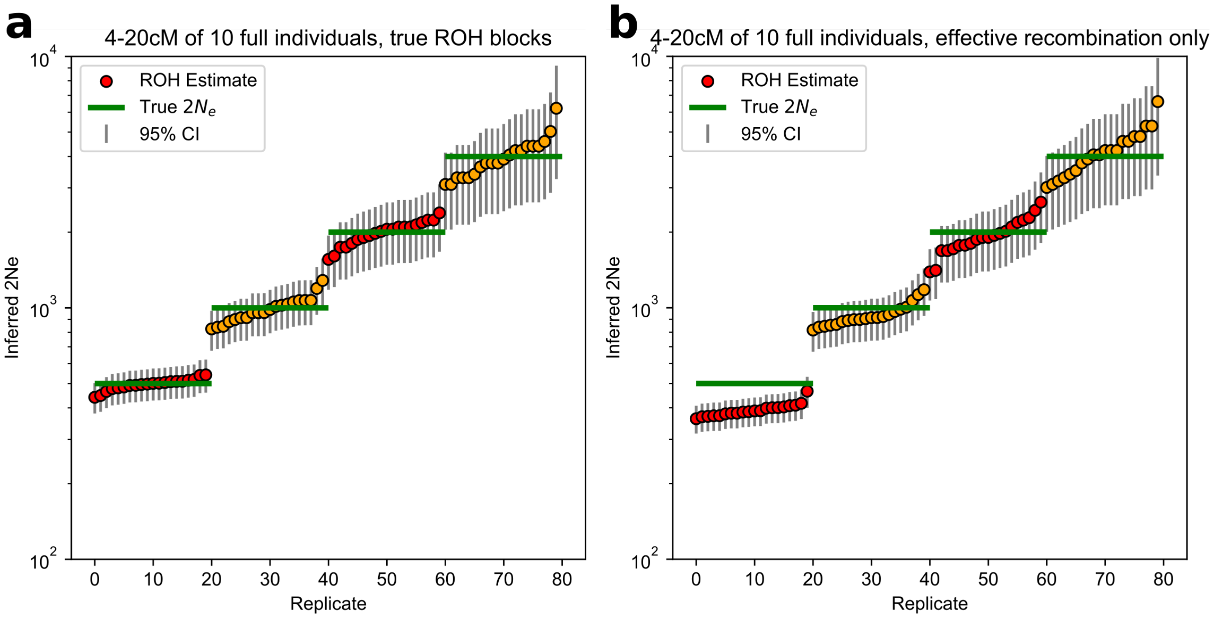

**Figure S19: Inferred Ne for various population sizes.** We inferred Ne using 10 individuals for 2N= 500, 1000, 2000 and 4000 (green lines). Each dot represents one replicate experiment of simulation and inferred data, vertical bars represent error bars. **(a)** Using all ROH defined by all recombination breakpoints (using the ARG). **(b)** ROH is defined as stretches with coalescent times less than 100 generations ago.

#### SI7 - Clade Grouping and Substructure Analysis with *qpWave, Treemix* and *$f_4$-statistics*

We used a five-step framework involving *qpWave*, Treemix, and *$f_4$*-statistics to group sites and individuals, and considered this information together with admixture profiles and proportions from *qpAdm* (see Supplementary Information section 8) to create groupings for genetic analysis independent from archaeologically-based material culture assignments. We recognize that people can be grouped in many different ways, and that these imposed groups may or may not reflect the true nature of organization in the past. However, as many genetic analyses require genetic information from more than one individual, we systematically grouped people together to facilitate analyses that require us to distinguish people who were relatively genetically closer to each other from people who were more genetically distinct. We emphasize that all groups or sub-clades presented in this study are composed of individuals from a broadly homogeneous genetic pool, also demonstrated by low pairwise $F_{ST}$ distances (Extended Data Fig. 1), and they represent a gradient of genetic relationships that may change with additional data that provides increased analytical power.

##### Step 1: Initial assignment of major clades with *qpWave*

For the first step, we ran pairwise *qpWave* tests between all sites included in this study, treating sites as units based on observed intra-site homogeneity in PCA and ADMIXTURE analyses (Fig. 2a, b; Figure S20). One outlier individual was identified through PCA, ADMIXTURE, and direct C14 dating. This individual, I10126 from Andrés, Dominican Republic, was directly dated to the Archaic Age and had a genomic profile clearly different from all other individuals from the predominantly Ceramic-associated Andrés site; for these reasons, I10126 was analysed independently. In Step 1, we identified three major clades, where all sites or individuals (in cases where a site was represented by only a single individual) within each clade were statistically consistent with being descended from an ancestral population that was genetically homogeneous since their separation from every other clade:

A) *GreaterAntilles_Archaic* - composed of a site (Canímar Abajo) from Cuba and I10126;

B) *Caribbean_Ceramic* - composed of sites from The Bahamas, Haiti, Dominican Republic, Puerto Rico, and Curaçao (although the high number of failed comparisons might suggest that the latter might not be a clade with a pool of the others, discussed in greater detail below and in Supplementary Information section 8);

C) *Venezuela_Ceramic* - composed of a site (Las Locas) from Venezuela.

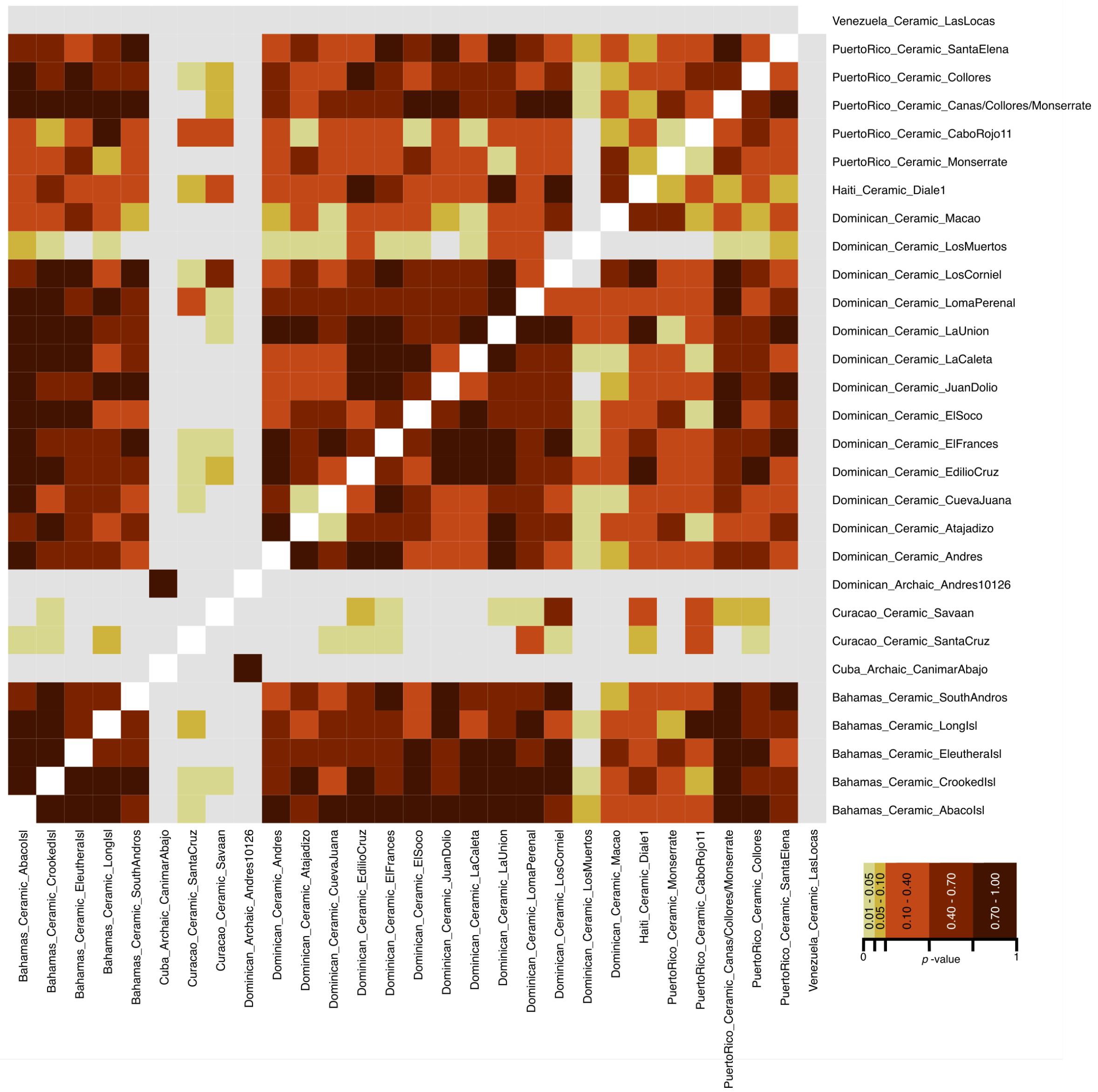

**Figure S20:** Pairwise *qpWave* Step 1 results for all sites in this study, using a threshold of $p>0.01$. Colors represent $p$-values of *qpWave* tests.

##### Step 2: Exploring substructure within Step 1 clades using a model competition approach

Step two consisted of repetition of pairwise models between members of the three major clades, but this time using a "model competition" approach, using the related software *qpAdm*. With this approach, a group on the "Left" ("sources") is moved to the "Right" ("references") if it is not currently being used on the "Left" (Lazaridis et al. 2016, Narasimhan et al. 2019, Fernandes et al. 2020). Sites were merged if they formed an exclusive clade with each other. Following this rule, we merged the individuals from Cuba (all of whom were from the same Archaic-associated site) with I10126 (who had an Archaic ancestry profile and was directly dated to the Archaic Age) to form *GreaterAntilles_Archaic* (intentionally maintaining the same name as the Step 1 clade; Figure S21). The individuals from two Ceramic-associated sites from Curaçao (de Savaan and Santa Cruz) did not form a clade with any of the other individuals from other sites in the *Caribbean_Ceramic* clade, but were in principle consistent with forming a clade with each other (p=0.017), so they were merged as *Curacao_Ceramic* and removed from *Caribbean_Ceramic*. In Supplementary Information section 8 we show how this behavior is explained by the presence of admixture in Curaçao. Individuals from the site of Diale 1 in Haiti also did not form a clade with any other site (again due to the presence of admixture), and were therefore merged them as *Haiti_Ceramic* and removed them from the *Caribbean_Ceramic* clade.

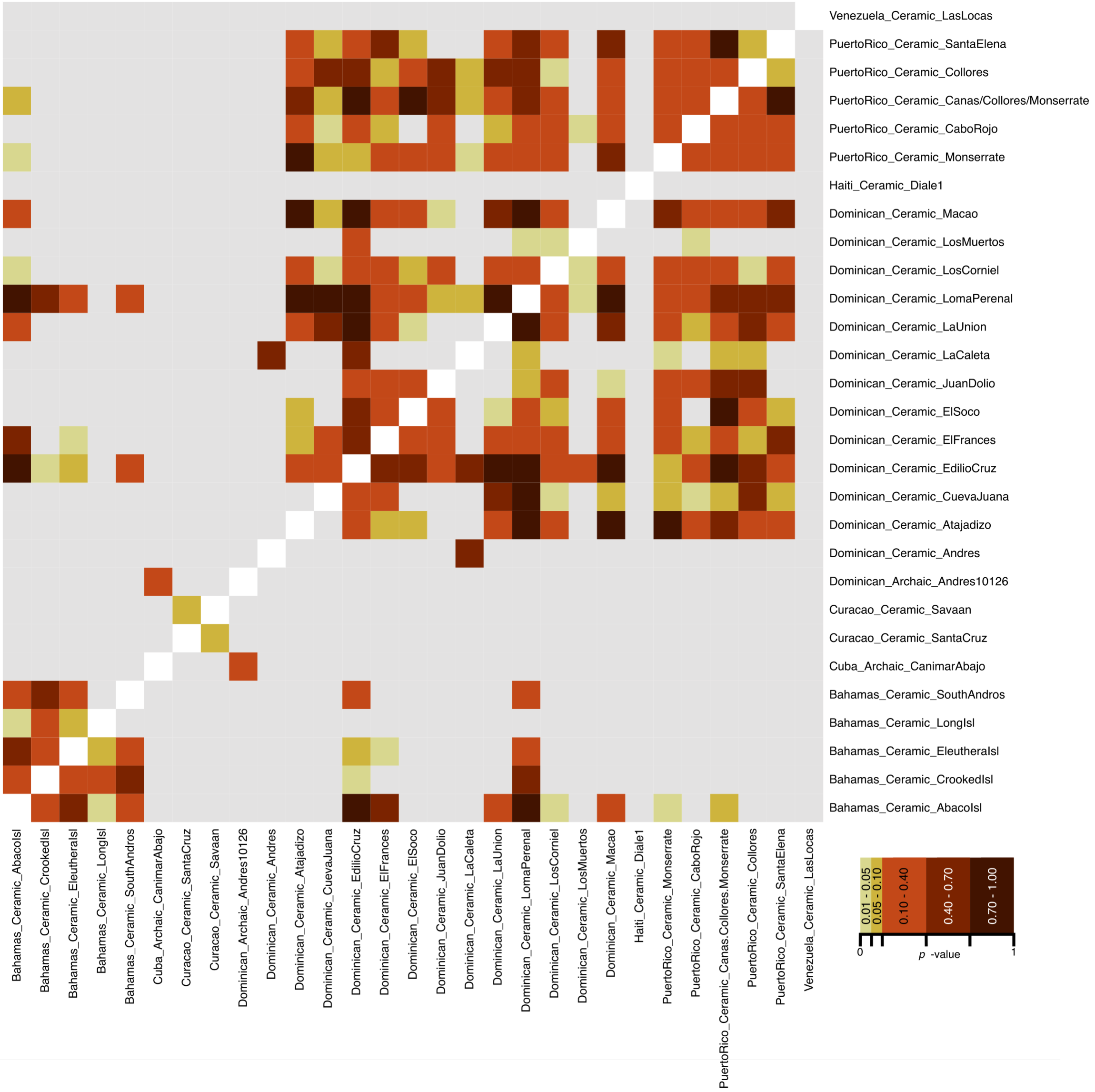

**Figure S21:** Pairwise *qpWave* Step 2 results, as a repetition of the tests from Step 1, but now using a model competition approach. Threshold of $p$>0.01 used. Colors represent $p$-values of *qpWave* tests.

**Step 3: Exploring structure using Treemix in *Caribbean_Ceramic***

For step three, we investigated structure within *Caribbean_Ceramic* (the only major clade composed of more than 2 sites). We first ran Treemix on the populations from the Illumina dataset allowing no migration events (Figure S22a). The residuals indicated a non-optimal fit for *Haiti_Ceramic* (Figure S22b) so we re-ran Treemix allowing up to as many admixture events as necessary until admixture was modelled into *Haiti_Ceramic* (Figures S22, S23). At "-m 1" Treemix identified a migration event between *Haiti_Ceramic* and *GreaterAntilles_Archaic* and the maximum residuals were reduced from 13.9 to 5.1 standard errors (Figure S23).

Next, we identified which sites were placed at a consistent location in the two tree fits (representing 0-1 migration events) relative to the other ancient sites, and explored if they formed statistically significant clades using $f_4$-statistics. Those sites were:

- All of The Bahamas sites/islands (forming a multi-site sub-group);

- La Caleta, Andrés, Juan Dolio, and El Soco (forming a multi-site sub-group);

- Cueva Juana

The majority of the sites that moved between different locations in the trees when we altered the number of migration events comprised single individuals and therefore have a lower resolution.

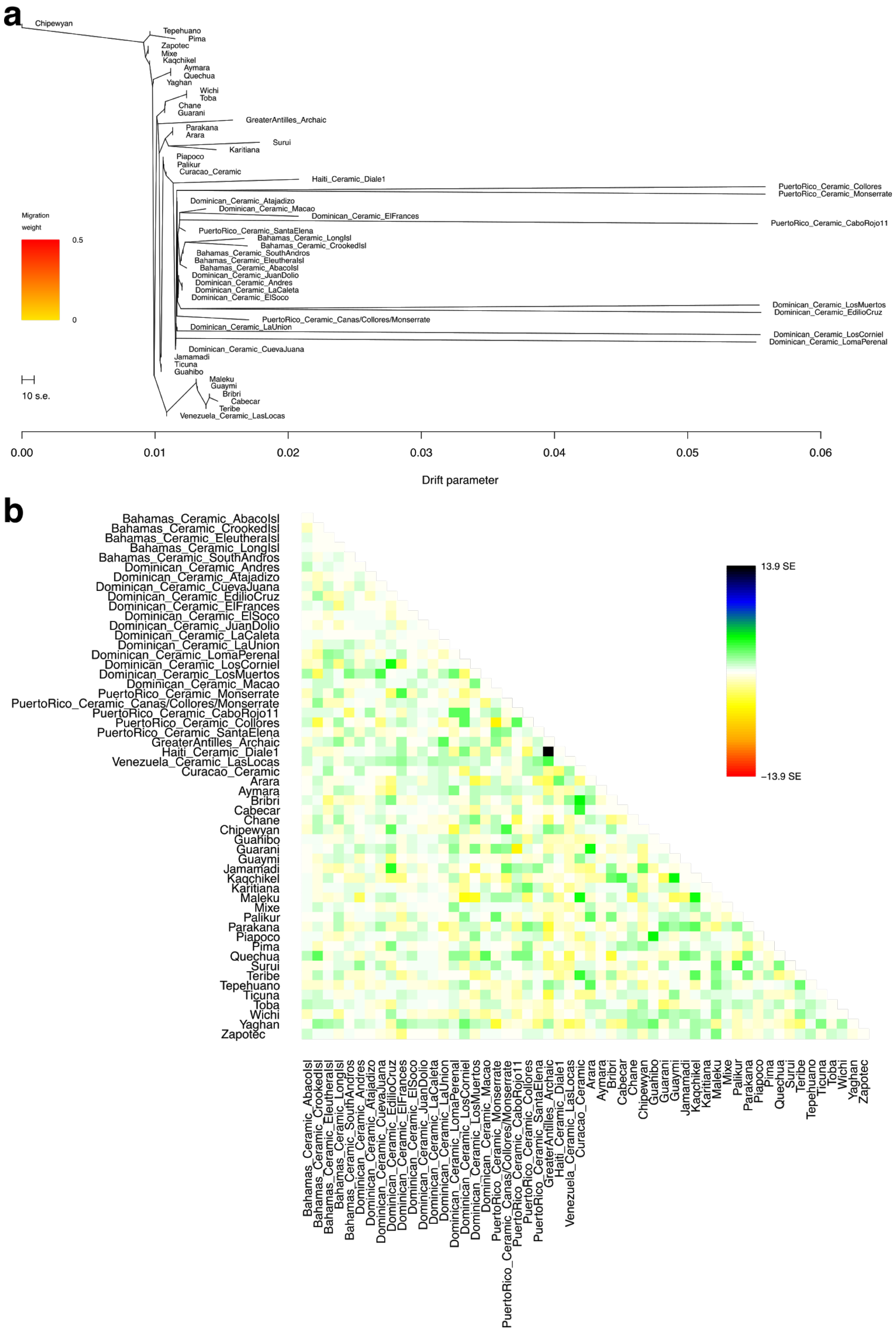

**Figure S22:** Treemix results with no migration/admixture events allowed (a), and corresponding fit's residuals (b).

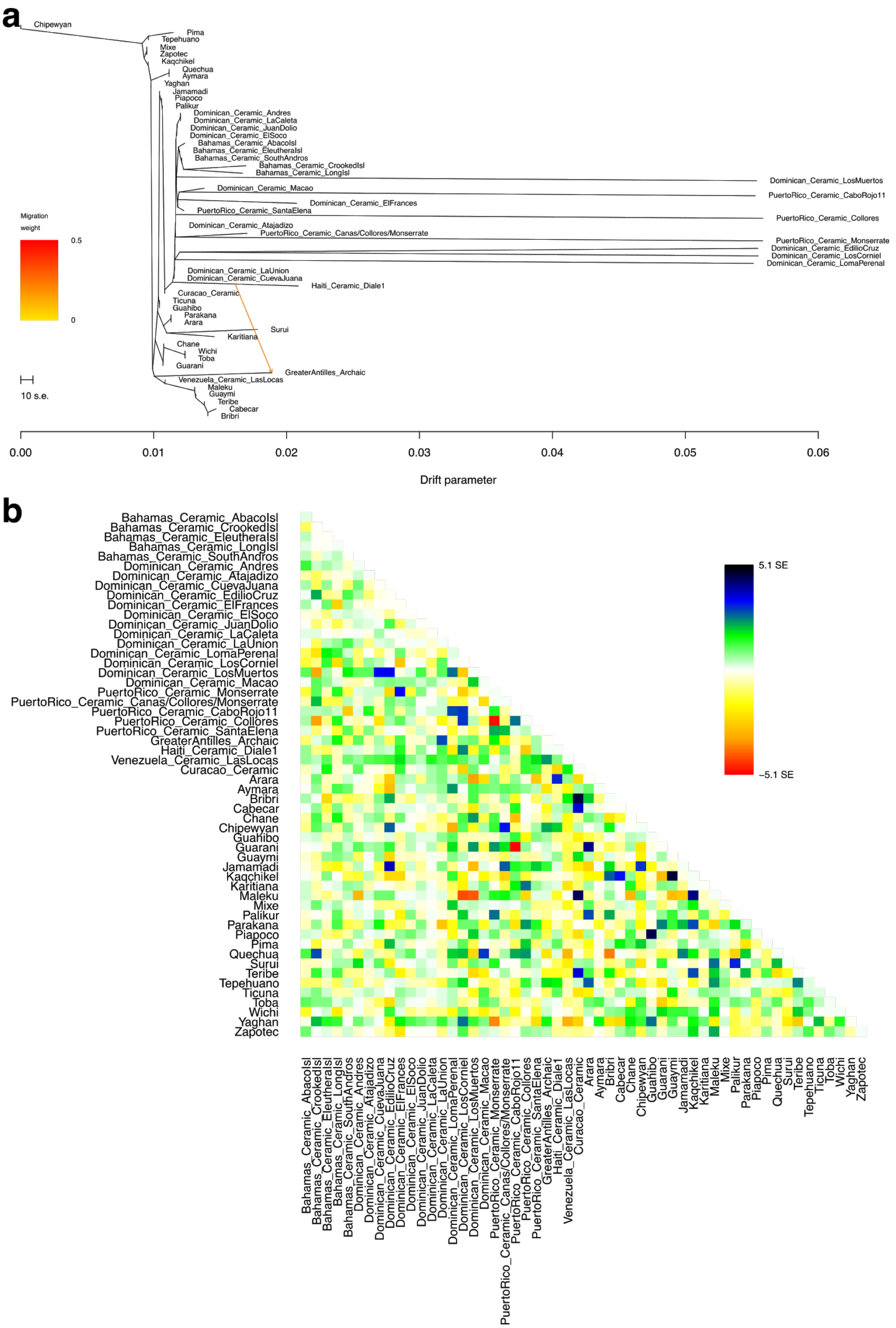

**Figure S23:** Treemix results with 1 migration/admixture events allowed (a), and corresponding fit's residuals (b).

**Step 4: Using $f_4$-statistics to confirm Treemix's structure**

For each unchanged branch identified in step 3 we performed *f*-statistics to evaluate if they formed clades to the exclusion of pools of other sites, hence suggesting structure within the *Caribbean_Ceramic* clade.

All tests involving islands from The Bahamas produced a |Z|<2.8 for the first test $f_4$(*Mbuti, Pool; Test1; Test2*) and |Z|>2.8 for at least one of the other two $f_4$-statistics (as described above and in Table S3), supporting their formation of a sub-clade within *Caribbean_Ceramic*. We therefore merged all sites from the Bahamas into *Bahamas_Ceramic*. Some possible structure is also observed for the sites from the Crooked and Long islands, as they are shown to be closer to each other than to some of the other sites from the Bahamas (Pools 1, 2, and 3 in the corresponding section of Table S3).

The Dominican sites of La Caleta and Andrés also clearly formed a sub-clade to the exclusion of the other tested nodes/pools, whereas we found contradictory results for El Soco and Juan Dolio, with El Soco sharing significantly more alleles with Juan Dolio than with the node containing La Caleta and Andrés (|Z| = 3.462, Table S3). All four of these sites are located within a 50 kilometer stretch of coast in the southeast part of the Dominican Republic (Figure S10). We therefore further explored the relationship between these sites with $f_4$-statistics of the form $f_4$(*Mbuti, Site1; Site2, PoolOfAllOtherCaribbean_Ceramic*) on all possible permutations between the sites. The results show a west-to-east cline of relatedness (the sites from west to east are La Caleta, Andrés, Juan Dolio, and El Soco) as statistically significant tests (|Z|>2.8) were found for the pairs La Caleta-Andrés, La Caleta-Juan Dolio, Andrés-Juan Dolio, and Juan Dolio-El Soco (Table S4). While the three westernmost sites, La Caleta, Andrés, and Juan Dolio, always shared more alleles with each other than with the remaining *Caribbean_Ceramic* sites, the easternmost site, El Soco, only produced statistically significant results with the site geographically closest to it (Juan Dolio, located 22 kilometers to the west) (Table S4). Even though the results between El Soco and La Caleta or Andrés were not statistically significant (|Z| = 2.2 and 0.8, respectively), we merged the four sites as *SECoastDR_Ceramic* due to their geographical proximity and the existence of a genetic cline connecting El Soco with the remaining sites.

**Table S3**: $f_4$-statistics Z-scores of the triple tests for the branches with unchanged relative location as identified in step 3. Tests of the form *$f_4$(Mbuti, Pool; Test1, Test2)*, *$f_4$(Mbuti, Test1; Pool, Test2)*, *$f_4$(Mbuti, Test2; Test1, Pool)*, with respective Z-scores shown as Z1, Z2, and Z3.

| | Test1 | Test2 | Z-score | |
|---|---|---|---|---|
| | **Bahamas_Ceramic_CrookedIsl** | **Bahamas_Ceramic_LongIsl** | | |
| **Pool1** | Bahamas_Ceramic_SouthAndros | | 1.986<br>3.378<br>-1.295 | Z1<br>Z2<br>Z3 |
| **Pool2** | Bahamas_Ceramic_EleutheraIsl | | 1.793<br>3.627<br>-1.840 | Z1<br>Z2<br>Z3 |
| **Pool3** | Bahamas_Ceramic_AbacoIsl | | 1.777<br>3.732<br>-2.055 | Z1<br>Z2<br>Z3 |
| **Pool4** | Dominican_Ceramic_Macao, Dominican_Ceramic_ElFrances, PuertoRico_Ceramic_CaboRojo11, Dominican_Ceramic_LosMuertos, Dominican_Ceramic_JuanDolio, Dominican_Ceramic_LaCaleta, Dominican_Ceramic_Andres, Dominican_Ceramic_ElSoco, Dominican_Ceramic_Atajadizo, PuertoRico_Ceramic_SantaElena, PuertoRico_Ceramic_Canas/Collores/Monserrate, PuertoRico_Ceramic_Monserrate, Dominican_Ceramic_LosCorniel, Dominican_Ceramic_EdilioCruz, PuertoRico_Ceramic_Collores, Dominican_Ceramic_LomaPerenal, Dominican_Ceramic_LaUnion, Dominican_Ceramic_CuevaJuana | | 1.623<br>6.897<br>-5.099 | Z1<br>Z2<br>Z3 |
| | **Test1** | **Test2** | | |
| | **Bahamas_Ceramic_SouthAndros** | **Bahamas_Ceramic_CrookedIsl, Bahamas_Ceramic_LongIsl** | | |
| **Pool1** | Bahamas_Ceramic_EleutheraIsl | | 0.612<br>1.797<br>-1.175 | Z1<br>Z2<br>Z3 |
| **Pool2** | Bahamas_Ceramic_AbacoIsl | | -0.051<br>0.996<br>-1.011 | Z1<br>Z2<br>Z3 |
| **Pool3** | Dominican_Ceramic_Macao, Dominican_Ceramic_ElFrances, PuertoRico_Ceramic_CaboRojo11, Dominican_Ceramic_LosMuertos, Dominican_Ceramic_JuanDolio, Dominican_Ceramic_LaCaleta, Dominican_Ceramic_Andres, Dominican_Ceramic_ElSoco, Dominican_Ceramic_Atajadizo, PuertoRico_Ceramic_SantaElena, PuertoRico_Ceramic_Canas/Collores/Monserrate, PuertoRico_Ceramic_Monserrate, Dominican_Ceramic_LosCorniel, Dominican_Ceramic_EdilioCruz, PuertoRico_Ceramic_Collores, Dominican_Ceramic_LomaPerenal, Dominican_Ceramic_LaUnion, Dominican_Ceramic_CuevaJuana | | 1.323<br>6.275<br>-4.815 | Z1<br>Z2<br>Z3 |
| | **Test1** | **Test2** | | |
| | **Bahamas_Ceramic_EleutheraIsl** | **Bahamas_Ceramic_SouthAndros, Bahamas_Ceramic_CrookedIsl, Bahamas_Ceramic_LongIsl** | | |
| **Pool1** | Bahamas_Ceramic_AbacoIsl | | 0.747<br>0.576<br>0.121 | Z1<br>Z2<br>Z3 |
| **Pool2** | Dominican_Ceramic_Macao, Dominican_Ceramic_ElFrances, PuertoRico_Ceramic_CaboRojo11, Dominican_Ceramic_LosMuertos, Dominican_Ceramic_JuanDolio, Dominican_Ceramic_LaCaleta, Dominican_Ceramic_Andres, Dominican_Ceramic_ElSoco, Dominican_Ceramic_Atajadizo, PuertoRico_Ceramic_SantaElena, PuertoRico_Ceramic_Canas/Collores/Monserrate, PuertoRico_Ceramic_Monserrate, Dominican_Ceramic_LosCorniel, Dominican_Ceramic_EdilioCruz, PuertoRico_Ceramic_Collores, Dominican_Ceramic_LomaPerenal, Dominican_Ceramic_LaUnion, Dominican_Ceramic_CuevaJuana | | 0.853<br>6.244<br>-4.111 | Z1<br>Z2<br>Z3 |
| | **Test1** | **Test2** | | |
| | **Bahamas_Ceramic_AbacoIsl** | **Bahamas_Ceramic_SouthAndros, Bahamas_Ceramic_EleutheraIsl, Bahamas_Ceramic_LongIsl, Bahamas_Ceramic_CrookedIsl** | | |
| **Pool1** | Dominican_Ceramic_Macao, Dominican_Ceramic_ElFrances, PuertoRico_Ceramic_CaboRojo11, Dominican_Ceramic_LosMuertos, Dominican_Ceramic_JuanDolio, Dominican_Ceramic_LaCaleta, Dominican_Ceramic_Andres, Dominican_Ceramic_ElSoco, Dominican_Ceramic_Atajadizo, PuertoRico_Ceramic_SantaElena, PuertoRico_Ceramic_Canas/Collores/Monserrate, PuertoRico_Ceramic_Monserrate, Dominican_Ceramic_LosCorniel, Dominican_Ceramic_EdilioCruz, PuertoRico_Ceramic_Collores, Dominican_Ceramic_LomaPerenal, Dominican_Ceramic_LaUnion, Dominican_Ceramic_CuevaJuana | | -0.357<br>5.126<br>-4.029 | Z1<br>Z2<br>Z3 |
| | **Test1** | **Test2** | | |
| | **Dominican_Ceramic_LaCaleta** | **Dominican_Ceramic_Andres** | | |
| **Pool1** | Dominican_Ceramic_JuanDolio | | 2.244<br>3.082<br>-1.076 | Z1<br>Z2<br>Z3 |
| **Pool2** | Dominican_Ceramic_ElSoco | | 1.309<br>4.313<br>-3.864 | Z1<br>Z2<br>Z3 |
| **Pool3** | Dominican_Ceramic_LosMuertos, Bahamas_Ceramic_SouthAndros, Bahamas_Ceramic_EleutheraIsl, Bahamas_Ceramic_LongIsl, Bahamas_Ceramic_CrookedIsl, Bahamas_Ceramic_AbacoIsl, Dominican_Ceramic_Macao, PuertoRico_Ceramic_CaboRojo11, Dominican_Ceramic_ElFrances, Dominican_Ceramic_Atajadizo, PuertoRico_Ceramic_SantaElena, PuertoRico_Ceramic_Canas/Collores/Monserrate, PuertoRico_Ceramic_Monserrate, Dominican_Ceramic_LosCorniel, Dominican_Ceramic_EdilioCruz, PuertoRico_Ceramic_Collores, Dominican_Ceramic_LomaPerenal, Dominican_Ceramic_LaUnion, Dominican_Ceramic_CuevaJuana | | 2.288<br>6.440<br>-6.837 | Z1<br>Z2<br>Z3 |
| | **Test1** | **Test2** | | |
| | **Dominican_Ceramic_JuanDolio** | **Dominican_Ceramic_LaCaleta, Dominican_Ceramic_Andres** | | |
| **Pool1** | Dominican_Ceramic_ElSoco | | -3.462<br>-2.014<br>-1.523 | Z1<br>Z2<br>Z3 |
| **Pool2** | Dominican_Ceramic_LosMuertos, Bahamas_Ceramic_SouthAndros, Bahamas_Ceramic_EleutheraIsl, Bahamas_Ceramic_LongIsl, Bahamas_Ceramic_CrookedIsl, Bahamas_Ceramic_AbacoIsl, Dominican_Ceramic_Macao, PuertoRico_Ceramic_CaboRojo11, Dominican_Ceramic_ElFrances, Dominican_Ceramic_Atajadizo, PuertoRico_Ceramic_SantaElena, PuertoRico_Ceramic_Canas/Collores/Monserrate, PuertoRico_Ceramic_Monserrate, Dominican_Ceramic_LosCorniel, Dominican_Ceramic_EdilioCruz, PuertoRico_Ceramic_Collores, Dominican_Ceramic_LomaPerenal, Dominican_Ceramic_LaUnion, Dominican_Ceramic_CuevaJuana | | -2.693<br>1.100<br>-3.542 | Z1<br>Z2<br>Z3 |

| | Test1 | Test2 |
|---|---|---|

| | | | |
|---|---|---|---|
| **Dominican_Ceramic_ElSoco** | **Dominican_Ceramic_JuanDolio, Dominican_Ceramic_LaCaleta, Dominican_Ceramic_Andres** | | |
| **Pool1** | Dominican_Ceramic_LosMuertos, Bahamas_Ceramic_SouthAndros, Bahamas_Ceramic_EleutheraIsl, Bahamas_Ceramic_LongIsl, Bahamas_Ceramic_CrookedIsl, Bahamas_Ceramic_AbacoIsl, Dominican_Ceramic_Macao, PuertoRico_Ceramic_CaboRojo11, Dominican_Ceramic_ElFrances, Dominican_Ceramic_Atajadizo, PuertoRico_Ceramic_SantaElena, PuertoRico_Ceramic_Canas/Collores/Monserrate, PuertoRico_Ceramic_Monserrate, Dominican_Ceramic_LosCorniel, Dominican_Ceramic_EdilioCruz, PuertoRico_Ceramic_Collores, Dominican_Ceramic_LomaPerenal, Dominican_Ceramic_LaUnion, Dominican_Ceramic_CuevaJuana | -1.932<br>0.197<br>-2.166 | Z1<br>Z2<br>Z3 |

**Table S4**: Testing the relationship between the 4 coastal sites from southeastern Dominican Republic, against a pool of other *Caribbean_Ceramic* sites.

| Pop X | Pop Y | Pop W | Pop Z | f4 | SE | Z | SNPs |
|---|---|---|---|---|---|---|---|
| *Mbuti* | *Dominican_Ceramic_Andres* | *Dominican_Ceramic_LaCaleta* | *Ceramic_Pool* | -0.000605 | 0.000088 | -6.837 | 982655 |
| *Mbuti* | *Dominican_Ceramic_Andres* | *Dominican_Ceramic_ElSoco* | *Ceramic_Pool* | -0.000102 | 0.000125 | -0.811 | 972718 |
| *Mbuti* | *Dominican_Ceramic_Andres* | *Dominican_Ceramic_JuanDolio* | *Ceramic_Pool* | -0.000453 | 0.000143 | -3.181 | 955522 |
| *Mbuti* | *Dominican_Ceramic_LaCaleta* | *Dominican_Ceramic_ElSoco* | *Ceramic_Pool* | -0.000212 | 0.000099 | -2.150 | 1061854 |
| *Mbuti* | *Dominican_Ceramic_LaCaleta* | *Dominican_Ceramic_JuanDolio* | *Ceramic_Pool* | -0.000393 | 0.000113 | -3.493 | 1023549 |
| *Mbuti* | *Dominican_Ceramic_ElSoco* | *Dominican_Ceramic_JuanDolio* | *Ceramic_Pool* | -0.000424 | 0.000129 | -3.280 | 1011045 |

The site composition of the final sub-clades created after these 4 steps are as follows, and are represented in Fig. 2c:

*GreaterAntilles_Archaic* - Canímar Abajo, Andrés (I10126)

*SECoastDR_Ceramic* - La Caleta, Andrés, Juan Dolio, El Soco

*Bahamas_Ceramic* - Abaco Island (Bill Johnson's Cave, Hopetown, Randy's Cave, Imperial Lighthouse Cave), Crooked Island (Burial Cave #1, Gordon Hill Cave, unknown site), Eleuthera Island (Garden Cave, Preacher's Cave, Valentine's Blue Hole, Wemyss Bight Cave), Long Island (Rolling Heads Site, Cave near Clarence Town), South Andros Island (Sanctuary Blue Hole, Stargate Blue Hole)

*EasternGreaterAntilles_Ceramic* - El Francés, Edilio Cruz, La Unión, Loma Perenal, Macao, Collores, Monserrate, Cabo Rojo 11, Santa Elena, Canas/Collores/Monserrate, Atajadizo, Cueva Juana, Los Corniel, Los Muertos

*Venezuela_Ceramic* - Las Locas

*Haiti_Ceramic* - Diale 1

*Curacao_Ceramic* - de Savaan, Santa Cruz

After identifying all sub-clades, we used the intra-sub-clade statistic $f_4$(*Mbuti*, *GreaterAntilles_Archaic*; *Individual*, *SubClade-Without-Individual*), with *Individual* as every individual from within a sub-clade, and *SubClade-Without-Individual* as their sub-clade minus the individual being tested, to assess if any individuals had more Archaic-related ancestry than the remainder of their sub-clade (significant results in Table S5, all results in Supplementary Data 6). For any statistically significant case (Z<-2.8) we removed the individual from the sub-clade and labeled them as their sub-clade plus their sample ID as a suffix (e.g. *Sub_Clade8888*). Any individuals determined to have excess Archaic-related ancestry were not included in further analyses of their sub-clade.

**Table S5**: Testing each individual's archaic-related affinities within each sub-clade. Results for |Z|>2.8 are presented; complete table found in Supplementary Data 6.

| Pop X | Pop Y | Pop W | Pop Z | f4 | SE | Z | SNPs |
|---|---|---|---|---|---|---|---|
| *Mbuti* | *GreaterAntilles_Archaic* | *SECoastDR_Ceramic 16539* | *SECoastDR_Ceramic _ excl_I16539* | -0.002158 | 0.000408 | -5.288 | 687900 |
| *Mbuti* | *GreaterAntilles_Archaic* | *EasternGreaterAntilles_ Ceramic7969* | *EasternGreaterAntilles_ Ceramic_excl_I7969* | -0.001054 | 0.000368 | -2.867 | 805059 |

As shown above, two individuals produced statistically significant statistics that suggested having higher levels of archaic-related ancestry when compared to their sub-clade: I16539 from the site of La Caleta in the *SECoastDR_Ceramic* sub-clade (Z = -5.288) and I7969 from the site of La Union in the *EasternGreaterAntilles_Ceramic* sub-clade (Z = -2.867). In Supplementary Information section 8, and before performing any other analyses, we estimated the amount of this Archaic-related ancestry in these two individuals using *qpAdm*. Because we were not able to reject the formation of a clade between I7969 and *EasternGreaterAntilles_Ceramic* (even with *GreaterAntilles_Archaic* on the "Right") we did not remove this individual from the sub-clade; however, I16539 was renamed as *SECoastDR_Ceramic16539*.

**Step 5: Exploring substructure with $f_4$-statistics**

We finally investigated the existence of substructure within the *Caribbean_Ceramic* sub-clades using $f_4$-statistics. Substructure was interpreted as present if a site produced statistically significant results (average pairwise Z-score > 2.8) with at least 50% of the remaining sites from that sub-clade.

For the *Bahamas_Ceramic* group, all islands had an average Z-score above 2.8 with at least 50% of the remaining sites, suggesting the presence of substructure in this clade (Supplementary Data 5).

For S*ECoastDR_Ceramic*, the individuals from La Caleta shared more alleles with each other than with individuals from El Soco and Juan Dolio, but not with individuals from the geographically more proximate site of Andrés. This further supports the genetic cline in this sub-clade (discussed above, with data in Table S4).

Within the large *EasternGreaterAntilles_Ceramic* sub-clade, 13 pairwise tests had a Z > 2.8, but only the Dominican site of Macao showed significant signs of substructure, according to the rules set above. We note that it is possible that this result is driven at least in part by the relatedness of the studied individuals from Macao, as we observe one pair of 2nd-degree relatives and two pairs of 2nd- or 3rd-degree relatives among the four individuals studied (Supplementary Information section 6).

For Fig. 2c we allowed as many migration/admixture events ('- m') as necessary until all the admixture events identified for our (sub-)clades in other analysis such as *f*-statistics and *qpAdm* (see sections below) were introduced by Treemix. This happened at '-m 5' where individual I16539 received admixture from GreaterAntilles_Archaic, reducing also the standard error of the residuals from |11| to |4.4|.

#### SI8 - Admixture Modelling and Estimates of Ancestry Proportions

We first used *qpAdm* to estimate the amount of Archaic-associated ancestry in individuals I16539 and I7969, both of whom showed higher affinities to *GreaterAntilles_Archaic* than the rest of their sub-clades (although marginally for I7969). We then attempted to understand if the exclusion of *Haiti_Ceramic* and *Curacao_Ceramic* from the major *Caribbean_Ceramic* clade could possibly be explained by admixture, after finding that these two sub-clades initially formed a clade with other ceramic-period sites in *qpWave*, but no longer grouped with them upon the application of the model competition approach. In the tables in this section "Anc_X" represents the proportion of ancestry from a source population, and "SE_X" the associated standard error.

##### *SECoastDR_Ceramic16539* and *EasternGreaterAntilles_Ceramic7969*

Two individuals from different sub-clades within the *Caribbean_Ceramic* major clade (I16539 and I7969) had evidence of significant proportions of archaic-related ancestry. *SECoastDR_Ceramic16539* falls on the PCA at the border of the *Caribbean_Ceramic* cluster (in the direction of the *GreaterAntilles_Archaic* cluster), and is identified in Fig. 2a as a PCA outlier. We started by confirming that these individuals formed a clade with the sub-clade they initially belonged to, and then added *GreaterAntilles_Archaic* to the "Right." If the latter contributed ancestry to the outlier individuals, it is expected that the previously-working model would now fail (i.e. $p<0.05$, although we allow a buffer zone between 0.01 and 0.05 in cases where standard errors are substantially reduced during model competition). We also tested both *Cuba_CanimarAbajo_Archaic* and *Andres_Archaic10126* separately as sources of Archaic-associated ancestry, with the source not used on the "Left" added to the "Right."

Individual I16539 was modeled as having between 12.1±2.0% and 14.8±3.0% Archaic-related ancestry, depending on the Archaic source used (*Andres_Archaic10126* and *Cuba_CanimarAbajo_Archaic*, respectively), and the remainder *SECoastDR_Ceramic*-associated (Table S6**)**. We report the model with the lower standard error in the main manuscript.

In contrast, we could not reject the 1-way model using only *EasternGreaterAntilles_Ceramic* as a source for I7969 (even with *GreaterAntilles_Archaic* on the "Right"), most likely due to a weak signal of 4.10±2.7% to 6.40±1.9% Archaic-related ancestry from *Andres_Archaic10126* and *Cuba_CanimarAbajo_Archaic*, respectively (Table S6). We therefore merged individual I7969 back into the *EasternGreaterAntilles_Ceramic* sub-clade for all subsequent analyses.

**Table S6**: Modeling the Ceramic- and Archaic-related ancestries of *SECoastDR_Ceramic16539* and *EasternGreaterAntilles_Ceramic7969*.

| *Test* | A | B | Sources on "Right" | p-value | Anc_A | Anc_B | SE_A | SE_B |
|---|---|---|---|---|---|---|---|---|
| *SECoastDR_Ceramic 16539* | *SECoastDR_Ceramic* | - | - | 0.890 | 1.000 | - | 0.000 | - |
| *SECoastDR_Ceramic 16539* | *SECoastDR_Ceramic* | - | *GreaterAntilles_Archaic* | 1.80E-05 | 1.000 | - | 0.000 | - |
| *SECoastDR_Ceramic 16539* | *SECoastDR_Ceramic* | *Cuba_CanimarAbajo _Archaic* | *Andres_Archaic10126* | 0.901 | 0.852 | 0.148 | 0.030 | 0.030 |
| *SECoastDR_Ceramic 16539* | *SECoastDR_Ceramic* | *Andres_Archaic10126* | *Cuba_CanimarAbajo _Archaic* | 0.836 | 0.879 | 0.121 | 0.020 | 0.020 |
| *EasternGreaterAntilles_ Ceramic7969* | *EasternGreaterAntilles_ Ceramic* | - | - | 0.756 | 1.000 | - | 0.000 | - |
| *EasternGreaterAntilles_ Ceramic7969* | *EasternGreaterAntilles_ Ceramic* | - | *GreaterAntilles_Archaic* | 0.151 | 1.000 | - | 0.000 | - |
| *EasternGreaterAntilles_ Ceramic7969* | *EasternGreaterAntilles_ Ceramic* | *Cuba_CanimarAbajo_ Archaic* | *Andres_Archaic10126* | 0.692 | 0.959 | 0.041 | 0.027 | 0.027 |
| *EasternGreaterAntilles_ Ceramic7969* | *EasternGreaterAntilles_ Ceramic* | *Andres_Archaic10126* | *Cuba_CanimarAbajo _Archaic* | 0.675 | 0.936 | 0.064 | 0.019 | 0.019 |

##### *Haiti_Ceramic*

We started by identifying the lowest-rank valid models using the three major clades as sources, and then individually added each of the remaining clades to the "Right." As above, if those clades contributed ancestry to the test population, it is expected that the previously-working model would now fail. A 1-way model for *Haiti_Ceramic* with *Caribbean_Ceramic* as the source was initially valid, but the addition of *GreaterAntilles_Archaic* to the "Right" caused it to fail, suggesting the presence of Archaic-related ancestry in *Haiti_Ceramic* (Table S7). Therefore, we added *GreaterAntilles_Archaic* as a source together with *Caribbean_Ceramic* and tested a 2-way model, which we found to be valid even with *Venezuela_Ceramic* on the "Right." The final admixture model for *Haiti_Ceramic* based on the 3 major clades/ancestries was between 85.0±8.2% and 89.4±8.1% *Caribbean_Ceramic*-related ancestry and 15.0±8.2% and 10.6±8.1% *GreaterAntilles_Archaic*-related ancestry (p=0.199; see below for more precise mixture proportion estimates). This admixture explains why they do not form a clade with any other group during the *qpWave* analysis.

**Table S7**: Modeling *Haiti_Ceramic* with three major ancestry components - *Caribbean_Ceramic*, *GreaterAntilles_Archaic*, and *Venezuela_Ceramic*.

| A | B | Sources on "Right" | p-value | Anc_A | Anc_B | SE_A | SE_B |
|---|---|---|---|---|---|---|---|
| *Caribbean_Ceramic* | - | - | 0.158 | 1.000 | - | 0.000 | - |
| *Caribbean_Ceramic* | - | *GreaterAntilles_Archaic* | 6.22E-19 | 1.000 | - | 0.000 | - |
| *Caribbean_Ceramic* | - | *Venezuela_Ceramic* | 0.079 | 1.000 | - | 0.000 | - |
| *Caribbean_Ceramic* | *GreaterAntilles_Archaic* | - | 0.334 | 0.850 | 0.150 | 0.082 | 0.082 |
| *Caribbean_Ceramic* | *GreaterAntilles_Archaic* | *Venezuela_Ceramic* | 0.199 | 0.894 | 0.106 | 0.081 | 0.081 |
| *GreaterAntilles_Archaic* | - | - | 1.795E-12 | 1.000 | - | 0.000 | - |
| *Venezuela_Ceramic* | - | - | 1.30E-21 | 1.000 | - | 0.000 | - |

We then explored the possibility of improving model fit using the specific sub-clades within the two major clades identified as sources for *Haiti_Ceramic* (*Caribbean_Ceramic* and *GreaterAntilles_Archaic*). 2-way models using the sub-clades *Bahamas_Ceramic*, *EasternGreaterAntilles_Ceramic*, and *SECoastDR_Ceramic* within *Caribbean_Ceramic* and either

*Cuba_CanimarAbajo_Archaic* or *Andres_Archaic10126*, produced valid models with p>0.083 (**Table S8**). The addition of the three unused sources to the "Right" of the working 2-way models caused the *p*-values to decrease to below the threshold of 0.05, but always above 0.01 for the models with *Cuba_CanimarAbajo_Archaic* as a source. Standard errors were also substantially reduced. The model with *EasternGreaterAntilles_Ceramic* as the Ceramic-related source, for example, had a 4X decrease in the standard errors and the highest *p*-value (0.045). We therefore consider these model competition results as successful but caveat that the p-value <0.05 might reflect less optimal models, despite the very low standard errors. In Fig. 2c we report the averages of the 3 model competition models above 0.01 (81.4±2.1% and 18.57±2.1%), as they all constitute the *Caribbean_Ceramic* clade.

**Table S8**: Modeling *Haiti_Ceramic* with groups from within the two major clades identified in Supplementary Information section 7.

| A | B | Sources on "Right" | p-value | Anc_A | Anc_B | SE_A | SE_B |
|---|---|---|---|---|---|---|---|
| *Bahamas_Ceramic* | *Cuba_CanimarAbajo_ Archaic* | - | 0.379 | 0.847 | 0.153 | 0.088 | 0.088 |
| *EasternGreaterAntilles _Ceramic* | *Cuba_CanimarAbajo_ Archaic* | - | 0.331 | 0.853 | 0.147 | 0.087 | 0.087 |
| *SECoastDR_Ceramic* | *Cuba_CanimarAbajo_ Archaic* | - | 0.387 | 0.843 | 0.157 | 0.085 | 0.085 |
| *Bahamas_Ceramic* | *Andres_Archaic10126* | - | 0.117 | 0.881 | 0.119 | 0.107 | 0.107 |
| *EasternGreaterAntilles _Ceramic* | *Andres_Archaic10126* | - | 0.083 | 0.877 | 0.123 | 0.105 | 0.105 |
| *SECoastDR_Ceramic* | *Andres_Archaic10126* | - | 0.111 | 0.871 | 0.129 | 0.100 | 0.100 |
| *Bahamas_Ceramic* | *Cuba_CanimarAbajo_ Archaic* | *EasternGreaterAntilles _Ceramic, SECoastDR_Ceramic, Andres_Archaic10126* | 0.027 | 0.809 | 0.191 | 0.021 | 0.021 |
| *Bahamas_Ceramic* | *Andres_Archaic10126* | *EasternGreaterAntilles _Ceramic, SECoastDR_Ceramic, Cuba_CanimarAbajo_ Archaic* | 0.003 | 0.817 | 0.183 | 0.016 | 0.016 |
| *EasternGreaterAntilles _Ceramic* | *Cuba_CanimarAbajo_ Archaic* | *Bahamas_Ceramic, SECoastDR_Ceramic, Andres_Archaic10126* | 0.045 | 0.814 | 0.186 | 0.021 | 0.021 |
| *EasternGreaterAntilles _Ceramic* | *Andres_Archaic10126* | *Bahamas_Ceramic, SECoastDR_Ceramic, Cuba_CanimarAbajo_ Archaic* | 0.006 | 0.818 | 0.182 | 0.015 | 0.015 |
| *SECoastDR_Ceramic* | *Cuba_CanimarAbajo_ Archaic* | *Bahamas_Ceramic, EasternGreaterAntilles _Ceramic, Andres_Archaic10126* | 0.036 | 0.820 | 0.180 | 0.021 | 0.021 |
| *SECoastDR_Ceramic* | *Andres_Archaic10126* | *Bahamas_Ceramic, EasternGreaterAntilles _Ceramic, Cuba_CanimarAbajo_ Archaic* | 0.012 | 0.820 | 0.180 | 0.015 | 0.015 |

##### *Curacao_Ceramic*

No 1-way models were valid for *Curacao_Ceramic*, but a 2-way model using *Caribbean_Ceramic* (68.1±5.1%) and *Venezuela_Ceramic* (31.9±5.1%) produced a good fit (p=0.317) (Table S9). The addition of the unused source, *GreaterAntilles_Archaic*, to the "Right" reduced the model's *p*-value to between 0.01 and 0.05, possibly due to excess allele sharing and/or gene flow with one of the source populations. This is shown by the highly positive *gendstats* results from *qpAdm*'s output for *GreaterAntilles_Archaic* (Z-score average=2.576, median=2.556, maximum=3.549, minimum=0.930), based on statistics of the form $f_4$(*Curacao_Ceramic*, *Caribbean_Ceramic+Venezuela_Ceramic*; *Right1*, *GreaterAntilles_Archaic*), where *Right1* are the 13 modern populations from the base "Right" set, and *Caribbean_Ceramic+Venezuela_Ceramic* represents a modeled mixture of the two sources. These results suggest some caution in the use of *GreaterAntilles_Archaic* as an outgroup in these tests, as *qpAdm* requires that gene flow between "Left" and "Right" populations occurs closer to the source populations than the mixture into the test population.

**Table S9**: Modeling *Curacao_Ceramic* with three major ancestry components - *Caribbean_Ceramic*, *GreaterAntilles_Archaic*, and *Venezuela_Ceramic*.

| A | B | Sources on "Right" | p-value | Anc_A | Anc_B | SE_A | SE_B |
|---|---|---|---|---|---|---|---|
| *Caribbean_Ceramic* | - | - | 2.17E-07 | 1.000 | - | 0.000 | - |
| *Caribbean_Ceramic* | *GreaterAntilles_Archaic* | - | 9.19E-06 | 0.742 | 0.258 | 0.108 | 0.108 |
| *Caribbean_Ceramic* | *Venezuela_Ceramic* | - | 0.317 | 0.681 | 0.319 | 0.051 | 0.051 |
| *Caribbean_Ceramic* | *Venezuela_Ceramic* | *GreaterAntilles_Archaic* | 0.031 | 0.704 | 0.296 | 0.051 | 0.051 |
| *GreaterAntilles_Archaic* | - | - | 6.22E-14 | 1.000 | - | 0.000 | - |
| *GreaterAntilles_Archaic* | *Venezuela_Ceramic* | - | 1.23E-13 | 1.108 | -0.108 | 0.355 | 0.355 |
| *Venezuela_Ceramic* | - | - | 8.16E-22 | 1.000 | - | 0.000 | - |

Using the final *Caribbean_Ceramic* sub-clades to improve model fits, we find that all fit equally well (Table S10), again consistent with the high levels of genetic homogeneity observed within the *Caribbean_Ceramic* clade. With model competition the only model in the buffer zone of 0.01-0.05 is the one for *EasternGreaterAntilles_Ceramic* (*p*=0.012), and since it provides lower standard error estimates we reported these admixture proportions in the manuscript.

**Table S10**: Modeling *Curacao_Ceramic* with groups from within the two major ancestry clades identified in Table S9.

| A | B | Sources on "Right" | p-value | Anc_A | Anc_B | SE_A | SE_B |
|---|---|---|---|---|---|---|---|
| *Bahamas_Ceramic* | *Venezuela_Ceramic* | - | 0.429 | 0.703 | 0.297 | 0.054 | 0.054 |
| *EasternGreaterAntilles_Ceramic* | *Venezuela_Ceramic* | - | 0.295 | 0.679 | 0.321 | 0.052 | 0.052 |
| *SECoastDR_Ceramic* | *Venezuela_Ceramic* | - | 0.328 | 0.679 | 0.321 | 0.052 | 0.052 |
| *Bahamas_Ceramic* | *Venezuela_Ceramic* | *SECoastDR_Ceramic, EasternGreaterAntilles_Ceramic* | 0.004 | 0.496 | 0.504 | 0.026 | 0.026 |
| *EasternGreaterAntilles_Ceramic* | *Venezuela_Ceramic* | *Bahamas_Ceramic, SECoastDR_Ceramic* | 0.012 | 0.506 | 0.494 | 0.026 | 0.026 |
| *SECoastDR_Ceramic* | *Venezuela_Ceramic* | *Bahamas_Ceramic, EasternGreaterAntilles_Ceramic* | 0.002 | 0.475 | 0.525 | 0.025 | 0.025 |

**Distal modeling with literature samples**

For the distal analysis we used the following previously published individuals/populations (Lindo et al. 2018; Moreno-Mayar et al. 2018; Scheib et al. 2018; Posth et al. 2018) with more than 100,000 SNPs as possible sources for the three major *qpWave* clades:

*Argentina_ArroyoSeco2_7700BP, Argentina_LagunaChica_6800BP, Belize_MayahakCabPek_9300BP, Belize_SakiTzul_7400BP, Brazil_LapaDoSanto_9600BP, Brazil_Laranjal_6700BP, Brazil_Moraes_5800BP, Brazil_Sumidouro_10100BP, Chile_LosRieles_12000BP, Chile_LosRieles_5100BP, Chile_PuntaSantaAna_7300BP, Chile_Ayayema_5100BP, Peru_Cuncaicha_3300BP, Peru_Cuncaicha_4200BP, Peru_Cuncaicha_9000BP, Peru_LaGalgada_4100BP, Peru_Lauricocha_3500BP, Peru_Lauricocha_5800BP, Peru_Lauricocha_8600BP, Peru_SoroMikayaPatjxa_6800BP, USA_CA_Early_SanNicolas*

We first identified which source populations were involved in passing models of the lowest possible rank (i.e., minimum number of sources) with a threshold of p>0.05. We then re-ran the same tests moving the unused of these sources to the "Right" to increase the power of the tests (Table S11).

For *GreaterAntilles_Archaic*, due to the shared ancestry between *Brazil_Moraes_5800BP* and *Brazil_Laranjal_6700BP* their previously working 1-way models failed during model competition, so we removed the lower coverage sample (*Brazil_Moraes_5800BP*) and repeated these analyses. In this new model competition step, the 1-way model for *Brazil_Laranjal_6700BP* (p=0.343) was the only model with p>0.05, suggesting an affinity between *GreaterAntilles_Archaic* and ancient Brazilian populations. However, statistics of the form $f_4$(*Mbuti*, *GreaterAntilles_Archaic*, *Brazil_Laranjal_6700BP*, *Test*), with *Test* as the six other sources, do not support this finding (|Z|<1.451), as expected if it were due to a statistical fluctuation due to limited data for *Brazil_Laranjal_6700BP* (Table S12).

For *Caribbean_Ceramic*, we found no valid models, whereas for *Venezuela_Ceramic* only one 2-way model worked, but with an extremely large standard error of 39.6% that was larger than the smallest ancestry proportion (Table S11). We attempted to add the other sources to the "Right," one at a time, to try to reduce the error. The only improvement was seen when using *Peru_Cuncaicha_4200BP* on the "Right," reducing the error to 20.2% and the ancestry proportion of *Peru_SoroMikayaPatjxa_6800BP* to 41.0% (Table S11).

**Table S11**: Results of *qpAdm* modeling using ancient published data for our 3 major clades.

| Test | A | B | Sources on "Right" | p-value | Anc_A | Anc_B | SE_A | SE_B |
|---|---|---|---|---|---|---|---|---|
| *GreaterAntilles_Archaic* | *Belize_SakiTzul_7400BP* | - | - | 0.220 | 1.000 | - | 0.000 | - |
| *GreaterAntilles_Archaic* | *Brazil_LapaDoSanto_9600BP* | - | - | 0.090 | 1.000 | - | 0.000 | - |
| *GreaterAntilles_Archaic* | *Brazil_Laranjal_6700BP* | - | - | 0.109 | 1.000 | - | 0.000 | - |
| *GreaterAntilles_Archaic* | *Brazil_Moraes_5800BP* | - | - | 0.412 | 1.000 | - | 0.000 | - |
| *GreaterAntilles_Archaic* | *Chile_Ayayema_5100BP* | - | - | 0.404 | 1.000 | - | 0.000 | - |
| *GreaterAntilles_Archaic* | *Chile_PuntaSantaAna_7300BP* | - | - | 0.220 | 1.000 | - | 0.000 | - |
| *GreaterAntilles_Archaic* | *Chile_LosRieles_12000BP* | - | - | 0.055 | 1.000 | - | 0.000 | - |
| *GreaterAntilles_Archaic* | *Belize_SakiTzul_7400BP* | - | *Brazil_LapaDoSanto_9600BP, Brazil_Laranjal_6700BP, Chile_Ayayema_5100BP, Chile_PuntaSantaAna_7300BP, Chile_LosRieles_12000BP* | 0.036 | 1.000 | - | 0.000 | - |
| *GreaterAntilles_Archaic* | *Brazil_LapaDoSanto_9600BP* | - | *Belize_SakiTzul_7400BP, Brazil_Laranjal_6700BP, Chile_Ayayema_5100BP, Chile_PuntaSantaAna_7300BP, Chile_LosRieles_12000BP* | 0.001 | 1.000 | - | 0.000 | - |
| *GreaterAntilles_Archaic* | *Brazil_Laranjal_6700BP* | - | *Brazil_LapaDoSanto_9600BP, Belize_SakiTzul_7400BP, Chile_Ayayema_5100BP, Chile_PuntaSantaAna_7300BP, Chile_LosRieles_12000BP* | 0.343 | 1.000 | - | 0.000 | - |
| *GreaterAntilles_Archaic* | *Chile_Ayayema_5100BP* | - | *Brazil_LapaDoSanto_9600BP, Brazil_Laranjal_6700BP, Belize_SakiTzul_7400BP, Chile_PuntaSantaAna_7300BP, Chile_LosRieles_12000BP* | 4.56E-15 | 1.000 | - | 0.000 | - |
| *GreaterAntilles_Archaic* | *Chile_PuntaSantaAna_7300BP* | - | *Brazil_LapaDoSanto_9600BP, Brazil_Laranjal_6700BP, Chile_Ayayema_5100BP, Belize_SakiTzul_7400BP, Chile_LosRieles_12000BP* | 8.89E-17 | 1.000 | - | 0.000 | - |
| *GreaterAntilles_Archaic* | *Chile_LosRieles_12000BP* | - | *Brazil_LapaDoSanto_9600BP, Brazil_Laranjal_6700BP, Chile_Ayayema_5100BP, Chile_PuntaSantaAna_7300BP, Belize_SakiTzul_7400BP* | 0.007 | 1.000 | - | 0.000 | - |
| *Venezuela_Ceramic* | *Brazil_Moraes_5800BP* | *Peru_SoroMikayaPatjxa_6800BP* | - | 0.217 | 0.670 | 0.330 | 0.396 | 0.396 |
| *Venezuela_Ceramic* | *Brazil_Moraes_5800BP* | *Peru_SoroMikayaPatjxa_6800BP* | *Peru_Cuncaicha_4200BP* | 0.212 | 0.590 | 0.410 | 0.202 | 0.202 |

**Table S12**: Results of statistics of the form $f_4$(*Mbuti*, *GreaterAntilles_Archaic*, *Brazil_Laranjal_6700BP, Test*), with Test as the other 6 possible sources for *GreaterAntilles_Archaic* as in Table S11.

| Pop X | Pop Y | Pop W | Pop Z | f4 | SE | Z | SNPs |
|---|---|---|---|---|---|---|---|
| *Mbuti* | *GreaterAntilles_Archaic* | *Brazil_Laranjal_6700BP* | *Belize_SakiTzul_7400BP* | 0.000013 | 0.00051 | 0.025 | 377425 |
| *Mbuti* | *GreaterAntilles_Archaic* | *Brazil_Laranjal_6700BP* | *Brazil_LapaDoSanto_9600BP* | -0.000433 | 0.000365 | -1.186 | 550383 |
| *Mbuti* | *GreaterAntilles_Archaic* | *Brazil_Laranjal_6700BP* | *Brazil_Moraes_5800BP* | 0.000437 | 0.000551 | 0.793 | 203023 |
| *Mbuti* | *GreaterAntilles_Archaic* | *Brazil_Laranjal_6700BP* | *Chile_Ayayema_5100BP* | -0.000374 | 0.000523 | -0.715 | 565841 |
| *Mbuti* | *GreaterAntilles_Archaic* | *Brazil_Laranjal_6700BP* | *Chile_PuntaSantaAna_7300BP* | -0.000191 | 0.000508 | -0.377 | 444234 |
| *Mbuti* | *GreaterAntilles_Archaic* | *Brazil_Laranjal_6700BP* | *Chile_LosRieles_12000BP* | 0.000825 | 0.000569 | 1.451 | 494119 |

#### Distal modeling using present-day populations

We also ran *qpAdm* for the three major clades using present-day populations representing different language groups and regions. As we had been using these populations as the "Right" of all previous *qpAdm* and *qpWave* analyses, here we tested all 1-, 2-, and 3-way models of the populations from the "Right", moving them to the "Left" as necessary. We did not use Chipewyan in these models as a source due to their distance from ancient Caribbean groups (as shown in previous *f*-statistics results), and also excluded Wayuu and Yukpa due to evidence of admixture. We present the results for both 1- and 2-way models as, although all our three major clades had working 1-way models, the models from the rank above can provide insights into the validity of the lower ranks in poorly modeled situations (Table S13). We also confirmed the results by using the same but unmasked/unadmixed populations from the Illumina dataset (Reich et al. 2012), to the best possible match; only Apalai and Mixtec are missing from this dataset (Table S14).

**Table S13**: Results of 1- and 2-way *qpAdm* modeling using present-day populations from the Human Origins dataset for our 3 major clades.

| Test | A | B | p-value | Anc_A | Anc_B | SE_A | SE_B |
|---|---|---|---|---|---|---|---|
| *Caribbean_Ceramic* | Piapoco | - | 0.519 | 1.000 | - | 0.000 | - |
| *Caribbean_Ceramic* | Piapoco | Quechua | 0.549 | 0.934 | 0.066 | 0.051 | 0.051 |
| *GreaterAntilles_Archaic* | Cabecar | | 0.083 | 1.000 | - | 0.000 | - |
| *GreaterAntilles_Archaic* | Cabecar | Quechua | 0.202 | 0.620 | 0.380 | 0.233 | 0.233 |
| *GreaterAntilles_Archaic* | Karitiana | Quechua | 0.248 | 0.180 | 0.820 | 0.052 | 0.052 |
| *GreaterAntilles_Archaic* | Piapoco | Quechua | 0.186 | 0.292 | 0.708 | 0.071 | 0.071 |
| *GreaterAntilles_Archaic* | Quechua | Apalai | 0.110 | 0.787 | 0.213 | 0.055 | 0.055 |
| *GreaterAntilles_Archaic* | Quechua | Arara | 0.099 | 0.817 | 0.183 | 0.048 | 0.048 |
| *GreaterAntilles_Archaic* | Surui | Cabecar | 0.070 | 0.092 | 0.908 | 0.061 | 0.061 |
| *GreaterAntilles_Archaic* | Surui | Quechua | 0.178 | 0.177 | 0.823 | 0.048 | 0.048 |
| *Venezuela_Ceramic* | Cabecar | - | 0.144 | 1.000 | - | 0.000 | - |
| *Venezuela_Ceramic* | Cabecar | Apalai | 0.165 | 0.911 | 0.089 | 0.064 | 0.064 |
| *Venezuela_Ceramic* | Cabecar | Arara | 0.122 | 0.931 | 0.069 | 0.053 | 0.053 |
| *Venezuela_Ceramic* | Cabecar | Karitiana | 0.095 | 0.933 | 0.067 | 0.060 | 0.060 |
| *Venezuela_Ceramic* | Cabecar | Surui | 0.134 | 0.924 | 0.076 | 0.049 | 0.049 |

*Caribbean_Ceramic* could be modeled as a 1-way model with the Arawak-speaking Piapoco (p=0.519). Upon looking at the working 2-way models, we find support for this result as Piapoco still contributes the great majority of ancestry to *Caribbean_Ceramic* (93.4±5.1%). The results were unchanged when we ran the same tests on the unmasked and unadmixed populations from the Illumina dataset (Reich et al. 2012) (Table S14). These results are consistent with the prevalent theory of an origin of *Caribbean_Ceramic* in Arawak-speaking groups from northern South America, as also shown in our Treemix results. When considered in parallel with the results from outgroup-$f_3$ (Fig. 3a; Supplementary Information section 10; Supplementary Data 8 and 9) it is possible that a more coastal Arawak-speaking population, such as the Palikur, is a more plausible proxy source, although the Piapoco do occupy inland regions of the Orinoco basin.

A single 1-way model with the Chibchan-speaking Cabécar was found for *GreaterAntilles_Archaic*, but in this case most of the working 2-way models did not involve Chibchan-related ancestry, and instead modeled *GreaterAntilles_Archaic* as combinations of present-day populations speaking the Carib, Andean, Tupi, and Arawak languages. The results on the Illumina dataset support this by identifying a valid 1-way model also for the Quechua (Table S14). These results are consistent with the lack of affinity between *GreaterAntilles_Archaic* and populations from a particular language family (Fig. 3a,b; Supplementary Information section 10; Supplementary Data 8 and 9) and with Treemix results (Fig. 3c), all of which point to an early split position relative to other South and Central American populations.

*Venezuela_Ceramic* can also be modeled successfully with Cabécar as the single source, a result that is better supported by our data as 1) all 2-way models require most ancestry to be derived from the Chibchan-speaking Cabécar (91.1±6.4% to 93.3±6.0%), 2) the same result was found on the Illumina dataset populations (**Table S14**), and 3) *Venezuela_Ceramic* clearly shares more drift with Chibchan-speaking groups, as shown by *f*-statistics (Fig. 3a,b; Supplementary Information section 10; Supplementary Data 8 and 9).

**Table S14**: Results of 1-way *qpAdm* models using present-day populations from the Illumina dataset for our 3 major clades.

| Test | A | B | p-value | Anc_A | Anc_B | SE_A | SE_B |
|---|---|---|---|---|---|---|---|
| *Caribbean_Ceramic* | Piapoco | - | 0.249 | 1.000 | - | 0.000 | - |
| *GreaterAntilles_Archaic* | Cabecar | - | 0.586 | 1.000 | - | 0.000 | - |
| *GreaterAntilles_Archaic* | Quechua | - | 0.639 | 1.000 | - | 0.000 | - |
| *Venezuela_Ceramic* | Cabecar | - | 0.450 | 1.000 | - | 0.000 | - |

#### Comparative Analysis of Archaic-related Ancestry in the Ceramic Clades

We tested for the presence of archaic-related ancestry in the final sub-clades with the statistic $f_4$(*China_Tianyuan*, *GreaterAntilles_Archaic*; *Piapoco*, *qpWave_Clade*), which used *China_Tianyuan*, a 40,000-year-old individual from China as an outgroup (Yang et al. 2017), and whole-genome sequencing data from six Piapoco individuals from Bergström et al. (2020), to compare an excess of *GreaterAntilles_Archaic* ancestry in any one of our clades compared to an Arawak-speaking South American population potentially related to *Caribbean_Ceramic* (according to ADMIXTURE (Fig. 2b) and Treemix (Fig. 3c); also see Supplementary Information section 1) and expected to have no admixture with *GreaterAntilles_Archaic*. We used *China_Tianyuan* as an outgroup to avoid artifactually positive results of using a present-day population along with Piapoco. The results suggest a lack of statistically significant proportions of Archaic-related ancestry in most of the Ceramic-associated groups, with clearly identifiable signals in the outlier I16539 (Z = 5.833) and also in the admixed individuals from *Haiti_Ceramic* (Z = 7.736) (Table S15). These results agree with previous tests in identifying no significant Archaic-related ancestry in most of these individuals/groups, except in I16539 and *Haiti_Ceramic*. We caution, however, that if Piapoco has any amount of Archaic-related admixture, such signals would be masked.

**Table S15**: Investigating the overall presence of *GreaterAntilles_Archaic*-related ancestry in the *Caribbean_Ceramic* sub-clades.

| Pop X | Pop Y | Pop W | Pop Z | f4 | SE | Z | SNPs |
|---|---|---|---|---|---|---|---|
| *China_Tianyuan* | *GreaterAntilles_Archaic* | *Piapoco* | *Bahamas_Ceramic* | 0.000035 | 0.000260 | 0.136 | 846136 |
| *China_Tianyuan* | *GreaterAntilles_Archaic* | *Piapoco* | *EasternGreaterAntilles_Ceramic* | 0.000073 | 0.000238 | 0.308 | 847759 |
| *China_Tianyuan* | *GreaterAntilles_Archaic* | *Piapoco* | *SECoastDR_Ceramic* | 0.000111 | 0.000237 | 0.471 | 847851 |
| *China_Tianyuan* | *GreaterAntilles_Archaic* | *Piapoco* | *SECoastDR_Ceramic16539* | 0.003055 | 0.000524 | 5.833 | 631311 |
| *China_Tianyuan* | *GreaterAntilles_Archaic* | *Piapoco* | *Curacao_Ceramic* | -0.000408 | 0.000342 | -1.191 | 820022 |
| *China_Tianyuan* | *GreaterAntilles_Archaic* | *Piapoco* | *Haiti_Ceramic* | 0.003359 | 0.000434 | 7.736 | 825570 |
| *China_Tianyuan* | *GreaterAntilles_Archaic* | *Piapoco* | *Venezuela_Ceramic* | -0.000026 | 0.000312 | -0.085 | 811339 |

We then used the statistic $f_4$(*Mbuti*, *GreaterAntilles_Archaic*, *Sub_Clade1*, *Sub_Clade2*) to directly compare if any specific *Caribbean_Ceramic* sub-clade had more archaic-related ancestry than another, but found no tests producing a Z-score above 2.8, with the exception of I16539 (Table S16).

**Table S16**: Comparing the affinities of *GreaterAntilles_Archaic* to the *Caribbean_Ceramic* sub-clades, after excluding the identified outliers.

| Pop X | Pop Y | Pop W | Pop Z | f4 | SE | Z | SNPs |
|---|---|---|---|---|---|---|---|
| *Mbuti* | *GreaterAntilles_Archaic* | *Bahamas_Ceramic* | *SECoastDR_Ceramic* | 0.000088 | 0.000113 | 0.779 | 1000894 |

| | | | | | | | |
|---|---|---|---|---|---|---|---|
| *Mbuti* | *GreaterAntilles_Archaic* | *Bahamas_Ceramic* | *SECoastDR_Ceramic16539* | 0.002216 | 0.000420 | 5.280 | 687228 |
| *Mbuti* | *GreaterAntilles_Archaic* | *Bahamas_Ceramic* | *EasternGreaterAntilles_Ceramic* | 0.000138 | 0.000120 | 1.153 | 1000233 |
| *Mbuti* | *GreaterAntilles_Archaic* | *SECoastDR_Ceramic* | *SECoastDR_Ceramic16539* | 0.002158 | 0.000120 | 5.288 | 6879007 |
| *Mbuti* | *GreaterAntilles_Archaic* | *SECoastDR_Ceramic* | *EasternGreaterAntilles_Ceramic* | 0.000049 | 0.000408 | 0.572 | 1007247 |
| *Mbuti* | *GreaterAntilles_Archaic* | *SECoastDR_Ceramic16539* | *EasternGreaterAntilles_Ceramic* | -0.002112 | 0.000408 | -5.178 | 687836 |

#### SI9 - Uniparental Haplogroups

##### Mitochondrial (mtDNA) haplogroups

We determined mtDNA haplogroups for the 184 individuals included in this study (described in Methods, with data provided in Supplementary Data 7). Figure S24 illustrates the observed distribution of mtDNA haplogroups across the seven sub-clades in the Caribbean and Venezuela.

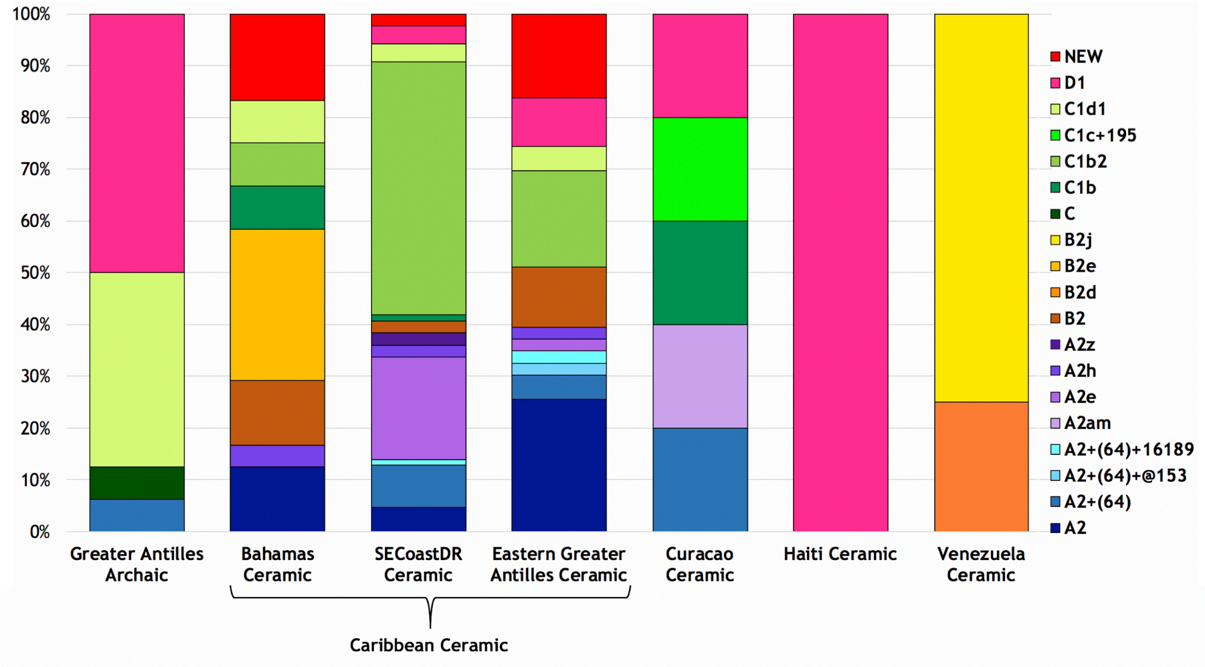

**Figure S24:** Distribution of mitochondrial haplogroups.

We find evidence of all four major pan-American haplogroups (A2, B2, C1, and D1; Schurr and Shery 2004; Achilli et al. 2008) among the samples in our dataset. Lineages from all four of these haplogroups are present in the Caribbean Ceramic-associated samples (the earliest of which dates to ~1800 cal. yr BP), and are not differentially distributed within the *Caribbean_Ceramic* sub-clades defined in this work. Haplogroup B2 is absent in the Caribbean prior to the Ceramic Age and is not identified in any Archaic-associated people who lived during the Ceramic Age, suggesting that this could be somewhat associated with Ceramic-associated peoples and possibly introduced during the Ceramic migrations (more Archaic Age data from the Caribbean will be needed to test this hypothesis). The B2d and B2j lineages are unique in our dataset to Venezuela, and are identified there ~2350 cal. yr BP at the beginning of the Caribbean Ceramic Age; however, we do not find evidence of their introduction into the Caribbean. B2j has been previously reported to have evolved in Venezuela ~3,900 years ago (Gómez-Carballa et al. 2012), consistent with its presence in six out of eight individuals from Las Locas.

As in Nieves-Colón et al. (2020), the most common haplogroup to which the individuals in our dataset belong to is C1b2, with this haplogroup restricted to Ceramic-associated individuals and over one-third of them belonging to it. C1b haplogroups identified in the Greater Antilles have been previously reported to represent links to South America (Vilar et al. 2014), with C1b2 identified in Amazonian populations from interior South America as well as Colombia (Williams et al. 2002; Noguera-Santamaría, et al. 2015). It has been proposed that the C1b2 lineage has been present in Puerto-Rico from the pre-contact era through the present day (Nieves-Colon et al. 2020) where it is currently the most common C1 lineage found (Vilar et al. 2014), and we confirm its ancient presence on the island through its detection in an individual (I13538) from the Ceramic-associated site of Santa Elena. Despite its high frequency, the C1b2 haplogroup is not found in any admixed individuals, including those from Haiti and I16539 who have Archaic-related ancestry, or those from Curaçao who have ancient Venezuela-related ancestry. Admixed individuals from Haiti and I16539 share haplogroups with Ceramic-associated people from other parts of the Caribbean, although we note that C1c+195 and A2am lineages are only found in Curaçao, represented in one individual each.

Ten individuals from our dataset belong to the B2 haplogroup, which was detected in an ancient individual from The Bahamas and is relatively rare in the Caribbean today (Schroeder et al. 2018). While Schroeder and colleagues propose that this haplogroup may also have been relatively rare in the Caribbean in the past, our finding of this haplogroup in ten Ceramic Age individuals suggests that it is more likely that limited pre-Columbian haplotypes persist into the present day (Moreno-Estrada et al. 2013). One observation is that B2e is unique to The Bahamas, where it is found in the Abaco Islands, Long Island, and Andros Island, but not on Crooked Island or Eleuthera.

Our data broadly mirror a present-day pattern in the Caribbean characterized by high frequencies of haplogroups A2 and C1 and lower frequencies of D1 (Mendizabal et al. 2008; Vilar, et al. 2014; Benn-Torres, et al. 2015), with A2 accounting for ~10% of haplogroup calls in our dataset, and lineages of A2 accounting for over 30%, while C1 lineages (C1b, C1b2, C1c+195, and C1d1) accounted for an additional ~40%; however, we do find that D1 is represented at a frequency of ~10%, a non-negligible presence of this haplogroup in the ancient Caribbean. Therefore it is possible that its scarcity in the present-day Caribbean is again a consequence of limited pre-Columbian haplotypes persisting into the present day or persisting in different proportions than in the past.

Out of the 184 samples for which we called mtDNA haplogroups, 13 were ambiguous in their haplogroup assignment (I7974, I7976, I7973, I7972, I14991, I13206, I14990, I14992, I15600, I14879, I13738, I14920, I14921). Therefore, we applied more stringent quality control, restricting to reads with MAPQ ≥ 30 and base quality ≥ 30, and trimming two base pairs to remove deamination artifacts. Further investigation showed that these 13 samples all supported mutations not present in any other samples in our dataset: G5237A, G6429A, T7630C, C8625A, C11509T, C12346T, G13578A, C15381T, A16335G. Of these, three mutations (T7630C C8625A C11509T; shown in Figure S25) were not only exclusive to these 13 samples but were also absent from Phylotree version 17. We identified a single Puerto Rican individual from the 1000 Genomes Project dataset (HG01248) that also had these three mutations. In our ancient samples, these mutations showed good support from multiple reads and were resilient to aggressive filtering. Further, these mutations are not associated with known differences between rCRS or RSRS mitogenomes, suggesting this is not a mitochondrial reference bias artifact.

T7630C

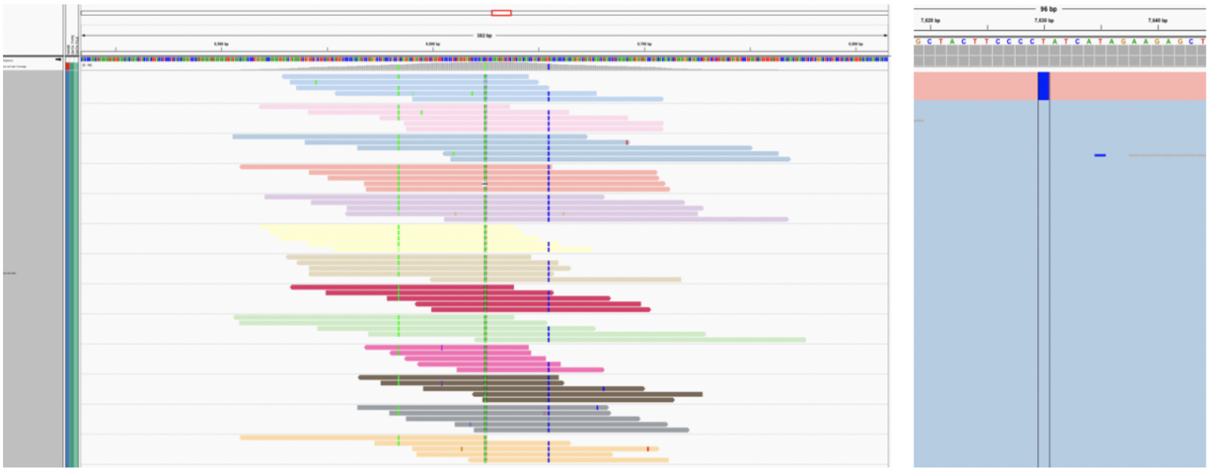

C8625A

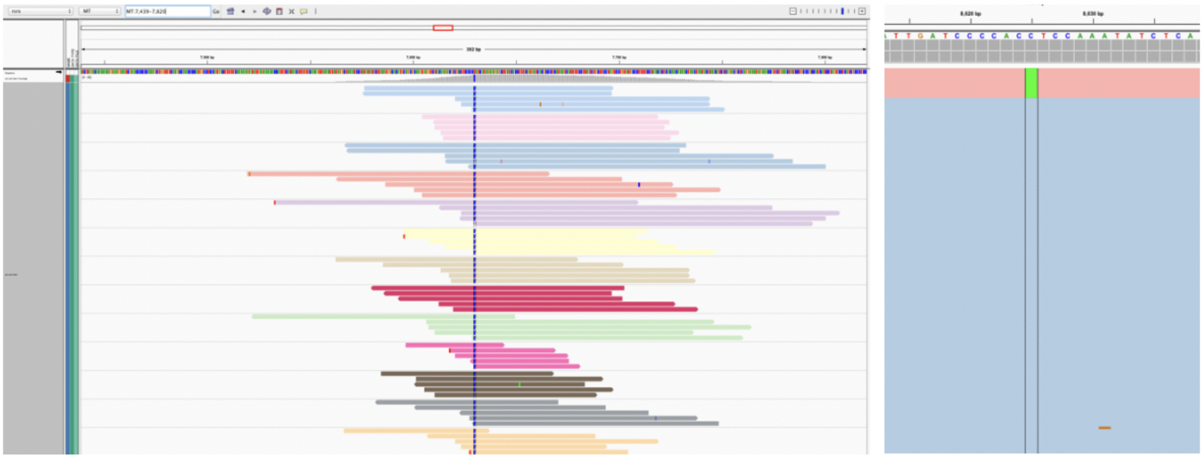

C11509T

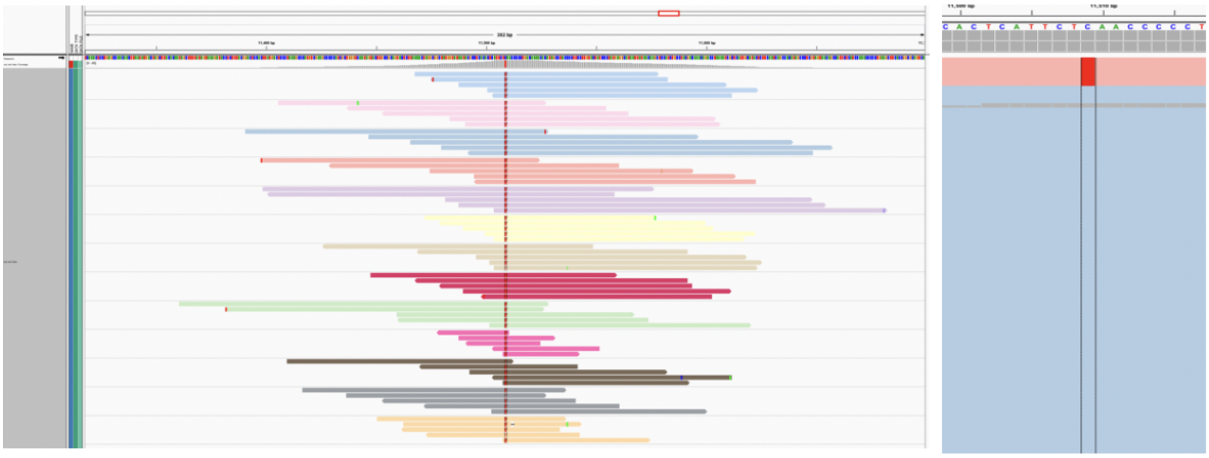

**Figure S25**: For each of the three mutations exclusive to the 13 samples in our dataset and not found in Phylotree, two panels are shown. On the left, a subset of five reads for each of the thirteen samples carrying the novel haplogroup are shown, each colored uniquely. Unambiguous support for the proposed mutations is shown, not associated with indels, proximity to the ends of the reads or other potential confounders. On the right, mitogenomes for all samples in this study are shown (picking alleles based on a majority rule of pass-filter bases); the thirteen samples are shown in pink; other samples are shown in blue.

These 13 individuals were assigned by HaploGrep2 as belonging to haplogroup C, with C1d as the second position call based on the presence of the A16051G mutation diagnostic of this haplogroup. Considering this diagnostic mutation with the additional unique mutations, these data provide evidence of a previously unobserved haplogroup which is a variant of C1d. To date, no scans of other mitogenomes show these mutations.

**Y chromosome haplogroups**

We determined Y chromosome haplogroups for the 99 male individuals included in this study (described in Methods). Figure S26 illustrates the observed distribution of Y chromosome haplogroups across the seven sub-clades in the Caribbean and Venezuela.

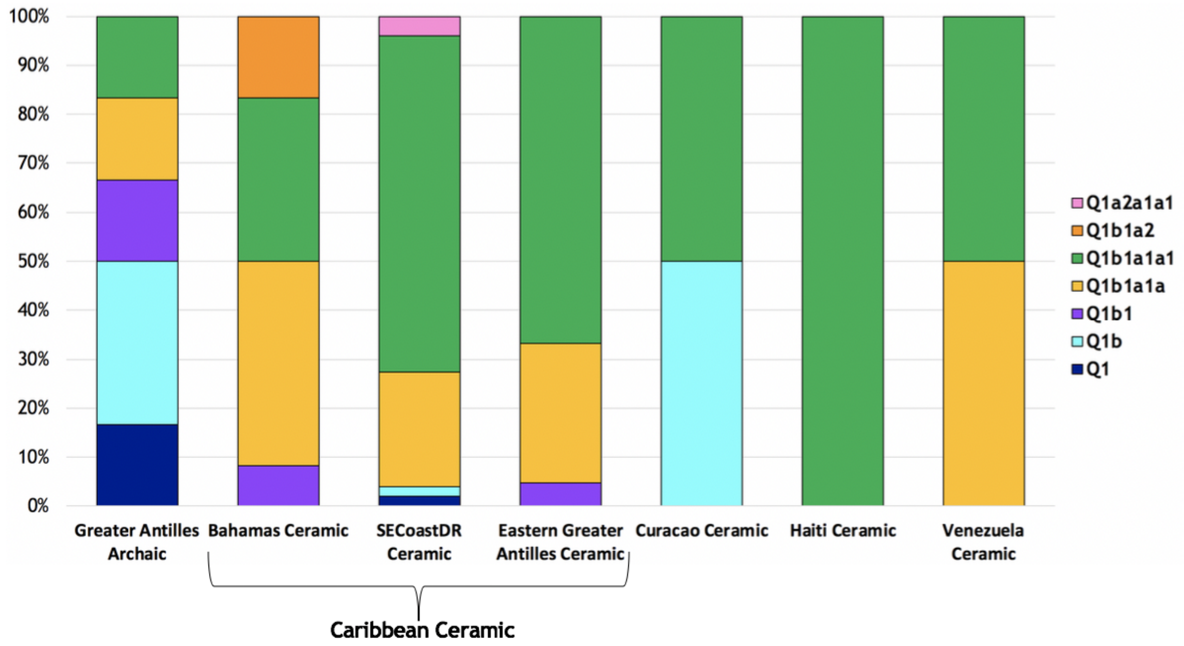

**Figure S26:** Distribution of Y chromosome haplogroups across seven sub-clades in the Caribbean and Venezuela.

Out of 99 Y chromosome haplogroups called, 88 individuals from all seven sub-clades were assigned to the Q-M3 lineage, with 28 male individuals belonging to Q1b1a1a and 60 to Q1b1a1a1. Q-M3 is strongly and deeply linked to Indigenous peoples of the Americas (Underhill et al. 1996; Kivisild 2017), where it is still found today (Bortolini et al. 2003; Battaglia et al. 2013; Benn Torres et al. 2015), showing the persistence of Y chromosome ancestry despite significant population turnover that resulted from European contact.

Two individuals, both from the Bahamas, belong to Q1b1a2, with one individual sharing the same derived mutation as found in Anzick (Q-M971; Rasmussen et al. 2014; Kivisild 2017). This suggests that not only did this Anzick-related lineage reach South America, but that it also spread into the Caribbean as well.

#### SI10 - *f*-statistics and Relatedness to Modern Language Groups

The Ceramic migration into the Caribbean has been attributed to populations related to modern Indigenous groups from the Macro-Arawakan language family (Supplementary Information section 1). We investigated the affinities of the ancient individuals from this study to present-day populations representing the different language groups from Central and South America. Based on the results of outgroup-$f_3$ analysis for each of the three major clades defined in SI11 we identified the top 10 populations with whom they shared the most drift, and their corresponding language group (Supplementary Data 8). Then we performed symmetry tests on the populations from these language groups using statistics of the form $f_4$(*Mbuti*, *Test*; *LanguageGroupA_Pop*, *LanguageGroupB_Pop*), with *Test* as the *qpWave* clades, and *LanguageGroupA/B_Pop* as all pairwise combinations of the populations identified in outgroup-$f_3$, and shown in Extended Data Table 4. The individual test results are presented in Supplementary Data 9.

The modern Indigenous populations with which ancient Caribbeans shared the highest drift belong to seven main language groups: Cariban, Chibchan, Chocoan, Macro-Arawakan, Guajiboan, Mataco-Guaicuru and Tupian. For *GreaterAntilles_Archaic* we found no specific affinities to any language group (Fig. 3).*Venezuela_Ceramic* showed clear shared ancestry with Chibchan-speaking populations. Finally, *Caribbean_Ceramic* was found to have a significantly closer relationship to Macro-Arawakan/Guajiboan/Tupian/Cariban-speakers than the other groups. The admixture events identified in other analyses help explain the positions of *Curacao_Ceramic* and *Haiti_Ceramic*, which were found in intermediate positions relative to the other ancient populations from the same geographical region.

#### SI11- *qpGraph*

We investigated where *GreaterAntilles_Archaic* would fit in the skeleton tree of ancient American populations previously published by Posth and colleagues (2018) using *qpGraph*. We found that *GreaterAntilles_Archaic* fit in several locations of the tree without increasing the maximum individual f4-statistics above |Z|=3.5, suggesting a lack of power of currently available data to provide a good fit for this ancestry (**Figure S27**). This was particularly true for any position after *Canada_Lucier_4800BP-500BP* (*ASO_SG*) that followed through to the branch that included ancient Brazilian and Argentinian populations.

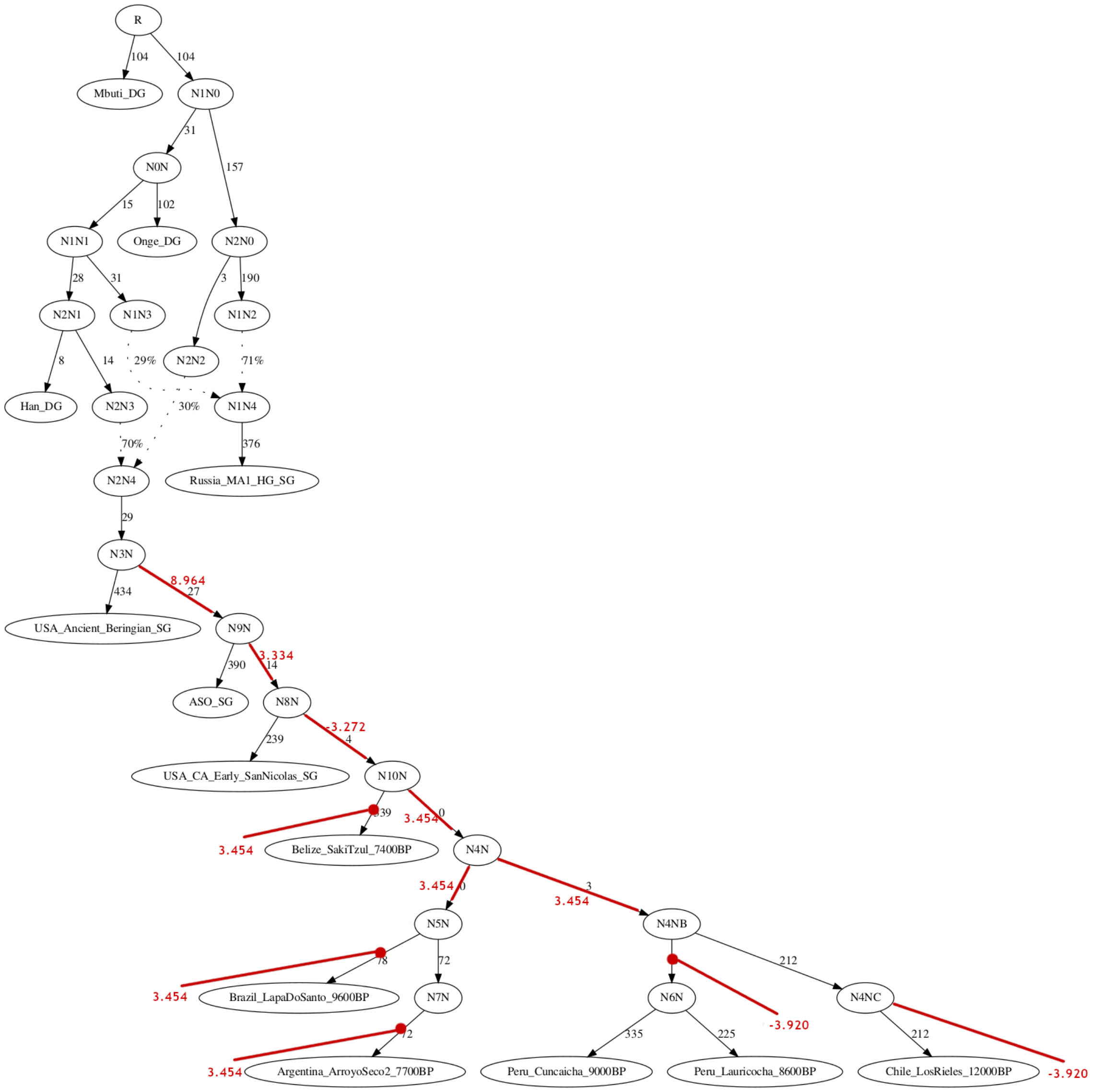

**Figure S27**: Admixture graph modelling of *GreaterAntilles_Archaic* with previously sequenced ancient individuals/populations. Lines in red represent locations where *GreaterAntilles_Archaic* was inserted, with the corresponding graph's worst Z-score shown in red.

#### SI12 - Ability to detect a Carib migration into the Caribbean: simulations and qpWave

Using facial morphology, Ross and colleagues (2020) found support for a movement of Carib peoples originating in the northwestern Amazon basin into the Greater Antilles around 800 CE. We therefore looked for evidence of a specific Carib-related genetic contribution to Ceramic Age individuals. Since our *Venezuela_Ceramic* individuals from Las Locas inhabit the centre of the region suggested by Ross and colleagues (2020) as the source of Carib-related ancestry, and predate the hypothetical arrival of ancestry from this region in the Greater Antilles, we also tested them as a possible proxy. Using admixture simulations (as described in the Methods section of the manuscript) we found that *qpWave* is able to detect Carib-related and *Venezuela_Ceramic*-related ancestries when they are present at levels that range between 2 and 8% in the genome of a *Caribbean_Ceramic* individual (Figure S28). We therefore cannot completely rule out a possible migration of individuals with these ancestries into the Greater Antilles at around 800 CE; however, we can attest that if this migration did occur, the contribution of this specific ancestry to peoples of the Greater Antilles analyzed here was equal to or less than 8%. An additional caveat to this analysis is the requirement that the present-day Cariban-speaking Arara or the ancient individuals in *Venezuela_Ceramic* closely represent the ancient Carib ancestry.

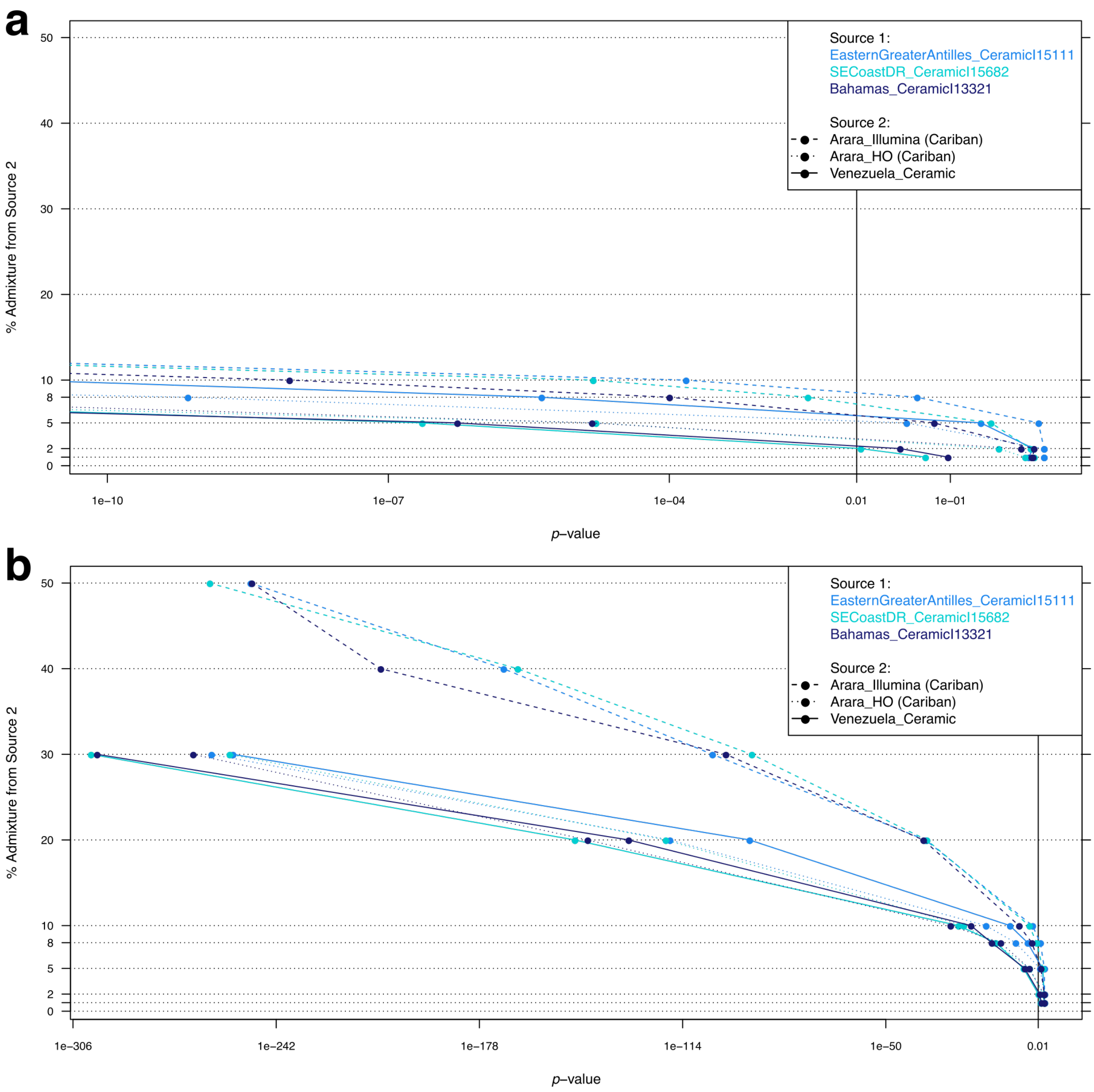

**Figure S28:** Minimum admixture detection threshold for Carib- and *Venezuela_Ceramic*-related ancestries (Source 2) using *qpWave* and simulated admixed individuals with primarily *Caribbean_Ceramic* ancestry (Source 1). The *qpWave* *p*-value threshold used was 0.01. Zoomed (**a**) and complete (**b**) plots shown. Logarithmic scale used for the X axis.

#### SI13- DATES

For two individuals from the site of Diale 1 in Haiti and the outlier individual I16539 from La Caleta, Dominican Republic (*SECoastDR_Ceramic_16539*), we used *GreaterAntilles_Archaic* (composed of all individuals from the Cuban site of Canímar Abajo and I10126), *Caribbean_Ceramic* (composed of *Bahamas_Ceramic*, *EasternGreaterAntilles_Ceramic* and *SECoastDR_Ceramic*) as a reference pair and estimated that admixture occurred an average of ~11±5 generations (or ~310±130 years) before the individuals from Haiti lived, and 39±14 generations (or ~1,100±400 years) before the individual I16539 lived, assuming a generation time of 28 years (Moorjani et al. 2016) (Table S17**)**. Using an average context date of 950 years BP for *Haiti_Ceramic*, this places admixture as occurring ~1,400-1,100 years BP, while using an average context date of 850 years BP for *SECoastDR_Ceramic_16539*, admixture is estimated to have occurred ~2,300-1,550 years BP. We note that the dates obtained with this method are based on a model of a single pulse of admixture and thus reflect an intermediate value if the true history includes multiple waves or continuous admixture. We also note that the Z-scores for the differences of the date estimates from zero were >2.0 but <2.8 for these two tests.

We were unable to obtain feasible dates for the Ceramic-associated individuals from Curacao using any combination of the main clades defined in this study as reference populations.

**Table S17**: Admixture date estimation with DATES for *Haiti_Ceramic* and *SECoastDR_Ceramic_16539*.

| Admixed Pop | Reference Pop 1 | Reference Pop 2 | Estimate (generations) | Std Err (generations) | Z-score |
|---|---|---|---|---|---|
| *Haiti_Ceramic* | *GreaterAntilles_Archaic* | *Caribbean_Ceramic* | *10.87* | 4.848 | 2.24 |
| *SECoastDR_Ceramic_I16539* | *GreaterAntilles_Archaic* | *Caribbean_Ceramic* | *38.74* | 13.99 | 2.77 |

#### SI14 – Relatedness of ancient individuals to present-day admixed Caribbean populations

After computing relative allele sharing between present-day admixed Caribbean populations and the ancient individuals, we performed an empirical power analysis for the present-day Cubans by leveraging our ancient data to study whether we would expect to be able to detect differentiation in indigenous ancestry between Archaic-related and ceramic-related. As a baseline, we had observed a significance level of Z = 0.5 for the statistic $f_4$(*European*, *Cuban*; *Cuba_Archaic*, *Caribbean_Ceramic*). We then merged the Archaic-associated individual I10126 into the present-day Cuban test population and recomputed the statistic, obtaining a significantly lower value (Z = -2.4). Thus, we can conclude that we have sufficient power to detect one full genome from the *GreaterAntilles_Archaic* clade when added to the second position in the $f_4$-statistic, even though I10126 has incomplete sequencing coverage and is plausibly more differentiated from *Cuba_Archaic* than persisting Archaic-related ancestry in Cuba would be.

We estimated 2.9-4.2% Indigenous ancestry in the present-day Cuban individuals using qpAdm (proxy sources consisting of *Caribbean_Ceramic*, Europeans [1000 Genomes CEU], and West Africans [1000 Genomes YRI], and reference groups drawn either from 1000 Genomes [PEL, PJL, JPT, and MSL] or Mallick et al. (2016) [Karitiana, Mixe, Yakut, Ulchi, Papuan, Mursi, and Mbuti]) (Extended Data Table 2). For 55 individuals, this equates to a total of 1.6-2.3 genomes' worth of Indigenous ancestry. Given that we observe sufficient power with a single genome under conservative assumption, we expect that if the actual Indigenous ancestry were (all) from an Archaic-related source, we would be able to detect it as well.

We then performed an analogous test with Ceramic-related ancestry in which we combined one Ceramic-associated individual (I14992 from Los Muertos) with the present-day Cubans. In this case, the $f_4$-statistic increases slightly to Z = 1.3. The difference from the baseline in this case is not statistically significant, and in fact, even if we extrapolate from a single genome to the high end of the indigenous ancestry estimate, we would not necessarily expect to have power to detect a greater affinity of the present-day Cubans to *Caribbean_Ceramic*. Thus, while we can rule entirely Archaic-related ancestry contributing to the indigenous ancestry in present-day Cubans, we cannot rule out entirely ceramic-related ancestry (or a mixture of both).

We note that all three provinces and three of the five municipalities (Yara, Banes, and Puerto Padre) in Cuba in which we identify weakly ceramic-related ancestry (Supplementary Data 11; Extended Data Table 3) are located in the eastern portion of the island. Indigenous ancestry is more prevalent in this region of Cuba compared to the central and western parts of the country (Fortes-Lima et al. 2018), allowing our statistics more power. While these results are not novel (having been previously discussed in Fortes-Lima et al. 2018), they are presented here to highlight the presence of multiple different ancestry types to present-day Caribbean people. It is also possible that ancient groups in eastern Cuba, in contrast to our sampled individuals from Canímar Abajo from the northwest, harbored Ceramic-related ancestry prior to European colonization.

#### SI15 - Ethics Statement

*Permissions for this study*

Permissions to carry out ancient DNA analysis of the human skeletal remains in this study were documented through authorization letters and Memorandums of Understanding (MOU) signed by a custodian who assumed responsibility for the skeletal remains collected from a specific geographic region or site. The authorization letter established permission to perform ancient DNA analysis (as well as radiocarbon dating, isotopic analysis, and other bioarchaeological analysis) on the ancient skeletal material and stated that 1) all skeletal samples were recovered during past archaeological excavations and archived for scientific analysis; 2) the institution on behalf of which the custodian signed provided explicit permission to perform the analysis carried out in this study; and 3) the institution on behalf of which the custodian signed was supportive of publishing results of this study in a scientific journal.

- **Bahamas:** Authorization letter signed by Michael Pateman (Director, Turks and Caicos National Museum); letter of permission for the export of samples from The Bahamas and for ancient DNA, isotopic, and 14C dating analyses signed by Keith L. Tinker (Director, The National Museum of The Bahamas);
- **Cuba:** Authorization letter signed by Ercilio Vento Canosa (Matanzas City Historian, paleopathologist for the Provincial Monument Commission, and Dean of the Chair of Paleopathology at the University of Medical Sciences in Matanzas);
- **Curaçao**: Authorization letter signed by Dmitri Cloose (Director, National Archaeological Anthropological Memory Management, NAAM);
- **Dominican Republic**: Authorization letter signed by Arq. Christian Martinez Villanueva (Director General. Museo del Hombre Dominicano);
- **Venezuela**: Authorization letter signed by Prof. Carlos García Sívoli (on behalf of the Instituto de Investigaciones Bioantropológicas y Arqueológicas de la Universidad de Los Andes);
- **Haiti**: Permission for the generation of full genome data using human skeletal remains was provided by the Peabody Museum of Natural History at Yale University;
- **Puerto Rico**: Permission for the generation of full genome data using human skeletal remains was provided by the Peabody Museum of Natural History at Yale University; we note that NAGPRA compliance is not relevant for the ancient individuals from Puerto Rico included in our study as the law has no jurisdiction and no federally recognized tribe has claimed affiliation.

*Collaboration and outreach*

In this study, we test hypotheses surrounding the ancient peoples of the Caribbean islands using paleogenomic data as one line of evidence that is useful for studying the past. Anthropologists, archaeologists, and other scholars from each geographic region included in this study played an integral role in the interpretation of data and formulation of results and were included as co-authors.

We also consulted multiple other Caribbean-based stakeholders throughout the course of the project to solicit critical feedback and local perspectives. A major goal was to ensure that local narratives were considered during the process of contextualizing these data, and thus the involvement and feedback of local scholars was crucial.

The inferences that we make as part of this project are intended to provide new information about the genetic ancestry of the people of the Caribbean prior to the colonization by Europeans in the late 15th century and to explore the genetic contribution from the ancient peoples of this region to the genomes of present-day Caribbean people. We note that our study of genetic ancestry of people in this region should not be conflated with individual and/or community perceptions of identity; genetic ancestry should not be conflated with perceptions of identity, which cannot be defined by genetics alone. Genetic data are one form of knowledge that contributes to understanding the past, and oral traditions and other forms of Indigenous knowledge can coexist with scientific data.

*Protection of archaeological sites*

Latitude and longitude coordinates for the archaeological sites from which the skeletal remains of individuals examined were excavated are reported in Supplementary Data 1. The Society for American Archaeology's Ethical Principle #6 states that, "An interest in preserving and protecting in situ archaeological sites must be taken into account when publishing and distributing information about their nature and location"; thus, we have provided latitude and longitude coordinates to two decimal degree digits in an attempt to balance protecting archaeological site integrity against publishing the geographic data needed to test genetic correlations with distance measures and meet open science replicability standards.

de Boyrie Moya, E. Cinco años de arqueología dominicana. (Anales de la Universidad de Santo Domingo, 1960).

de las Casas, B. Apologética historia sumaria. (Alianza, 1992).

de Moya E., B. & Herrera Fritot, R. Primera Exposicion de arte indigena autoctono. vol. 5 (Universidad de Santo Domingo, 1948).

Dobyns, H. F. Their Number Become Thinned: Native American Population Dynamics in Eastern North America. (University of Tennessee Press, 1983).

Drew, R. A survey of Pre-Columbian human remains from Puerto Rico. in Proceedings of the Twentieth International Congress for Caribbean Archaeology (eds. Tavares Maria, G. & Arevalo, M. G.) 522–534 (Museo del Hombre Dominicano and Fundacion Garcia Arevalo, 2005).

Drew, R. From Saladoid to Taíno: Human Behavior from Human Remains in the Greater Antilles. Bulletin of the Peabody Museum of Natural History 50, 167–185 (2009).

Drusini, A., Businaro, F. & Calderòn, F. L. Skeletal biology of the Taino: a preliminary report. International Journal of Anthropology 2, 247–254 (1987).

Dumas León, D. Historia y osteología de los restos óseos humanos exhumados entre 1984 y 1987 en Canímar Abajo, Matanzas, Cuba. (University of Havana, 2009).

Febles, J. Estudio tipológico y tecnológico del material de piedra tallada del sitio arqueológico Canímar I, Matanzas, Cuba. (Editorial Academia, 1982).

Federmann, N. Indianische Historia: Ein Schöne Kurtzweilige Historia Niclaus Federmanns des Jüngern von Ulmerster Raise. (Hagenau, 1557).

Fernandes, D. M. et al. The spread of steppe and Iranian-related ancestry in the islands of the western Mediterranean. Nat Ecol Evol 4, 334–345 (2020).

Figueredo, A. Manioc dethroned and maize triumphant: Interpretations on the etnohistory and archaeology of the Bahamas. Journal of Caribbean Archaeology 15, 120–134 (2015).

Fitzpatrick, S. & Rick, T. Local marine reservoir effect corrections for the Caribbean. in 26th Internation Congress for Caribbean Archaeology (2015).

Fitzpatrick, S. M. & Keegan, W. F. Human impacts and adaptations in the Caribbean Islands: an historical ecology approach. Earth and Environmental Science Transactions of the Royal Society of Edinburgh 98, 29–45 (2007).

Fitzpatrick, S. M., Kappers, M. & Giovas, C. The Southward Route Hypothesis: Examining Carriacou's chronological position in Antillean prehistory. Island Shores, Distant Pasts: Archaeological and Biological Approaches to the Pre-Columbian Settlement of the Caribbean 163–176 (2010).

Fortes-Lima, C. et al. Exploring Cuba's population structure and demographic history using genome-wide data. Sci. Rep. 8, 11422 (2018).

Frankham, R. Effective population size/adult population size ratios in wildlife: a review. Genet. Res. 89, 491–503 (2007).

Fu, Q. et al. A revised timescale for human evolution based on ancient mitochondrial genomes. Curr. Biol. 23, 553–559 (2013).

Fu, Q. et al. An early modern human from Romania with a recent Neanderthal ancestor. Nature 524, 216–219 (2015).

Fu, Q. et al. The genetic history of Ice Age Europe. Nature 534, 200–205 (2016).

Gansauge, M.-T. et al. Single-stranded DNA library preparation from highly degraded DNA using T4 DNA ligase. Nucleic Acids Res. 45, e79 (2017).

Garcia Arévalo, M. A. La Arqueologia Indo-Hispama en santo Domingo. in Unidad y vauedad: Ensayos Anthropologicos en Homenaje a José M. Cruxent (eds. Wagner, E. & Zucchi, A.) 77–127 (Istituto Venezolano de Investigaciones Cientificas, 1978).

Garcia Arévalo, M. A. Transculturation in Contact Period and Contemporary Hispaniola. in Archaeological and Historical Perspectives on the Spanish Borderlands East (ed. Thomas, D. H.) 269–280 (Smithsonian Institution Press, 1990).

Gillin, J. P. Tribes of the Guianas and the Left Amazon Tributaries. Handbook of South American Indians 3, 799–860 (1948).

Goggin, J. M. The Spanish Olive Jar an Introductory Study. (Yale University, 1960).

Gómez-Carballa, A. et al. A melting pot of multicontinental mtDNA lineages in admixed Venezuelans. Am. J. Phys. Anthropol. 147, 78–87 (2012).

Granberry, J. & Vescelius, G. Languages of the Pre-Columbian Antilles. (University of Alabama Press, 2004).

Granberry, J. A Survey of Bahamian Archaeology. (University of Florida, 1955).

Granberry, J. Lucayan Toponyms. Journal of the Bahamas Historical Society 13, 3–12 (1991).

Granberry, J. The Gordon Hill Site, Crooked Island, Bahamas. Journal of the Virgin Islands Archaeological Society 6, 32–44 (1978).

Greenberg, J. H. Language in the Americas. (Stanford University Press, 1987).

Haak, W. et al. Massive migration from the steppe was a source for Indo-European languages in Europe. Nature 522, 207–211 (2015).

Harney, É. et al. Ancient DNA from Chalcolithic Israel reveals the role of population mixture in cultural transformation. Nat. Commun. 9, 3336 (2018).

Hastings, A. K., Krigbaum, J., Steadman, D. W. & Albury, N. A. Domination by Reptiles in a Terrestrial Food Web of the Bahamas Prior to Human Occupation. Journal of Herpetology 48, 380–388 (2014).

Haviser, J. 1987 Amerindian Cultural Geography on Curaçao. (Leiden University, 1987).

Heckenberger, M. & Neves, E. G. Amazonian Archaeology. Annual Review of Anthropology 38,

251–266 (2009).

Heckenberger, M. J. Rethinking the Arawakan diaspora. in Comparative Arawakan histories: Rethinking language family and culture area in Amazonia (eds. Hill, J. D. & Santos-Granero, F.) 99–122 (University of Illinois, 2002).

Henige, D. On the Contact Population of Hispaniola: History as Higher Mathematics. Hispanic American Historical Review 58, 217–237 (1978).

Hernández de Alba, G. Tribes of Northwestern Venezuela. in The Circum-Caribbean Tribes (ed. Steward, J. H.) vol. 22 469–474 (Smithsonian Institution, 1948).

Herrera Fritot, R. & Youmans, C. L. La Caleta: Joya Arqueológica Antillana. (La Habana, 1946).

Hoffman, C. A. Petroglyphs on Crooked Island, Bahamas. Proceedings of the Fourth International Congress for the Study of the Pre-Columbian Cultures of the Lesser Antilles, St. Lucia Archaeological and Historical Society 9–12 (1973).

Hofman, C. L. & Reid, B. A. The Saladoid. in Encyclopedia of Caribbean archaeology (eds. Reid, B. & Gilmore, G.) 300–303 (University of Florida Press, 2014).

Hofman, C. L. & van Duijvenbode, A. Communities in Contact: Essays in Archaeology, Ethnohistory & Ethnography of the Amerindian Circum-Caribbean. (Sidestone Press, 2011).

Hofman, C. L., Ulloa Hung, J. & Jacobs, L. Juntando las piezas del rompecabezas: Dándole sentido a la cronología cerámica del este de la República Dominicana. El Caribe Arqueológico 10, 104–115 (2007).

Hoogland, M. L. P. & Hofman, C. L. Expansion of the Taino cacicazgos towards the Lesser Antilles. Journal de la Société des Américanistes 85, 93–113 (1999).

Hung, J. U. & Rojas, R. V. Cerámica temprana en el centro del oriente de Cuba. (Videograph, 2002).

Keegan, W. F. & DeNiro, M. J. Stable Carbon- and Nitrogen-Isotope Ratios of Bone Collagen Used to Study Coral-Reef and Terrestrial Components of Prehistoric Bahamian Diet. American Antiquity 53, 320–336 (1988).

Keegan, W. F. & Hofman, C. L. The Caribbean before Columbus. (Oxford University Press, 2017).

Keegan, W. F. Archaic Influences in the Origins and Development of Taino Societies. Caribbean Journal of Science 42, 1–10 (2006).

Keegan, W. F. Islands of Chaos. in The Late Ceramic Age Societies in the Eastern Caribbean (eds. Delpuech, A. & Hofman, C. L.) vol. 1273 33–44 (British Archaeological Reports, 2004).

Keegan, W. F. Lucayan Cave Burials from The Bahamas. Journal of New World Archaeology 5, 57–65 (1982).

Keegan, W. F. Modeling dispersal in the prehistoric West Indies. World Archaeology 26, 400–420 (1995).

Keegan, W. F. Situating Jamaica. in Early Settlers of the Insular Caribbean. Dearchaizing the Archaic (eds. Hofman, C. L. & Antczak, A. T.) 191–200 (Sidestone Press Academics, 2019).

Keegan, W. F. The Development and Extinction of Lucayan Society. Terrae Incognitae 24, 43–53 (1992).

Keegan, W. F., Hofman, C. L. & Ramos, R. R. The Oxford Handbook of Caribbean Archaeology. (Oxford University Press, 2013).

Keegan, W. F., Pateman, M. P. & White, M. Garden Cave Skeleton Recovery Project, Eleuthera, The Bahamas. (Report submitted to the Antiquities, Monuments and Museum Corporation, 2018).

Keegan, W. The People Who Discovered Columbus. (University Press of Florida, 1992).

Keehnen Floris, W. M. Treating 'Trifles': the Indigenous Adoption of European Material Goods in Early Colonial Hispaniola (1492–1550). in Material Encounters and Indigenous Transformations in the Early Colonial Americas (eds. Hofman, C. L. & Keehnen, F. W. M.) (Brill, 2019).

Kelleher, J., Etheridge, A. M. & McVean, G. Efficient Coalescent Simulation and Genealogical Analysis for Large Sample Sizes. PLoS Comput. Biol. 12, e1004842 (2016).

Kivisild, T. The study of human Y chromosome variation through ancient DNA. Human Genetics 136, 529–546 (2017).

Kjellmark, E. Late Holocene climate change and human disturbance on Andros Island, Bahamas. Journal of Paleolimnology 15, (1996).

Korlević, P. et al. Reducing microbial and human contamination in DNA extractions from ancient bones and teeth. Biotechniques 59, 87–93 (2015).

Korneliussen, T. S., Albrechtsen, A. & Nielsen, R. ANGSD: Analysis of Next Generation Sequencing Data. BMC Bioinformatics 15, 356 (2014).

Kraan, C. et al. New Archaeological Research on Curaçao. The Journal of Island and Coastal Archaeology 12, 138–144 (2016).

Krieger, H. W. Aboriginal Indian pottery of the Dominican Republic. 1–165 (Smithsonian Institution United States National Museum Bulletin 156, 1931).

Lalueza-Fox, C., Calderón, F. L., Calafell, F., Morera, B. & Bertranpetit, J. MtDNA from extinct Tainos and the peopling of the Caribbean. Ann. Hum. Genet. 65, 137–151 (2001).

Lalueza-Fox, C., Gilbert, M. T. P., Martínez-Fuentes, A. J., Calafell, F. & Bertranpetit, J. Mitochondrial DNA from pre-Columbian Ciboneys from Cuba and the prehistoric colonization of the Caribbean. Am. J. Phys. Anthropol. 121, 97–108 (2003).

Lathrap, D. W. Archäologische Untersuchungen am Mittlern Amazonas. (Dietrich Reimer, 1970).

Lazaridis, I. et al. Ancient human genomes suggest three ancestral populations for present-day Europeans. Nature 513, 409–413 (2014).

Lazaridis, I. et al. Genomic insights into the origin of farming in the ancient Near East. Nature 536, 419–424 (2016).

LeFebvre, M. J., deFrance, S. D., Kamenov, G. D., Keegan, W. F. & Krigbaum, J. The zooarchaeology and isotopic ecology of the Bahamian hutia (Geocapromys ingrahami):

Evidence for pre-Columbian anthropogenic management. PLoS One 14, e0220284 (2019).

Li, H. & Durbin, R. Fast and accurate long-read alignment with Burrows-Wheeler transform. Bioinformatics 26, 589–595 (2010).

Lindo, J. et al. The genetic prehistory of the Andean highlands 7000 years BP though European contact. Sci Adv 4, eaau4921 (2018).

Lleonart, R., Riego, E., Suárez, R. R., Ruiz, R. T. & de la Fuente, J. Analyses of DNA from ancient bones of a pre-Columbian Cuban woman and a child. Genetics and Molecular Biology 22, 285–289 (1999).

Lohse, J. C., Culleton, B. J., Black, S. L. & Kennett, D. J. A Precise Chronology of Middle to Late Holocene Bison Exploitation in the Far Southern Great Plains. Journal of Texas Archeology and History 1, 94–126 (2014).

Lovén, S. E. Origins of the Tainan Culture, West Indies. (Elanders Bokfryckeri Akfiebolag, 1935).

Lowie, R. H. The tropical forest cultures: an introduction. in Handbook of South American Indians, Vol. 3: The tropical forest tribes (ed. Steward, J. H.) 1–56 (Smithsonian Institution Bureau of American Ethnology, 1948a).

Lowie, R. H. Property among the tropical forest and marginal tribes. in Handbook of South American Indians, Vol. 5: The comparative ethnology of South American Indians (ed. Steward, J. H.) 351–367 (Smithsonian Institution Bureau of American Ethnology, 1948b).

Luna Calderon, F. Antropología Y Paleopatologia de los Pobladores del Soco. in Tenth International Congress for the Study of Pre-Columbian Cultures of the Lesser Antilles (1985).

Luna Calderón, F. Informe preliminar del cemeterio indigena de 'El Atajadizo', Republica Dominicana. Boletin del Museo del Hombre Dominicano 7, 67–95 (1976).

Luna Calderón, F. Preliminary report on the Indian cemetary 'El Atajadizo,' Dominican Republic. in Proceedings of the Sixth International Congress for the Study of Pre-Columbian Cultures of the Lesser Antilles 295–303 (1976).

Mallick, S. et al. The Simons Genome Diversity Project: 300 genomes from 142 diverse populations. Nature 538, 201–206 (2016).

Martínez-Cruzado, J. C. The history of Amerindian mitochondrial DNA lineages in Puerto Rico. in Island shores, distant pasts: archaeological and biological approaches to the Pre-Columbian settlement of the Caribbean (eds. Fitzpatrick, S. M. & Ross, A. H.) 92–100 (University Press of Florida, 2010).

Martínez-Cruzado, J. The DNA Evidence for the Human Colonization and Spread Across the Americas. Oxford Handbooks Online (2013) doi:10.1093/oxfordhb/9780195392302.013.0135.

Martínez-López, J. G., Arredondo Antúnez, C., Rodríguez Suárez, R. & Díaz-Franco, S. Aproximación Tafonómica en los depósitos humanos del sitio arqueológico Canímar Abajo, Matanzas, Cuba. Arqueologia Iberoamericana 4, 5-21 (2009).

Martínez-López, J. G., Díaz-Franco, S. & Morales, Y. D. Valoraciones tafonómicas sobre el sitio arqueológico Canímar Abajo, Matanzas, Cuba. II Seminario Inter- nacional de Arqueología (La Habana, Cuba) (2007).

Mathieson, I. et al. Genome-wide patterns of selection in 230 ancient Eurasians. Nature 528, 499–503 (2015).

McClure, S. B., Puchol, O. G. & Culleton, B. J. Ams Dating of Human Bone from Cova De La Pastora: New Evidence of Ritual Continuity in the Prehistory of Eastern Spain. Radiocarbon 52, 25–32 (2010).

Meggers, B. J. Biologist Philosopher: A Study of the Life and Writings of Alfred Russel Wallace . Wilma George. American Anthropologist 67, (1965).

Mendisco, F. et al. Where are the Caribs? Ancient DNA from ceramic period human remains in the Lesser Antilles. Philos. Trans. R. Soc. Lond. B Biol. Sci. 370, 20130388 (2015).

Mendizabal, I. et al. Genetic origin, admixture, and asymmetry in maternal and paternal human lineages in Cuba. BMC Evol. Biol. 8, 213 (2008).

Metraux, A. Weapons. in Handbook of South American Indians, Vol. 5: The comparative ethnology of South American Indians (ed. Steward, J. H.) 229–261 (Smithsonian Institution Bureau of American Ethnology, 1948a).

Metraux, A. Boys' initiation rites. in Handbook of South American Indians, Vol. 5: The comparative ethnology of South American Indians (ed. Steward, J. H.) 275–382 (Smithsonian Institution Bureau of American Ethnology, 1948b).

Metraux, A. Religion and shamanism. in Handbook of South American Indians, Vol. 5: The comparative ethnology of South American Indian (ed. Steward, J. H.) 559–599. (Smithsonian Institution Bureau of American Ethnology, 1948c).

Moorjani, P., Gao, Z. & Przeworski, M. Human Germline Mutation and the Erratic Evolutionary Clock. PLoS Biol. 14, e2000744 (2016).

Morales Valdes, D. Caracterización osteológica de aborígenes adultos exhumados en Canímar Abajo, durante la campaña de excavación del 2006. (University of Havana, 2009).

Morbán Laucer, F. Ritos Funerarios Acción del Fuego y Medio Ambiente en las Osamentas Precolombinas. vol. 2 (Academia de Ciencias de la República Dominicana, 1979).

Moreno-Estrada, A. et al. Reconstructing the population genetic history of the Caribbean. PLoS Genet. 9, e1003925 (2013).

Moreno-Mayar, J. V. et al. Early human dispersals within the Americas. Science 362, (2018).

Mylroie, J. E. & Mylroie, J. R. Flank Margin Caves in Carbonate Islands and the Effects of Sea Level. Treatise on Geomorphology 351–362 (2013).

Nakatsuka, N. et al. A Paleogenomic Reconstruction of the Deep Population History of the Andes. Cell (2020) doi:10.1016/j.cell.2020.04.015.

Napolitano, M. F. et al. Reevaluating human colonization of the Caribbean using chronometric hygiene and Bayesian modeling. Sci Adv 5, eaar7806 (2019).

Narasimhan, V. M. et al. The formation of human populations in South and Central Asia. Science 365, (2019).

Nieves-Colón, M. A. et al. Ancient DNA reconstructs the genetic legacies of pre-contact Puerto

Rico communities. Molecular Biology and Evolution 37, 611–626 (2020).

Noguera-Santamaría, M. C. et al. Mitochondrial DNA analysis suggests a Chibchan migration into Colombia. Universitas Scientiarum 20, 261 (2015).

Olalde, I. et al. The genomic history of the Iberian Peninsula over the past 8000 years. Science 363, 1230–1234 (2019).

Oliver, J. R. The archaeological, linguistic and ethnohistorical evidence for the expansion of Arawakan into northwestern Venezuela and northeastern Colombia. (University of Illinois at Urbana-Champaign, 1989).

Oliver, J. R. The Taino cosmos. in Indigenous People of the Caribbean (ed. Wilson, S. M.) vol. 27 140–153 (University Press of Florida, 1997).

Olsen, H. B. Investigaciones Arqueológicas Punta Macao. Manuscript on file. (Museo Del Hombre Dominicano, 2004).

Ortega, E. & Fondeur, C. Estudio de la cerámica del período indo-hispano de la antigua Concepción de La Vega. (Taller, 1978).

Ortega, E. Compendio general arqueológico de Santo Domingo. (Academia de Ciencias de la República Dominicana, 2005).

Pagán-Jiménez, J. R. Human–Plant Dynamics in the Precolonial Antilles. in The Oxford handbook of Caribbean archaeology (eds. Keegan, W. F., Hofman, C. L. & Rodríguez Ramos, R.) 391–406 (Oxford University Press, 2013).

Pagán-Jiménez, J. R., Rodríguez-Ramos, R., Reid, B. A., van den Bel, M. & Hofman, C. L. Early dispersals of maize and other food plants into the Southern Caribbean and Northeastern South America. Quaternary Science Reviews 123, 231–246 (2015).

Palmer, R. Hydrology and speleogenesis beneath Andros Island. Cave Science 13, 7–12 (1986).

Palmer, R. The Blue Holes of the Bahamas. Cave Science 13, 3–6 (1986).

Pantel, A. G. Precolumbian Flaked Stone Assemblages in the West Indies. (University of Tennessee, 1998).

Passariello, I. et al. Characterization of Different Chemical Procedures for 14C Dating of Buried, Cremated, and Modern Bone Samples at Circe. Radiocarbon 54, 867–877 (2012).

Passes, A. The Anthropology of Love and Anger. (Routledge, 2002).

Pateman, M. P. & Keegan, W. F. Lucayan Burials in the Bahama Archipelago. in 84th annual meeting of the Society for American Archaeology (2019).

Pateman, M. P. Reconstructing Lucayan Mortuary Practices Through Skeletal Analysis. Journal of the Bahamas Historical Society 5–10 (2007).

Patterson, N. et al. Ancient admixture in human history. Genetics 192, 1065–1093 (2012).

Patterson, N., Price, A. L. & Reich, D. Population structure and eigenanalysis. PLoS Genet. 2, e190 (2006).

Persons, B. Pottery, People, and Place: Examining the Emergence of Political Authority in Late Ceramic Age Cuba. (University of Alabama, 2013).

Pickrell, J. K. & Pritchard, J. K. Inference of population splits and mixtures from genome-wide allele frequency data. PLoS Genet. 8, e1002967 (2012).

Pinhasi, R. et al. Optimal Ancient DNA Yields from the Inner Ear Part of the Human Petrous Bone. PLoS One 10, e0129102 (2015).

Pinhasi, R., Fernandes, D. M., Sirak, K. & Cheronet, O. Isolating the human cochlea to generate bone powder for ancient DNA analysis. Nat. Protoc. 14, 1194–1205 (2019).

Posth, C. et al. Reconstructing the Deep Population History of Central and South America. Cell 175, 1185–1197.e22 (2018).

Raghavan, M. et al. Genomic evidence for the Pleistocene and recent population history of Native Americans. Science 349, aab3884 (2015).

Rainey, F. G. Diary beginning January 22, 1934: Upon arrival in Port-Au-Prince. vol. 50 (Manuscript on file at Yale Peabody Museum, 1934).

Rainey, F. G. Excavations in the Ft. Liberté Region, Haiti. (Yale University, 1941).

Rainey, F. G. Porto Rican Archaeology. Scientific Survey of Porto Rico and the Virgin Islands. N. Y. Academy of Sciences 18, (1940).

Ralph, P. & Coop, G. The geography of recent genetic ancestry across Europe. PLoS Biol. 11, e1001555 (2013).

Randolph, L. An Ethnobotanical Investigation of Andros, Island, Bahamas. (Miami University, 1994).

Rasmussen, M. et al. The genome of a Late Pleistocene human from a Clovis burial site in western Montana. Nature 506, 225–229 (2014).

Reich, D. et al. Reconstructing Native American population history. Nature 488, 370–374 (2012).

Reimer, P. J. et al. IntCal13 and Marine13 Radiocarbon Age Calibration Curves 0–50,000 Years cal BP. Radiocarbon 55, 1869–1887 (2013).

Rímoli, R. O. Restos alimenticios en Atajadizo y Guayabal. in Arqueología de Yuma, República Dominicana. Santo Domingo (eds. Veloz Maggiolo, M., Vargas, I., Sanoja, M. & Calderón, F.) 341–344 (Taller, 1976).

Ringbauer, H., Coop, G. & Barton, N. H. Inferring Recent Demography from Isolation by Distance of Long Shared Sequence Blocks. Genetics 205, 1335–1351 (2017).

Ringbauer, H., Novembre, J. & Steinrucken, M. Detecting runs of homozygosity from low-coverage ancient DNA. bioRxiv.org (2020).

Rivera-Collazo, I. Palaeoecology and human occupation during the Mid-Holocene in Puerto Rico: the case of Angostura. in Communities in Contact: Essays in archaeology, ethnohistory & ethnography of the Amerindian circum-Caribbean (eds. Hofman, C. & van Duijvenbode, A.) 407–420 (Sidestone Press, 2011).

Rodriguez Ramos, R. From the Guanahatabey to the Archaic of Puerto Rico: The Nonevident Evidence. Ethnohistory 55, 393–415 (2008).

Rodríguez Ramos, R. Isthmo-Antillean Engagements. in Oxford Handbook of Caribbean Archaeology (eds. Keegan, W. F., Hofman, C. L. & Rodríguez Ramos, R.) 155–170 (Oxford University Press, 2013).

Rodríguez Ramos, R. Rethinking Puerto Rican Precolonial History. (University of Alabama Press, 2010).

Rodríguez Ramos, R. The Crab-Shell Dichotomy Revisited: The Lithics Speak Out. in Ancient Borinquen: Archaeology and Ethnohistory of Native Puerto Rico (ed. Siegel, P. E.) 1–54 (University of Alabama Press, 2005).

Rodríguez Suárez, R. et al. 5000 años de ocupación prehispánica en Canímar Abajo, Matanzas, Cuba. in (Memorias del congreso Antropología, 2006).

Rodríguez Suárez, R., Buhay, B. & Roksandic, M. New dates from Canimar Abajo. Biological and cultural landscapes of indigenous people in pre- and post-contact Cuba. (2010).

Rohland, N., Glocke, I., Aximu-Petri, A. & Meyer, M. Extraction of highly degraded DNA from ancient bones, teeth and sediments for high-throughput sequencing. Nat. Protoc. 13, 2447–2461 (2018).

Rohland, N., Harney, E., Mallick, S., Nordenfelt, S. & Reich, D. Partial uracil-DNA-glycosylase treatment for screening of ancient DNA. Philos. Trans. R. Soc. Lond. B Biol. Sci. 370, 20130624 (2015).

Roksandic, I. & Roksandic, M. Peopling of the Caribbean. 199–223 (Kerns Verlag, 2018).

Roksandic, M. et al. Radiocarbon and Stratigraphic Chronology of Canímar Abajo, Matanzas, Cuba. Radiocarbon 57, 755–763 (2015).

Roksandic, M., Alarie, K., Rodríguez Suárez, R., Huebner, E. & Roksandic, I. Not of African Descent: Dental Modification among Indigenous Caribbean People from Canímar Abajo, Cuba. PLoS One 11, e0153536 (2016).

Ross, A. H., Keegan, W. F., Pateman, M. P. & Young, C. B. Faces Divulge the Origins of Caribbean Prehistoric Inhabitants. Sci. Rep. 10, 147 (2020).

Rothschild, B. M., Calderon, F. L., Coppa, A. & Rothschild, C. First European exposure to syphilis: the Dominican Republic at the time of Columbian contact. Clin. Infect. Dis. 31, 936–941 (2000).

Rouse, I. & Cruxent, J. M. Venezuelan Archaeology. (New Haven : Yale University Press, 1963).

Rouse, I. Introduction to prehistory: A systematic approach. (McGraw-Hill, 1972).

Rouse, I. Migrations in Prehistory: Inferring Population Movement from Cultural Remains. (Yale University Press, 1986).

Rouse, I. Porto Rican Prehistory: Introduction: Excavations in the West and North. Scientific Survey of Porto Rico and the Virgin Islands. vol. XVIII-Part 3 (New York Academy of Sciences, 1952a).

Rouse, I. Porto Rican Prehistory: Excavations in the Interior, South and East: Chronological Implications. vol. XVIII-Part 4 (New York Academy of Sciences, 1952b).

Rouse, I. The Classification of Artifacts in Archaeology. American Antiquity 25, 313–323 (1960).

Rouse, I. The Tainos: Rise & Decline of the People who Greeted Columbus. (Yale University Press, 1992).

Russell, N., Cook, G. T., Ascough, P. L. & Marian Scott, E. A period of calm in Scottish seas: A comprehensive study of ΔR values for the northern British Isles coast and the consequent implications for archaeology and oceanography. Quaternary Geochronology 30, 34–41 (2015).

Salzano, F. M., Hutz, M. H., Salamoni, S. P., Rohr, P. & Callegari-Jacques, S. M. Genetic Support for Proposed Patterns of Relationship among Lowland South American Languages. Current Anthropology 46, S121–S128 (2005).

Sanoja, M. & Vargas, I. Las Sociedades Formativas del Noroeste de Venezuela y el Orinoco Medio. International Journal of South American Archaelogy 1, 14–23 (2007).

Sanoja, M. & Vargas, I. Proyecto de Arqueología del occidente de Venezuela. Primer Informe General. Revista de Economía y Ciencias Sociales 2, 25–60 (1967).

Schaffer, W. C. A Reappraisal of Prehistoric Human Skeletal Remains from The Bahamas Housed at the Yale Peabody Museum of Natural History. Journal of Caribbean Archaeology 15, 134–156 (2015).

Scheib, C. L. et al. Ancient human parallel lineages within North America contributed to a coastal expansion. Science 360, 1024–1027 (2018).

Schroeder, H. et al. Origins and genetic legacies of the Caribbean Taino. Proc. Natl. Acad. Sci. U. S. A. 115, 2341–2346 (2018).

Schulting, R. J. et al. Seven centuries of adaptation to a challenging island environment: AMS 14C dating and isotopic analyses of pre-Columbian human remains from the Bahamian archipelago reveal dietary trends. (In preparation).

Schurr, T. G. & Sherry, S. T. Mitochondrial DNA and Y chromosome diversity and the peopling of the Americas: evolutionary and demographic evidence. Am. J. Hum. Biol. 16, 420–439 (2004).

Sealey, N. E. Bahamian Landscapes: An Introduction to the Geography of the Bahamas. (Media Publishing Ltd., 1985).

Sears, W. H. & Sullivan, S. O. Bahamas Prehistory. American Antiquity 43, 3–25 (1978).

Siegel, P. E. et al. Paleoenvironmental evidence for first human colonization of the eastern Caribbean. Quaternary Science Reviews 129, 275–295 (2015).

Sirak, K. et al. Human auditory ossicles as an alternative optimal source of ancient DNA. Genome Res. 30, 427–436 (2020).

Skoglund, P. et al. Genetic evidence for two founding populations of the Americas. Nature 525, 104–108 (2015).

Skoglund, P. et al. Genomic insights into the peopling of the Southwest Pacific. Nature 538,

510–513 (2016).

Smith, D. G. et al. Straddling the subsistence divide. in The archaeology of Caribbean and Circum-Caribbean farmers (6000 BC-AD 1500) (ed. Reid, B. A.) 98–112 (Routledge, 2018).

Steadman, D. W. et al. Exceptionally well preserved late Quaternary plant and vertebrate fossils from a blue hole on Abaco, The Bahamas. Proc. Natl. Acad. Sci. U. S. A. 104, 19897–19902 (2007).

Steadman, D. W., Albury, N. A., Mead, J. I., Angel Soto-Centeno, J. & Franklin, J. Holocene vertebrates from a dry cave on Eleuthera Island, Commonwealth of The Bahamas. The Holocene 28, 806–813 (2017).

Steward, J. H. & Faron, L. C. Native Peoples of South America. (McGraw-Hill, 1959).

Stokes, A. V. A biogeographic survey of prehistoric human diet in the West Indies using stable isotopes. (University of Florida, 1998).

Tacoma, J. Skeletal remains from de Savaan Region, Curaçao. in Proceedings of the XI International Congress for Caribbean Archaeology 217–223 (1990).

Tavarez María, G. Informe de Antropología Física del Cementerio de Macao. (Museo del Hombre Dominicano, 2004).

Taylor, R. E. Six Decades of Radiocarbon Dating in New World Archaeology. Radiocarbon 51, 173–212 (2009).

Toro-Labrador, G., Wever, O. R. & Martinez-Cruzado, J. C. Mitochondrial DNA Analysis in Aruba: Strong Maternal Ancestry of Closely Related Amerindians and Implications for the Peopling of Northwestern Venezuela. Caribbean Journal of Science 39, (2003).

Underhill, P. A., Jin, L., Zemans, R., Oefner, P. J. & Cavalli-Sforza, L. L. A pre-Columbian Y chromosome-specific transition and its implications for human evolutionary history. Proceedings of the National Academy of Sciences 93, 196–200 (1996).

Valcárcel Rojas, R. Banes Precolombino. La ocupacion agricultora. (Ediciones Holguin, 2002).

Veloz Maggiolo, M. & Ortega, E. J. Excavaciones en Macao, República Dominicana. Boletín del Museo del Hombre Dominicano 2, 157 (1972).

Veloz Maggiolo, M. & Ortega., E. J. Punta Canay el origen de la agricultura en la isla de Santo Domingo. in Ponencias del Primer Seminario de Ar-queología del Caribe (eds. Veloz Maggiolo, M. & Caba Fuentes, A.) 5–11 (Museo Arqueológico Regional Altos de Chavón, 1996).

Veloz Maggiolo, M. Arqueología prehistórica de Santo Domingo. (McGraw-Hill Far Eastern Publishers Ltd, 1972).

Veloz Maggiolo, M. La isla de Santo Domingo antes de Colón. (Banco Central de la Republica Dominicana, 1993).

Veloz Maggiolo, M. Medioambiente y adaptación humana en la prehistoria de Santo Domingo. Tomo 2. vol. 30 (Editora de la Universidad Autonoma de santo Domingo, Coleccion Historia y Sociedad, 1977).

Veloz Maggiolo, M., Ortega, E. & Sanoja, M. Preliminary Report on Archaeological

Investigations at El Atajadizo, Dominican Republic. in Proceedings of the Sixth International Congress for the Study of Pre-Columbian Cultures of the Lesser Antilles 283–294 (1976).

Veloz Maggiolo, M., Ortega, E., Peña, P. P., Rimoli, R. & Calderón, F. El cementerio del La 'Union' Provincia de Puerto Plata. Buletin del Museo del Hombre Dominicano 2, 130–156 (1972).

Veloz Maggiolo, M., Ortega, E., Rimoli, R. & Calderón, F. Estudio comparative y preliminar de dos cementerios neo-indios:La Cucama y La Union, Repubblica Dominicana. Buletin del Museo del Hombre Dominicano 3, 11–47 (1973).

Veloz Maggiolo, M., Vargas, I., Sanoja, M. & Luna Calderón, F. Arqueología de Yuma, República Dominicana. (Taller, 1976).

Versteeg, A. H. & Schinkel, K. The Archaeology of St. Eustatius: The Golden Rock Site. (Publication of the St. Eustatius Historical Foundation 2 and Publication of the Foundation for Scientific Research in the Caribbean Region 131, 1992).

Vilar, M. G. et al. Genetic diversity in Puerto Rico and its implications for the peopling of the Island and the West Indies. Am. J. Phys. Anthropol. 155, 352–368 (2014).

Walker, R. S., Wichmann, S., Mailund, T. & Atkisson, C. J. Cultural phylogenetics of the Tupi language family in lowland South America. PLoS One 7, e35025 (2012).

Welsch, R. L., Terrell, J. & Nadolski, J. A. Language and Culture on the North Coast of New Guinea. American Anthropologist 94, 568–600 (1992).

Willey, G. R. An Introduction to American Archaeology Volume 2: South America. (Prentice Hall, 1971).

Williams, S. R., Chagnon, N. A. & Spielman, R. S. Nuclear and mitochondrial genetic variation in the Yanomamö: a test case for ancient DNA studies of prehistoric populations. Am. J. Phys. Anthropol. 117, 246–259 (2002).

Wilson, S. M. The Archaeology of the Caribbean. (Cambridge University Press, 2007).

Wilson, S. M., Iceland, H. B. & Hester, T. R. Preceramic Connections between Yucatan and the Caribbean. Latin American Antiquity 9, 342–352 (1998).

Yang, M. A. et al. 40,000-Year-Old Individual from Asia Provides Insight into Early Population Structure in Eurasia. Curr. Biol. 27, 3202–3208.e9 (2017).

Zucchi, A. A new model of the northern Arawakan expansion. in Comparative Arawakan histories: Rethinking language family and culture area in Amazonia (eds. Hill, J. D. & Santos-Granero, F.) vol. 43 199–222 (University of Illinois, 2002).

Zucchi, A. La Serie Meillacoide y sus Relaciones con la Cuenca del Orinoco. (Puerto Rico, 1985).
